# The biochemical impact of extracting an embedded adenylate kinase domain using circular permutation

**DOI:** 10.1101/2023.10.25.564053

**Authors:** Tom Coleman, John Shin, Jonathan J. Silberg, Yousif Shamoo, Joshua T. Atkinson

## Abstract

Adenylate kinases (AKs) are phosphotransferases that are frequently employed as models to investigate protein structure-function relationships. Prior studies have shown that AK homologs of different stabilities retain cellular activity in cells following circular permutation that split the AMP binding domain into fragments coded at different ends of the primary structure, such that this domain was no longer embedded as a continuous polypeptide within the core domain. Herein, we show mesophilic and thermophilic AKs having this topological restructuring retain activity and substrate-binding characteristics of the parental AK. While permutation decreased the activity of both AK homologs at physiological temperatures, the catalytic activity of the thermophilic AK increased upon permutation when assayed >30°C below the melting temperature of the native AK. The thermostabilities of the permuted AKs were uniformly lower than native AKs, and they exhibited multi-phasic unfolding transitions, unlike the native AKs, which presented cooperative thermal unfolding. In addition, proteolytic digestion revealed that permutation destabilized each AK, and mass spectrometry suggested that the new termini within the AMP binding domain were responsible for the increased proteolysis sensitivity. These findings illustrate how changes in contact order can be used to tune enzyme activity and alter folding dynamics in multidomain enzymes.

**Graphical abstract:** 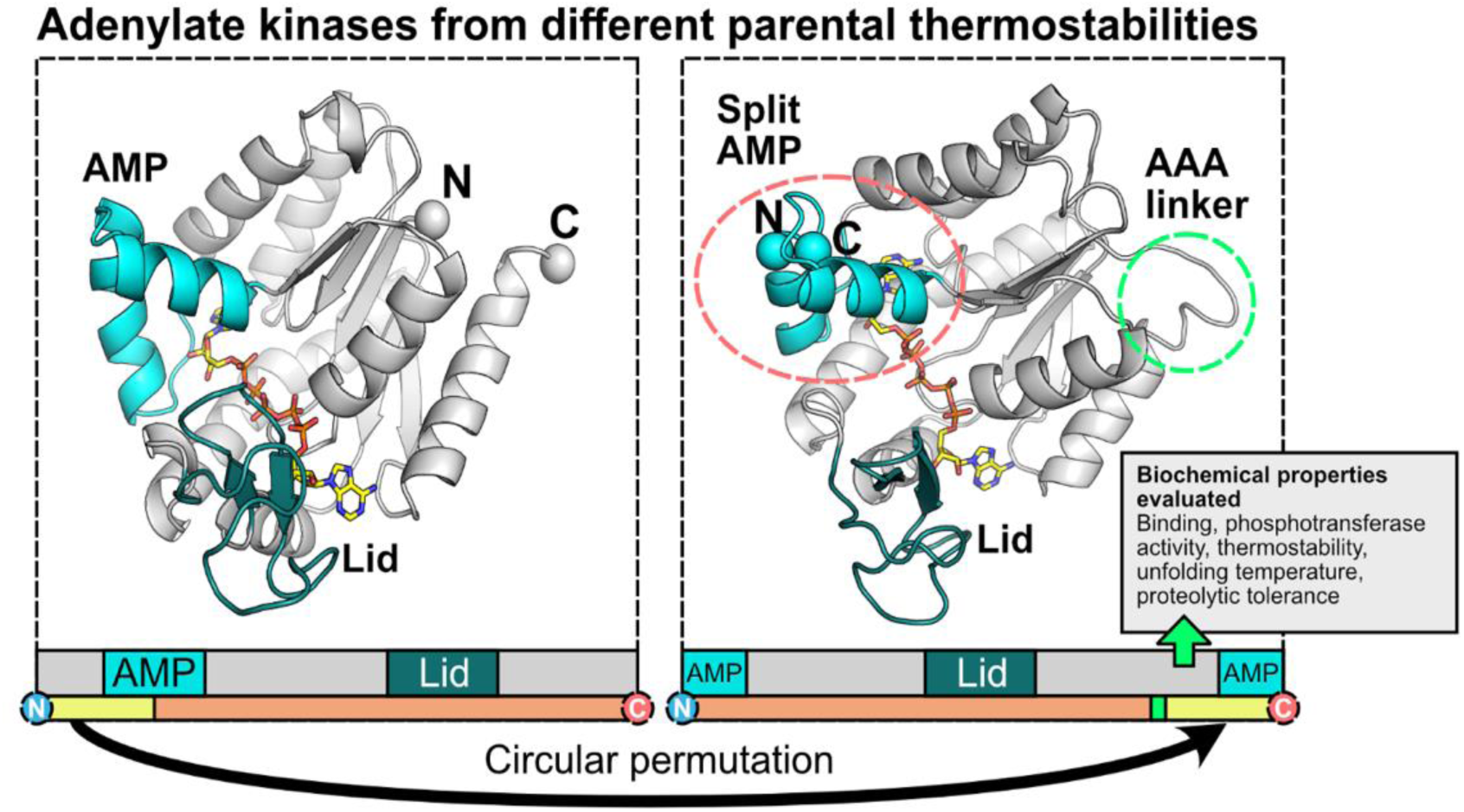

## INTRODUCTION

During evolution, proteins can gain new functions through mechanisms such as gene fusion and recombination, resulting in enzymes with multiple domains. Following events such as gene shuffling, multiple domains may become embedded within each other.^1–3^ Adenylate kinases (AKs) are good models for proteins with embedded domains, as computational studies have implicated a role for their inserted domains facilitating cooperative folding.^4^ AKs are Mg^2+^-dependent phosphotransferases that modulate adenylate homeostasis by catalyzing the reversible reaction AMP + ATP ⇄ 2 ADP.^5–7^ AKs are comprised of core, lid, and AMP binding domains. The lid and AMP binding domains are embedded at different locations within a discontinuous core domain, which contains the catalytic residues that carry out the phosphoryl transfer reaction. During catalysis, the lid and AMP binding domains close over the site of phosphoryl transfer, with lid opening representing the enzyme’s rate limiting step.^8^ Initially studied due to the crucial cellular role across all domains of life, AKs have since become an established model system for the study of protein structure, function and dynamics, through extensive structural, mutational, and computational efforts.^4, 5, 9–18^

AKs are found in organisms that grow optimally across a range of environments, including organisms that live in the human gut, soils, sediments, and oceans to those that persist in extremes of temperature, acidity, salinity, and radiation.^19–21^ These latter extremotolerant organisms, termed extremophiles, produce enzymes with specialized properties to support fitness in extreme environments. AK homologs from extremophiles have been compared with those from mesophiles to study protein adaptation as well as the relationship between protein dynamics and function.^19, 22–25^ The manipulation of AK genes in microorganisms has been additionally employed to demonstrate fitness adaptation to a variety of selection pressures.^26–28^ Structural and kinetic studies performed on AKs from thermophilic, mesophilic, and psychrophilic organisms – *Geobacillus stearothermophilus* (*Gs*AK), *Bacillus subtilis* (*Bs*AK), *Escherichia coli* (*Ec*AK), and *Bacillus globisporus* (*Bg*AK), respectively – have revealed similar overall structures.^13, 14, 29^ However, their thermostabilities and optimum temperatures for activity are markedly different, highlighting the significance of protein adaptation within a specific environmental context.^29–31^

A computational study has suggested that the embedded domains in AKs support cooperative folding and stabilize structure, even though the individual domains vary in stability.^4^ One way to directly probe this idea is to alter AK contact order using circular permutation, creating variants where one of the domains is no longer embedded. With this topological restructuring, the original N- and C- termini of the protein are fused and new termini are created elsewhere in the protein, such as in the middle of an embedded domain.^32,33^ There is evidence that proteins have evolved in nature through this type of topological mutation.^33^ This dramatic topological restructuring is thought to arise when a gene is duplicated, fused into a tandem repeat, and then subjected to fission.^32, 34^ In some proteins, this type of structural rearrangement may confer an evolutionary advantage. In the lab, the use of circular permutation to generate new proteins can permit the study of the biochemical impact of altering contact order and at times improve catalytic activity.^35, 36^ Combinatorial experiments have also shown that functional tolerance to permutation correlates with thermostability.^37^ When a trio of AK homologs were targeted for random circular permutation, all three homologs retained function at 40°C when permutation created new N- and C-termini within the AMP binding domain.^37^ While this study quantified the cellular fitness of homologous permuted AKs, the biochemical and biophysical properties of these proteins were not directly characterized.

To better understand the biochemical advantages of embedding a domain, we characterized a pair of AKs where the embedded AMP-binding domain was extracted via circular permutation. We expressed, purified, and characterized circularly permutated Aks from a mesophile (*Bs*AK) and thermophile (*Gs*AK), where the protein sequence began translation at residue 45, designated herein as cp45 (Figure 1). Because residue 45 resides in the center of the AMP binding domain,^37^ backbone fission at this location has the potential to alter substrate binding, phosphotransfer kinetics, and protein thermostability. Both the mesophilic and thermophilic cp45 retained phosphotransferase activities, albeit with decreased binding affinities and thermostabilities. Further, these enzymes presented an increased sensitivity to proteolytic digestion which suggested changes to protein folding dynamics, the extent of protein folding, and surface accessibility of proteolytic sites. Notably, the new cp45 enzymes were still catalytically active. While the AKs presented decreased catalytic activities at physiological temperatures, the thermophilic AK (*Gs*AK) presented increased steady-state activity when assayed at temperatures consistent with growth of mesophiles rather than the ancestral thermophile.

**Figure 1.**
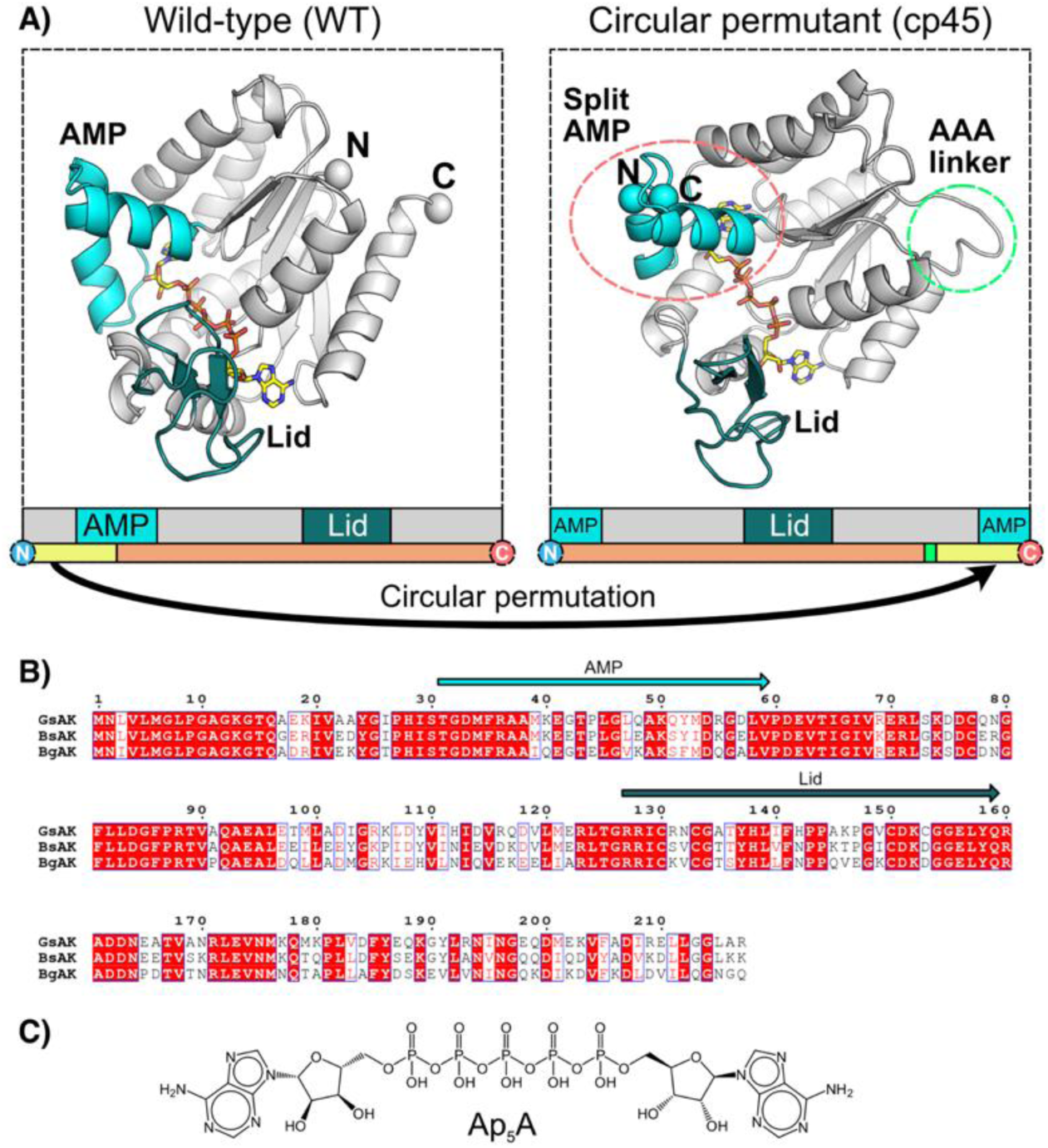
Effect of changing contact order on AK structure. (**A**) Schematic of AK before and after permutation that yields cp45. The core, (grey), AMP binding (cyan), and lid (dark blue) domains are colored differently. (*left*) The *Gs*AK (PDB: 1ZIP) structure is compared with (*right*) a model of cp45 structure generated using ColabFold.^38^ The N- and C-termini are shown as spheres. The location of the AAA linker is highlighted (green circle). (**B**) Sequence alignment of the mesophilic (*Bs*AK), thermophilic (*Gs*AK), and psychrophilic (*Bg*AK) discussed herein. The AMP and lid domains are indicated with arrows. (**C**) The structure of Ap_5_A, an AK inhibitor.

## MATERIALS & METHODS

### Molecular cloning

The genes encoding *Bacillus globisporus*, *Bacillus subtilis*, and *Geobacillus stearothermophilus* AK were PCR amplified from pET26b-BgAK, pET26b- BsAK, pET26b-GsAK. The DNA for the wild-type and permutant AK enzymes were amplified from constructs reported previously and combined using Golden Gate assembly with pET28b.^37^ Plasmid DNA was sequence-verified using Sanger sequencing (Genewiz, NJ, U.S.). The N- and C- termini of the wild type enzyme were linked by the peptide Ala-Ala-Ala.

### Protein expression

Protein-expressing plasmids (WT, cp45 for each AK) were transformed using standard methods into competent *E. coli* cells. All expressing cultures were subcultured (1:100 inoculum) from overnight growths in lysogeny broth (LB) medium with 50 µg/mL kanamycin at 37 °C with shaking at 225 rpm. For *Gs*AK protein expression, Lemo21 DE3 competent cells expressing the pET28-*Gs*AK-WT plasmid were incubated in 1 L of LB medium with 50 µg/mL kanamycin at 37 °C with shaking at 225 rpm. *Gs*AK-cp45 protein expression was achieved in Tuner DE3 competent cells expressing the pET28-*Gs*AK-cp45 plasmid in similar conditions. For expression of *Bs*AK and *Bs*AK-cp45 proteins, Tuner DE3 competent cells transformed with the respective plasmids were incubated in LB medium supplemented with 1 mM zinc acetate, 1 mM magnesium chloride, and 50 µg/mL kanamycin at 37 °C, with shaking at 225 rpm. Once the cells reached an optical density (600 nm) of 0.6, 50 µM of IPTG was added to the cultures to induce protein production. After 18 hours of induction at 16 °C with shaking at 150 rpm, the cells were harvested by centrifugation (6000 rpm, 4 °C, 10 min) and the pellet was frozen at -80 °C for later purification of protein.

### Enzyme purification

All chromatography columns were from GE Healthcare. Cells were resuspended in buffer A (5 mL per g of wet cell mass) containing 50 mM tris pH 7.5, 25 mM NaCl, 10 mM MgCl2, 1 mM EDTA, and protease inhibitor cocktail (Roche), at a ratio of 25 μL of tablet solution (from a stock in 1 mL) per 10 mL of buffer. Cells were lysed using sonication, and lysate was clarified by centrifugation at 48,000 g for 40 min at 4 °C, and the cell-free extract was filtered through a 0.22 μM PES filter (Millipore). The sample was applied to a Hitrap Blue HP column, and eluted using stepwise increases in NaCl concentration (180 mM, 750 mM, 1.5 M). The fractions containing protein (as analyzed by SDS-PAGE) were pooled, and buffer exchanged into buffer A. The sample was applied to a Hitrap Q XL column (GE Healthcare) which was previously equilibrated with buffer A and eluted using a gradient of NaCl (25-375 mM). The protein peak was collected and concentrated using a 10 kDa Amicon Ultra 15 concentrator cell (Millipore), and then applied to a Superdex 200 16/600 pg size exclusion column equilibrated with 25 mM Tris pH 7.5, 150 mM NaCl. The protein was eluted at 1 mL/min. For *Gs*AK, *Gs*AK-cp45 and *Bs*AK-cp45, analysis by SDS-PAGE indicated the protein was > 95% pure and was suitable for downstream experiments. Additional contaminants were present in the *Bs*AK sample. In this case, the sample was adjusted to 1 M (NH4)3SO4, centrifuged and applied to a Hitrap Phenyl HP equilibrated with 50 mM Tris pH 7.5, 1 M (NH4)3SO4. Protein was eluted using a linear gradient from 1 M – 0 M (NH4)3SO4; analysis of the resultant peak indicated the protein was > 95% pure. Protein samples were concentrated, aliquoted and flash-frozen in liquid N2 to store at -80 °C for later use. *Bg*AK- cp45 was unable to be overexpressed in a soluble form despite expression condition testing.

### Binding assays

Binding experiments were performed using microscale thermophoresis (MST).^39^ Experiments were performed on a Monolith NT.115 (NanoTemper) using 20% light-emitting diode power. Proteins were labelled in NHS labelling buffer using the Red NHS 2^nd^ generation amine labelling kit (Nanotemper, MO-L011), then buffer exchanged into 10 mM potassium phosphate buffer, pH 7.0. Binding was tested using 50 nM protein in the presence of increasing Ap5A concentrations. Data collection was performed with Nanotemper software v1.5.41. Analysis was performed using Origin Pro 9 (OriginLab).

### Activity assays

AK activity was measured using an endpoint-coupled assay as previously optimized and employed in our laboratory.^40^ The assay mixture contained buffer (25mM phosphate buffer pH 7.2, 5 mM MgCl2, 65 mM KCl), 1.4 mM AMP, and various ATP concentrations (5, 10, 25, 50, 100, 250, 500, 1000 and 1400 μM). The reaction was started by the addition of AK enzyme to a final concentration of 10 nM (with the exception of *Bs*AK- cp45, where 50 nM was used). The reaction mixture tube and AK sample tube were incubated at the desired temperature (25 – 80 °C) in a water bath for 5 min before addition of AK. Time- point measurements were taken at 20, 40, and 60 sec after initiation, by quickly transferring 90 µL of reaction mixture from the tube to 30 uL of Ap5A (1 mM) maintained on ice. Each reaction was performed as biological triplicates. The time-point samples were sealed and kept on ice until the amount of ADP was ready to be measured for all samples simultaneously. ADP was quantitated by a secondary assay in which 5 enzyme units (U) of pyruvate kinase, 0.3 mM NADH, and 0.5 mM phosphoenol pyruvate (PEP) were added to 100 μL of quenched reaction. Absorbance at 340 nm was measured before and after the addition of 5 U of lactate dehydrogenase. The conversion of NADH to NAD^+^ and resulting decrease in A340 was used as a measure of the ADP produced. The slope of [ADP] versus time for these three time points was used as the rate of product formation to determine the activity of each AK enzyme. Data were analyzed and plotted using Origin Pro 9 (OriginLab).

### Spectropolarimetry

The circular dichroism (CD) spectrum of each protein was recorded from 190-300 nm using a Jasco J-815 CD spectrometer with temperature control at 30 °C. Measurements were performed using 400 µL of 2 μM protein in a 2 mm pathlength quartz cuvette. The unfolding of AK protein was calculated by monitoring the α-helical structure at 220 nm as a function of temperature. AK enzyme was diluted from freshly thawed aliquots into 10 mM potassium phosphate buffer, pH 7.0. Samples of each AK isoform (each WT and cp45 enzyme, 5 μM) with and without 250 μM of Ap5A (Diadenosine Pentaphosphate) were incubated on ice for 10 min. Melting curves were measured from 20-98 °C using 400 μL of sample in a 2 mm pathlength quartz cuvette. Data were analyzed using OriginPro software.

### Proteoloysis assays

A time-course assay employing trypsin (Sigma Aldrich) was devised to investigate the degradation of the adenylate kinases. Separate solutions of AK enzyme (10 μg total mass; 50 μL from a 1 mg/mL stock) and trypsin (1 mg/mL stock in H2O) were incubated at 30 °C for 5 min. Trypsin (1 μg total mass; 5 μL from stock solution) was then added to initiate digestion of the AK enzyme. At several time intervals (1, 2, 5, 10 min), samples were taken by pipetting 11 μL of reaction mixture into 3.5 μL of 4 x SDS-PAGE sample buffer that was pre-chilled on ice. Immediately after this, each sample was flash-frozen in liquid N2 and stored at -80 °C until analysis could be performed on all samples. The same experiment was also performed with 250 μM of Ap5A substrate present in the AK sample tube (5 μL of a 4 mM stock), allowing samples to sit at 30°C for 10 min to allow for binding of Ap5A before the addition of trypsin. When all samples had been collected and frozen at -80 °C, they were analyzed using SDS-PAGE.

### Mass spectrometry

Samples for protein MS were prepared by excising gel bands from the trypsin digestion experiments, according to the commonly employed in-gel trypsin digestion protocol.^41^ LC-MS/MS Analysis was conducted using a Shimadzu Prominence LC- 20AD XR UPLC system interfaced to a Shimadzu Ion Trap – Time of Flight (IT-ToF) Mass Spectrometer (MS) through an Electrospray Ionization (ESI) source. Separations were achieved using a 1.0 mm ID x 150 mm, 2.7 micron particle size Ascentis Express Peptide ES- C18 column. The injection volume was 25 μL. The column was operated at a flow rate of 0.175 mL/min with mobile phase A: 0.1% (v/v) formic acid in water and mobile phase B: 0.1% (v/v) formic acid in acetonitrile. The LC gradient for the analytical separation was operated as follows; 5% B (IC), 5% B to 30%B (0.0 min to 12.5 min), 30%B to 50%B (12.5 min to 14.5 min), and 50%B to 95%B (14.5 min to 16.0 min). The IT-ToF MS was operated in the positive ion mode with data dependent MS/MS acquisition enabled. MS spectra were acquired over the range of m/z 300 to m/z 1500 and MS/MS spectra were acquired over the range of m/z 150 to m/z 1500. Protein identification analysis was conducted using Mascot v. 2.7.^42^ Peptides were searched at significance threshold of p < 0.05 against the sequences of the desired AK enzymes as well as against bacterial enzymes, such as AK from *E. coli,* and common contaminating peptides. Proteins were considered detected with ≥ 3 observed peptide sequences, including oxidations and modifications.

### Structure prediction

Structure prediction models of circular permutants of adenylate kinases were produced using the simplified algorithm for AlphaFold2, titled ColabFold, freely available online at https://github.com/sokrypton/ColabFold.^38^

## RESULTS

### Binding of Ap5A to permuted Aks

A prior study found that cp45 permutants of three AK homologs (*Gs*AK, *Bs*AK, and *Bg*AK) retained function in cells at 40°C (Figure 1B),^37^ but this study did not explore the effects of circular permutation on their activity, structure, and thermostability. We were able to express and purify two AK homologs (*Gs*AK and *Bs*AK) and their respective cp45 permutants at sufficient levels to characterize biochemically. While a cp45 homolog derived from *Bg*AK was also found to retain cellular function in a prior circular permutation study,^37^ it was excluded from our study as this variant could not be produced at sufficient levels as a soluble protein when overexpressed in *E. coli*.

When new N- and C-termini are introduced into the AMP binding domain, an increase in conformational flexibility is expected, which could alter the binding affinity for substrates and the ability of the enzyme to modulate the AMP + ATP ⇄ 2 ADP equilibrium. To test these ideas, we investigated the binding affinity of each AK for the non-hydrolyzable transition state analog (Figure 1C), Ap5A, using microscale thermophoresis (Figure 2).^39^ This analysis revealed that native *Gs*AK binds Ap5A ∼4-fold tighter (Kd = 17.0 ± 5.7 nM) than *Gs*AK-cp45 (Kd = 69.0 ± 14.3 nM). We observed the same trend with *Bs*AK. Native *Bs*AK binds ∼5-fold more tightly to Ap5A (Kd = 25.1 ± 1.1 nM) compared to the *Bs*AK-cp45 (Kd = 116 ± 20 nM). These findings show that circular permutation of both AKs, which creates new N- and C-termini within the α-helix at position 45, does not dramatically alter Ap5A binding, although it decreases the affinities modestly and to similar extents.

**Figure 2.**
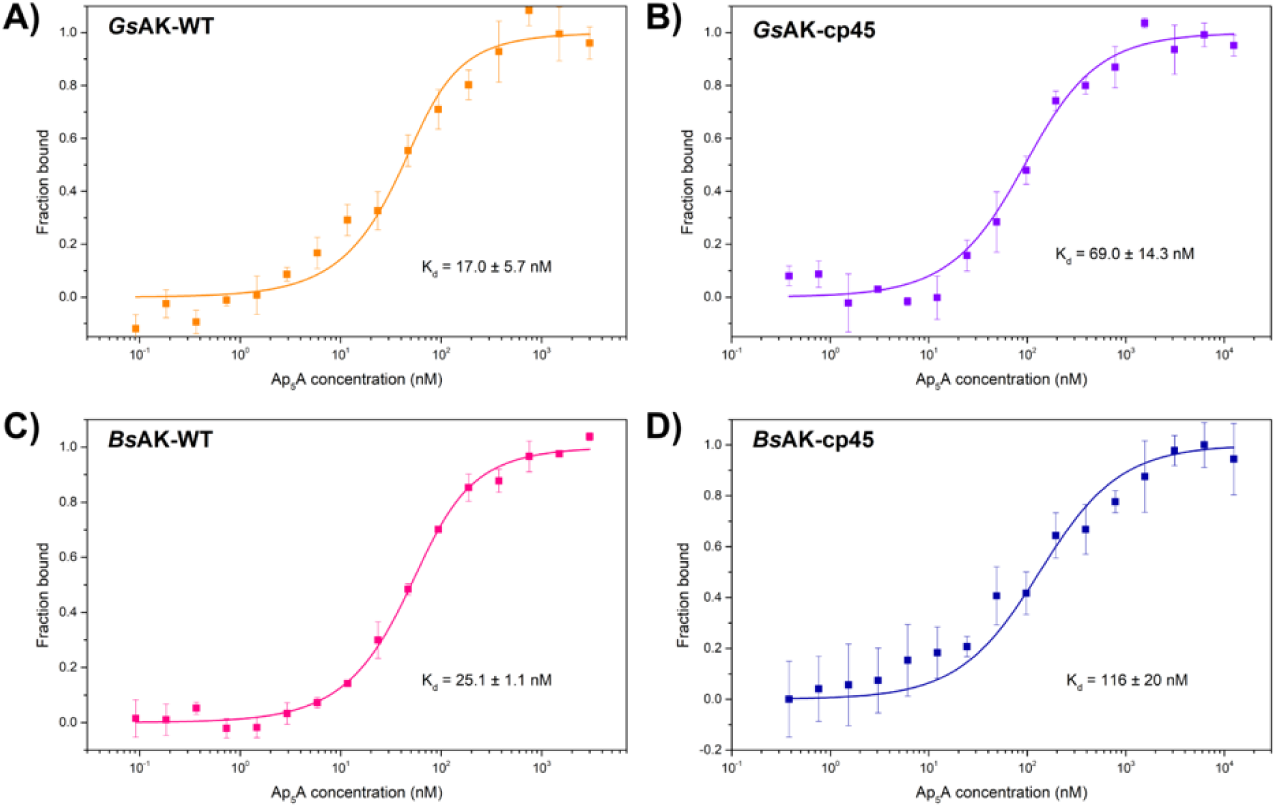
Effect of circular permutation on AK and Ap_5_A binding. Microscale thermophoresis analysis of Ap5A binding to (**A**) *Gs*AK, (**B**) *Gs*AK-cp45, (**C**) *Bs*AK, and (**D**) *Bs*AK-cp45. All measurements were made in triplicate at 20 °C, and data are plotted with error bars that represent ±1 standard deviation.

### Protein unfolding measurements

The native AK residues coding for the AMP binding and lid domains are continuous, while the residues coding for the core domain are found in three discontinuous regions of the primary structure. In contrast, with cp45, the core domain is coded in two different regions of the primary structure, and the AMP binding domain is split so that it is also coded in different regions of primary structure. We hypothesized that this change in domain contact order might affect protein folding. To test this idea, we used circular dichroism (CD) spectroscopy to investigate the extent to which AK folding and thermostability was affected by circular permutation. Prior studies showed that AKs exhibit melting curves consistent with the two-state model due to cooperative folding.^22, 27, 29^

CD spectra suggested that each protein was folded at 30 °C, exhibiting characteristic peaks at ∼210 and 220 nm as expected for structures containing α-helices (Figure S1).^43^ The melting temperatures of each AK was estimated by fitting either a single or double logistic function to temperature ramping experiments. The melting temperature of *Gs*AK was 75.9 °C (Figure 3A), in agreement with prior estimates (74.5 °C), and this protein presented a simple two-state transition.^29, 44^ In the presence of Ap5A, the melting temperature increased to 88.5 °C. *Gs*AK-cp45 was less thermostable than *Gs*AK (Figure 3B). In addition, *Gs*AK-cp45 exhibited multiple unfolding transitions making a direct estimate of the Tm impossible using a two-state model. A fit to this data to a double logistic function yielded melting transitions with midpoints at 62.3 and 67.3°C (Table 1). The presence of multiple transitions for *Gs*AK-cp45 also occurred in the presence of Ap5A. With bound Ap5A, the first transition for *Gs*AK-cp45 was largely unchanged, while the second transition occurred at a higher temperature than with the substrate-free curve. These data suggest that introduction of new N- and C-termini in the AMP binding domain of *Gs*AK leads to discrete unfolding of domains at different temperatures, and they show that Ap5A binding stabilizes *Gs*AK-cp45 like native *Gs*AK.

**Figure 3.**
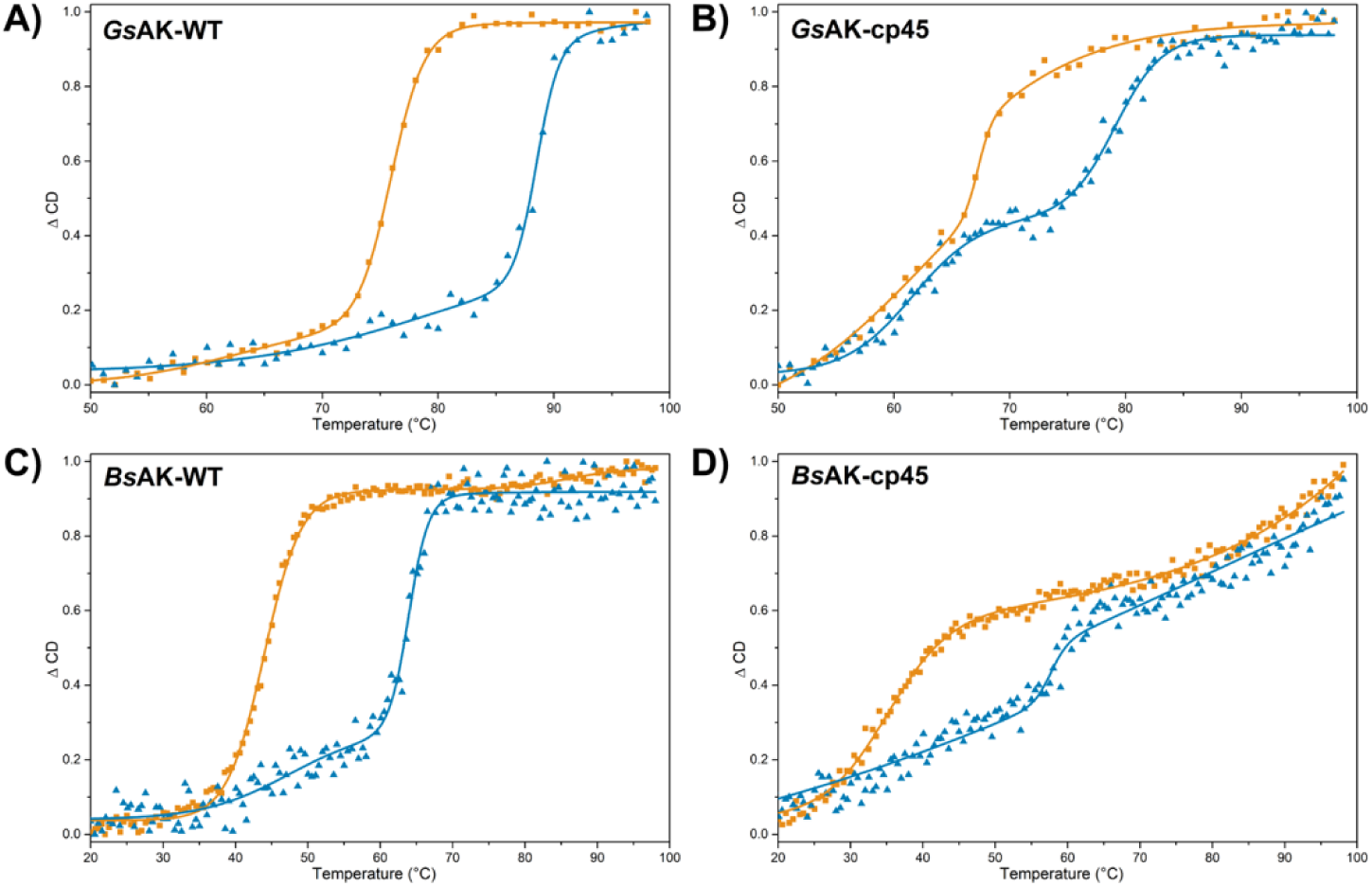
Effect of temperature on AK structure. Melting curves for: (**A**) *Gs*AK, (**B**) *Gs*AK- cp45, (**C**) *Bs*AK, and (**D**) *Bs*AK-cp45 in the absence (orange) and presence (blue) of Ap_5_A. Unfolding curves were fit to either a single or double logistic functions. The midpoint transitions from each fit are provided in Table 1.

**Table 1.** Melting temperature (T_m_) determination for AK enzymes using the change in ellipticity at 220 nm. T_m_ was estimated by fitting the data to the double logistic function. Two numbers are shown when two transitions were calculated when fitting the data. Prior values reported in the literature values are in brackets.^29, 44^ R^2^ > 0.99 in all cases.

| AK | $T_m$ (-Ap<sub>5</sub>A) | $T_m$ (+Ap<sub>5</sub>A) | Apparent $\Delta T$ on binding<sup>i</sup> |
| --- | --- | --- | --- |
| GsAK | 75.9 [74.5]<sup>ii</sup> | 88.5 [$> 70.0$]<sup>ii</sup> | 12.6 |
| GsAK-cp45 | 62.3, 67.3 | 61.6, 79.1 | 11.8 |
| BsAK | 43.9 [47.6]<sup>ii</sup> | 47.4, 63.9 [66.0]<sup>ii</sup> | 20.0 |
| BsAK-cp45 | 35.0 | 57.7 | n.d.<sup>iii</sup> |

<sup>i</sup> Change in apparent unfolding temperature upon binding of Ap<sub>5</sub>A. The transition with the highest slope is here reported. <sup>ii</sup> For GsAK, changes in the CD signal were not observed up to the temperature tested and the experiment was stopped early.<sup>29</sup> <sup>iii</sup> BsAK-cp45 appeared to unfold steadily so $\Delta T$ was not calculated.

*Bs*AK had an unfolding transition at 43.9°C (Figure 3C), which increased to 63.9°C in the presence of Ap5A. This Ap5A stabilization was larger than that observed with *Gs*AK. With *Bs*AK, a fit of the data to a double logistic function showed a shallow transition before the larger unfolding transition. With *Bs*AK-cp45 (Figure 3D), the substrate-free melting curve could not be fit to a two-state unfolding model. The melting temperature was estimated to be 35.0°C for the steepest transition (Table 1). Upon binding of Ap5A, *Bs*AK-cp45 appeared to unfold continuously over the assayed temperature range, with a small sharp transition at 57.7°C. We investigated how increasing the Ap5A concentrations affects these transitions, which revealed that the enzyme was saturated at 250 μM Ap5A (*data not shown*). These data show that circular permutation decreases *Bs*AK stability as observed with *Gs*AK, and they illustrate how the unfolding transitions are affected by permutation.

### Effect of permutation on activity

While both cp45 homologs presented phosphotransferase activity in a cellular assay at 40°C,^37^ the effect of permutation on their catalytic activity is not known. To directly examine the catalytic activities of these circularly permuted kinases, we examined their ability to convert ATP and AMP to ADP across a range of temperatures and ATP concentrations as previously described.^40^ We first studied the maximal activity of each AK variant over a range of temperatures (Figure 4A), using constant AMP and ATP starting concentrations (Activity units: μM-ADP/min/nM-AK, henceforth written as min^-1^). With *Gs*AK, maximal activity was observed at 70°C (38.5 min^-1^). In contrast, *Gs*AK-cp45 presented maximal activity at 60°C (20.1 min^-1^). *Gs*AK and *Gs*AK-cp45 also had similar activities at 40°C. While *Gs*AK activity was consistently higher than *Gs*AK-cp45 above this temperature, *Gs*AK-cp45 was slightly higher than *Gs*AK below this temperature. With BsAK, the maximal activity (12.3 min^-1^) occurred at 45°C, while *Bs*AK-cp45 had maximal activity (2.2 min^-1^) at 35°C. These results show that circular permutation does not abolish activity in either homolog, and they implicate circular permutation as increasing *Gs*AK activity below 40°C.

**Figure 4.**
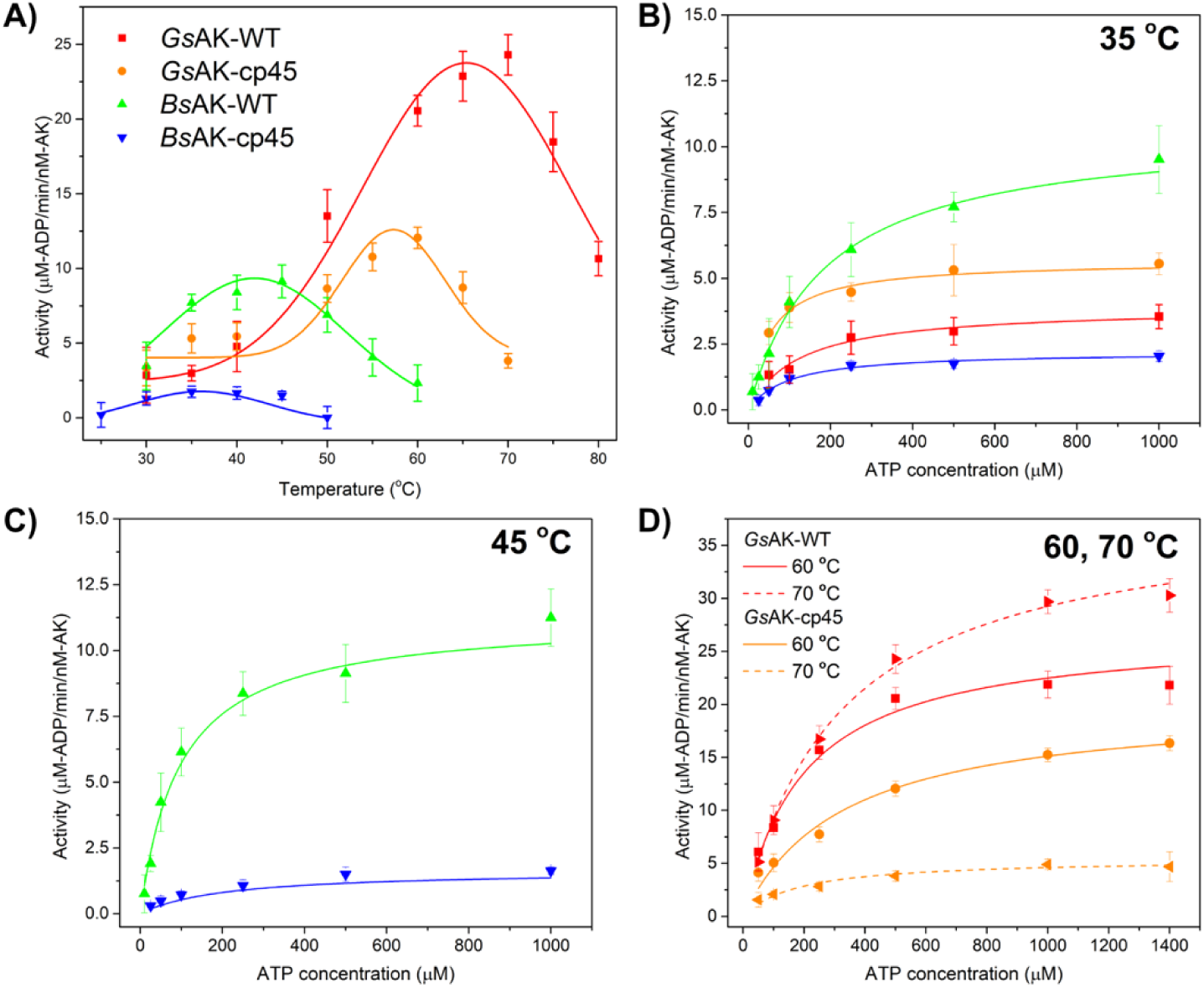
Kinetic analysis of each AK. (**A**) Effect of temperature on *Gs*AK (red), *Gs*AK-cp45 (orange), *Bs*AK (green), and *Bs*AK-cp45 (blue) in the presence of 500 uM ATP and 1.4 mM AMP. Each point represents three or more measurements, with values reported as μM-ADP/min/nM-AK. (**B- D**) Effect of varying ATP concentration on the kinetic parameters at four different temperatures. Error bars represent ±1σ.

The Michaelis-Menten kinetic parameters were determined at 35°C for all AKs (Figure 4B), at 45°C for *Bs*AK and *Bs*AK-cp45 (Figure 4C), and at 60 and 70°C for *Gs*AK and *Gs*AK- cp45 (Figure 4D). Table 2 summarizes this analysis. At 35°C, *Bs*AK had the highest *kcat/Km*, which doubled at 45°C. In contrast, the *kcat/Km* for *Bs*AK-cp45 at 45°C was lower compared with that at 35°C. The *kcat/Km* for *Gs*AK and *Gs*AK-cp45 were maximal at 60°C, although the measured *kcat* for *Gs*AK was higher at 70°C. The *Km* values also varied with temperature. At the lowest temperature (35°C), permutation decreased the *Km* value by ∼50%. At 45°C, *Bs*AK and *Bs*AK-cp45 had similar *Km*. At 60°C, permutation of *Gs*AK led to a ∼50% increase in *Km*, but at 70°C this trend was reversed. While the permuted AKs presented lower activities at most temperatures assayed, the activity of *Gs*AK-cp45 was higher than *Gs*AK at 35°C across all substrate concentrations assayed. These findings show that permutation decreases the maximal activities of AKs when assayed near temperatures that present the maximum activity. They also show that *Gs*AK-cp45 presents enhanced activity when assayed at a temperature that is 30°C lower than the optimal temperature of the *Gs*AK.

**Table 2.** Kinetic parameters determined for AK enzyme variants studied. *k_cat_* units are μM- ADP/min/nM-AK, or min^-1^; *K_m_* are μM; *k_cat_/K_m_* values are (M-AK)^-1^min^-1^.

| Temperature | Enzyme | $k_{cat}$ [$\text{min}^{-1}$] | $K_m$ [$\mu\text{M}$] | $k_{cat}/K_m$ [$(\text{M-AK})^{-1}\text{min}^{-1}$] |
| --- | --- | --- | --- | --- |
| 35 °C | GsAK | $3.90 \pm 0.24$ | $123 \pm 27$ | $3.17 \pm 0.891 \times 10^7$ |
| | GsAK-cp45 | $5.66 \pm 0.21$ | $49.9 \pm 8.5$ | $1.13 \pm 0.235 \times 10^8$ |
| | BsAK | $10.7 \pm 0.4$ | $183 \pm 18$ | $5.85 \pm 0.794 \times 10^7$ |
| | BsAK-cp45 | $2.21 \pm 0.10$ | $95 \pm 15$ | $2.33 \pm 0.473 \times 10^7$ |
| 45 °C | BsAK | $12.3 \pm 0.7$ | $124 \pm 16$ | $9.92 \pm 1.84 \times 10^7$ |
| | BsAK-cp45 | $1.85 \pm 0.07$ | $139 \pm 15$ | $1.33 \pm 0.841 \times 10^7$ |
| 60 °C | GsAK | $26.9 \pm 1.6$ | $197 \pm 32$ | $1.37 \pm 0.303 \times 10^8$ |
| | GsAK-cp45 | $20.1 \pm 1.4$ | $331 \pm 69$ | $6.07 \pm 1.69 \times 10^7$ |
| 70 °C | GsAK | $38.5 \pm 1.0$ | $316 \pm 24$ | $1.22 \pm 0.124 \times 10^8$ |
| | GsAK-cp45 | $5.43 \pm 0.44$ | $180 \pm 41$ | $3.02 \pm 0.932 \times 10^7$ |

### Effect of permutation on structure

We speculated that the creation of new N- and C-termini within the AMP-binding domain would destabilize this domain and decrease the overall stability of each AK. To test this hypothesis, we investigated if the enzymes varied in their kinetics of degradation when exposed to trypsin. We posited that partial trypsin proteolysis would reveal localized differences in folding and stability, due to changes in the accessibility of lysine and arginine (R/K) residues arising from permutation. The presence or absence of peptides near these R/K cleavage sites may also indicate the stability of each domain. The WT and cp45 variants each contain the same number of Arg and Lys residues and thus the same number of potential trypsinization sites. *Gs*AK and *Bs*AK each contain 4 such sites in the AMP-binding domain, and these are in conserved positions (Figure S2). Each AK was incubated with trypsin for varying durations (1, 2, 5, 10 min), snap-frozen in liquid N2, and then analyzed using SDS-PAGE. Control reactions with the trypsin omitted showed that the SDS-PAGE buffer and liquid N2 did not degrade the AK enzymes (lanes marked C, Figure 5).

**Figure 5.**
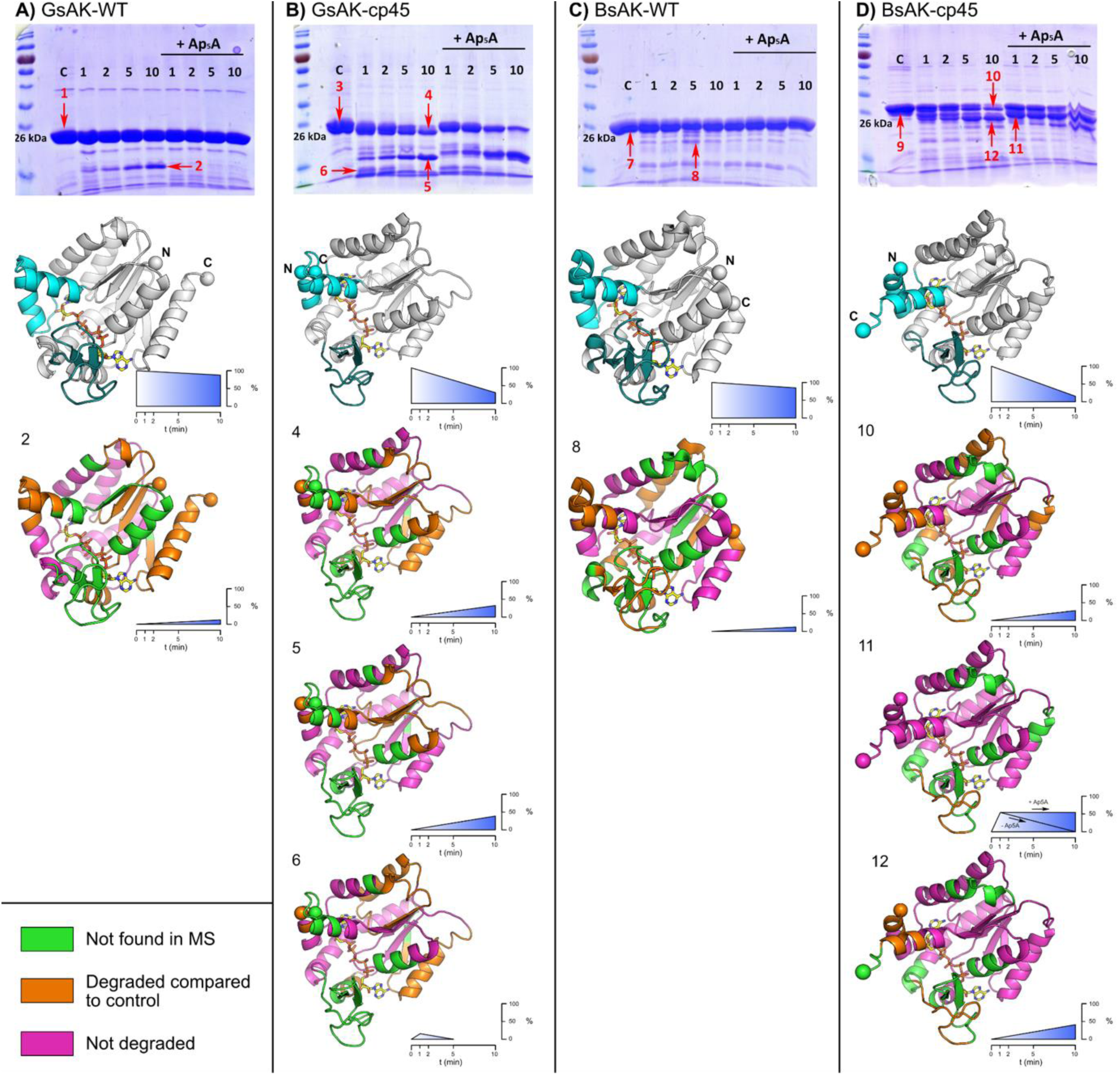
AK sensitivity to trypsin. (**A**) *Gs*AK, (**B**) *Gs*AK-cp45, (**C**) *Bs*AK, and (**D**) *Bs*AK-cp45 were subjected to trypsin digestion and analyzed using SDS-PAGE and MS. *Top of each panel:* SDS-PAGE gel for each AK. The control reaction (noted as C) lacking trypsin is followed by samples lacking and containing Ap_5_A Ap_5_A (250 μM), with the time of trypsin incubation noted (1, 2, 5, 10 min). Red arrows indicate bands analyzed using MS. *First structure in each subpanel:* reference model of each AK enzyme showing the core (grey), AMP (cyan), and lid (dark blue) domains, which were generated using PDB 1ZIP and 1P3J. The cp45 variant models were generated using ColabFold.^38^ The N- and C- termini are shown as spheres, and Ap_5_A is shown with standard CPK atom coloring (yellow carbons). *Numbered structures:* Mass spec fragments mapped to AK structures, revealing the regions being degraded over time by trypsin. Green regions indicate peptides that were not detected in the full-length control, orange indicates regions which were not observed following trypsinization, indicating domains that were sensitive to trypsin cleavage. Pink regions indicate those observed in both digested and control samples, indicating stability to trypsin. The blue color scales represent the change in the species’ abundance over time, estimated visually as percentage of the protein bands on the SDS-PAGE. Mascot search reports are provided as a supplemental file.

The digestion results indicated that *Gs*AK is highly resistant to 10 min trypsinization (Figure 5A, top), with a single degradation product appearing on the SDS-PAGE gel. The addition of Ap5A further decreased degradation. In contrast, *Gs*AK-cp45 was significantly more prone to degradation in the absence and presence of Ap5A (Figure 5B, top), with two major degradation species increasing in intensity as the full length protein decreased in intensity. A similar trend was observed with *Bs*AK (Figure 5C, top) and *Bs*AK-cp45 (Figure 5D, top). This analysis shows that localized disruption of the AMP-binding domain increases the overall sensitivity of these AKs to trypsinization.

We next investigated whether the pattern of proteolysis arises from backbone cleavage proximal to native residue 45 or if the cleavage sensitivity was more global. Protein fragments of interest from SDS-PAGE were identified using standard in-gel trypsinization and LC-MS.^41^ In total, twelve different species (indicated with red arrows) and the control bands were analyzed. Detected peptides were compared to sequences of the target adenylate kinase, common bacterial contaminants, and trypsin; the latter does not have overlapping regions with the AK using Clustal Omega.^45^ MS analysis of Species 1 (*Gs*AK) detected ∼70% of the total sequence (Table S1), including most of the core and AMP binding domains (Figure 5A, bottom). Peptides from the lid domain were not detected. The major *Gs*AK degradation product (Species 2) was comprised of peptides within the core domain and a smaller AMP-binding domain fragment (residues 50-59); the remainder of this domain was not observed. This degradation was not observed when adding trypsin to Ap5A-bound *Gs*AK. These finding show that the AMP-binding domain in *Gs*AK can be detected and is stabilized by Ap5A binding.

*Gs*AK-cp45 was degraded to a greater extent (Figure 5B, bottom), with little stabilization conferred by Ap5A. In the control band (Species 3), peptides for much of the core domain were observed, as well as a portion of the AMP binding domain (residues 21-33 and 225-230 but not 231-240). There were three major degradation bands observed, two of which increased with time (Species 4 and 5) as Species 3 disappeared. Species 4 and 5 were missing more of the N-terminal AMP binding domain, and most of the C-terminal AMP domain, with further core domain degradation. Interestingly, in Species 4 and 6, the AAA linker was observed, while the lid domain was not. In addition, both mobile domains as well as the N- and C-termini were degraded before the core domain, and the split in the AMP domain significantly exacerbated this process, presumably by reducing the stabilizing effect from Ap5A binding. These findings illustrate differences in the patterns of degradation for *Gs*AK-cp45 and *Gs*AK, and they show that the split AMP-binding domain in *Gs*AK-cp45 presents increased sensitivity to digestion near the new protein termini.

*Bs*AK showed proteolytic sensitivity that was distinct from *Gs*AK. Without Ap5A, very little degradation was observed (Figure 5C, bottom) and ∼71% sequence coverage was obtained (Species 7). All residues in the AMP-binding domain were observed, while only half of the lid domain was detected (residues 146-159). Only one degradation product (Species 8) was detected, which was predominantly the core domain with most of the AMP domain. The lid domain was not observed in Species 8, suggesting it was digested in the absence of Ap5A. These findings reveal that the *Bs*AK lid domain, like that of *Gs*AK, is the most sensitive to digestion.

*Bs*AK-cp45 was degraded to three major products with similar apparent molecular masses (Figure 5D, bottom). Over the 10 min reaction, Species 10 and 12 accumulated as the full-length enzyme (Species 9) decreased. Species 11 appeared at the beginning of the degradation reaction and then disappeared after 10 minutes. In Species 9, the entirety of the AMP domain was observed (residues 21-33 and 225-240). Furthermore, half of the lid domain was observed, similarly to *Bs*AK (residues 119-133). Species 10 and 12 increased over time when Ap5A was omitted. In both Species 10 and 12, a small portion of the N-terminus AMP domain was absent (residues 21-24), and a significant section of the core domain was absent in Species 10 (residues 63-97). The lid domain was also digested in both Species 10 and 12. When Ap5A was included with *Bs*AK-cp45, the distribution moved to an accumulation of Species 11. Notably, the two halves of the AMP domain were intact in Species 11. These findings show that Ap5A positioned the split AMP domain of *Bs*AK-cp45 such that trypsin has poorer access to cleavage sites.

## DISCUSSION

Circular permutation is broadly tolerated by a range of enzymes,^46^ but it is difficult to predict how this structural rearrangement affects protein structure and function, even within model proteins like AKs whose sequence-structure-function relationships have been extensively characterized.^4, 5, 8–18, 22, 47^ In AKs, the AMP-domain presents local energetic frustration,^37^ and local unfolding of this domain has been implicated as a rate-controlling motion for catalytic turnover.^48^ This observation has implicated conformational flexibility as a control dial on the AK enzymatic reaction cycle. While a prior study showed that mutational lesions that alter flexibility and conformational freedom are uniformly tolerated across AK homologs when generated in the AMP-binding domain,^37^ it was unclear how this type of sequence change affects AK function. Herein, we show the biochemical impact of altering conformational flexibility in the AMP-binding domain. At most temperatures assayed, the changes in binding affinity, activity, and temperature optimum were comparable in variants derived from the thermophilic (*Gs*AK) and mesophilic (*Bs*AK) protein homologs. One exception was when the activity of the thermophilic *Gs*AK was assayed at low temperatures. In this case, there was in a modest increase in catalytic activity at mesophilic temperatures (<45°C) compared with the parental AK. This result is similar to a prior study on chimeric AK enzymes. which revealed that swapping the domains among AK homologs with differing stabilities yielded chimeric AK where the stability was controlled by the core domain and the activity level was controlled by the source of the mobile domains (AMP and lid).^47^ Similar to our findings, chimeric AKs with mobile domains from mesophiles fused to thermophilic core domains presented increased catalytic activity at mesophile growth temperatures relative to the thermophilic parent protein. Taken together with our results, these findings suggest a mechanism by which increased conformational flexibility can be introduced into thermophilic proteins to increase activity at low temperatures.

In addition to affecting catalysis,^35, 49^ circular permutation can impact protein folding,^49–51^ energetic frustration,^52^ and stability.^53–55^ The effects on folding are especially evident in multidomain proteins with complex topologies where domains are embedded within discontinuous domains. For example, experimental studies of permutants of a two-domain protein, T4 lysozyme,^50^ showed that changes in the topology of the polypeptide backbone by relocating an embedded domain critically impacted folding cooperativity between the two domains of this protein. AK, like T4 lysozyme, is a multidomain protein with a complex topology consisting of two continuous domains (AMP and lid) that are embedded within a discontinuous domain (core). A previous computational study of AK suggested that this complex domain topology facilities cooperative folding.^4^ With the permuted proteins characterized herein, the creation of new protein termini within the middle of the AMP-binding domain makes this domain discontinuous because it is no longer embedded within the primary structure of the core domain. Our thermal denaturation study of *Gs*AK-cp45 and *Bs*AK-cp45 reveals that this permuted protein no longer presents a fully cooperative folding pathway, and instead has a multiphasic unfolding process when Ap5A is bound. We speculate that the observed multiphasic transitions arise from two distinct unfolding events, one where the discontinuous AMP-binding domain begins to unfold and the other corresponding to unfolding of the rest of the enzyme. Taken together with previous results,^4, 50^ this finding shows how altering the continuity of embedded domains can disrupt cooperative unfolding in multidomain proteins.

Thermostability has been demonstrated previously to be one possible mechanism to protect proteins against disruptions like random mutations and backbone fission.^56–58^ Therefore, it might be expected that AKs with lower thermostability will be more sensitive to circular permutation. Here, our findings suggest that the core domain is not significantly disrupted by extraction of the AMP domain, and thus the magnitude in reduction to melting temperature is similar regardless of parental thermostabilities. Conversely, the change in mobility of the AMP domain manifested as dramatic changes in susceptibility to trypsinization. This showed that the cooperativity of folding had been decreased, thereby increasing the exposure of the core domain of each enzyme. This result did not appear to correlate strongly to the thermostability of each AK, instead suggesting that individual domains can deviate from this behavior. The permuted enzymes may furthermore exist in a dynamic range of partially unfolded/folded states that increased susceptibility to trypsin. These experiments revealed that the AMP domain was more stable in the WT variants than the cp45 variants, which suggested different conformations of the domain. Taken together, these results demonstrate that modifications to a domain involved in catalysis can yield dramatic changes in kinetic and thermodynamic characteristics.

## Supporting information

Supplemental File - Mascot search reports

## AUTHOR CONTRIBUTION STATEMENTS

Tom Coleman performed experiments, designed the study, analyzed data, and wrote the manuscript. John Shin performed experiments and wrote the manuscript. Joshua T. Atkinson designed the study, analyzed data, and wrote the manuscript. Jonathan J. Silberg wrote the manuscript. Yousif Shamoo designed the study, analyzed data and wrote the manuscript. All authors approved the final submitted manuscript.

## ACKNOWLEDGEMENTS

This project was supported by DOE grant DE-SC0014462 (to J.J.S.). J.T.A. was supported by a Loideska Stockbridge Vaughn Fellowship and then the NSF Postdoctoral Research Fellowships in Biology Program under Grant No. 2010604. The authors gratefully acknowledge the support of the Rice SEA Proteomics Core. MS samples were kindly run and analyzed by Dr. Christopher Pennington.

## ABBREVIATIONS

AK: Adenylate Kinase
cp45: circular permutant at position 45
WT: Wild type
Ap5A: Diadenosine pentaphosphate
*Gs*AK: *Geobacillus stearothermophilus* Adenylate Kinase
*Bs*AK: *Bacillus subtilis* Adenylate Kinase
*Bg*AK: *Bacillus globisporus* Adenylate Kinase
AMP: Adenosine monophosphate
ADP: Adenosine diphosphate
ATP: Adenosine triphosphate

## Notes

### Competing Interest Statement

The authors have declared no competing interest.

