## Supporting Information for "The biochemical impact of extracting an embedded adenylate kinase domain using circular permutation"

Supporting information includes:

Figure S1. Ellipticity data for native AKs

Figure S2. Trypsin digestion sites (R/K) mapped onto AK sequences

Table S1. Collection statistics for MS samples

Supplemental File. Mascot search reports

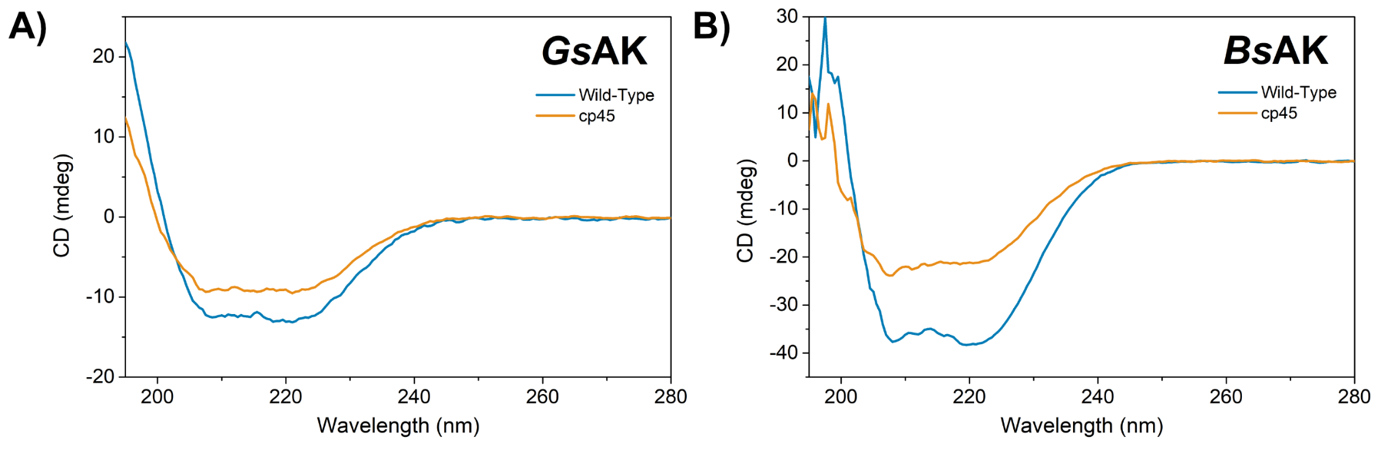

**Figure S1.** Circular dichroism spectroscopy scans of each AK investigated in this work. (**A**) a comparison of *Gs*AK and the cp45 variant derived from *Gs*AK, and (**B**) *Bs*AK and the cp45 variant derived from *Bs*AK. In both cases, the cpAK presents decreased ellipticity.

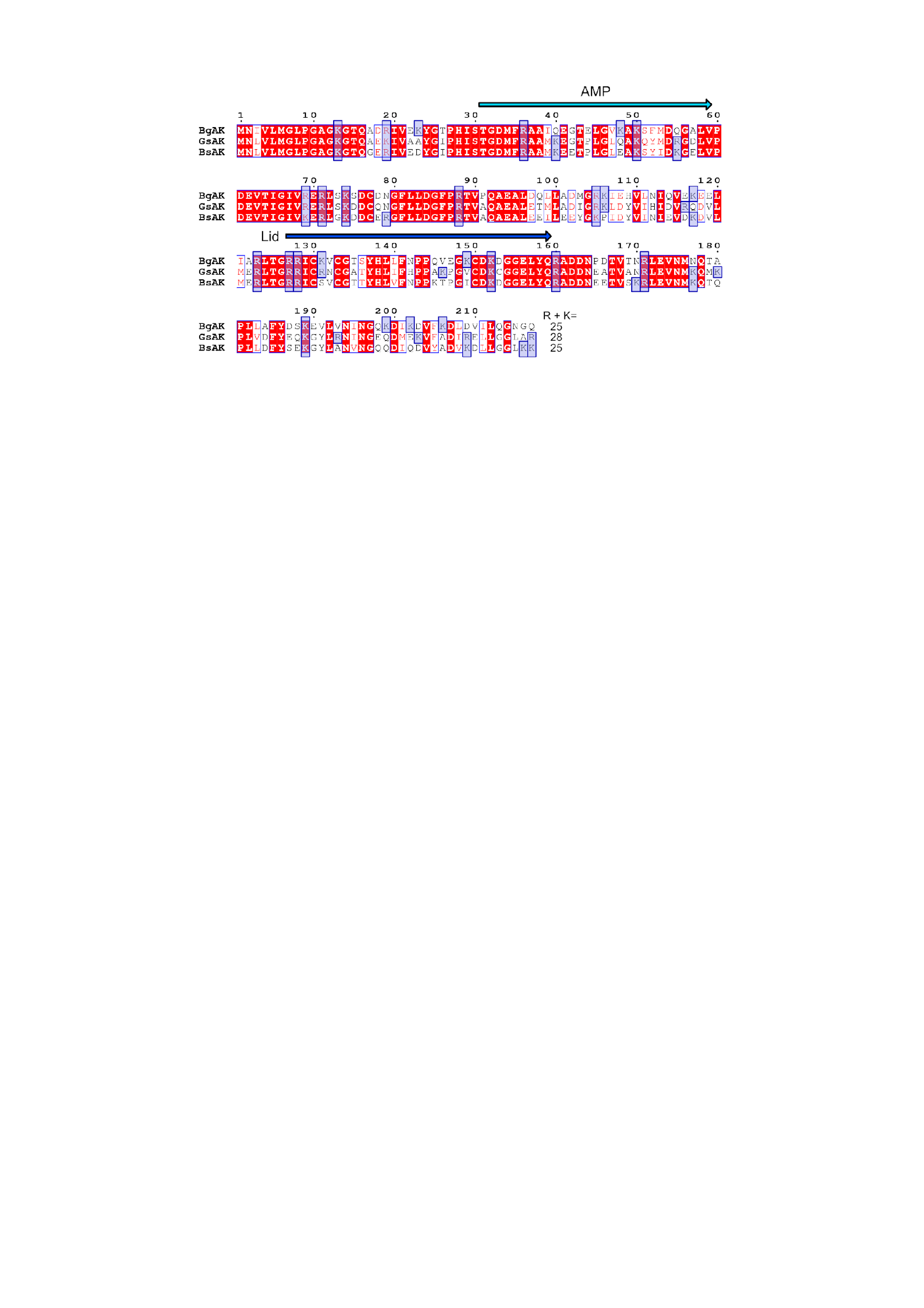

**Figure S2.** **Trypsin digestion sites (R/K) mapped onto AK sequences.** Blue boxes indicate potential cleavage sites. The AMP-binding and lid domains are shown with arrows.

**Table S1. Collection statistics for MS samples.** The Peptide MW is the calculated molecular mass by considering all observed peptides with ≥ 3 observations having a mass error ≤ 25 ppm, including modifications (such as oxidation). The control (C) band data is also provided which represents the analysis of the full-length enzyme.

| **AK enzyme** | **Species** | **Coverage % ^i^** | **Peptide MW (kDa, Total length) ^ii^** |
| --- | --- | --- | --- |
| *Gs*AK-WT C | 1 | 69 | 16.81 (151aa) |
| *Gs*AK-WT | 2 | 35 | 8.68 (76aa) |
| *Gs*AK-cp45 C | 3 | 70 | 18.67 (169aa) |
| *Gs*AK-cp45 | 4 | 38 | 10.18 (92aa) |
| *Gs*AK-cp45 | 5 | 44 | 12.22 (107aa) |
| *Gs*AK-cp45 | 6 | 38 | 10.23 (92aa) |
| *Bs*AK-WT C | 7 | 71 | 17.47 (156aa) |
| *Bs*AK-WT | 8 | 35 | 8.65 (77aa) |
| *Bs*AK-cp45 C | 9 | 75 | 19.75 (180aa) |
| *Bs*AK-cp45 | 10 | 38 | 10.15 (93aa) |
| *Bs*AK-cp45 | 11 | 65 | 17.29 (157aa) |
| *Bs*AK-cp45 | 12 | 59 | 15.86 (143aa) |

^i^ Sequence coverage as reported using Mascot search. A score of approx. 70 % provides high confidence of the full-length target enzyme in the original sample. ^ii^ Calculated using all observed peptides, using the Expasy MW and p_i_ calculator (<https://web.expasy.org/compute_pi/>).
