## Supplemental File - Mascot search reports for "The biochemical impact of extracting an embedded adenylate kinase domain using circular permutation"

**SUPPLEMENTAL DATA**

Mascot search report files for each Species in Figure 5.

Page 2 - Species 1

Page 8 - Species 2

Page 11 - Species 3

Page 16 - Species 4

Page 19 - Species 5

Page 23 - Species 6

Page 25 - Species 7

Page 30 - Species 8

Page 32 - Species 9

Page 37 - Species 10

Page 40 - Species 11

Page 44 - Species 12

### Species 1

#### MATRIX SCIENCE MASCOT Search Results

##### Protein View: QNL33371.1

gsAK [Escherichia coli]

| | |
|---|---|
| Database: | NCBIcustTC |
| Score: | 4617 |
| Monoisotopic mass ($M_r$): | 24127 |
| Calculated pI: | 5.86 |

Sequencesimilarity is availableas an NCBI BLAST search of QNL33371.1 against nr.

##### Search parameters

| | |
|---|---|
| MS data file: | 09182020_TC-1_001.temp.mgf |
| Enzyme: | Trypsin/P: cuts C-term side of KR. |
| Variable modifications: | Oxidation (M), Propionamide (C), Propionamide (K), Propionamide (N-term) |

##### Protein sequence coverage: 69%

Matched peptides shown in **bold red**.

```
  1 MNLVLMGLPG AGKGTQAEKI VAAYGIPHIS TGDMFRAAMK EGTPLGLQAK
 51 QYMDRGDLVP DEVTIGIVRE RLSKDDCQNG FLLDGFPRTV AQAEALETML
101 ADIGRKLDYV IHIDVRQDVL MERLTGRRIC RNCGATYHLI FHPPAKPGVC
151 DKCGGELYQR ADDNEATVAN RLEVNMKQMK PLVDFYEQKG YLRNINGEQD
201 MEKVFADIRE LLGGLAR
```

Unformatted sequence string: 217 residues (for pasting into other applications).

Sort by (●) residue number ( ) increasing mass ( ) decreasing mass
Show (●) matched peptides only ( ) predicted peptides also
Show (●) uncorrected delta ( ) delta corrected for 13C

| Query | Start - End | Observed | Mr(expt) | Mr(calc) | ppm | M | Score | Expect | Rank | U | Peptide |
|---|---|---|---|---|---|---|---|---|---|---|---|
| 637 | 1 - 13 | 650.8512 | 1299.6879 | 1299.7043 | -12.6 | 0 | 29 | 0.0012 | 1 | U | -.MNLVLMGLPGAGK.G |
| 638 | 1 - 13 | 650.8513 | 1299.6880 | 1299.7043 | -12.6 | 0 | 23 | 0.0048 | 1 | U | -.MNLVLMGLPGAGK.G |
| 639 | 1 - 13 | 650.8568 | 1299.6990 | 1299.7043 | -4.08 | 0 | 19 | 0.011 | 1 | U | -.MNLVLMGLPGAGK.G |
| 640 | 1 - 13 | 650.8581 | 1299.7016 | 1299.7043 | -2.06 | 0 | 39 | 0.00012 | 1 | U | -.MNLVLMGLPGAGK.G |
| 641 | 1 - 13 | 650.8582 | 1299.7018 | 1299.7043 | -1.91 | 0 | 4 | 0.37 | 1 | U | -.MNLVLMGLPGAGK.G |
| 642 | 1 - 13 | 650.8608 | 1299.7069 | 1299.7043 | 2.05 | 0 | 17 | 0.019 | 1 | U | -.MNLVLMGLPGAGK.G |
| 643 | 1 - 13 | 650.8620 | 1299.7094 | 1299.7043 | 3.94 | 0 | 38 | 0.00015 | 1 | U | -.MNLVLMGLPGAGK.G |
| 644 | 1 - 13 | 658.8495 | 1315.6845 | 1315.6992 | -11.2 | 0 | 38 | 0.00021 | 1 | U | -.MNLVLMGLPGAGK.G + Oxidation (M) |
| 645 | 1 - 13 | 658.8499 | 1315.6852 | 1315.6992 | -10.7 | 0 | 49 | 1.7e-005 | 1 | U | -.MNLVLMGLPGAGK.G + Oxidation (M) |
| 646 | 1 - 13 | 658.8502 | 1315.6859 | 1315.6992 | -10.1 | 0 | 1 | 1 | 1 | U | -.MNLVLMGLPGAGK.G + Oxidation (M) |
| 647 | 1 - 13 | 658.8506 | 1315.6866 | 1315.6992 | -9.57 | 0 | 43 | 6e-005 | 1 | U | -.MNLVLMGLPGAGK.G + Oxidation (M) |
| 648 | 1 - 13 | 658.8514 | 1315.6882 | 1315.6992 | -8.34 | 0 | 59 | 1.3e-006 | 1 | U | -.MNLVLMGLPGAGK.G + Oxidation (M) |
| 649 | 1 - 13 | 658.8523 | 1315.6900 | 1315.6992 | -6.97 | 0 | 31 | 0.00095 | 1 | U | -.MNLVLMGLPGAGK.G + Oxidation (M) |
| 650 | 1 - 13 | 658.8524 | 1315.6903 | 1315.6992 | -6.78 | 0 | 15 | 0.034 | 1 | U | -.MNLVLMGLPGAGK.G + Oxidation (M) |
| 651 | 1 - 13 | 658.8528 | 1315.6911 | 1315.6992 | -6.12 | 0 | 48 | 1.8e-005 | 1 | U | -.MNLVLMGLPGAGK.G + Oxidation (M) |
| 652 | 1 - 13 | 658.8528 | 1315.6911 | 1315.6992 | -6.12 | 0 | 45 | 3.2e-005 | 1 | U | -.MNLVLMGLPGAGK.G + Oxidation (M) |
| 653 | 1 - 13 | 658.8532 | 1315.6918 | 1315.6992 | -5.59 | 0 | 28 | 0.0018 | 1 | U | -.MNLVLMGLPGAGK.G + Oxidation (M) |
| 654 | 1 - 13 | 658.8533 | 1315.6920 | 1315.6992 | -5.45 | 0 | 1 | 0.86 | 1 | U | -.MNLVLMGLPGAGK.G + Oxidation (M) |
| 655 | 1 - 13 | 658.8534 | 1315.6922 | 1315.6992 | -5.30 | 0 | 42 | 7e-005 | 1 | U | -.MNLVLMGLPGAGK.G + Oxidation (M) |
| 656 | 1 - 13 | 658.8535 | 1315.6924 | 1315.6992 | -5.15 | 0 | 30 | 0.001 | 1 | U | -.MNLVLMGLPGAGK.G + Oxidation (M) |
| 657 | 1 - 13 | 658.8537 | 1315.6929 | 1315.6992 | -4.80 | 0 | 22 | 0.0074 | 1 | U | -.MNLVLMGLPGAGK.G + Oxidation (M) |
| 658 | 1 - 13 | 658.8546 | 1315.6946 | 1315.6992 | -3.51 | 0 | 12 | 0.068 | 1 | U | -.MNLVLMGLPGAGK.G + Oxidation (M) |
| 659 | 1 - 13 | 658.8549 | 1315.6952 | 1315.6992 | -3.05 | 0 | 47 | 2.2e-005 | 1 | U | -.MNLVLMGLPGAGK.G + Oxidation (M) |
| 668 | 1 - 13 | 666.8477 | 1331.6809 | 1331.6941 | -9.94 | 0 | 20 | 0.012 | 1 | U | -.MNLVLMGLPGAGK.G + 2 Oxidation (M) |
| 669 | 1 - 13 | 666.8492 | 1331.6838 | 1331.6941 | -7.71 | 0 | 27 | 0.0027 | 1 | U | -.MNLVLMGLPGAGK.G + 2 Oxidation (M) |
| 670 | 1 - 13 | 666.8493 | 1331.6841 | 1331.6941 | -7.52 | 0 | 35 | 0.00043 | 1 | U | -.MNLVLMGLPGAGK.G + 2 Oxidation (M) |
| 671 | 1 - 13 | 666.8495 | 1331.6844 | 1331.6941 | -7.26 | 0 | 33 | 0.00061 | 1 | U | -.MNLVLMGLPGAGK.G + 2 Oxidation (M) |

| Query | Start – End | Observed | Mr(expt) | Mr(calc) | ppm | M | Score | Expect | Rank | U | Peptide |
|---|---|---|---|---|---|---|---|---|---|---|---|
| 672 | 1 – 13 | 666.8499 | 1331.6853 | 1331.6941 | -6.60 | 0 | 48 | 1.7e-005 | 1 | U | -.MNLVLMGLPGAGK.G + 2 Oxidation (M) |
| 673 | 1 – 13 | 666.8503 | 1331.6860 | 1331.6941 | -6.12 | 0 | 34 | 0.0005 | 1 | U | -.MNLVLMGLPGAGK.G + 2 Oxidation (M) |
| 674 | 1 – 13 | 666.8503 | 1331.6860 | 1331.6941 | -6.12 | 0 | 37 | 0.00022 | 1 | U | -.MNLVLMGLPGAGK.G + 2 Oxidation (M) |
| 675 | 1 – 13 | 666.8503 | 1331.6860 | 1331.6941 | -6.11 | 0 | 41 | 8.7e-005 | 1 | U | -.MNLVLMGLPGAGK.G + 2 Oxidation (M) |
| 676 | 1 – 13 | 666.8503 | 1331.6860 | 1331.6941 | -6.11 | 0 | 45 | 3.7e-005 | 1 | U | -.MNLVLMGLPGAGK.G + 2 Oxidation (M) |
| 677 | 1 – 13 | 666.8504 | 1331.6863 | 1331.6941 | -5.90 | 0 | 45 | 3.5e-005 | 1 | U | -.MNLVLMGLPGAGK.G + 2 Oxidation (M) |
| 678 | 1 – 13 | 666.8505 | 1331.6864 | 1331.6941 | -5.82 | 0 | 28 | 0.0017 | 1 | U | -.MNLVLMGLPGAGK.G + 2 Oxidation (M) |
| 679 | 1 – 13 | 666.8505 | 1331.6864 | 1331.6941 | -5.82 | 0 | 47 | 2.3e-005 | 1 | U | -.MNLVLMGLPGAGK.G + 2 Oxidation (M) |
| 680 | 1 – 13 | 666.8505 | 1331.6864 | 1331.6941 | -5.79 | 0 | 36 | 0.00026 | 1 | U | -.MNLVLMGLPGAGK.G + 2 Oxidation (M) |
| 681 | 1 – 13 | 666.8505 | 1331.6864 | 1331.6941 | -5.76 | 0 | 24 | 0.005 | 1 | U | -.MNLVLMGLPGAGK.G + 2 Oxidation (M) |
| 776 | 37 – 50 | 715.8794 | 1429.7442 | 1429.7599 | -11.0 | 1 | 3 | 0.49 | 1 | U | R.AAMKEGTPLGLQAK.Q + Oxidation (M) |
| 777 | 37 – 50 | 715.8812 | 1429.7478 | 1429.7599 | -8.45 | 1 | 4 | 0.38 | 1 | U | R.AAMKEGTPLGLQAK.Q + Oxidation (M) |
| 778 | 37 – 50 | 715.8827 | 1429.7507 | 1429.7599 | -6.38 | 1 | 1 | 0.87 | 1 | U | R.AAMKEGTPLGLQAK.Q + Oxidation (M) |
| 781 | 37 – 50 | 715.8863 | 1429.7579 | 1429.7599 | -1.35 | 1 | 12 | 0.059 | 1 | U | R.AAMKEGTPLGLQAK.Q + Oxidation (M) |
| 782 | 37 – 50 | 715.8868 | 1429.7590 | 1429.7599 | -0.63 | 1 | 31 | 0.00084 | 1 | U | R.AAMKEGTPLGLQAK.Q + Oxidation (M) |
| 783 | 37 – 50 | 715.8885 | 1429.7625 | 1429.7599 | 1.84 | 1 | 4 | 0.44 | 1 | U | R.AAMKEGTPLGLQAK.Q + Oxidation (M) |
| 337 | 41 – 50 | 507.2779 | 1012.5413 | 1012.5553 | -13.9 | 0 | 43 | 5.2e-005 | 1 | U | K.EGTPLGLQAK.Q |
| 338 | 41 – 50 | 507.2794 | 1012.5443 | 1012.5553 | -10.8 | 0 | 23 | 0.0055 | 1 | U | K.EGTPLGLQAK.Q |
| 339 | 41 – 50 | 507.2796 | 1012.5446 | 1012.5553 | -10.6 | 0 | 32 | 0.00058 | 1 | U | K.EGTPLGLQAK.Q |
| 340 | 41 – 50 | 507.2799 | 1012.5452 | 1012.5553 | -9.93 | 0 | 32 | 0.00061 | 1 | U | K.EGTPLGLQAK.Q |
| 341 | 41 – 50 | 507.2803 | 1012.5461 | 1012.5553 | -9.12 | 0 | 34 | 0.00039 | 1 | U | K.EGTPLGLQAK.Q |
| 342 | 41 – 50 | 507.2805 | 1012.5465 | 1012.5553 | -8.66 | 0 | 26 | 0.0028 | 1 | U | K.EGTPLGLQAK.Q |
| 343 | 41 – 50 | 507.2811 | 1012.5476 | 1012.5553 | -7.63 | 0 | 20 | 0.01 | 1 | U | K.EGTPLGLQAK.Q |
| 344 | 41 – 50 | 507.2820 | 1012.5494 | 1012.5553 | -5.82 | 0 | 34 | 0.00036 | 1 | U | K.EGTPLGLQAK.Q |
| 345 | 41 – 50 | 507.2830 | 1012.5513 | 1012.5553 | -3.90 | 0 | 17 | 0.021 | 1 | U | K.EGTPLGLQAK.Q |
| 346 | 41 – 50 | 507.2847 | 1012.5548 | 1012.5553 | -0.46 | 0 | 13 | 0.05 | 1 | U | K.EGTPLGLQAK.Q |
| 347 | 41 – 50 | 507.2854 | 1012.5562 | 1012.5553 | 0.90 | 0 | 21 | 0.0078 | 1 | U | K.EGTPLGLQAK.Q |
| 1346 | 51 – 69 | 726.0345 | 2175.0816 | 2175.0994 | -8.20 | 1 | 25 | 0.0039 | 1 | U | K.QYMDRGDLVPDEVTIGIVR.E |
| 1347 | 51 – 69 | 726.0349 | 2175.0829 | 2175.0994 | -7.60 | 1 | 44 | 5.2e-005 | 1 | U | K.QYMDRGDLVPDEVTIGIVR.E |
| 1348 | 51 – 69 | 726.0355 | 2175.0848 | 2175.0994 | -6.72 | 1 | 31 | 0.00095 | 1 | U | K.QYMDRGDLVPDEVTIGIVR.E |
| 1349 | 51 – 69 | 726.0367 | 2175.0883 | 2175.0994 | -5.12 | 1 | 31 | 0.00099 | 1 | U | K.QYMDRGDLVPDEVTIGIVR.E |
| 1350 | 51 – 69 | 726.0397 | 2175.0972 | 2175.0994 | -1.04 | 1 | 34 | 0.00055 | 1 | U | K.QYMDRGDLVPDEVTIGIVR.E |
| 1388 | 51 – 69 | 1096.5414 | 2191.0682 | 2191.0943 | -11.9 | 1 | 4 | 0.44 | 1 | U | K.QYMDRGDLVPDEVTIGIVR.E + Oxidation (M) |
| 1389 | 51 – 69 | 731.3644 | 2191.0714 | 2191.0943 | -10.5 | 1 | 46 | 3e-005 | 1 | U | K.QYMDRGDLVPDEVTIGIVR.E + Oxidation (M) |
| 1391 | 51 – 69 | 731.3652 | 2191.0737 | 2191.0943 | -9.41 | 1 | 47 | 2.4e-005 | 1 | U | K.QYMDRGDLVPDEVTIGIVR.E + Oxidation (M) |
| 1392 | 51 – 69 | 731.3656 | 2191.0751 | 2191.0943 | -8.77 | 1 | 21 | 0.01 | 1 | U | K.QYMDRGDLVPDEVTIGIVR.E + Oxidation (M) |
| 1393 | 51 – 69 | 731.3659 | 2191.0758 | 2191.0943 | -8.47 | 1 | 50 | 1.2e-005 | 1 | U | K.QYMDRGDLVPDEVTIGIVR.E + Oxidation (M) |
| 1394 | 51 – 69 | 731.3662 | 2191.0769 | 2191.0943 | -7.96 | 1 | 60 | 1.2e-006 | 1 | U | K.QYMDRGDLVPDEVTIGIVR.E + Oxidation (M) |
| 1395 | 51 – 69 | 731.3664 | 2191.0773 | 2191.0943 | -7.78 | 1 | 59 | 1.8e-006 | 1 | U | K.QYMDRGDLVPDEVTIGIVR.E + Oxidation (M) |
| 1396 | 51 – 69 | 731.3673 | 2191.0801 | 2191.0943 | -6.48 | 1 | 34 | 0.0005 | 1 | U | K.QYMDRGDLVPDEVTIGIVR.E + Oxidation (M) |
| 1397 | 51 – 69 | 1096.5486 | 2191.0826 | 2191.0943 | -5.33 | 1 | 16 | 0.036 | 1 | U | K.QYMDRGDLVPDEVTIGIVR.E + Oxidation (M) |
| 1398 | 51 – 69 | 731.3683 | 2191.0830 | 2191.0943 | -5.19 | 1 | 59 | 1.8e-006 | 1 | U | K.QYMDRGDLVPDEVTIGIVR.E + Oxidation (M) |
| 1399 | 51 – 69 | 731.3686 | 2191.0839 | 2191.0943 | -4.78 | 1 | 57 | 2.4e-006 | 1 | U | K.QYMDRGDLVPDEVTIGIVR.E + Oxidation (M) |
| 1400 | 51 – 69 | 731.3690 | 2191.0851 | 2191.0943 | -4.21 | 1 | 25 | 0.0039 | 1 | U | K.QYMDRGDLVPDEVTIGIVR.E + Oxidation (M) |
| 1401 | 51 – 69 | 1096.5507 | 2191.0868 | 2191.0943 | -3.41 | 1 | 11 | 0.1 | 1 | U | K.QYMDRGDLVPDEVTIGIVR.E + Oxidation (M) |
| 1402 | 51 – 69 | 731.3696 | 2191.0869 | 2191.0943 | -3.39 | 1 | 46 | 3.6e-005 | 1 | U | K.QYMDRGDLVPDEVTIGIVR.E + Oxidation (M) |
| 1403 | 51 – 69 | 1096.5530 | 2191.0914 | 2191.0943 | -1.32 | 1 | 5 | 0.4 | 1 | U | K.QYMDRGDLVPDEVTIGIVR.E + Oxidation (M) |
| 1404 | 51 – 69 | 1096.5532 | 2191.0918 | 2191.0943 | -1.13 | 1 | 16 | 0.029 | 1 | U | K.QYMDRGDLVPDEVTIGIVR.E + Oxidation (M) |
| 1405 | 51 – 69 | 1096.5569 | 2191.0992 | 2191.0943 | 2.24 | 1 | 5 | 0.37 | 1 | U | K.QYMDRGDLVPDEVTIGIVR.E + Oxidation (M) |
| 800 | 56 – 69 | 741.9045 | 1481.7944 | 1481.8090 | -9.82 | 0 | 4 | 0.43 | 1 | U | R.GDLVPDEVTIGIVR.E |
| 801 | 56 – 69 | 741.9063 | 1481.7981 | 1481.8090 | -7.36 | 0 | 21 | 0.0076 | 1 | U | R.GDLVPDEVTIGIVR.E |
| 802 | 56 – 69 | 741.9066 | 1481.7987 | 1481.8090 | -6.96 | 0 | 50 | 1.1e-005 | 1 | U | R.GDLVPDEVTIGIVR.E |
| 803 | 56 – 69 | 741.9066 | 1481.7987 | 1481.8090 | -6.96 | 0 | 62 | 6.7e-007 | 1 | U | R.GDLVPDEVTIGIVR.E |
| 804 | 56 – 69 | 741.9066 | 1481.7987 | 1481.8090 | -6.96 | 0 | 42 | 5.7e-005 | 1 | U | R.GDLVPDEVTIGIVR.E |
| 805 | 56 – 69 | 741.9071 | 1481.7997 | 1481.8090 | -6.25 | 0 | 53 | 5e-006 | 1 | U | R.GDLVPDEVTIGIVR.E |
| 806 | 56 – 69 | 741.9077 | 1481.8008 | 1481.8090 | -5.51 | 0 | 34 | 0.00041 | 1 | U | R.GDLVPDEVTIGIVR.E |
| 807 | 56 – 69 | 741.9079 | 1481.8013 | 1481.8090 | -5.19 | 0 | 56 | 2.2e-006 | 1 | U | R.GDLVPDEVTIGIVR.E |
| 808 | 56 – 69 | 741.9080 | 1481.8014 | 1481.8090 | -5.08 | 0 | 57 | 1.9e-006 | 1 | U | R.GDLVPDEVTIGIVR.E |
| 809 | 56 – 69 | 741.9082 | 1481.8018 | 1481.8090 | -4.84 | 0 | 11 | 0.081 | 1 | U | R.GDLVPDEVTIGIVR.E |
| 810 | 56 – 69 | 741.9084 | 1481.8022 | 1481.8090 | -4.58 | 0 | 26 | 0.0023 | 1 | U | R.GDLVPDEVTIGIVR.E |
| 811 | 56 – 69 | 741.9090 | 1481.8034 | 1481.8090 | -3.74 | 0 | 31 | 0.00078 | 1 | U | R.GDLVPDEVTIGIVR.E |
| 812 | 56 – 69 | 741.9094 | 1481.8042 | 1481.8090 | -3.24 | 0 | 48 | 1.7e-005 | 1 | U | R.GDLVPDEVTIGIVR.E |
| 813 | 56 – 69 | 741.9096 | 1481.8046 | 1481.8090 | -2.97 | 0 | 16 | 0.023 | 1 | U | R.GDLVPDEVTIGIVR.E |
| 814 | 56 – 69 | 741.9096 | 1481.8047 | 1481.8090 | -2.91 | 0 | 56 | 2.5e-006 | 1 | U | R.GDLVPDEVTIGIVR.E |
| 815 | 56 – 69 | 741.9096 | 1481.8047 | 1481.8090 | -2.91 | 0 | 50 | 9.2e-006 | 1 | U | R.GDLVPDEVTIGIVR.E |
| 816 | 56 – 69 | 741.9096 | 1481.8047 | 1481.8090 | -2.91 | 0 | 55 | 3e-006 | 1 | U | R.GDLVPDEVTIGIVR.E |
| 817 | 56 – 69 | 741.9099 | 1481.8053 | 1481.8090 | -2.50 | 0 | 38 | 0.00017 | 1 | U | R.GDLVPDEVTIGIVR.E |

| Query | Start - End | Observed | Mr(expt) | Mr(calc) | ppm | M | Score | Expect | Rank | U | Peptide |
|---|---|---|---|---|---|---|---|---|---|---|---|
| 818 | 56 - 69 | 741.9099 | 1481.8053 | 1481.8090 | -2.50 | 0 | 54 | 4.3e-006 | 1 | U | R.GDLVPDEVTIGIVR.E |
| 819 | 56 - 69 | 741.9099 | 1481.8053 | 1481.8090 | -2.50 | 0 | 57 | 1.9e-006 | 1 | U | R.GDLVPDEVTIGIVR.E |
| 820 | 56 - 69 | 741.9104 | 1481.8062 | 1481.8090 | -1.85 | 0 | 50 | 1.1e-005 | 1 | U | R.GDLVPDEVTIGIVR.E |
| 821 | 56 - 69 | 741.9112 | 1481.8078 | 1481.8090 | -0.77 | 0 | 19 | 0.012 | 1 | U | R.GDLVPDEVTIGIVR.E |
| 1183 | 72 - 88 | 962.9514 | 1923.8883 | 1923.9149 | -13.8 | 1 | 16 | 0.025 | 1 | U | R.LSKDDCQNGFLLDGFPR.T |
| 1184 | 72 - 88 | 962.9532 | 1923.8918 | 1923.9149 | -12.0 | 1 | 5 | 0.35 | 1 | U | R.LSKDDCQNGFLLDGFPR.T |
| 1190 | 72 - 88 | 665.9831 | 1994.9273 | 1994.9520 | -12.4 | 1 | 7 | 0.24 | 1 | U | R.LSKDDCQNGFLLDGFPR.T + Propionamide (C) |
| 1191 | 72 - 88 | 665.9850 | 1994.9333 | 1994.9520 | -9.38 | 1 | 15 | 0.036 | 1 | U | R.LSKDDCQNGFLLDGFPR.T + Propionamide (C) |
| 1192 | 72 - 88 | 665.9878 | 1994.9414 | 1994.9520 | -5.31 | 1 | 3 | 0.55 | 1 | U | R.LSKDDCQNGFLLDGFPR.T + Propionamide (C) |
| 1193 | 72 - 88 | 666.3174 | 1995.9305 | 1994.9520 | 490 | 1 | 24 | 0.0036 | 1 | U | R.LSKDDCQNGFLLDGFPR.T + Propionamide (C) |
| 1194 | 72 - 88 | 666.3187 | 1995.9342 | 1994.9520 | 492 | 1 | 13 | 0.052 | 1 | U | R.LSKDDCQNGFLLDGFPR.T + Propionamide (C) |
| 1195 | 72 - 88 | 666.3189 | 1995.9348 | 1994.9520 | 493 | 1 | 7 | 0.2 | 1 | U | R.LSKDDCQNGFLLDGFPR.T + Propionamide (C) |
| 900 | 75 - 88 | 798.8696 | 1595.7247 | 1595.7039 | 13.0 | 0 | 42 | 8.1e-005 | 1 | U | K.DDCQNGFLLDGFPR.T |
| 901 | 75 - 88 | 798.8735 | 1595.7324 | 1595.7039 | 17.9 | 0 | 33 | 0.00076 | 1 | U | K.DDCQNGFLLDGFPR.T |
| 1070 | 89 - 105 | 894.9530 | 1787.8915 | 1787.9087 | -9.64 | 0 | 82 | 5.9e-009 | 1 | U | R.TVAQAEALETMLADIGR.K |
| 1071 | 89 - 105 | 894.9530 | 1787.8915 | 1787.9087 | -9.62 | 0 | 25 | 0.0032 | 1 | U | R.TVAQAEALETMLADIGR.K |
| 1072 | 89 - 105 | 894.9543 | 1787.8941 | 1787.9087 | -8.18 | 0 | 63 | 5.5e-007 | 1 | U | R.TVAQAEALETMLADIGR.K |
| 1073 | 89 - 105 | 894.9547 | 1787.8949 | 1787.9087 | -7.73 | 0 | 20 | 0.0098 | 1 | U | R.TVAQAEALETMLADIGR.K |
| 1074 | 89 - 105 | 894.9553 | 1787.8961 | 1787.9087 | -7.04 | 0 | 66 | 2.4e-007 | 1 | U | R.TVAQAEALETMLADIGR.K |
| 1075 | 89 - 105 | 894.9562 | 1787.8979 | 1787.9087 | -6.03 | 0 | 66 | 2.5e-007 | 1 | U | R.TVAQAEALETMLADIGR.K |
| 1076 | 89 - 105 | 894.9566 | 1787.8987 | 1787.9087 | -5.60 | 0 | 66 | 2.7e-007 | 1 | U | R.TVAQAEALETMLADIGR.K |
| 1077 | 89 - 105 | 894.9569 | 1787.8992 | 1787.9087 | -5.33 | 0 | 76 | 2.8e-008 | 1 | U | R.TVAQAEALETMLADIGR.K |
| 1078 | 89 - 105 | 596.9739 | 1787.9000 | 1787.9087 | -4.87 | 0 | 31 | 0.00079 | 1 | U | R.TVAQAEALETMLADIGR.K |
| 1079 | 89 - 105 | 596.9740 | 1787.9002 | 1787.9087 | -4.75 | 0 | 18 | 0.015 | 1 | U | R.TVAQAEALETMLADIGR.K |
| 1080 | 89 - 105 | 596.9742 | 1787.9008 | 1787.9087 | -4.43 | 0 | 30 | 0.00099 | 1 | U | R.TVAQAEALETMLADIGR.K |
| 1081 | 89 - 105 | 894.9584 | 1787.9022 | 1787.9087 | -3.63 | 0 | 66 | 2.8e-007 | 1 | U | R.TVAQAEALETMLADIGR.K |
| 1082 | 89 - 105 | 596.9757 | 1787.9053 | 1787.9087 | -1.91 | 0 | 26 | 0.0026 | 1 | U | R.TVAQAEALETMLADIGR.K |
| 1083 | 89 - 105 | 894.9612 | 1787.9078 | 1787.9087 | -0.51 | 0 | 65 | 3.3e-007 | 1 | U | R.TVAQAEALETMLADIGR.K |
| 1084 | 89 - 105 | 596.9771 | 1787.9094 | 1787.9087 | 0.37 | 0 | 20 | 0.01 | 1 | U | R.TVAQAEALETMLADIGR.K |
| 1085 | 89 - 105 | 894.9646 | 1787.9145 | 1787.9087 | 3.26 | 0 | 34 | 0.00037 | 1 | U | R.TVAQAEALETMLADIGR.K |
| 1086 | 89 - 105 | 602.2998 | 1803.8775 | 1803.9036 | -14.5 | 0 | 38 | 0.00014 | 1 | U | R.TVAQAEALETMLADIGR.K + Oxidation (M) |
| 1087 | 89 - 105 | 602.3015 | 1803.8827 | 1803.9036 | -11.6 | 0 | 39 | 0.00012 | 1 | U | R.TVAQAEALETMLADIGR.K + Oxidation (M) |
| 1088 | 89 - 105 | 602.3018 | 1803.8837 | 1803.9036 | -11.0 | 0 | 1 | 0.87 | 1 | U | R.TVAQAEALETMLADIGR.K + Oxidation (M) |
| 1089 | 89 - 105 | 602.3025 | 1803.8858 | 1803.9036 | -9.89 | 0 | 56 | 2.4e-006 | 1 | U | R.TVAQAEALETMLADIGR.K + Oxidation (M) |
| 1090 | 89 - 105 | 602.3026 | 1803.8859 | 1803.9036 | -9.84 | 0 | 16 | 0.024 | 1 | U | R.TVAQAEALETMLADIGR.K + Oxidation (M) |
| 1091 | 89 - 105 | 602.3037 | 1803.8893 | 1803.9036 | -7.93 | 0 | 35 | 0.00029 | 1 | U | R.TVAQAEALETMLADIGR.K + Oxidation (M) |
| 1092 | 89 - 105 | 602.3043 | 1803.8912 | 1803.9036 | -6.90 | 0 | 35 | 0.00034 | 1 | U | R.TVAQAEALETMLADIGR.K + Oxidation (M) |
| 1093 | 89 - 105 | 902.9530 | 1803.8914 | 1803.9036 | -6.80 | 0 | 74 | 3.9e-008 | 1 | U | R.TVAQAEALETMLADIGR.K + Oxidation (M) |
| 1094 | 89 - 105 | 902.9539 | 1803.8933 | 1803.9036 | -5.72 | 0 | 59 | 1.1e-006 | 1 | U | R.TVAQAEALETMLADIGR.K + Oxidation (M) |
| 1095 | 89 - 105 | 902.9542 | 1803.8938 | 1803.9036 | -5.43 | 0 | 18 | 0.015 | 1 | U | R.TVAQAEALETMLADIGR.K + Oxidation (M) |
| 1096 | 89 - 105 | 602.3054 | 1803.8943 | 1803.9036 | -5.17 | 0 | 35 | 0.0003 | 1 | U | R.TVAQAEALETMLADIGR.K + Oxidation (M) |
| 1097 | 89 - 105 | 902.9547 | 1803.8948 | 1803.9036 | -4.87 | 0 | 81 | 7.9e-009 | 1 | U | R.TVAQAEALETMLADIGR.K + Oxidation (M) |
| 1098 | 89 - 105 | 902.9550 | 1803.8955 | 1803.9036 | -4.50 | 0 | 38 | 0.00016 | 1 | U | R.TVAQAEALETMLADIGR.K + Oxidation (M) |
| 1099 | 89 - 105 | 602.3058 | 1803.8957 | 1803.9036 | -4.41 | 0 | 32 | 0.00058 | 1 | U | R.TVAQAEALETMLADIGR.K + Oxidation (M) |
| 1100 | 89 - 105 | 902.9556 | 1803.8967 | 1803.9036 | -3.87 | 0 | 47 | 2.1e-005 | 1 | U | R.TVAQAEALETMLADIGR.K + Oxidation (M) |
| 1101 | 89 - 105 | 902.9561 | 1803.8976 | 1803.9036 | -3.32 | 0 | 86 | 2.2e-009 | 1 | U | R.TVAQAEALETMLADIGR.K + Oxidation (M) |
| 1102 | 89 - 105 | 902.9562 | 1803.8978 | 1803.9036 | -3.21 | 0 | 77 | 1.8e-008 | 1 | U | R.TVAQAEALETMLADIGR.K + Oxidation (M) |
| 1103 | 89 - 105 | 902.9562 | 1803.8979 | 1803.9036 | -3.18 | 0 | 37 | 0.0002 | 1 | U | R.TVAQAEALETMLADIGR.K + Oxidation (M) |
| 1104 | 89 - 105 | 902.9563 | 1803.8980 | 1803.9036 | -3.10 | 0 | 65 | 3e-007 | 1 | U | R.TVAQAEALETMLADIGR.K + Oxidation (M) |
| 1105 | 89 - 105 | 902.9569 | 1803.8993 | 1803.9036 | -2.42 | 0 | 34 | 0.0004 | 1 | U | R.TVAQAEALETMLADIGR.K + Oxidation (M) |
| 1106 | 89 - 105 | 602.3071 | 1803.8996 | 1803.9036 | -2.24 | 0 | 45 | 3.3e-005 | 1 | U | R.TVAQAEALETMLADIGR.K + Oxidation (M) |
| 1107 | 89 - 105 | 902.9583 | 1803.9020 | 1803.9036 | -0.93 | 0 | 35 | 0.00034 | 1 | U | R.TVAQAEALETMLADIGR.K + Oxidation (M) |
| 1108 | 89 - 105 | 903.4531 | 1804.8917 | 1803.9036 | 548 | 0 | 14 | 0.05 | 1 | U | R.TVAQAEALETMLADIGR.K + Oxidation (M) |
| 1109 | 89 - 105 | 903.4532 | 1804.8919 | 1803.9036 | 548 | 0 | 5 | 0.34 | 1 | U | R.TVAQAEALETMLADIGR.K + Oxidation (M) |
| 724 | 106 - 116 | 685.8764 | 1369.7382 | 1369.7718 | -24.5 | 1 | 0 | 0.91 | 1 | U | R.KLDYVIHIDVR.Q |
| 725 | 106 - 116 | 685.8771 | 1369.7395 | 1369.7718 | -23.5 | 1 | 9 | 0.14 | 1 | U | R.KLDYVIHIDVR.Q |
| 727 | 106 - 116 | 457.5910 | 1369.7513 | 1369.7718 | -15.0 | 1 | 20 | 0.01 | 1 | U | R.KLDYVIHIDVR.Q |
| 728 | 106 - 116 | 685.8835 | 1369.7524 | 1369.7718 | -14.2 | 1 | 24 | 0.0043 | 1 | U | R.KLDYVIHIDVR.Q |
| 729 | 106 - 116 | 457.5915 | 1369.7528 | 1369.7718 | -13.9 | 1 | 5 | 0.33 | 1 | U | R.KLDYVIHIDVR.Q |
| 730 | 106 - 116 | 457.5916 | 1369.7529 | 1369.7718 | -13.8 | 1 | 19 | 0.012 | 1 | U | R.KLDYVIHIDVR.Q |
| 731 | 106 - 116 | 457.5919 | 1369.7538 | 1369.7718 | -13.1 | 1 | 27 | 0.0019 | 1 | U | R.KLDYVIHIDVR.Q |
| 732 | 106 - 116 | 457.5922 | 1369.7547 | 1369.7718 | -12.5 | 1 | 2 | 0.64 | 1 | U | R.KLDYVIHIDVR.Q |
| 734 | 106 - 116 | 457.5925 | 1369.7556 | 1369.7718 | -11.8 | 1 | 27 | 0.0021 | 1 | U | R.KLDYVIHIDVR.Q |
| 735 | 106 - 116 | 457.5926 | 1369.7560 | 1369.7718 | -11.5 | 1 | 16 | 0.028 | 1 | U | R.KLDYVIHIDVR.Q |
| 737 | 106 - 116 | 457.5931 | 1369.7576 | 1369.7718 | -10.4 | 1 | 33 | 0.00045 | 1 | U | R.KLDYVIHIDVR.Q |
| 738 | 106 - 116 | 457.5932 | 1369.7578 | 1369.7718 | -10.2 | 1 | 7 | 0.21 | 1 | U | R.KLDYVIHIDVR.Q |
| 739 | 106 - 116 | 457.5935 | 1369.7586 | 1369.7718 | -9.61 | 1 | 19 | 0.013 | 1 | U | R.KLDYVIHIDVR.Q |

| Query | Start – End | Observed | Mr(expt) | Mr(calc) | ppm | M | Score | Expect | Rank | U | Peptide |
|---|---|---|---|---|---|---|---|---|---|---|---|
| 740 | 106 – 116 | 457.5936 | 1369.7590 | 1369.7718 | -9.30 | 1 | 17 | 0.021 | 1 | U | R.KLDYVIHIDVR.Q |
| 741 | 106 – 116 | 685.8869 | 1369.7593 | 1369.7718 | -9.10 | 1 | 23 | 0.0051 | 1 | U | R.KLDYVIHIDVR.Q |
| 742 | 106 – 116 | 685.8870 | 1369.7594 | 1369.7718 | -9.06 | 1 | 4 | 0.36 | 1 | U | R.KLDYVIHIDVR.Q |
| 744 | 106 – 116 | 457.5938 | 1369.7596 | 1369.7718 | -8.89 | 1 | 16 | 0.024 | 1 | U | R.KLDYVIHIDVR.Q |
| 745 | 106 – 116 | 685.8872 | 1369.7599 | 1369.7718 | -8.65 | 1 | 24 | 0.0044 | 1 | U | R.KLDYVIHIDVR.Q |
| 746 | 106 – 116 | 457.5940 | 1369.7601 | 1369.7718 | -8.54 | 1 | 11 | 0.084 | 1 | U | R.KLDYVIHIDVR.Q |
| 747 | 106 – 116 | 685.8875 | 1369.7604 | 1369.7718 | -8.28 | 1 | 41 | 8.4e-005 | 1 | U | R.KLDYVIHIDVR.Q |
| 748 | 106 – 116 | 685.8876 | 1369.7607 | 1369.7718 | -8.09 | 1 | 40 | 0.00012 | 1 | U | R.KLDYVIHIDVR.Q |
| 749 | 106 – 116 | 685.8885 | 1369.7625 | 1369.7718 | -6.75 | 1 | 27 | 0.0019 | 1 | U | R.KLDYVIHIDVR.Q |
| 750 | 106 – 116 | 457.5949 | 1369.7629 | 1369.7718 | -6.50 | 1 | 25 | 0.003 | 1 | U | R.KLDYVIHIDVR.Q |
| 751 | 106 – 116 | 685.8889 | 1369.7632 | 1369.7718 | -6.24 | 1 | 5 | 0.36 | 1 | U | R.KLDYVIHIDVR.Q |
| 752 | 106 – 116 | 457.5951 | 1369.7634 | 1369.7718 | -6.13 | 1 | 10 | 0.11 | 1 | U | R.KLDYVIHIDVR.Q |
| 753 | 106 – 116 | 685.8891 | 1369.7636 | 1369.7718 | -5.99 | 1 | 29 | 0.0012 | 1 | U | R.KLDYVIHIDVR.Q |
| 754 | 106 – 116 | 685.8901 | 1369.7657 | 1369.7718 | -4.41 | 1 | 32 | 0.00073 | 1 | U | R.KLDYVIHIDVR.Q |
| 755 | 106 – 116 | 685.8914 | 1369.7683 | 1369.7718 | -2.56 | 1 | 29 | 0.0014 | 1 | U | R.KLDYVIHIDVR.Q |
| 756 | 106 – 116 | 457.5972 | 1369.7699 | 1369.7718 | -1.35 | 1 | 15 | 0.03 | 1 | U | R.KLDYVIHIDVR.Q |
| 757 | 106 – 116 | 685.8923 | 1369.7700 | 1369.7718 | -1.29 | 1 | 49 | 1.4e-005 | 1 | U | R.KLDYVIHIDVR.Q |
| 310 | 117 – 123 | 453.7160 | 905.4175 | 905.4277 | -11.2 | 0 | 19 | 0.014 | 1 | U | R.QDVLMER.L + Oxidation (M) |
| 311 | 117 – 123 | 453.7163 | 905.4180 | 905.4277 | -10.7 | 0 | 8 | 0.17 | 1 | U | R.QDVLMER.L + Oxidation (M) |
| 312 | 117 – 123 | 453.7164 | 905.4182 | 905.4277 | -10.4 | 0 | 29 | 0.0012 | 1 | U | R.QDVLMER.L + Oxidation (M) |
| 313 | 117 – 123 | 453.7173 | 905.4201 | 905.4277 | -8.38 | 0 | 12 | 0.058 | 1 | U | R.QDVLMER.L + Oxidation (M) |
| 314 | 117 – 123 | 453.7177 | 905.4209 | 905.4277 | -7.43 | 0 | 14 | 0.039 | 1 | U | R.QDVLMER.L + Oxidation (M) |
| 315 | 117 – 123 | 453.7184 | 905.4223 | 905.4277 | -5.95 | 0 | 18 | 0.016 | 1 | U | R.QDVLMER.L + Oxidation (M) |
| 317 | 117 – 123 | 453.7192 | 905.4238 | 905.4277 | -4.21 | 0 | 8 | 0.16 | 1 | U | R.QDVLMER.L + Oxidation (M) |
| 318 | 117 – 123 | 453.7193 | 905.4241 | 905.4277 | -3.92 | 0 | 16 | 0.026 | 1 | U | R.QDVLMER.L + Oxidation (M) |
| 319 | 117 – 123 | 906.4321 | 905.4248 | 905.4277 | -3.12 | 0 | 10 | 0.095 | 1 | U | R.QDVLMER.L + Oxidation (M) |
| 547 | 161 – 171 | 588.2566 | 1174.4987 | 1174.5214 | -19.3 | 0 | 4 | 0.44 | 1 | U | R.ADDNEATVANR.L |
| 548 | 161 – 171 | 588.2566 | 1174.4987 | 1174.5214 | -19.3 | 0 | 17 | 0.019 | 1 | U | R.ADDNEATVANR.L |
| 549 | 161 – 171 | 588.2582 | 1174.5018 | 1174.5214 | -16.7 | 0 | 16 | 0.026 | 1 | U | R.ADDNEATVANR.L |
| 550 | 161 – 171 | 588.2589 | 1174.5032 | 1174.5214 | -15.5 | 0 | 10 | 0.11 | 1 | U | R.ADDNEATVANR.L |
| 551 | 161 – 171 | 588.2593 | 1174.5041 | 1174.5214 | -14.7 | 0 | 10 | 0.1 | 1 | U | R.ADDNEATVANR.L |
| 552 | 161 – 171 | 588.2602 | 1174.5059 | 1174.5214 | -13.2 | 0 | 17 | 0.02 | 1 | U | R.ADDNEATVANR.L |
| 553 | 161 – 171 | 588.2607 | 1174.5067 | 1174.5214 | -12.5 | 0 | 44 | 4.1e-005 | 1 | U | R.ADDNEATVANR.L |
| 554 | 161 – 171 | 588.2609 | 1174.5071 | 1174.5214 | -12.2 | 0 | 39 | 0.00013 | 1 | U | R.ADDNEATVANR.L |
| 555 | 161 – 171 | 588.2609 | 1174.5073 | 1174.5214 | -12.0 | 0 | 15 | 0.033 | 1 | U | R.ADDNEATVANR.L |
| 556 | 161 – 171 | 588.2611 | 1174.5076 | 1174.5214 | -11.8 | 0 | 27 | 0.0021 | 1 | U | R.ADDNEATVANR.L |
| 557 | 161 – 171 | 588.2613 | 1174.5080 | 1174.5214 | -11.4 | 0 | 24 | 0.0038 | 1 | U | R.ADDNEATVANR.L |
| 558 | 161 – 171 | 588.2617 | 1174.5089 | 1174.5214 | -10.7 | 0 | 18 | 0.015 | 1 | U | R.ADDNEATVANR.L |
| 559 | 161 – 171 | 588.2621 | 1174.5097 | 1174.5214 | -9.97 | 0 | 33 | 0.00045 | 1 | U | R.ADDNEATVANR.L |
| 560 | 161 – 171 | 588.2622 | 1174.5098 | 1174.5214 | -9.91 | 0 | 50 | 1e-005 | 1 | U | R.ADDNEATVANR.L |
| 561 | 161 – 171 | 588.2624 | 1174.5102 | 1174.5214 | -9.58 | 0 | 15 | 0.033 | 1 | U | R.ADDNEATVANR.L |
| 562 | 161 – 171 | 588.2629 | 1174.5112 | 1174.5214 | -8.68 | 0 | 33 | 0.00053 | 1 | U | R.ADDNEATVANR.L |
| 563 | 161 – 171 | 588.2631 | 1174.5117 | 1174.5214 | -8.27 | 0 | 41 | 8.5e-005 | 1 | U | R.ADDNEATVANR.L |
| 564 | 161 – 171 | 588.2632 | 1174.5118 | 1174.5214 | -8.17 | 0 | 12 | 0.061 | 1 | U | R.ADDNEATVANR.L |
| 565 | 161 – 171 | 588.2632 | 1174.5119 | 1174.5214 | -8.08 | 0 | 29 | 0.0013 | 1 | U | R.ADDNEATVANR.L |
| 566 | 161 – 171 | 588.2633 | 1174.5120 | 1174.5214 | -8.05 | 0 | 34 | 0.00038 | 1 | U | R.ADDNEATVANR.L |
| 567 | 161 – 171 | 588.2635 | 1174.5125 | 1174.5214 | -7.62 | 0 | 23 | 0.0052 | 1 | U | R.ADDNEATVANR.L |
| 568 | 161 – 171 | 588.2637 | 1174.5128 | 1174.5214 | -7.35 | 0 | 37 | 0.00018 | 1 | U | R.ADDNEATVANR.L |
| 569 | 161 – 171 | 588.2637 | 1174.5129 | 1174.5214 | -7.23 | 0 | 18 | 0.015 | 1 | U | R.ADDNEATVANR.L |
| 570 | 161 – 171 | 588.2638 | 1174.5130 | 1174.5214 | -7.18 | 0 | 33 | 0.00047 | 1 | U | R.ADDNEATVANR.L |
| 571 | 161 – 171 | 588.2641 | 1174.5137 | 1174.5214 | -6.59 | 0 | 28 | 0.0017 | 1 | U | R.ADDNEATVANR.L |
| 572 | 161 – 171 | 588.2642 | 1174.5138 | 1174.5214 | -6.52 | 0 | 21 | 0.0072 | 1 | U | R.ADDNEATVANR.L |
| 573 | 161 – 171 | 588.2645 | 1174.5144 | 1174.5214 | -5.99 | 0 | 37 | 0.00019 | 1 | U | R.ADDNEATVANR.L |
| 574 | 161 – 171 | 588.2645 | 1174.5144 | 1174.5214 | -5.96 | 0 | 19 | 0.013 | 1 | U | R.ADDNEATVANR.L |
| 575 | 161 – 171 | 588.2648 | 1174.5150 | 1174.5214 | -5.44 | 0 | 39 | 0.00013 | 1 | U | R.ADDNEATVANR.L |
| 576 | 161 – 171 | 588.2648 | 1174.5151 | 1174.5214 | -5.41 | 0 | 3 | 0.55 | 1 | U | R.ADDNEATVANR.L |
| 577 | 161 – 171 | 588.2649 | 1174.5152 | 1174.5214 | -5.33 | 0 | 34 | 0.00042 | 1 | U | R.ADDNEATVANR.L |
| 579 | 161 – 171 | 588.2650 | 1174.5154 | 1174.5214 | -5.12 | 0 | 37 | 0.00022 | 1 | U | R.ADDNEATVANR.L |
| 580 | 161 – 171 | 588.2650 | 1174.5154 | 1174.5214 | -5.12 | 0 | 20 | 0.01 | 1 | U | R.ADDNEATVANR.L |
| 581 | 161 – 171 | 588.2652 | 1174.5159 | 1174.5214 | -4.68 | 0 | 32 | 0.00063 | 1 | U | R.ADDNEATVANR.L |
| 582 | 161 – 171 | 588.2654 | 1174.5161 | 1174.5214 | -4.49 | 0 | 33 | 0.00054 | 1 | U | R.ADDNEATVANR.L |
| 583 | 161 – 171 | 588.2658 | 1174.5171 | 1174.5214 | -3.66 | 0 | 33 | 0.00055 | 1 | U | R.ADDNEATVANR.L |
| 584 | 161 – 171 | 588.2662 | 1174.5178 | 1174.5214 | -3.08 | 0 | 38 | 0.00015 | 1 | U | R.ADDNEATVANR.L |
| 585 | 161 – 171 | 588.2663 | 1174.5181 | 1174.5214 | -2.82 | 0 | 11 | 0.083 | 1 | U | R.ADDNEATVANR.L |
| 586 | 161 – 171 | 588.2677 | 1174.5208 | 1174.5214 | -0.52 | 0 | 33 | 0.00051 | 1 | U | R.ADDNEATVANR.L |
| 1115 | 161 – 177 | 630.6315 | 1888.8726 | 1888.8949 | -11.8 | 1 | 23 | 0.0058 | 1 | U | R.ADDNEATVANRLEVNMK.Q |
| 1117 | 161 – 177 | 630.6336 | 1888.8789 | 1888.8949 | -8.42 | 1 | 43 | 5.3e-005 | 1 | U | R.ADDNEATVANRLEVNMK.Q |

| Query | Start - End | Observed | Mr(expt) | Mr(calc) | ppm | M | Score | Expect | Rank | U | Peptide |
|---|---|---|---|---|---|---|---|---|---|---|---|
| 1118 | 161 - 177 | 630.6346 | 1888.8819 | 1888.8949 | -6.87 | 1 | 20 | 0.011 | 1 | U | R.ADDNEATVANRLEVNMK.Q |
| 1119 | 161 - 177 | 630.6349 | 1888.8829 | 1888.8949 | -6.34 | 1 | 17 | 0.024 | 1 | U | R.ADDNEATVANRLEVNMK.Q |
| 1120 | 161 - 177 | 630.6361 | 1888.8865 | 1888.8949 | -4.42 | 1 | 3 | 0.85 | 1 | U | R.ADDNEATVANRLEVNMK.Q |
| 1137 | 161 - 177 | 635.9638 | 1904.8695 | 1904.8898 | -10.6 | 1 | 50 | 1.2e-005 | 1 | U | R.ADDNEATVANRLEVNMK.Q + Oxidation (M) |
| 1138 | 161 - 177 | 635.9639 | 1904.8698 | 1904.8898 | -10.5 | 1 | 14 | 0.044 | 1 | U | R.ADDNEATVANRLEVNMK.Q + Oxidation (M) |
| 1139 | 161 - 177 | 635.9640 | 1904.8703 | 1904.8898 | -10.2 | 1 | 35 | 0.00033 | 1 | U | R.ADDNEATVANRLEVNMK.Q + Oxidation (M) |
| 1140 | 161 - 177 | 635.9641 | 1904.8704 | 1904.8898 | -10.2 | 1 | 1 | 0.91 | 1 | U | R.ADDNEATVANRLEVNMK.Q + Oxidation (M) |
| 1141 | 161 - 177 | 635.9643 | 1904.8710 | 1904.8898 | -9.83 | 1 | 20 | 0.011 | 1 | U | R.ADDNEATVANRLEVNMK.Q + Oxidation (M) |
| 1142 | 161 - 177 | 635.9647 | 1904.8721 | 1904.8898 | -9.27 | 1 | 33 | 0.00051 | 1 | U | R.ADDNEATVANRLEVNMK.Q + Oxidation (M) |
| 1143 | 161 - 177 | 635.9647 | 1904.8724 | 1904.8898 | -9.14 | 1 | 26 | 0.0028 | 1 | U | R.ADDNEATVANRLEVNMK.Q + Oxidation (M) |
| 1144 | 161 - 177 | 635.9647 | 1904.8724 | 1904.8898 | -9.11 | 1 | 59 | 1.3e-006 | 1 | U | R.ADDNEATVANRLEVNMK.Q + Oxidation (M) |
| 1145 | 161 - 177 | 635.9652 | 1904.8737 | 1904.8898 | -8.41 | 1 | 35 | 0.00033 | 1 | U | R.ADDNEATVANRLEVNMK.Q + Oxidation (M) |
| 1146 | 161 - 177 | 635.9655 | 1904.8748 | 1904.8898 | -7.88 | 1 | 28 | 0.0016 | 1 | U | R.ADDNEATVANRLEVNMK.Q + Oxidation (M) |
| 1148 | 161 - 177 | 953.4453 | 1904.8761 | 1904.8898 | -7.18 | 1 | 17 | 0.02 | 1 | U | R.ADDNEATVANRLEVNMK.Q + Oxidation (M) |
| 1149 | 161 - 177 | 953.4468 | 1904.8791 | 1904.8898 | -5.61 | 1 | 7 | 0.23 | 1 | U | R.ADDNEATVANRLEVNMK.Q + Oxidation (M) |
| 1150 | 161 - 177 | 635.9670 | 1904.8792 | 1904.8898 | -5.55 | 1 | 39 | 0.00016 | 1 | U | R.ADDNEATVANRLEVNMK.Q + Oxidation (M) |
| 1151 | 161 - 177 | 953.4478 | 1904.8810 | 1904.8898 | -4.61 | 1 | 17 | 0.025 | 1 | U | R.ADDNEATVANRLEVNMK.Q + Oxidation (M) |
| 1152 | 161 - 177 | 953.4485 | 1904.8824 | 1904.8898 | -3.84 | 1 | 4 | 0.5 | 1 | U | R.ADDNEATVANRLEVNMK.Q + Oxidation (M) |
| 1153 | 161 - 177 | 953.4489 | 1904.8833 | 1904.8898 | -3.38 | 1 | 14 | 0.045 | 1 | U | R.ADDNEATVANRLEVNMK.Q + Oxidation (M) |
| 1154 | 161 - 177 | 953.4507 | 1904.8868 | 1904.8898 | -1.53 | 1 | 17 | 0.025 | 1 | U | R.ADDNEATVANRLEVNMK.Q + Oxidation (M) |
| 1155 | 161 - 177 | 953.4545 | 1904.8945 | 1904.8898 | 2.48 | 1 | 25 | 0.0033 | 1 | U | R.ADDNEATVANRLEVNMK.Q + Oxidation (M) |
| 1165 | 161 - 177 | 636.2948 | 1905.8626 | 1904.8898 | 511 | 1 | 2 | 0.64 | 1 | U | R.ADDNEATVANRLEVNMK.Q + Oxidation (M) |
| 866 | 178 - 189 | 514.5860 | 1540.7361 | 1540.7596 | -15.2 | 1 | 23 | 0.0094 | 1 | U | K.QMKPLVDFYEQK.G + Oxidation (M) |
| 867 | 178 - 189 | 514.5861 | 1540.7365 | 1540.7596 | -14.9 | 1 | 13 | 0.096 | 1 | U | K.QMKPLVDFYEQK.G + Oxidation (M) |
| 868 | 178 - 189 | 514.5865 | 1540.7376 | 1540.7596 | -14.2 | 1 | 17 | 0.041 | 1 | U | K.QMKPLVDFYEQK.G + Oxidation (M) |
| 872 | 178 - 189 | 514.5889 | 1540.7448 | 1540.7596 | -9.59 | 1 | 13 | 0.098 | 1 | U | K.QMKPLVDFYEQK.G + Oxidation (M) |
| 873 | 178 - 189 | 514.5895 | 1540.7468 | 1540.7596 | -8.28 | 1 | 7 | 0.42 | 1 | U | K.QMKPLVDFYEQK.G + Oxidation (M) |
| 875 | 178 - 189 | 771.3820 | 1540.7494 | 1540.7596 | -6.56 | 1 | 25 | 0.0068 | 1 | U | K.QMKPLVDFYEQK.G + Oxidation (M) |
| 876 | 178 - 189 | 514.5904 | 1540.7495 | 1540.7596 | -6.53 | 1 | 19 | 0.026 | 1 | U | K.QMKPLVDFYEQK.G + Oxidation (M) |
| 877 | 178 - 189 | 514.5907 | 1540.7504 | 1540.7596 | -5.93 | 1 | 15 | 0.069 | 1 | U | K.QMKPLVDFYEQK.G + Oxidation (M) |
| 878 | 178 - 189 | 771.3825 | 1540.7505 | 1540.7596 | -5.90 | 1 | 33 | 0.001 | 1 | U | K.QMKPLVDFYEQK.G + Oxidation (M) |
| 879 | 178 - 189 | 771.3833 | 1540.7521 | 1540.7596 | -4.86 | 1 | 23 | 0.0095 | 1 | U | K.QMKPLVDFYEQK.G + Oxidation (M) |
| 880 | 178 - 189 | 771.3834 | 1540.7522 | 1540.7596 | -4.79 | 1 | 31 | 0.0013 | 1 | U | K.QMKPLVDFYEQK.G + Oxidation (M) |
| 881 | 178 - 189 | 771.3843 | 1540.7540 | 1540.7596 | -3.61 | 1 | 35 | 0.00053 | 1 | U | K.QMKPLVDFYEQK.G + Oxidation (M) |
| 882 | 178 - 189 | 771.3848 | 1540.7551 | 1540.7596 | -2.88 | 1 | 25 | 0.0053 | 1 | U | K.QMKPLVDFYEQK.G + Oxidation (M) |
| 883 | 178 - 189 | 771.3852 | 1540.7559 | 1540.7596 | -2.38 | 1 | 44 | 5.3e-005 | 1 | U | K.QMKPLVDFYEQK.G + Oxidation (M) |
| 884 | 178 - 189 | 771.3864 | 1540.7582 | 1540.7596 | -0.89 | 1 | 18 | 0.022 | 1 | U | K.QMKPLVDFYEQK.G + Oxidation (M) |
| 885 | 178 - 189 | 771.3873 | 1540.7601 | 1540.7596 | 0.37 | 1 | 2 | 0.85 | 1 | U | K.QMKPLVDFYEQK.G + Oxidation (M) |
| 886 | 178 - 189 | 771.3889 | 1540.7633 | 1540.7596 | 2.41 | 1 | 14 | 0.048 | 1 | U | K.QMKPLVDFYEQK.G + Oxidation (M) |
| 599 | 194 - 203 | 589.2545 | 1176.4945 | 1176.5081 | -11.6 | 0 | 2 | 0.62 | 1 | U | R.NINGEQDMEK.V |
| 600 | 194 - 203 | 589.2570 | 1176.4994 | 1176.5081 | -7.35 | 0 | 22 | 0.0064 | 1 | U | R.NINGEQDMEK.V |
| 601 | 194 - 203 | 589.2597 | 1176.5049 | 1176.5081 | -2.71 | 0 | 4 | 0.37 | 1 | U | R.NINGEQDMEK.V |
| 607 | 194 - 203 | 597.2489 | 1192.4833 | 1192.5030 | -16.5 | 0 | 6 | 0.24 | 1 | U | R.NINGEQDMEK.V + Oxidation (M) |
| 608 | 194 - 203 | 597.2489 | 1192.4833 | 1192.5030 | -16.5 | 0 | 7 | 0.19 | 1 | U | R.NINGEQDMEK.V + Oxidation (M) |
| 610 | 194 - 203 | 597.2505 | 1192.4864 | 1192.5030 | -13.9 | 0 | 3 | 0.47 | 1 | U | R.NINGEQDMEK.V + Oxidation (M) |
| 611 | 194 - 203 | 597.2516 | 1192.4886 | 1192.5030 | -12.0 | 0 | 1 | 0.75 | 1 | U | R.NINGEQDMEK.V + Oxidation (M) |
| 613 | 194 - 203 | 597.2534 | 1192.4922 | 1192.5030 | -9.08 | 0 | 10 | 0.089 | 1 | U | R.NINGEQDMEK.V + Oxidation (M) |
| 616 | 194 - 203 | 597.2540 | 1192.4934 | 1192.5030 | -8.02 | 0 | 9 | 0.13 | 1 | U | R.NINGEQDMEK.V + Oxidation (M) |
| 619 | 194 - 203 | 597.7497 | 1193.4849 | 1192.5030 | 823 | 0 | 22 | 0.0059 | 1 | U | R.NINGEQDMEK.V + Oxidation (M) |
| 620 | 194 - 203 | 597.7522 | 1193.4897 | 1192.5030 | 827 | 0 | 3 | 0.49 | 1 | U | R.NINGEQDMEK.V + Oxidation (M) |
| 1125 | 194 - 209 | 632.2972 | 1893.8699 | 1893.8891 | -10.1 | 1 | 0 | 0.97 | 1 | U | R.NINGEQDMEKVFADIR.E + Oxidation (M) |
| 1126 | 194 - 209 | 632.2974 | 1893.8703 | 1893.8891 | -9.93 | 1 | 5 | 0.35 | 1 | U | R.NINGEQDMEKVFADIR.E + Oxidation (M) |
| 1127 | 194 - 209 | 632.2978 | 1893.8716 | 1893.8891 | -9.23 | 1 | 3 | 0.52 | 1 | U | R.NINGEQDMEKVFADIR.E + Oxidation (M) |
| 1128 | 194 - 209 | 632.2989 | 1893.8749 | 1893.8891 | -7.47 | 1 | 1 | 0.73 | 1 | U | R.NINGEQDMEKVFADIR.E + Oxidation (M) |
| 1129 | 194 - 209 | 632.2989 | 1893.8750 | 1893.8891 | -7.44 | 1 | 4 | 0.37 | 1 | U | R.NINGEQDMEKVFADIR.E + Oxidation (M) |
| 1132 | 194 - 209 | 632.6301 | 1894.8684 | 1893.8891 | 517 | 1 | 10 | 0.098 | 1 | U | R.NINGEQDMEKVFADIR.E + Oxidation (M) |
| 1133 | 194 - 209 | 632.6305 | 1894.8696 | 1893.8891 | 518 | 1 | 2 | 0.57 | 1 | U | R.NINGEQDMEKVFADIR.E + Oxidation (M) |
| 101 | 210 - 217 | 414.7466 | 827.4785 | 827.4865 | -9.59 | 0 | 22 | 0.0061 | 1 | U | R.ELLGGLAR.- |
| 103 | 210 - 217 | 414.7484 | 827.4823 | 827.4865 | -4.99 | 0 | 25 | 0.0034 | 1 | U | R.ELLGGLAR.- |
| 105 | 210 - 217 | 828.4907 | 827.4834 | 827.4865 | -3.68 | 0 | 3 | 0.52 | 1 | U | R.ELLGGLAR.- |
| 106 | 210 - 217 | 828.4914 | 827.4842 | 827.4865 | -2.79 | 0 | 11 | 0.074 | 1 | U | R.ELLGGLAR.- |
| 107 | 210 - 217 | 828.4915 | 827.4842 | 827.4865 | -2.72 | 0 | 12 | 0.07 | 1 | U | R.ELLGGLAR.- |
| 108 | 210 - 217 | 828.4924 | 827.4851 | 827.4865 | -1.68 | 0 | 4 | 0.38 | 1 | U | R.ELLGGLAR.- |
| 109 | 210 - 217 | 414.7504 | 827.4862 | 827.4865 | -0.35 | 0 | 24 | 0.0041 | 1 | U | R.ELLGGLAR.- |

Error: try setting browser cache to automatic.

---

```
LOCUS       QNL33371                  48 aa            linear   BCT 04-SEP-2020
DEFINITION  Hypothetical protein (plasmid) [Escherichia coli].
ACCESSION   QNL33371
VERSION     QNL33371.1
DBSOURCE    accession MT180430.1
KEYWORDS    .
SOURCE      Escherichia coli
  ORGANISM  Escherichia coli
            Bacteria; Proteobacteria; Gammaproteobacteria; Enterobacterales;
            Enterobacteriaceae; Escherichia.
REFERENCE   1  (residues 1 to 48)
  AUTHORS   Tarabai,H., Wyrsch,E.R., Bitar,I., Djordjevic,S.P. and Dolejska,M.
  TITLE     Comparative analysis of multidrug resistant Escherichia coli ST216
            isolates from silver gulls in Australia
  JOURNAL   Unpublished
REFERENCE   2  (residues 1 to 48)
  AUTHORS   Tarabai,H., Wyrsch,E.R., Bitar,I., Djordjevic,S.P. and Dolejska,M.
  TITLE     Direct Submission
  JOURNAL   Submitted (09-MAR-2020) Department of Biology and Wildlife
            Diseases, University of Veterinary and Pharmaceutical Sciences
            Brno, Palackeho tr. 1946/1, Brno, Brno 61242, Czech Republic
COMMENT     ##Assembly-Data-START##
            Assembly Method       :: SMRT Link v. 8.0
            Assembly Name         :: pCE1681-A
            Coverage              :: 326X
            Sequencing Technology :: PacBio
            ##Assembly-Data-END##
FEATURES             Location/Qualifiers
     source          1..48
                     /organism="Escherichia coli"
                     /strain="CE1681"
                     /host="Chroicocephalus novaehollandiae"
                     /db_xref="taxon:562"
                     /plasmid="pCE1681-A"
                     /country="Australia"
                     /collection_date="2012"
                     /note="Closed IncHI2-ST3/IncN fusion plasmid;
                     type: ST216"
     Protein         1..48
                     /product="Hypothetical protein"
     CDS             1..48
                     /coded_by="complement(MT180430.1:247923..248069)"
                     /note="Hypothetical protein"
                     /transl_table=11
                     /db_xref="SEED:fig|6666666.506717.peg.328"
```

### Species 2

#### MATRIX SCIENCE MASCOT Search Results

##### Protein View: QNL33371.1

gsAK [Escherichia coli]

| | |
|---|---|
| Database: | NCBIcustTC |
| Score: | 478 |
| Monoisotopic mass ($M_r$): | 24127 |
| Calculated pI: | 5.86 |

Sequencesimilarity is availableas an NCBI BLAST search of QNL33371.1 against nr.

##### Search parameters

| | |
|---|---|
| MS data file: | 09182020_TC-3_001.temp.mgf |
| Enzyme: | Trypsin/P: cuts C-term side of KR. |
| Variable modifications: | Oxidation (M), Propionamide (C), Propionamide (K), Propionamide (N-term) |

##### Protein sequence coverage: 35%

Matched peptides shown in **bold red**.

```
  1 MNLVLMGLPG AGKGTQAEKI VAAYGIPHIS TGDMFRAAMK EGTPLGLQAK
 51 QYMDRGDLVP DEVTIGIVRE RLSKDDCQNG FLLDGFPRTV AQAEALETML
101 ADIGRKLDYV IHIDVRQDVL MERLTGRRIC RNCGATYHLI FHPPAKPGVC
151 DKCGGELYQR ADDNEATVAN RLEVNMKQMK PLVDFYEQKG YLRNINGEQD
201 MEKVFADIRE LLGGLAR
```

Unformatted sequencestring: 217 residues (for pasting into other applications).

Sort by ◉ residue number ○ increasing mass ○ decreasing mass
Show ◉ matched peptides only ○ predicted peptides also
Show ◉ uncorrected delta ○ delta corrected for 13C

| Query | Start – End | Observed | Mr(expt) | Mr(calc) | ppm | M | Score | Expect | Rank | U | Peptide |
|---|---|---|---|---|---|---|---|---|---|---|---|
| 563 | 51 – 69 | 731.3592 | 2191.0559 | 2191.0943 | -17.5 | 1 | 8 | 0.17 | 1 | U | K.QYMDRGDLVPDEVTIGIVR.E + Oxidation (M) |
| 565 | 51 – 69 | 731.3649 | 2191.0728 | 2191.0943 | -9.81 | 1 | 27 | 0.0023 | 1 | U | K.QYMDRGDLVPDEVTIGIVR.E + Oxidation (M) |
| 566 | 51 – 69 | 731.3656 | 2191.0750 | 2191.0943 | -8.82 | 1 | 22 | 0.0083 | 1 | U | K.QYMDRGDLVPDEVTIGIVR.E + Oxidation (M) |
| 567 | 51 – 69 | 731.3664 | 2191.0774 | 2191.0943 | -7.74 | 1 | 19 | 0.018 | 1 | U | K.QYMDRGDLVPDEVTIGIVR.E + Oxidation (M) |
| 569 | 51 – 69 | 731.3676 | 2191.0809 | 2191.0943 | -6.12 | 1 | 15 | 0.043 | 1 | U | K.QYMDRGDLVPDEVTIGIVR.E + Oxidation (M) |
| 570 | 51 – 69 | 731.3703 | 2191.0892 | 2191.0943 | -2.36 | 1 | 5 | 0.41 | 1 | U | K.QYMDRGDLVPDEVTIGIVR.E + Oxidation (M) |
| 282 | 56 – 69 | 741.9047 | 1481.7949 | 1481.8090 | -9.49 | 0 | 34 | 0.00036 | 1 | U | R.GDLVPDEVTIGIVR.E |
| 283 | 56 – 69 | 741.9062 | 1481.7978 | 1481.8090 | -7.52 | 0 | 30 | 0.001 | 1 | U | R.GDLVPDEVTIGIVR.E |
| 284 | 56 – 69 | 741.9072 | 1481.7999 | 1481.8090 | -6.15 | 0 | 20 | 0.011 | 1 | U | R.GDLVPDEVTIGIVR.E |
| 285 | 56 – 69 | 741.9081 | 1481.8017 | 1481.8090 | -4.93 | 0 | 32 | 0.00058 | 1 | U | R.GDLVPDEVTIGIVR.E |
| 286 | 56 – 69 | 741.9087 | 1481.8028 | 1481.8090 | -4.15 | 0 | 43 | 5.3e-005 | 1 | U | R.GDLVPDEVTIGIVR.E |
| 287 | 56 – 69 | 741.9088 | 1481.8030 | 1481.8090 | -4.00 | 0 | 32 | 0.00064 | 1 | U | R.GDLVPDEVTIGIVR.E |
| 288 | 56 – 69 | 741.9091 | 1481.8037 | 1481.8090 | -3.54 | 0 | 43 | 4.9e-005 | 1 | U | R.GDLVPDEVTIGIVR.E |
| 289 | 56 – 69 | 741.9091 | 1481.8037 | 1481.8090 | -3.54 | 0 | 47 | 1.8e-005 | 1 | U | R.GDLVPDEVTIGIVR.E |
| 399 | 89 – 105 | 894.9485 | 1787.8824 | 1787.9087 | -14.7 | 0 | 18 | 0.015 | 1 | U | R.TVAQAEALETMLADIGR.K |
| 400 | 89 – 105 | 894.9530 | 1787.8914 | 1787.9087 | -9.67 | 0 | 30 | 0.00092 | 1 | U | R.TVAQAEALETMLADIGR.K |
| 401 | 89 – 105 | 894.9530 | 1787.8914 | 1787.9087 | -9.67 | 0 | 18 | 0.015 | 1 | U | R.TVAQAEALETMLADIGR.K |
| 402 | 89 – 105 | 894.9536 | 1787.8927 | 1787.9087 | -8.97 | 0 | 4 | 0.44 | 1 | U | R.TVAQAEALETMLADIGR.K |
| 403 | 89 – 105 | 894.9565 | 1787.8984 | 1787.9087 | -5.77 | 0 | 14 | 0.036 | 1 | U | R.TVAQAEALETMLADIGR.K |
| 404 | 89 – 105 | 602.3003 | 1803.8792 | 1803.9036 | -13.6 | 0 | 16 | 0.024 | 1 | U | R.TVAQAEALETMLADIGR.K + Oxidation (M) |
| 405 | 89 – 105 | 602.3029 | 1803.8870 | 1803.9036 | -9.23 | 0 | 17 | 0.022 | 1 | U | R.TVAQAEALETMLADIGR.K + Oxidation (M) |
| 406 | 89 – 105 | 902.9510 | 1803.8874 | 1803.9036 | -9.02 | 0 | 25 | 0.0029 | 1 | U | R.TVAQAEALETMLADIGR.K + Oxidation (M) |
| 407 | 89 – 105 | 602.3031 | 1803.8875 | 1803.9036 | -8.95 | 0 | 13 | 0.047 | 1 | U | R.TVAQAEALETMLADIGR.K + Oxidation (M) |
| 408 | 89 – 105 | 902.9513 | 1803.8880 | 1803.9036 | -8.64 | 0 | 47 | 2e-005 | 1 | U | R.TVAQAEALETMLADIGR.K + Oxidation (M) |
| 409 | 89 – 105 | 602.3037 | 1803.8892 | 1803.9036 | -7.98 | 0 | 5 | 0.31 | 1 | U | R.TVAQAEALETMLADIGR.K + Oxidation (M) |
| 411 | 89 – 105 | 902.9539 | 1803.8932 | 1803.9036 | -5.81 | 0 | 52 | 6.7e-006 | 1 | U | R.TVAQAEALETMLADIGR.K + Oxidation (M) |
| 412 | 89 – 105 | 902.9551 | 1803.8956 | 1803.9036 | -4.43 | 0 | 43 | 4.6e-005 | 1 | U | R.TVAQAEALETMLADIGR.K + Oxidation (M) |
| 413 | 89 – 105 | 902.9551 | 1803.8957 | 1803.9036 | -4.41 | 0 | 28 | 0.0015 | 1 | U | R.TVAQAEALETMLADIGR.K + Oxidation (M) |

| Query | Start – End | Observed | Mr(expt) | Mr(calc) | ppm | M | Score | Expect | Rank | U | Peptide |
|---|---|---|---|---|---|---|---|---|---|---|---|
| 414 | 89 – 105 | 902.9553 | 1803.8961 | 1803.9036 | -4.16 | 0 | 43 | 4.9e-005 | 1 | U | R.TVAQAEALETMLADIGR.K + Oxidation (M) |
| 415 | 89 – 105 | 902.9555 | 1803.8964 | 1803.9036 | -3.99 | 0 | 17 | 0.021 | 1 | U | R.TVAQAEALETMLADIGR.K + Oxidation (M) |
| 416 | 89 – 105 | 602.3070 | 1803.8991 | 1803.9036 | -2.49 | 0 | 3 | 0.46 | 1 | U | R.TVAQAEALETMLADIGR.K + Oxidation (M) |
| 417 | 89 – 105 | 902.9597 | 1803.9049 | 1803.9036 | 0.68 | 0 | 17 | 0.022 | 1 | U | R.TVAQAEALETMLADIGR.K + Oxidation (M) |
| 265 | 106 – 116 | 457.5923 | 1369.7550 | 1369.7718 | -12.2 | 1 | 13 | 0.053 | 1 | U | R.KLDYVIHIDVR.Q |
| 269 | 106 – 116 | 457.5933 | 1369.7579 | 1369.7718 | -10.1 | 1 | 4 | 0.36 | 1 | U | R.KLDYVIHIDVR.Q |
| 427 | 161 – 177 | 635.9629 | 1904.8669 | 1904.8898 | -12.0 | 1 | 28 | 0.0016 | 1 | U | R.ADDNEATVANRLEVNMK.Q + Oxidation (M) |
| 428 | 161 – 177 | 635.9629 | 1904.8669 | 1904.8898 | -12.0 | 1 | 17 | 0.022 | 1 | U | R.ADDNEATVANRLEVNMK.Q + Oxidation (M) |
| 429 | 161 – 177 | 635.9633 | 1904.8681 | 1904.8898 | -11.4 | 1 | 3 | 0.59 | 1 | U | R.ADDNEATVANRLEVNMK.Q + Oxidation (M) |
| 430 | 161 – 177 | 635.9649 | 1904.8728 | 1904.8898 | -8.90 | 1 | 0 | 0.96 | 1 | U | R.ADDNEATVANRLEVNMK.Q + Oxidation (M) |
| 431 | 161 – 177 | 635.9654 | 1904.8743 | 1904.8898 | -8.13 | 1 | 13 | 0.057 | 1 | U | R.ADDNEATVANRLEVNMK.Q + Oxidation (M) |
| 432 | 161 – 177 | 635.9657 | 1904.8753 | 1904.8898 | -7.61 | 1 | 14 | 0.045 | 1 | U | R.ADDNEATVANRLEVNMK.Q + Oxidation (M) |
| 433 | 161 – 177 | 635.9657 | 1904.8753 | 1904.8898 | -7.61 | 1 | 25 | 0.0032 | 1 | U | R.ADDNEATVANRLEVNMK.Q + Oxidation (M) |
| 434 | 161 – 177 | 635.9659 | 1904.8760 | 1904.8898 | -7.25 | 1 | 31 | 0.00077 | 1 | U | R.ADDNEATVANRLEVNMK.Q + Oxidation (M) |
| 435 | 161 – 177 | 635.9659 | 1904.8760 | 1904.8898 | -7.25 | 1 | 17 | 0.021 | 1 | U | R.ADDNEATVANRLEVNMK.Q + Oxidation (M) |
| 436 | 161 – 177 | 635.9666 | 1904.8780 | 1904.8898 | -6.16 | 1 | 1 | 0.87 | 1 | U | R.ADDNEATVANRLEVNMK.Q + Oxidation (M) |
| 437 | 161 – 177 | 635.9671 | 1904.8795 | 1904.8898 | -5.41 | 1 | 12 | 0.079 | 1 | U | R.ADDNEATVANRLEVNMK.Q + Oxidation (M) |
| 438 | 161 – 177 | 953.4480 | 1904.8814 | 1904.8898 | -4.40 | 1 | 8 | 0.2 | 1 | U | R.ADDNEATVANRLEVNMK.Q + Oxidation (M) |
| 314 | 178 – 189 | 771.3815 | 1540.7484 | 1540.7596 | -7.23 | 1 | 1 | 1.5 | 1 | U | K.QMKPLVDFYEQK.G + Oxidation (M) |

Error: try setting browser cache to automatic.

---

```
LOCUS       QNL33371                  48 aa            linear   BCT 04-SEP-2020
DEFINITION  Hypothetical protein (plasmid) [Escherichia coli].
ACCESSION   QNL33371
VERSION     QNL33371.1
DBSOURCE    accession MT180430.1
KEYWORDS    .
SOURCE      Escherichia coli
  ORGANISM  Escherichia coli
            Bacteria; Proteobacteria; Gammaproteobacteria; Enterobacterales;
            Enterobacteriaceae; Escherichia.
REFERENCE   1  (residues 1 to 48)
  AUTHORS   Tarabai,H., Wyrsch,E.R., Bitar,I., Djordjevic,S.P. and Dolejska,M.
  TITLE     Comparative analysis of multidrug resistant Escherichia coli ST216
            isolates from silver gulls in Australia
  JOURNAL   Unpublished
REFERENCE   2  (residues 1 to 48)
  AUTHORS   Tarabai,H., Wyrsch,E.R., Bitar,I., Djordjevic,S.P. and Dolejska,M.
  TITLE     Direct Submission
  JOURNAL   Submitted (09-MAR-2020) Department of Biology and Wildlife
            Diseases, University of Veterinary and Pharmaceutical Sciences
            Brno, Palackeho tr. 1946/1, Brno, Brno 61242, Czech Republic
COMMENT     ##Assembly-Data-START##
            Assembly Method       :: SMRT Link v. 8.0
            Assembly Name         :: pCE1681-A
            Coverage              :: 326X
            Sequencing Technology :: PacBio
            ##Assembly-Data-END##
FEATURES             Location/Qualifiers
     source          1..48
                     /organism="Escherichia coli"
                     /strain="CE1681"
                     /host="Chroicocephalus novaehollandiae"
                     /db_xref="taxon:562"
                     /plasmid="pCE1681-A"
                     /country="Australia"
                     /collection_date="2012"
                     /note="Closed IncHI2-ST3/IncN fusion plasmid;
                     type: ST216"
     Protein         1..48
                     /product="Hypothetical protein"
     CDS             1..48
```

```
/coded_by="complement(MT180430.1:247923..248069)"
/note="Hypothetical protein"
/transl_table=11
/db_xref="SEED:fig|6666666.506717.peg.328"
```

Mascot: http://www.matrixscience.com/

### Species 3

#### MASCOT Search Results

##### Protein View: QNL33370.1

gsAK-cp45 [Escherichia coli]

| | |
|---|---|
| Database: | NCBIcustTC |
| Score: | 3060 |
| Monoisotopic mass ($M_r$): | 26627 |
| Calculated pI: | 6.45 |

Sequencesimilarity is availableas an NCBI BLAST search of QNL33370.1 against nr.

##### Search parameters

| | |
|---|---|
| MS data file: | 09182020_TC-4_001.temp.mgf |
| Enzyme: | Trypsin/P: cuts C-term side of KR. |
| Variable modifications: | Oxidation (M), Propionamide (C), Propionamide (K), Propionamide (N-term) |

##### Protein sequence coverage: 70%

Matched peptides shown in **bold red**.

```
  1 MGFRIYRETL SRFSCAAQLG LQAKQYMDRG DLVPDEVTIG IVRERLSKDD
 51 CQNGFLLDGF PRTVAQAEAL ETMLADIGRK LDYVIHIDVR QDVLMERLTG
101 RRICRNCGAT YHLIFHPPAK PGVCDKCGGE LYQRADDNEA TVANRLEVNM
151 KQMKPLVDFY EQKGYLRNIN GEQDMEKVFA DIRELLGGLA RAAAMNLVLM
201 GLPGAGKGTQ AEKIVAAYGI PHISTGDMFR AAMKEGTPLG
```

Unformatted sequencestring: 240 residues (for pasting into other applications).

Sort by ◉ residue number ○ increasing mass ○ decreasing mass
Show ◉ matched peptides only ○ predicted peptides also
Show ◉ uncorrected delta ○ delta corrected for 13C

| Query | Start – End | Observed | Mr(expt) | Mr(calc) | ppm | M | Score | Expect | Rank | U | Peptide |
|---|---|---|---|---|---|---|---|---|---|---|---|
| 360 | 13 – 24 | 618.8199 | 1235.6252 | 1235.6332 | -6.45 | 0 | 1 | 1.2 | 1 | U | R.FSCAAQLGLQAK.Q |
| 374 | 13 – 24 | 654.3359 | 1306.6573 | 1306.6703 | -9.98 | 0 | 30 | 0.0014 | 1 | U | R.FSCAAQLGLQAK.Q + Propionamide (C) |
| 375 | 13 – 24 | 654.3362 | 1306.6579 | 1306.6703 | -9.48 | 0 | 9 | 0.19 | 1 | U | R.FSCAAQLGLQAK.Q + Propionamide (C) |
| 376 | 13 – 24 | 654.3365 | 1306.6585 | 1306.6703 | -9.02 | 0 | 10 | 0.13 | 1 | U | R.FSCAAQLGLQAK.Q + Propionamide (C) |
| 377 | 13 – 24 | 654.3374 | 1306.6603 | 1306.6703 | -7.67 | 0 | 25 | 0.0042 | 1 | U | R.FSCAAQLGLQAK.Q + Propionamide (C) |
| 378 | 13 – 24 | 654.3416 | 1306.6686 | 1306.6703 | -1.31 | 0 | 32 | 0.00086 | 1 | U | R.FSCAAQLGLQAK.Q + Propionamide (C) |
| 379 | 13 – 24 | 654.3430 | 1306.6715 | 1306.6703 | 0.88 | 0 | 9 | 0.18 | 1 | U | R.FSCAAQLGLQAK.Q + Propionamide (C) |
| 898 | 25 – 43 | 726.0377 | 2175.0914 | 2175.0994 | -3.67 | 1 | 19 | 0.017 | 1 | U | K.QYMDRGDLVPDEVTIGIVR.E |
| 899 | 25 – 43 | 726.0384 | 2175.0935 | 2175.0994 | -2.74 | 1 | 18 | 0.022 | 1 | U | K.QYMDRGDLVPDEVTIGIVR.E |
| 900 | 25 – 43 | 726.0416 | 2175.1030 | 2175.0994 | 1.65 | 1 | 36 | 0.00037 | 1 | U | K.QYMDRGDLVPDEVTIGIVR.E |
| 902 | 25 – 43 | 731.3623 | 2191.0652 | 2191.0943 | -13.3 | 1 | 20 | 0.011 | 1 | U | K.QYMDRGDLVPDEVTIGIVR.E + Oxidation (M) |
| 903 | 25 – 43 | 731.3646 | 2191.0719 | 2191.0943 | -10.2 | 1 | 31 | 0.001 | 1 | U | K.QYMDRGDLVPDEVTIGIVR.E + Oxidation (M) |
| 904 | 25 – 43 | 731.3648 | 2191.0725 | 2191.0943 | -9.96 | 1 | 37 | 0.00028 | 1 | U | K.QYMDRGDLVPDEVTIGIVR.E + Oxidation (M) |
| 905 | 25 – 43 | 731.3648 | 2191.0725 | 2191.0943 | -9.96 | 1 | 53 | 6.5e-006 | 1 | U | K.QYMDRGDLVPDEVTIGIVR.E + Oxidation (M) |
| 906 | 25 – 43 | 731.3648 | 2191.0726 | 2191.0943 | -9.92 | 1 | 61 | 1.1e-006 | 1 | U | K.QYMDRGDLVPDEVTIGIVR.E + Oxidation (M) |
| 907 | 25 – 43 | 731.3649 | 2191.0729 | 2191.0943 | -9.78 | 1 | 49 | 1.6e-005 | 1 | U | K.QYMDRGDLVPDEVTIGIVR.E + Oxidation (M) |
| 908 | 25 – 43 | 731.3649 | 2191.0729 | 2191.0943 | -9.78 | 1 | 53 | 5.9e-006 | 1 | U | K.QYMDRGDLVPDEVTIGIVR.E + Oxidation (M) |
| 909 | 25 – 43 | 731.3650 | 2191.0731 | 2191.0943 | -9.71 | 1 | 49 | 1.6e-005 | 1 | U | K.QYMDRGDLVPDEVTIGIVR.E + Oxidation (M) |
| 910 | 25 – 43 | 731.3650 | 2191.0733 | 2191.0943 | -9.60 | 1 | 18 | 0.02 | 1 | U | K.QYMDRGDLVPDEVTIGIVR.E + Oxidation (M) |
| 911 | 25 – 43 | 731.3652 | 2191.0737 | 2191.0943 | -9.44 | 1 | 48 | 2.1e-005 | 1 | U | K.QYMDRGDLVPDEVTIGIVR.E + Oxidation (M) |
| 912 | 25 – 43 | 731.3652 | 2191.0737 | 2191.0943 | -9.44 | 1 | 53 | 6e-006 | 1 | U | K.QYMDRGDLVPDEVTIGIVR.E + Oxidation (M) |
| 913 | 25 – 43 | 731.3660 | 2191.0761 | 2191.0943 | -8.33 | 1 | 56 | 3.5e-006 | 1 | U | K.QYMDRGDLVPDEVTIGIVR.E + Oxidation (M) |
| 914 | 25 – 43 | 731.3662 | 2191.0769 | 2191.0943 | -7.96 | 1 | 41 | 9.4e-005 | 1 | U | K.QYMDRGDLVPDEVTIGIVR.E + Oxidation (M) |
| 915 | 25 – 43 | 731.3664 | 2191.0772 | 2191.0943 | -7.81 | 1 | 9 | 0.18 | 1 | U | K.QYMDRGDLVPDEVTIGIVR.E + Oxidation (M) |
| 916 | 25 – 43 | 731.3664 | 2191.0775 | 2191.0943 | -7.68 | 1 | 22 | 0.0076 | 1 | U | K.QYMDRGDLVPDEVTIGIVR.E + Oxidation (M) |

| Query | Start – End | Observed | Mr(expt) | Mr(calc) | ppm | M | Score | Expect | Rank | U | Peptide |
|---|---|---|---|---|---|---|---|---|---|---|---|
| 917 | 25 – 43 | 731.3677 | 2191.0813 | 2191.0943 | -5.96 | 1 | 41 | 0.00011 | 1 | U | K.QYMDRGDLVPDEVTIGIVR.E + Oxidation (M) |
| 463 | 30 – 43 | 741.9058 | 1481.7969 | 1481.8090 | -8.12 | 0 | 25 | 0.0033 | 1 | U | R.GDLVPDEVTIGIVR.E |
| 464 | 30 – 43 | 741.9058 | 1481.7971 | 1481.8090 | -8.02 | 0 | 47 | 2.1e-005 | 1 | U | R.GDLVPDEVTIGIVR.E |
| 465 | 30 – 43 | 741.9059 | 1481.7972 | 1481.8090 | -7.93 | 0 | 49 | 1.2e-005 | 1 | U | R.GDLVPDEVTIGIVR.E |
| 466 | 30 – 43 | 741.9071 | 1481.7997 | 1481.8090 | -6.25 | 0 | 47 | 2.1e-005 | 1 | U | R.GDLVPDEVTIGIVR.E |
| 467 | 30 – 43 | 741.9072 | 1481.7999 | 1481.8090 | -6.09 | 0 | 24 | 0.0043 | 1 | U | R.GDLVPDEVTIGIVR.E |
| 468 | 30 – 43 | 741.9073 | 1481.8000 | 1481.8090 | -6.06 | 0 | 25 | 0.0033 | 1 | U | R.GDLVPDEVTIGIVR.E |
| 469 | 30 – 43 | 741.9073 | 1481.8001 | 1481.8090 | -5.97 | 0 | 53 | 5.3e-006 | 1 | U | R.GDLVPDEVTIGIVR.E |
| 470 | 30 – 43 | 741.9076 | 1481.8007 | 1481.8090 | -5.59 | 0 | 57 | 2.1e-006 | 1 | U | R.GDLVPDEVTIGIVR.E |
| 471 | 30 – 43 | 741.9076 | 1481.8007 | 1481.8090 | -5.59 | 0 | 52 | 6.3e-006 | 1 | U | R.GDLVPDEVTIGIVR.E |
| 472 | 30 – 43 | 741.9079 | 1481.8011 | 1481.8090 | -5.28 | 0 | 61 | 7.4e-007 | 1 | U | R.GDLVPDEVTIGIVR.E |
| 473 | 30 – 43 | 741.9079 | 1481.8012 | 1481.8090 | -5.23 | 0 | 49 | 1.4e-005 | 1 | U | R.GDLVPDEVTIGIVR.E |
| 474 | 30 – 43 | 741.9082 | 1481.8019 | 1481.8090 | -4.76 | 0 | 50 | 1e-005 | 1 | U | R.GDLVPDEVTIGIVR.E |
| 475 | 30 – 43 | 741.9089 | 1481.8032 | 1481.8090 | -3.91 | 0 | 57 | 2.2e-006 | 1 | U | R.GDLVPDEVTIGIVR.E |
| 476 | 30 – 43 | 741.9113 | 1481.8081 | 1481.8090 | -0.59 | 0 | 5 | 0.28 | 1 | U | R.GDLVPDEVTIGIVR.E |
| 477 | 30 – 43 | 741.9144 | 1481.8142 | 1481.8090 | 3.56 | 0 | 9 | 0.12 | 1 | U | R.GDLVPDEVTIGIVR.E |
| 780 | 46 – 62 | 665.9809 | 1994.9208 | 1994.9520 | -15.7 | 1 | 8 | 0.15 | 1 | U | R.LSKDDCQNGFLLDGFPR.T + Propionamide (C) |
| 781 | 46 – 62 | 665.9822 | 1994.9249 | 1994.9520 | -13.6 | 1 | 15 | 0.033 | 1 | U | R.LSKDDCQNGFLLDGFPR.T + Propionamide (C) |
| 782 | 46 – 62 | 665.9844 | 1994.9314 | 1994.9520 | -10.3 | 1 | 19 | 0.015 | 1 | U | R.LSKDDCQNGFLLDGFPR.T + Propionamide (C) |
| 784 | 46 – 62 | 666.3142 | 1995.9206 | 1994.9520 | 486 | 1 | 17 | 0.021 | 1 | U | R.LSKDDCQNGFLLDGFPR.T + Propionamide (C) |
| 785 | 46 – 62 | 666.3170 | 1995.9292 | 1994.9520 | 490 | 1 | 30 | 0.0011 | 1 | U | R.LSKDDCQNGFLLDGFPR.T + Propionamide (C) |
| 786 | 46 – 62 | 666.3170 | 1995.9292 | 1994.9520 | 490 | 1 | 1 | 0.82 | 1 | U | R.LSKDDCQNGFLLDGFPR.T + Propionamide (C) |
| 787 | 46 – 62 | 666.3192 | 1995.9357 | 1994.9520 | 493 | 1 | 26 | 0.0026 | 1 | U | R.LSKDDCQNGFLLDGFPR.T + Propionamide (C) |
| 788 | 46 – 62 | 666.3192 | 1995.9357 | 1994.9520 | 493 | 1 | 7 | 0.25 | 1 | U | R.LSKDDCQNGFLLDGFPR.T + Propionamide (C) |
| 789 | 46 – 62 | 666.3196 | 1995.9371 | 1994.9520 | 494 | 1 | 29 | 0.0013 | 1 | U | R.LSKDDCQNGFLLDGFPR.T + Propionamide (C) |
| 790 | 46 – 62 | 666.3202 | 1995.9387 | 1994.9520 | 495 | 1 | 25 | 0.0035 | 1 | U | R.LSKDDCQNGFLLDGFPR.T + Propionamide (C) |
| 791 | 46 – 62 | 666.3314 | 1995.9725 | 1994.9520 | 512 | 1 | 14 | 0.046 | 1 | U | R.LSKDDCQNGFLLDGFPR.T + Propionamide (N-term) |
| 687 | 63 – 79 | 894.9537 | 1787.8929 | 1787.9087 | -8.83 | 0 | 25 | 0.0034 | 1 | U | R.TVAQAEALETMLADIGR.K |
| 688 | 63 – 79 | 894.9546 | 1787.8947 | 1787.9087 | -7.86 | 0 | 58 | 1.7e-006 | 1 | U | R.TVAQAEALETMLADIGR.K |
| 689 | 63 – 79 | 894.9551 | 1787.8957 | 1787.9087 | -7.30 | 0 | 33 | 0.0005 | 1 | U | R.TVAQAEALETMLADIGR.K |
| 690 | 63 – 79 | 894.9564 | 1787.8982 | 1787.9087 | -5.89 | 0 | 21 | 0.0081 | 1 | U | R.TVAQAEALETMLADIGR.K |
| 691 | 63 – 79 | 894.9565 | 1787.8984 | 1787.9087 | -5.76 | 0 | 16 | 0.025 | 1 | U | R.TVAQAEALETMLADIGR.K |
| 692 | 63 – 79 | 894.9617 | 1787.9088 | 1787.9087 | 0.037 | 0 | 58 | 1.5e-006 | 1 | U | R.TVAQAEALETMLADIGR.K |
| 693 | 63 – 79 | 602.3000 | 1803.8782 | 1803.9036 | -14.1 | 0 | 35 | 0.00033 | 1 | U | R.TVAQAEALETMLADIGR.K + Oxidation (M) |
| 694 | 63 – 79 | 602.3008 | 1803.8806 | 1803.9036 | -12.8 | 0 | 45 | 3.2e-005 | 1 | U | R.TVAQAEALETMLADIGR.K + Oxidation (M) |
| 695 | 63 – 79 | 602.3030 | 1803.8872 | 1803.9036 | -9.13 | 0 | 50 | 1.1e-005 | 1 | U | R.TVAQAEALETMLADIGR.K + Oxidation (M) |
| 696 | 63 – 79 | 602.3037 | 1803.8894 | 1803.9036 | -7.92 | 0 | 50 | 1e-005 | 1 | U | R.TVAQAEALETMLADIGR.K + Oxidation (M) |
| 697 | 63 – 79 | 602.3041 | 1803.8904 | 1803.9036 | -7.33 | 0 | 39 | 0.00012 | 1 | U | R.TVAQAEALETMLADIGR.K + Oxidation (M) |
| 698 | 63 – 79 | 902.9527 | 1803.8908 | 1803.9036 | -7.13 | 0 | 18 | 0.017 | 1 | U | R.TVAQAEALETMLADIGR.K + Oxidation (M) |
| 699 | 63 – 79 | 902.9527 | 1803.8909 | 1803.9036 | -7.05 | 0 | 73 | 5.4e-008 | 1 | U | R.TVAQAEALETMLADIGR.K + Oxidation (M) |
| 700 | 63 – 79 | 902.9536 | 1803.8927 | 1803.9036 | -6.08 | 0 | 78 | 1.7e-008 | 1 | U | R.TVAQAEALETMLADIGR.K + Oxidation (M) |
| 701 | 63 – 79 | 902.9545 | 1803.8944 | 1803.9036 | -5.13 | 0 | 66 | 2.5e-007 | 1 | U | R.TVAQAEALETMLADIGR.K + Oxidation (M) |
| 702 | 63 – 79 | 902.9549 | 1803.8953 | 1803.9036 | -4.63 | 0 | 57 | 2e-006 | 1 | U | R.TVAQAEALETMLADIGR.K + Oxidation (M) |
| 703 | 63 – 79 | 902.9550 | 1803.8955 | 1803.9036 | -4.50 | 0 | 69 | 1.2e-007 | 1 | U | R.TVAQAEALETMLADIGR.K + Oxidation (M) |
| 704 | 63 – 79 | 902.9552 | 1803.8957 | 1803.9036 | -4.38 | 0 | 59 | 1.3e-006 | 1 | U | R.TVAQAEALETMLADIGR.K + Oxidation (M) |
| 705 | 63 – 79 | 602.3062 | 1803.8968 | 1803.9036 | -3.77 | 0 | 46 | 2.7e-005 | 1 | U | R.TVAQAEALETMLADIGR.K + Oxidation (M) |
| 706 | 63 – 79 | 902.9564 | 1803.8983 | 1803.9036 | -2.96 | 0 | 41 | 8.3e-005 | 1 | U | R.TVAQAEALETMLADIGR.K + Oxidation (M) |
| 707 | 63 – 79 | 902.9566 | 1803.8986 | 1803.9036 | -2.78 | 0 | 63 | 5.2e-007 | 1 | U | R.TVAQAEALETMLADIGR.K + Oxidation (M) |
| 708 | 63 – 79 | 902.9576 | 1803.9007 | 1803.9036 | -1.64 | 0 | 66 | 2.4e-007 | 1 | U | R.TVAQAEALETMLADIGR.K + Oxidation (M) |
| 709 | 63 – 79 | 902.9577 | 1803.9009 | 1803.9036 | -1.52 | 0 | 21 | 0.0077 | 1 | U | R.TVAQAEALETMLADIGR.K + Oxidation (M) |
| 710 | 63 – 79 | 602.3081 | 1803.9024 | 1803.9036 | -0.71 | 0 | 1 | 0.76 | 1 | U | R.TVAQAEALETMLADIGR.K + Oxidation (M) |
| 711 | 63 – 79 | 602.3095 | 1803.9067 | 1803.9036 | 1.71 | 0 | 25 | 0.0034 | 1 | U | R.TVAQAEALETMLADIGR.K + Oxidation (M) |
| 410 | 80 – 90 | 685.8811 | 1369.7477 | 1369.7718 | -17.6 | 1 | 7 | 0.19 | 1 | U | R.KLDYVIHIDVR.Q |
| 412 | 80 – 90 | 457.5907 | 1369.7502 | 1369.7718 | -15.8 | 1 | 7 | 0.2 | 1 | U | R.KLDYVIHIDVR.Q |
| 413 | 80 – 90 | 457.5909 | 1369.7508 | 1369.7718 | -15.3 | 1 | 15 | 0.035 | 1 | U | R.KLDYVIHIDVR.Q |
| 414 | 80 – 90 | 457.5917 | 1369.7531 | 1369.7718 | -13.6 | 1 | 4 | 0.39 | 1 | U | R.KLDYVIHIDVR.Q |
| 415 | 80 – 90 | 457.5921 | 1369.7544 | 1369.7718 | -12.7 | 1 | 15 | 0.034 | 1 | U | R.KLDYVIHIDVR.Q |
| 417 | 80 – 90 | 457.5926 | 1369.7558 | 1369.7718 | -11.6 | 1 | 11 | 0.083 | 1 | U | R.KLDYVIHIDVR.Q |
| 418 | 80 – 90 | 457.5926 | 1369.7560 | 1369.7718 | -11.5 | 1 | 10 | 0.11 | 1 | U | R.KLDYVIHIDVR.Q |
| 419 | 80 – 90 | 457.5928 | 1369.7566 | 1369.7718 | -11.1 | 1 | 23 | 0.0055 | 1 | U | R.KLDYVIHIDVR.Q |
| 420 | 80 – 90 | 457.5929 | 1369.7567 | 1369.7718 | -11.0 | 1 | 19 | 0.013 | 1 | U | R.KLDYVIHIDVR.Q |
| 422 | 80 – 90 | 457.5931 | 1369.7573 | 1369.7718 | -10.6 | 1 | 18 | 0.017 | 1 | U | R.KLDYVIHIDVR.Q |
| 423 | 80 – 90 | 457.5932 | 1369.7578 | 1369.7718 | -10.2 | 1 | 5 | 0.35 | 1 | U | R.KLDYVIHIDVR.Q |
| 424 | 80 – 90 | 457.5937 | 1369.7594 | 1369.7718 | -9.06 | 1 | 3 | 0.47 | 1 | U | R.KLDYVIHIDVR.Q |

| Query | Start – End | Observed | Mr(expt) | Mr(calc) | ppm | M | Score | Expect | Rank | U | Peptide |
|---|---|---|---|---|---|---|---|---|---|---|---|
| 425 | 80 – 90 | 457.5938 | 1369.7595 | 1369.7718 | -9.00 | 1 | 16 | 0.027 | 1 | U | R.KLDYVIHIDVR.Q |
| 426 | 80 – 90 | 457.5938 | 1369.7595 | 1369.7718 | -9.00 | 1 | 9 | 0.11 | 1 | U | R.KLDYVIHIDVR.Q |
| 427 | 80 – 90 | 457.5938 | 1369.7597 | 1369.7718 | -8.84 | 1 | 27 | 0.0018 | 1 | U | R.KLDYVIHIDVR.Q |
| 428 | 80 – 90 | 685.8872 | 1369.7597 | 1369.7718 | -8.78 | 1 | 13 | 0.048 | 1 | U | R.KLDYVIHIDVR.Q |
| 429 | 80 – 90 | 457.5939 | 1369.7598 | 1369.7718 | -8.75 | 1 | 4 | 0.41 | 1 | U | R.KLDYVIHIDVR.Q |
| 430 | 80 – 90 | 685.8874 | 1369.7603 | 1369.7718 | -8.37 | 1 | 5 | 0.31 | 1 | U | R.KLDYVIHIDVR.Q |
| 431 | 80 – 90 | 457.5941 | 1369.7604 | 1369.7718 | -8.29 | 1 | 15 | 0.031 | 1 | U | R.KLDYVIHIDVR.Q |
| 432 | 80 – 90 | 457.5941 | 1369.7606 | 1369.7718 | -8.19 | 1 | 22 | 0.0066 | 1 | U | R.KLDYVIHIDVR.Q |
| 433 | 80 – 90 | 457.5944 | 1369.7614 | 1369.7718 | -7.57 | 1 | 18 | 0.016 | 1 | U | R.KLDYVIHIDVR.Q |
| 434 | 80 – 90 | 685.8880 | 1369.7615 | 1369.7718 | -7.51 | 1 | 39 | 0.00013 | 1 | U | R.KLDYVIHIDVR.Q |
| 435 | 80 – 90 | 685.8884 | 1369.7623 | 1369.7718 | -6.94 | 1 | 27 | 0.002 | 1 | U | R.KLDYVIHIDVR.Q |
| 436 | 80 – 90 | 685.8884 | 1369.7623 | 1369.7718 | -6.93 | 1 | 12 | 0.06 | 1 | U | R.KLDYVIHIDVR.Q |
| 437 | 80 – 90 | 685.8890 | 1369.7635 | 1369.7718 | -6.06 | 1 | 40 | 0.0001 | 1 | U | R.KLDYVIHIDVR.Q |
| 438 | 80 – 90 | 685.8891 | 1369.7635 | 1369.7718 | -6.01 | 1 | 24 | 0.0043 | 1 | U | R.KLDYVIHIDVR.Q |
| 439 | 80 – 90 | 685.8892 | 1369.7639 | 1369.7718 | -5.74 | 1 | 22 | 0.006 | 1 | U | R.KLDYVIHIDVR.Q |
| 440 | 80 – 90 | 685.8892 | 1369.7639 | 1369.7718 | -5.71 | 1 | 31 | 0.00091 | 1 | U | R.KLDYVIHIDVR.Q |
| 441 | 80 – 90 | 685.8895 | 1369.7645 | 1369.7718 | -5.30 | 1 | 16 | 0.027 | 1 | U | R.KLDYVIHIDVR.Q |
| 442 | 80 – 90 | 685.8900 | 1369.7654 | 1369.7718 | -4.68 | 1 | 11 | 0.085 | 1 | U | R.KLDYVIHIDVR.Q |
| 443 | 80 – 90 | 685.8900 | 1369.7655 | 1369.7718 | -4.57 | 1 | 26 | 0.0026 | 1 | U | R.KLDYVIHIDVR.Q |
| 445 | 80 – 90 | 685.8911 | 1369.7677 | 1369.7718 | -2.97 | 1 | 13 | 0.049 | 1 | U | R.KLDYVIHIDVR.Q |
| 446 | 80 – 90 | 685.8917 | 1369.7689 | 1369.7718 | -2.11 | 1 | 18 | 0.017 | 1 | U | R.KLDYVIHIDVR.Q |
| 447 | 80 – 90 | 685.8923 | 1369.7700 | 1369.7718 | -1.32 | 1 | 17 | 0.02 | 1 | U | R.KLDYVIHIDVR.Q |
| 361 | 81 – 90 | 621.8313 | 1241.6481 | 1241.6768 | -23.2 | 0 | 18 | 0.016 | 1 | U | K.LDYVIHIDVR.Q |
| 194 | 91 – 97 | 453.7156 | 905.4167 | 905.4277 | -12.0 | 0 | 18 | 0.016 | 1 | U | R.QDVLMER.L + Oxidation (M) |
| 195 | 91 – 97 | 453.7167 | 905.4189 | 905.4277 | -9.64 | 0 | 26 | 0.0023 | 1 | U | R.QDVLMER.L + Oxidation (M) |
| 196 | 91 – 97 | 453.7172 | 905.4199 | 905.4277 | -8.52 | 0 | 22 | 0.0067 | 1 | U | R.QDVLMER.L + Oxidation (M) |
| 198 | 91 – 97 | 453.7196 | 905.4247 | 905.4277 | -3.26 | 0 | 3 | 0.5 | 1 | U | R.QDVLMER.L + Oxidation (M) |
| 199 | 91 – 97 | 906.4336 | 905.4263 | 905.4277 | -1.48 | 0 | 11 | 0.074 | 1 | U | R.QDVLMER.L + Oxidation (M) |
| 200 | 91 – 97 | 906.4336 | 905.4263 | 905.4277 | -1.48 | 0 | 2 | 0.57 | 1 | U | R.QDVLMER.L + Oxidation (M) |
| 326 | 135 – 145 | 588.2551 | 1174.4957 | 1174.5214 | -21.9 | 0 | 25 | 0.003 | 1 | U | R.ADDNEATVANR.L |
| 327 | 135 – 145 | 588.2594 | 1174.5043 | 1174.5214 | -14.6 | 0 | 17 | 0.019 | 1 | U | R.ADDNEATVANR.L |
| 328 | 135 – 145 | 588.2594 | 1174.5043 | 1174.5214 | -14.6 | 0 | 24 | 0.0043 | 1 | U | R.ADDNEATVANR.L |
| 329 | 135 – 145 | 588.2599 | 1174.5052 | 1174.5214 | -13.8 | 0 | 37 | 0.00019 | 1 | U | R.ADDNEATVANR.L |
| 330 | 135 – 145 | 588.2603 | 1174.5061 | 1174.5214 | -13.0 | 0 | 38 | 0.00018 | 1 | U | R.ADDNEATVANR.L |
| 331 | 135 – 145 | 588.2606 | 1174.5067 | 1174.5214 | -12.6 | 0 | 34 | 0.00043 | 1 | U | R.ADDNEATVANR.L |
| 332 | 135 – 145 | 588.2614 | 1174.5083 | 1174.5214 | -11.1 | 0 | 21 | 0.0073 | 1 | U | R.ADDNEATVANR.L |
| 333 | 135 – 145 | 588.2616 | 1174.5087 | 1174.5214 | -10.8 | 0 | 11 | 0.08 | 1 | U | R.ADDNEATVANR.L |
| 334 | 135 – 145 | 588.2623 | 1174.5101 | 1174.5214 | -9.67 | 0 | 13 | 0.046 | 1 | U | R.ADDNEATVANR.L |
| 335 | 135 – 145 | 588.2626 | 1174.5107 | 1174.5214 | -9.12 | 0 | 25 | 0.0031 | 1 | U | R.ADDNEATVANR.L |
| 336 | 135 – 145 | 588.2630 | 1174.5115 | 1174.5214 | -8.44 | 0 | 27 | 0.002 | 1 | U | R.ADDNEATVANR.L |
| 337 | 135 – 145 | 588.2630 | 1174.5115 | 1174.5214 | -8.44 | 0 | 20 | 0.01 | 1 | U | R.ADDNEATVANR.L |
| 338 | 135 – 145 | 588.2633 | 1174.5121 | 1174.5214 | -7.95 | 0 | 34 | 0.00043 | 1 | U | R.ADDNEATVANR.L |
| 339 | 135 – 145 | 588.2639 | 1174.5133 | 1174.5214 | -6.89 | 0 | 30 | 0.00096 | 1 | U | R.ADDNEATVANR.L |
| 341 | 135 – 145 | 588.2650 | 1174.5154 | 1174.5214 | -5.14 | 0 | 17 | 0.02 | 1 | U | R.ADDNEATVANR.L |
| 342 | 135 – 145 | 588.2655 | 1174.5165 | 1174.5214 | -4.20 | 0 | 29 | 0.0013 | 1 | U | R.ADDNEATVANR.L |
| 343 | 135 – 145 | 588.2662 | 1174.5177 | 1174.5214 | -3.13 | 0 | 21 | 0.0078 | 1 | U | R.ADDNEATVANR.L |
| 344 | 135 – 145 | 588.2665 | 1174.5184 | 1174.5214 | -2.58 | 0 | 12 | 0.067 | 1 | U | R.ADDNEATVANR.L |
| 345 | 135 – 145 | 588.2681 | 1174.5216 | 1174.5214 | 0.19 | 0 | 30 | 0.00091 | 1 | U | R.ADDNEATVANR.L |
| 739 | 135 – 151 | 635.9618 | 1904.8635 | 1904.8898 | -13.8 | 1 | 3 | 0.51 | 1 | U | R.ADDNEATVANRLEVNMK.Q + Oxidation (M) |
| 740 | 135 – 151 | 635.9618 | 1904.8635 | 1904.8898 | -13.8 | 1 | 6 | 0.29 | 1 | U | R.ADDNEATVANRLEVNMK.Q + Oxidation (M) |
| 741 | 135 – 151 | 953.4399 | 1904.8652 | 1904.8898 | -12.9 | 1 | 3 | 0.48 | 1 | U | R.ADDNEATVANRLEVNMK.Q + Oxidation (M) |
| 742 | 135 – 151 | 635.9637 | 1904.8692 | 1904.8898 | -10.8 | 1 | 23 | 0.0047 | 1 | U | R.ADDNEATVANRLEVNMK.Q + Oxidation (M) |
| 743 | 135 – 151 | 635.9638 | 1904.8695 | 1904.8898 | -10.7 | 1 | 49 | 1.2e-005 | 1 | U | R.ADDNEATVANRLEVNMK.Q + Oxidation (M) |
| 744 | 135 – 151 | 635.9639 | 1904.8700 | 1904.8898 | -10.4 | 1 | 25 | 0.0036 | 1 | U | R.ADDNEATVANRLEVNMK.Q + Oxidation (M) |
| 745 | 135 – 151 | 635.9646 | 1904.8719 | 1904.8898 | -9.39 | 1 | 48 | 1.6e-005 | 1 | U | R.ADDNEATVANRLEVNMK.Q + Oxidation (M) |
| 747 | 135 – 151 | 953.4434 | 1904.8722 | 1904.8898 | -9.20 | 1 | 6 | 0.29 | 1 | U | R.ADDNEATVANRLEVNMK.Q + Oxidation (M) |
| 748 | 135 – 151 | 635.9647 | 1904.8724 | 1904.8898 | -9.14 | 1 | 20 | 0.011 | 1 | U | R.ADDNEATVANRLEVNMK.Q + Oxidation (M) |
| 749 | 135 – 151 | 635.9648 | 1904.8725 | 1904.8898 | -9.04 | 1 | 32 | 0.00074 | 1 | U | R.ADDNEATVANRLEVNMK.Q + Oxidation (M) |
| 751 | 135 – 151 | 635.9656 | 1904.8751 | 1904.8898 | -7.69 | 1 | 5 | 0.37 | 1 | U | R.ADDNEATVANRLEVNMK.Q + Oxidation (M) |
| 752 | 135 – 151 | 635.9658 | 1904.8756 | 1904.8898 | -7.42 | 1 | 37 | 0.00022 | 1 | U | R.ADDNEATVANRLEVNMK.Q + Oxidation (M) |
| 753 | 135 – 151 | 635.9664 | 1904.8775 | 1904.8898 | -6.45 | 1 | 42 | 6.5e-005 | 1 | U | R.ADDNEATVANRLEVNMK.Q + Oxidation (M) |
| 754 | 135 – 151 | 635.9675 | 1904.8808 | 1904.8898 | -4.73 | 1 | 9 | 0.14 | 1 | U | R.ADDNEATVANRLEVNMK.Q + Oxidation (M) |
| 755 | 135 – 151 | 953.4478 | 1904.8811 | 1904.8898 | -4.56 | 1 | 2 | 0.77 | 1 | U | R.ADDNEATVANRLEVNMK.Q + Oxidation (M) |
| 756 | 135 – 151 | 953.4478 | 1904.8811 | 1904.8898 | -4.54 | 1 | 11 | 0.093 | 1 | U | R.ADDNEATVANRLEVNMK.Q + Oxidation (M) |

| Query | Start – End | Observed | Mr(expt) | Mr(calc) | ppm | M | Score | Expect | Rank | U | Peptide |
|---|---|---|---|---|---|---|---|---|---|---|---|
| 757 | 135 – 151 | 953.4483 | 1904.8820 | 1904.8898 | -4.08 | 1 | 7 | 0.25 | 1 | U | R.ADDNEATVANRLEVNMK.Q + Oxidation (M) |
| 758 | 135 – 151 | 953.4487 | 1904.8828 | 1904.8898 | -3.68 | 1 | 11 | 0.087 | 1 | U | R.ADDNEATVANRLEVNMK.Q + Oxidation (M) |
| 759 | 135 – 151 | 635.9685 | 1904.8836 | 1904.8898 | -3.25 | 1 | 2 | 0.67 | 1 | U | R.ADDNEATVANRLEVNMK.Q + Oxidation (M) |
| 520 | 152 – 163 | 514.5872 | 1540.7397 | 1540.7596 | -12.9 | 1 | 0 | 1.8 | 1 | U | K.QMKPLVDFYEQK.G + Oxidation (M) |
| 521 | 152 – 163 | 514.5875 | 1540.7406 | 1540.7596 | -12.3 | 1 | 13 | 0.11 | 1 | U | K.QMKPLVDFYEQK.G + Oxidation (M) |
| 524 | 152 – 163 | 514.5881 | 1540.7426 | 1540.7596 | -11.0 | 1 | 1 | 1.5 | 1 | U | K.QMKPLVDFYEQK.G + Oxidation (M) |
| 525 | 152 – 163 | 514.5882 | 1540.7428 | 1540.7596 | -10.9 | 1 | 14 | 0.073 | 1 | U | K.QMKPLVDFYEQK.G + Oxidation (M) |
| 526 | 152 – 163 | 771.3791 | 1540.7437 | 1540.7596 | -10.3 | 1 | 16 | 0.052 | 1 | U | K.QMKPLVDFYEQK.G + Oxidation (M) |
| 527 | 152 – 163 | 514.5888 | 1540.7446 | 1540.7596 | -9.72 | 1 | 24 | 0.0086 | 1 | U | K.QMKPLVDFYEQK.G + Oxidation (M) |
| 531 | 152 – 163 | 514.5892 | 1540.7458 | 1540.7596 | -8.92 | 1 | 2 | 1.2 | 1 | U | K.QMKPLVDFYEQK.G + Oxidation (M) |
| 532 | 152 – 163 | 771.3803 | 1540.7461 | 1540.7596 | -8.73 | 1 | 10 | 0.19 | 1 | U | K.QMKPLVDFYEQK.G + Oxidation (M) |
| 533 | 152 – 163 | 514.5898 | 1540.7475 | 1540.7596 | -7.83 | 1 | 12 | 0.12 | 1 | U | K.QMKPLVDFYEQK.G + Oxidation (M) |
| 534 | 152 – 163 | 514.5901 | 1540.7484 | 1540.7596 | -7.21 | 1 | 17 | 0.042 | 1 | U | K.QMKPLVDFYEQK.G + Oxidation (M) |
| 536 | 152 – 163 | 514.5905 | 1540.7495 | 1540.7596 | -6.51 | 1 | 9 | 0.24 | 1 | U | K.QMKPLVDFYEQK.G + Oxidation (M) |
| 537 | 152 – 163 | 771.3822 | 1540.7498 | 1540.7596 | -6.31 | 1 | 13 | 0.088 | 1 | U | K.QMKPLVDFYEQK.G + Oxidation (M) |
| 538 | 152 – 163 | 771.3833 | 1540.7520 | 1540.7596 | -4.87 | 1 | 14 | 0.07 | 1 | U | K.QMKPLVDFYEQK.G + Oxidation (M) |
| 539 | 152 – 163 | 771.3839 | 1540.7533 | 1540.7596 | -4.04 | 1 | 14 | 0.07 | 1 | U | K.QMKPLVDFYEQK.G + Oxidation (M) |
| 540 | 152 – 163 | 771.3851 | 1540.7556 | 1540.7596 | -2.56 | 1 | 40 | 0.00015 | 1 | U | K.QMKPLVDFYEQK.G + Oxidation (M) |
| 541 | 152 – 163 | 771.3853 | 1540.7559 | 1540.7596 | -2.34 | 1 | 22 | 0.0084 | 1 | U | K.QMKPLVDFYEQK.G + Oxidation (M) |
| 542 | 152 – 163 | 771.3892 | 1540.7639 | 1540.7596 | 2.85 | 1 | 32 | 0.00077 | 1 | U | K.QMKPLVDFYEQK.G + Oxidation (M) |
| 732 | 168 – 183 | 632.2931 | 1893.8574 | 1893.8891 | -16.7 | 1 | 5 | 0.3 | 1 | U | R.NINGEQDMEKVFADIR.E + Oxidation (M) |
| 733 | 168 – 183 | 632.2962 | 1893.8668 | 1893.8891 | -11.8 | 1 | 3 | 0.47 | 1 | U | R.NINGEQDMEKVFADIR.E + Oxidation (M) |
| 734 | 168 – 183 | 632.2980 | 1893.8723 | 1893.8891 | -8.87 | 1 | 7 | 0.2 | 1 | U | R.NINGEQDMEKVFADIR.E + Oxidation (M) |
| 735 | 168 – 183 | 632.2990 | 1893.8753 | 1893.8891 | -7.27 | 1 | 17 | 0.019 | 1 | U | R.NINGEQDMEKVFADIR.E + Oxidation (M) |
| 737 | 168 – 183 | 632.6277 | 1894.8614 | 1893.8891 | 513 | 1 | 7 | 0.2 | 1 | U | R.NINGEQDMEKVFADIR.E + Oxidation (M) |
| 88 | 184 – 191 | 414.7436 | 827.4726 | 827.4865 | -16.8 | 0 | 29 | 0.0011 | 1 | U | R.ELLGGLAR.A |
| 89 | 184 – 191 | 414.7447 | 827.4749 | 827.4865 | -14.0 | 0 | 11 | 0.089 | 1 | U | R.ELLGGLAR.A |
| 90 | 184 – 191 | 828.4827 | 827.4754 | 827.4865 | -13.4 | 0 | 4 | 0.44 | 1 | U | R.ELLGGLAR.A |
| 91 | 184 – 191 | 414.7477 | 827.4809 | 827.4865 | -6.78 | 0 | 11 | 0.08 | 1 | U | R.ELLGGLAR.A |
| 93 | 184 – 191 | 828.4921 | 827.4848 | 827.4865 | -1.98 | 0 | 16 | 0.027 | 1 | U | R.ELLGGLAR.A |
| 94 | 184 – 191 | 828.4924 | 827.4851 | 827.4865 | -1.65 | 0 | 8 | 0.17 | 1 | U | R.ELLGGLAR.A |
| 95 | 184 – 191 | 828.4942 | 827.4870 | 827.4865 | 0.58 | 0 | 6 | 0.28 | 1 | U | R.ELLGGLAR.A |
| 96 | 184 – 191 | 828.4956 | 827.4884 | 827.4865 | 2.26 | 0 | 3 | 0.52 | 1 | U | R.ELLGGLAR.A |
| 479 | 192 – 207 | 757.4088 | 1512.8030 | 1512.8156 | -8.33 | 0 | 2 | 0.61 | 1 | U | R.AAAMNLVLMGLPGAGK.G |
| 499 | 192 – 207 | 765.4017 | 1528.7888 | 1528.8105 | -14.2 | 0 | 10 | 0.11 | 1 | U | R.AAAMNLVLMGLPGAGK.G + Oxidation (M) |
| 500 | 192 – 207 | 765.4034 | 1528.7921 | 1528.8105 | -12.0 | 0 | 22 | 0.0063 | 1 | U | R.AAAMNLVLMGLPGAGK.G + Oxidation (M) |
| 501 | 192 – 207 | 765.4049 | 1528.7953 | 1528.8105 | -9.96 | 0 | 4 | 0.41 | 1 | U | R.AAAMNLVLMGLPGAGK.G + Oxidation (M) |
| 502 | 192 – 207 | 765.4064 | 1528.7982 | 1528.8105 | -8.09 | 0 | 3 | 0.53 | 1 | U | R.AAAMNLVLMGLPGAGK.G + Oxidation (M) |
| 503 | 192 – 207 | 765.4065 | 1528.7984 | 1528.8105 | -7.93 | 0 | 13 | 0.045 | 1 | U | R.AAAMNLVLMGLPGAGK.G + Oxidation (M) |
| 504 | 192 – 207 | 765.4070 | 1528.7994 | 1528.8105 | -7.30 | 0 | 28 | 0.0015 | 1 | U | R.AAAMNLVLMGLPGAGK.G + Oxidation (M) |
| 506 | 192 – 207 | 765.4078 | 1528.8011 | 1528.8105 | -6.18 | 0 | 19 | 0.013 | 1 | U | R.AAAMNLVLMGLPGAGK.G + Oxidation (M) |
| 507 | 192 – 207 | 765.4079 | 1528.8013 | 1528.8105 | -6.06 | 0 | 10 | 0.1 | 1 | U | R.AAAMNLVLMGLPGAGK.G + Oxidation (M) |
| 508 | 192 – 207 | 765.4082 | 1528.8019 | 1528.8105 | -5.65 | 0 | 30 | 0.0011 | 1 | U | R.AAAMNLVLMGLPGAGK.G + Oxidation (M) |
| 509 | 192 – 207 | 765.4094 | 1528.8042 | 1528.8105 | -4.16 | 0 | 40 | 0.00011 | 1 | U | R.AAAMNLVLMGLPGAGK.G + Oxidation (M) |
| 510 | 192 – 207 | 765.4102 | 1528.8059 | 1528.8105 | -3.04 | 0 | 36 | 0.00026 | 1 | U | R.AAAMNLVLMGLPGAGK.G + Oxidation (M) |
| 511 | 192 – 207 | 765.4134 | 1528.8122 | 1528.8105 | 1.10 | 0 | 24 | 0.0036 | 1 | U | R.AAAMNLVLMGLPGAGK.G + Oxidation (M) |
| 544 | 192 – 207 | 773.3978 | 1544.7811 | 1544.8055 | -15.7 | 0 | 25 | 0.0042 | 1 | U | R.AAAMNLVLMGLPGAGK.G + 2 Oxidation (M) |
| 545 | 192 – 207 | 773.3998 | 1544.7851 | 1544.8055 | -13.2 | 0 | 30 | 0.0013 | 1 | U | R.AAAMNLVLMGLPGAGK.G + 2 Oxidation (M) |
| 546 | 192 – 207 | 773.4024 | 1544.7903 | 1544.8055 | -9.81 | 0 | 12 | 0.086 | 1 | U | R.AAAMNLVLMGLPGAGK.G + 2 Oxidation (M) |
| 547 | 192 – 207 | 773.4029 | 1544.7913 | 1544.8055 | -9.18 | 0 | 36 | 0.00041 | 1 | U | R.AAAMNLVLMGLPGAGK.G + 2 Oxidation (M) |
| 549 | 192 – 207 | 773.4031 | 1544.7917 | 1544.8055 | -8.92 | 0 | 47 | 3e-005 | 1 | U | R.AAAMNLVLMGLPGAGK.G + 2 Oxidation (M) |
| 550 | 192 – 207 | 773.4031 | 1544.7917 | 1544.8055 | -8.92 | 0 | 43 | 8.1e-005 | 1 | U | R.AAAMNLVLMGLPGAGK.G + 2 Oxidation (M) |
| 551 | 192 – 207 | 773.4037 | 1544.7929 | 1544.8055 | -8.11 | 0 | 37 | 0.00033 | 1 | U | R.AAAMNLVLMGLPGAGK.G + 2 Oxidation (M) |
| 552 | 192 – 207 | 773.4037 | 1544.7929 | 1544.8055 | -8.11 | 0 | 41 | 0.00012 | 1 | U | R.AAAMNLVLMGLPGAGK.G + 2 Oxidation (M) |
| 553 | 192 – 207 | 773.4039 | 1544.7932 | 1544.8055 | -7.94 | 0 | 36 | 0.00039 | 1 | U | R.AAAMNLVLMGLPGAGK.G + 2 Oxidation (M) |
| 554 | 192 – 207 | 773.4039 | 1544.7932 | 1544.8055 | -7.94 | 0 | 27 | 0.0029 | 1 | U | R.AAAMNLVLMGLPGAGK.G + 2 Oxidation (M) |
| 555 | 192 – 207 | 773.4042 | 1544.7938 | 1544.8055 | -7.56 | 0 | 36 | 0.00038 | 1 | U | R.AAAMNLVLMGLPGAGK.G + 2 Oxidation (M) |
| 556 | 192 – 207 | 773.4048 | 1544.7950 | 1544.8055 | -6.75 | 0 | 31 | 0.0014 | 1 | U | R.AAAMNLVLMGLPGAGK.G + 2 Oxidation (M) |
| 557 | 192 – 207 | 773.4048 | 1544.7950 | 1544.8055 | -6.75 | 0 | 54 | 6.4e-006 | 1 | U | R.AAAMNLVLMGLPGAGK.G + 2 Oxidation (M) |
| 558 | 192 – 207 | 773.4051 | 1544.7957 | 1544.8055 | -6.29 | 0 | 14 | 0.065 | 1 | U | R.AAAMNLVLMGLPGAGK.G + 2 Oxidation (M) |
| 559 | 192 – 207 | 773.4052 | 1544.7959 | 1544.8055 | -6.20 | 0 | 38 | 0.00022 | 1 | U | R.AAAMNLVLMGLPGAGK.G + 2 Oxidation (M) |
| 560 | 192 – 207 | 773.4052 | 1544.7959 | 1544.8055 | -6.18 | 0 | 26 | 0.0038 | 1 | U | R.AAAMNLVLMGLPGAGK.G + 2 Oxidation (M) |
| 561 | 192 – 207 | 773.4061 | 1544.7976 | 1544.8055 | -5.08 | 0 | 42 | 0.0001 | 1 | U | R.AAAMNLVLMGLPGAGK.G + 2 Oxidation (M) |
| 562 | 192 – 207 | 773.4061 | 1544.7976 | 1544.8055 | -5.08 | 0 | 28 | 0.0025 | 1 | U | R.AAAMNLVLMGLPGAGK.G + 2 Oxidation (M) |

| Query | Start - End | Observed | Mr(expt) | Mr(calc) | ppm | M | Score | Expect | Rank | U | Peptide |
|---|---|---|---|---|---|---|---|---|---|---|---|
| 563 | 192 - 207 | 773.4061 | 1544.7977 | 1544.8055 | -5.02 | 0 | 36 | 0.00042 | 1 | U | R.AAAMNLVLMGLPGAGK.G + 2 Oxidation (M) |
| 564 | 192 - 207 | 773.4077 | 1544.8008 | 1544.8055 | -2.98 | 0 | 35 | 0.00048 | 1 | U | R.AAAMNLVLMGLPGAGK.G + 2 Oxidation (M) |
| 565 | 192 - 207 | 773.4102 | 1544.8058 | 1544.8055 | 0.21 | 0 | 10 | 0.17 | 1 | U | R.AAAMNLVLMGLPGAGK.G + 2 Oxidation (M) |
| 716 | 214 - 230 | 621.9791 | 1862.9155 | 1862.9349 | -10.4 | 0 | 26 | 0.0023 | 1 | U | K.IVAAYGIPHISTGDMFR.A + Oxidation (M) |
| 717 | 214 - 230 | 621.9791 | 1862.9155 | 1862.9349 | -10.4 | 0 | 21 | 0.0077 | 1 | U | K.IVAAYGIPHISTGDMFR.A + Oxidation (M) |
| 718 | 214 - 230 | 621.9797 | 1862.9172 | 1862.9349 | -9.51 | 0 | 10 | 0.092 | 1 | U | K.IVAAYGIPHISTGDMFR.A + Oxidation (M) |
| 719 | 214 - 230 | 621.9803 | 1862.9190 | 1862.9349 | -8.53 | 0 | 8 | 0.15 | 1 | U | K.IVAAYGIPHISTGDMFR.A + Oxidation (M) |
| 720 | 214 - 230 | 621.9806 | 1862.9199 | 1862.9349 | -8.08 | 0 | 20 | 0.011 | 1 | U | K.IVAAYGIPHISTGDMFR.A + Oxidation (M) |
| 721 | 214 - 230 | 621.9809 | 1862.9208 | 1862.9349 | -7.57 | 0 | 23 | 0.0048 | 1 | U | K.IVAAYGIPHISTGDMFR.A + Oxidation (M) |
| 722 | 214 - 230 | 621.9843 | 1862.9311 | 1862.9349 | -2.03 | 0 | 12 | 0.058 | 1 | U | K.IVAAYGIPHISTGDMFR.A + Oxidation (M) |
| 723 | 214 - 230 | 621.9846 | 1862.9321 | 1862.9349 | -1.49 | 0 | 23 | 0.0046 | 1 | U | K.IVAAYGIPHISTGDMFR.A + Oxidation (M) |

Error: try setting browser cache to automatic.

---

```
LOCUS       QNL33370                  37 aa            linear   BCT 04-SEP-2020
DEFINITION  hypothetical protein (plasmid) [Escherichia coli].
ACCESSION   QNL33370
VERSION     QNL33370.1
DBSOURCE    accession MT180430.1
KEYWORDS    .
SOURCE      Escherichia coli
  ORGANISM  Escherichia coli
            Bacteria; Proteobacteria; Gammaproteobacteria; Enterobacterales;
            Enterobacteriaceae; Escherichia.
REFERENCE   1  (residues 1 to 37)
  AUTHORS   Tarabai,H., Wyrsch,E.R., Bitar,I., Djordjevic,S.P. and Dolejska,M.
  TITLE     Comparative analysis of multidrug resistant Escherichia coli ST216
            isolates from silver gulls in Australia
  JOURNAL   Unpublished
REFERENCE   2  (residues 1 to 37)
  AUTHORS   Tarabai,H., Wyrsch,E.R., Bitar,I., Djordjevic,S.P. and Dolejska,M.
  TITLE     Direct Submission
  JOURNAL   Submitted (09-MAR-2020) Department of Biology and Wildlife
            Diseases, University of Veterinary and Pharmaceutical Sciences
            Brno, Palackeho tr. 1946/1, Brno, Brno 61242, Czech Republic
COMMENT     ##Assembly-Data-START##
            Assembly Method       :: SMRT Link v. 8.0
            Assembly Name         :: pCE1681-A
            Coverage              :: 326X
            Sequencing Technology :: PacBio
            ##Assembly-Data-END##
FEATURES             Location/Qualifiers
     source          1..37
                     /organism="Escherichia coli"
                     /strain="CE1681"
                     /host="Chroicocephalus novaehollandiae"
                     /db_xref="taxon:562"
                     /plasmid="pCE1681-A"
                     /country="Australia"
                     /collection_date="2012"
                     /note="Closed IncHI2-ST3/IncN fusion plasmid;
                     type: ST216"
     Protein         1..37
                     /product="hypothetical protein"
     CDS             1..37
                     /coded_by="complement(MT180430.1:247749..247862)"
                     /note="hypothetical protein"
                     /transl_table=11
                     /db_xref="SEED:fig|6666666.506717.peg.327"
```

### Species 4

#### MATRIX SCIENCE MASCOT Search Results

##### Protein View: QNL33370.1

gsAK-cp45 [Escherichia coli]

| | |
|---|---|
| Database: | NCBIcustTC |
| Score: | 859 |
| Monoisotopic mass ($M_r$): | 26627 |
| Calculated pI: | 6.45 |

Sequencesimilarity is availableas an NCBI BLAST search of QNL33370.1 against nr.

##### Search parameters

| | |
|---|---|
| MS data file: | 09182020_TC-6_001.temp.mgf |
| Enzyme: | Trypsin/P: cuts C-term side of KR. |
| Variable modifications: | Oxidation (M), Propionamide (C), Propionamide (K), Propionamide (N-term) |

##### Protein sequence coverage: 38%

Matched peptides shown in **bold red**.

```
  1 MGFRIYRETL SRFSCAAQLG LQAKQYMDRG DLVPDEVTIG IVRERLSKDD
 51 CQNGFLLDGF PRTVAQAEAL ETMLADIGRK LDYVIHIDVR QDVLMERLTG
101 RRICRNCGAT YHLIFHPPAK PGVCDKCGGE LYQRADDNEA TVANRLEVNM
151 KQMKPLVDFY EQKGYLRNIN GEQDMEKVFA DIRELLGGLA RAAAMNLVLM
201 GLPGAGKGTQ AEKIVAAYGI PHISTGDMFR AAMKEGTPLG
```

Unformatted sequence string: 240 residues (for pasting into other applications).

Sort by ◉ residue number ○ increasing mass ○ decreasing mass
Show ◉ matched peptides only ○ predicted peptides also
Show ◉ uncorrected delta ○ delta corrected for 13C

| Query | Start – End | Observed | Mr(expt) | Mr(calc) | ppm | M | Score | Expect | Rank | U | Peptide |
|---|---|---|---|---|---|---|---|---|---|---|---|
| 500 | 25 – 43 | 731.3633 | 2191.0681 | 2191.0943 | -12.0 | 1 | 9 | 0.15 | 1 | U | K.QYMDRGDLVPDEVTIGIVR.E + Oxidation (M) |
| 501 | 25 – 43 | 731.3640 | 2191.0703 | 2191.0943 | -11.0 | 1 | 18 | 0.017 | 1 | U | K.QYMDRGDLVPDEVTIGIVR.E + Oxidation (M) |
| 502 | 25 – 43 | 731.3643 | 2191.0712 | 2191.0943 | -10.5 | 1 | 17 | 0.023 | 1 | U | K.QYMDRGDLVPDEVTIGIVR.E + Oxidation (M) |
| 503 | 25 – 43 | 731.3646 | 2191.0718 | 2191.0943 | -10.3 | 1 | 3 | 0.64 | 1 | U | K.QYMDRGDLVPDEVTIGIVR.E + Oxidation (M) |
| 504 | 25 – 43 | 731.3663 | 2191.0771 | 2191.0943 | -7.85 | 1 | 2 | 0.81 | 1 | U | K.QYMDRGDLVPDEVTIGIVR.E + Oxidation (M) |
| 505 | 25 – 43 | 731.3667 | 2191.0784 | 2191.0943 | -7.27 | 1 | 9 | 0.15 | 1 | U | K.QYMDRGDLVPDEVTIGIVR.E + Oxidation (M) |
| 506 | 25 – 43 | 731.3672 | 2191.0797 | 2191.0943 | -6.70 | 1 | 17 | 0.026 | 1 | U | K.QYMDRGDLVPDEVTIGIVR.E + Oxidation (M) |
| 273 | 30 – 43 | 741.9031 | 1481.7915 | 1481.8090 | -11.8 | 0 | 10 | 0.1 | 1 | U | R.GDLVPDEVTIGIVR.E |
| 274 | 30 – 43 | 741.9044 | 1481.7943 | 1481.8090 | -9.93 | 0 | 1 | 0.78 | 1 | U | R.GDLVPDEVTIGIVR.E |
| 275 | 30 – 43 | 741.9044 | 1481.7943 | 1481.8090 | -9.91 | 0 | 49 | 1.4e-005 | 1 | U | R.GDLVPDEVTIGIVR.E |
| 276 | 30 – 43 | 741.9047 | 1481.7949 | 1481.8090 | -9.49 | 0 | 17 | 0.018 | 1 | U | R.GDLVPDEVTIGIVR.E |
| 277 | 30 – 43 | 741.9051 | 1481.7957 | 1481.8090 | -8.95 | 0 | 25 | 0.0032 | 1 | U | R.GDLVPDEVTIGIVR.E |
| 278 | 30 – 43 | 741.9067 | 1481.7989 | 1481.8090 | -6.81 | 0 | 24 | 0.0039 | 1 | U | R.GDLVPDEVTIGIVR.E |
| 279 | 30 – 43 | 741.9068 | 1481.7990 | 1481.8090 | -6.71 | 0 | 23 | 0.0049 | 1 | U | R.GDLVPDEVTIGIVR.E |
| 280 | 30 – 43 | 741.9070 | 1481.7994 | 1481.8090 | -6.44 | 0 | 55 | 2.9e-006 | 1 | U | R.GDLVPDEVTIGIVR.E |
| 281 | 30 – 43 | 741.9070 | 1481.7994 | 1481.8090 | -6.44 | 0 | 44 | 4.2e-005 | 1 | U | R.GDLVPDEVTIGIVR.E |
| 282 | 30 – 43 | 741.9078 | 1481.8011 | 1481.8090 | -5.34 | 0 | 44 | 4.1e-005 | 1 | U | R.GDLVPDEVTIGIVR.E |
| 283 | 30 – 43 | 741.9078 | 1481.8011 | 1481.8090 | -5.34 | 0 | 43 | 5.4e-005 | 1 | U | R.GDLVPDEVTIGIVR.E |
| 284 | 30 – 43 | 741.9079 | 1481.8012 | 1481.8090 | -5.27 | 0 | 34 | 0.00039 | 1 | U | R.GDLVPDEVTIGIVR.E |
| 285 | 30 – 43 | 741.9079 | 1481.8012 | 1481.8090 | -5.27 | 0 | 29 | 0.0013 | 1 | U | R.GDLVPDEVTIGIVR.E |
| 286 | 30 – 43 | 741.9089 | 1481.8032 | 1481.8090 | -3.91 | 0 | 52 | 6e-006 | 1 | U | R.GDLVPDEVTIGIVR.E |
| 287 | 30 – 43 | 741.9090 | 1481.8034 | 1481.8090 | -3.73 | 0 | 59 | 1.3e-006 | 1 | U | R.GDLVPDEVTIGIVR.E |
| 288 | 30 – 43 | 741.9090 | 1481.8034 | 1481.8090 | -3.73 | 0 | 46 | 2.4e-005 | 1 | U | R.GDLVPDEVTIGIVR.E |
| 370 | 63 – 79 | 894.9550 | 1787.8955 | 1787.9087 | -7.41 | 0 | 17 | 0.021 | 1 | U | R.TVAQAEALETMLADIGR.K |
| 371 | 63 – 79 | 894.9552 | 1787.8958 | 1787.9087 | -7.20 | 0 | 38 | 0.00017 | 1 | U | R.TVAQAEALETMLADIGR.K |
| 372 | 63 – 79 | 602.3003 | 1803.8792 | 1803.9036 | -13.5 | 0 | 2 | 0.73 | 1 | U | R.TVAQAEALETMLADIGR.K + Oxidation (M) |
| 373 | 63 – 79 | 602.3024 | 1803.8853 | 1803.9036 | -10.2 | 0 | 13 | 0.058 | 1 | U | R.TVAQAEALETMLADIGR.K + Oxidation (M) |
| 374 | 63 – 79 | 602.3024 | 1803.8855 | 1803.9036 | -10.1 | 0 | 7 | 0.21 | 1 | U | R.TVAQAEALETMLADIGR.K + Oxidation (M) |

| Query | Start - End | Observed | Mr(expt) | Mr(calc) | ppm | M | Score | Expect | Rank | U | Peptide |
|---|---|---|---|---|---|---|---|---|---|---|---|
| 375 | 63 - 79 | 902.9507 | 1803.8869 | 1803.9036 | -9.28 | 0 | 31 | 0.00079 | 1 | U | R.TVAQAEALETMLADIGR.K + Oxidation (M) |
| 376 | 63 - 79 | 602.3040 | 1803.8903 | 1803.9036 | -7.42 | 0 | 15 | 0.035 | 1 | U | R.TVAQAEALETMLADIGR.K + Oxidation (M) |
| 377 | 63 - 79 | 602.3043 | 1803.8910 | 1803.9036 | -7.03 | 0 | 26 | 0.0025 | 1 | U | R.TVAQAEALETMLADIGR.K + Oxidation (M) |
| 378 | 63 - 79 | 902.9530 | 1803.8915 | 1803.9036 | -6.74 | 0 | 44 | 4.4e-005 | 1 | U | R.TVAQAEALETMLADIGR.K + Oxidation (M) |
| 379 | 63 - 79 | 902.9537 | 1803.8929 | 1803.9036 | -5.97 | 0 | 29 | 0.0012 | 1 | U | R.TVAQAEALETMLADIGR.K + Oxidation (M) |
| 380 | 63 - 79 | 902.9540 | 1803.8934 | 1803.9036 | -5.67 | 0 | 20 | 0.0099 | 1 | U | R.TVAQAEALETMLADIGR.K + Oxidation (M) |
| 381 | 63 - 79 | 602.3051 | 1803.8934 | 1803.9036 | -5.65 | 0 | 31 | 0.00082 | 1 | U | R.TVAQAEALETMLADIGR.K + Oxidation (M) |
| 382 | 63 - 79 | 902.9548 | 1803.8951 | 1803.9036 | -4.74 | 0 | 49 | 1.2e-005 | 1 | U | R.TVAQAEALETMLADIGR.K + Oxidation (M) |
| 383 | 63 - 79 | 902.9565 | 1803.8984 | 1803.9036 | -2.91 | 0 | 29 | 0.0014 | 1 | U | R.TVAQAEALETMLADIGR.K + Oxidation (M) |
| 384 | 63 - 79 | 902.9571 | 1803.8997 | 1803.9036 | -2.20 | 0 | 24 | 0.0041 | 1 | U | R.TVAQAEALETMLADIGR.K + Oxidation (M) |
| 385 | 63 - 79 | 902.9580 | 1803.9014 | 1803.9036 | -1.25 | 0 | 43 | 4.5e-005 | 1 | U | R.TVAQAEALETMLADIGR.K + Oxidation (M) |
| 260 | 80 - 90 | 457.5910 | 1369.7513 | 1369.7718 | -15.0 | 1 | 13 | 0.053 | 1 | U | R.KLDYVIHIDVR.Q |
| 261 | 80 - 90 | 457.5922 | 1369.7547 | 1369.7718 | -12.5 | 1 | 13 | 0.045 | 1 | U | R.KLDYVIHIDVR.Q |
| 265 | 80 - 90 | 457.5936 | 1369.7589 | 1369.7718 | -9.41 | 1 | 12 | 0.065 | 1 | U | R.KLDYVIHIDVR.Q |
| 268 | 80 - 90 | 685.8876 | 1369.7606 | 1369.7718 | -8.15 | 1 | 4 | 0.44 | 1 | U | R.KLDYVIHIDVR.Q |
| 392 | 135 - 151 | 635.9633 | 1904.8681 | 1904.8898 | -11.4 | 1 | 29 | 0.0013 | 1 | U | R.ADDNEATVANRLEVNMK.Q + Oxidation (M) |
| 393 | 135 - 151 | 635.9638 | 1904.8697 | 1904.8898 | -10.5 | 1 | 18 | 0.016 | 1 | U | R.ADDNEATVANRLEVNMK.Q + Oxidation (M) |
| 395 | 135 - 151 | 635.9641 | 1904.8704 | 1904.8898 | -10.2 | 1 | 13 | 0.055 | 1 | U | R.ADDNEATVANRLEVNMK.Q + Oxidation (M) |
| 396 | 135 - 151 | 635.9643 | 1904.8711 | 1904.8898 | -9.78 | 1 | 13 | 0.05 | 1 | U | R.ADDNEATVANRLEVNMK.Q + Oxidation (M) |
| 397 | 135 - 151 | 635.9648 | 1904.8726 | 1904.8898 | -9.00 | 1 | 1 | 0.76 | 1 | U | R.ADDNEATVANRLEVNMK.Q + Oxidation (M) |
| 398 | 135 - 151 | 635.9651 | 1904.8736 | 1904.8898 | -8.49 | 1 | 1 | 0.83 | 1 | U | R.ADDNEATVANRLEVNMK.Q + Oxidation (M) |
| 399 | 135 - 151 | 635.9653 | 1904.8741 | 1904.8898 | -8.23 | 1 | 12 | 0.06 | 1 | U | R.ADDNEATVANRLEVNMK.Q + Oxidation (M) |
| 400 | 135 - 151 | 635.9654 | 1904.8744 | 1904.8898 | -8.07 | 1 | 27 | 0.0022 | 1 | U | R.ADDNEATVANRLEVNMK.Q + Oxidation (M) |
| 401 | 135 - 151 | 635.9658 | 1904.8755 | 1904.8898 | -7.50 | 1 | 2 | 0.63 | 1 | U | R.ADDNEATVANRLEVNMK.Q + Oxidation (M) |
| 404 | 135 - 151 | 953.4492 | 1904.8837 | 1904.8898 | -3.16 | 1 | 2 | 0.68 | 1 | U | R.ADDNEATVANRLEVNMK.Q + Oxidation (M) |
| 310 | 152 - 163 | 771.3765 | 1540.7384 | 1540.7596 | -13.7 | 1 | 1 | 1.5 | 1 | U | K.QMKPLVDFYEQK.G + Oxidation (M) |
| 311 | 152 - 163 | 514.5868 | 1540.7385 | 1540.7596 | -13.7 | 1 | 3 | 1.1 | 1 | U | K.QMKPLVDFYEQK.G + Oxidation (M) |
| 312 | 152 - 163 | 514.5877 | 1540.7414 | 1540.7596 | -11.8 | 1 | 2 | 1.2 | 1 | U | K.QMKPLVDFYEQK.G + Oxidation (M) |
| 313 | 152 - 163 | 771.3796 | 1540.7446 | 1540.7596 | -9.71 | 1 | 4 | 0.8 | 1 | U | K.QMKPLVDFYEQK.G + Oxidation (M) |
| 316 | 152 - 163 | 771.3886 | 1540.7627 | 1540.7596 | 2.02 | 1 | 17 | 0.026 | 1 | U | K.QMKPLVDFYEQK.G + Oxidation (M) |
| 317 | 192 - 207 | 773.4012 | 1544.7878 | 1544.8055 | -11.4 | 0 | 30 | 0.0014 | 1 | U | R.AAAMNLVLMGLPGAGK.G + 2 Oxidation (M) |
| 318 | 192 - 207 | 773.4014 | 1544.7883 | 1544.8055 | -11.1 | 0 | 7 | 0.29 | 1 | U | R.AAAMNLVLMGLPGAGK.G + 2 Oxidation (M) |
| 319 | 192 - 207 | 773.4029 | 1544.7912 | 1544.8055 | -9.21 | 0 | 15 | 0.052 | 1 | U | R.AAAMNLVLMGLPGAGK.G + 2 Oxidation (M) |
| 320 | 192 - 207 | 773.4031 | 1544.7916 | 1544.8055 | -8.95 | 0 | 26 | 0.0041 | 1 | U | R.AAAMNLVLMGLPGAGK.G + 2 Oxidation (M) |
| 321 | 192 - 207 | 773.4034 | 1544.7923 | 1544.8055 | -8.49 | 0 | 37 | 0.0003 | 1 | U | R.AAAMNLVLMGLPGAGK.G + 2 Oxidation (M) |
| 322 | 192 - 207 | 773.4042 | 1544.7938 | 1544.8055 | -7.54 | 0 | 19 | 0.02 | 1 | U | R.AAAMNLVLMGLPGAGK.G + 2 Oxidation (M) |
| 323 | 192 - 207 | 773.4042 | 1544.7938 | 1544.8055 | -7.54 | 0 | 27 | 0.0033 | 1 | U | R.AAAMNLVLMGLPGAGK.G + 2 Oxidation (M) |
| 324 | 192 - 207 | 773.4049 | 1544.7953 | 1544.8055 | -6.58 | 0 | 40 | 0.00014 | 1 | U | R.AAAMNLVLMGLPGAGK.G + 2 Oxidation (M) |
| 325 | 192 - 207 | 773.4050 | 1544.7955 | 1544.8055 | -6.41 | 0 | 29 | 0.002 | 1 | U | R.AAAMNLVLMGLPGAGK.G + 2 Oxidation (M) |
| 326 | 192 - 207 | 773.4050 | 1544.7955 | 1544.8055 | -6.41 | 0 | 37 | 0.00033 | 1 | U | R.AAAMNLVLMGLPGAGK.G + 2 Oxidation (M) |
| 327 | 192 - 207 | 773.4051 | 1544.7957 | 1544.8055 | -6.32 | 0 | 16 | 0.037 | 1 | U | R.AAAMNLVLMGLPGAGK.G + 2 Oxidation (M) |
| 328 | 192 - 207 | 773.4052 | 1544.7959 | 1544.8055 | -6.19 | 0 | 33 | 0.00075 | 1 | U | R.AAAMNLVLMGLPGAGK.G + 2 Oxidation (M) |
| 329 | 192 - 207 | 773.4054 | 1544.7963 | 1544.8055 | -5.89 | 0 | 33 | 0.00073 | 1 | U | R.AAAMNLVLMGLPGAGK.G + 2 Oxidation (M) |
| 330 | 192 - 207 | 773.4054 | 1544.7963 | 1544.8055 | -5.89 | 0 | 23 | 0.0073 | 1 | U | R.AAAMNLVLMGLPGAGK.G + 2 Oxidation (M) |
| 331 | 192 - 207 | 773.4059 | 1544.7972 | 1544.8055 | -5.36 | 0 | 36 | 0.00038 | 1 | U | R.AAAMNLVLMGLPGAGK.G + 2 Oxidation (M) |

Error: try setting browser cache to automatic.

---

```
LOCUS       QNL33370                  37 aa            linear   BCT 04-SEP-2020
DEFINITION  hypothetical protein (plasmid) [Escherichia coli].
ACCESSION   QNL33370
VERSION     QNL33370.1
DBSOURCE    accession MT180430.1
KEYWORDS    .
SOURCE      Escherichia coli
  ORGANISM  Escherichia coli
            Bacteria; Proteobacteria; Gammaproteobacteria; Enterobacterales;
            Enterobacteriaceae; Escherichia.
REFERENCE   1  (residues 1 to 37)
```

```
  AUTHORS   Tarabai,H., Wyrsch,E.R., Bitar,I., Djordjevic,S.P. and Dolejska,M.
  TITLE     Comparative analysis of multidrug resistant Escherichia coli ST216
            isolates from silver gulls in Australia
  JOURNAL   Unpublished
REFERENCE   2  (residues 1 to 37)
  AUTHORS   Tarabai,H., Wyrsch,E.R., Bitar,I., Djordjevic,S.P. and Dolejska,M.
  TITLE     Direct Submission
  JOURNAL   Submitted (09-MAR-2020) Department of Biology and Wildlife
            Diseases, University of Veterinary and Pharmaceutical Sciences
            Brno, Palackeho tr. 1946/1, Brno, Brno 61242, Czech Republic
COMMENT     ##Assembly-Data-START##
            Assembly Method       :: SMRT Link v. 8.0
            Assembly Name         :: pCE1681-A
            Coverage              :: 326X
            Sequencing Technology :: PacBio
            ##Assembly-Data-END##
FEATURES             Location/Qualifiers
     source          1..37
                     /organism="Escherichia coli"
                     /strain="CE1681"
                     /host="Chroicocephalus novaehollandiae"
                     /db_xref="taxon:562"
                     /plasmid="pCE1681-A"
                     /country="Australia"
                     /collection_date="2012"
                     /note="Closed IncHI2-ST3/IncN fusion plasmid;
                     type: ST216"
     Protein         1..37
                     /product="hypothetical protein"
     CDS             1..37
                     /coded_by="complement(MT180430.1:247749..247862)"
                     /note="hypothetical protein"
                     /transl_table=11
                     /db_xref="SEED:fig|6666666.506717.peg.327"
```

### Species 5

#### MATRIX SCIENCE MASCOT Search Results

##### Protein View: QNL33370.1

gsAK-cp45 [Escherichia coli]

| | |
|---|---|
| Database: | NCBIcustTC |
| Score: | 1417 |
| Monoisotopic mass ($M_r$): | 26627 |
| Calculated pI: | 6.45 |

Sequence similarity is available as an NCBI BLAST search of QNL33370.1 against nr.

##### Search parameters

| | |
|---|---|
| MS data file: | 09182020_TC-7_001.temp.mgf |
| Enzyme: | Trypsin/P: cuts C-term side of KR. |
| Variable modifications: | Oxidation (M), Propionamide (C), Propionamide (K), Propionamide (N-term) |

##### Protein sequence coverage: 44%

Matched peptides shown in **bold red**.

```
  1 MGFRIYRETL SRFSCAAQLG LQAKQYMDRG DLVPDEVTIG IVRERLSKDD
 51 CQNGFLLDGF PRTVAQAEAL ETMLADIGRK LDYVIHIDVR QDVLMERLTG
101 RRICRNCGAT YHLIFHPPAK PGVCDKCGGE LYQRADDNEA TVANRLEVNM
151 KQMKPLVDFY EQKGYLRNIN GEQDMEKVFA DIRELLGGLA RAAAMNLVLM
201 GLPGAGKGTQ AEKIVAAYGI PHISTGDMFR AAMKEGTPLG
```

Unformatted sequence string: 240 residues (for pasting into other applications).

Sort by ◉ residue number ○ increasing mass ○ decreasing mass
Show ◉ matched peptides only ○ predicted peptides also
Show ◉ uncorrected delta ○ delta corrected for 13C

| Query | Start - End | Observed | Mr(expt) | Mr(calc) | ppm | M | Score | Expect | Rank | U | Peptide |
|---|---|---|---|---|---|---|---|---|---|---|---|
| 572 | 25 - 43 | 731.3637 | 2191.0692 | 2191.0943 | -11.5 | 1 | 4 | 0.43 | 1 | U | K.QYMDRGDLVPDEVTIGIVR.E + Oxidation (M) |
| 573 | 25 - 43 | 731.3655 | 2191.0746 | 2191.0943 | -9.01 | 1 | 7 | 0.26 | 1 | U | K.QYMDRGDLVPDEVTIGIVR.E + Oxidation (M) |
| 574 | 25 - 43 | 731.3655 | 2191.0747 | 2191.0943 | -8.94 | 1 | 21 | 0.01 | 1 | U | K.QYMDRGDLVPDEVTIGIVR.E + Oxidation (M) |
| 575 | 25 - 43 | 731.3659 | 2191.0758 | 2191.0943 | -8.48 | 1 | 21 | 0.01 | 1 | U | K.QYMDRGDLVPDEVTIGIVR.E + Oxidation (M) |
| 576 | 25 - 43 | 731.3660 | 2191.0763 | 2191.0943 | -8.22 | 1 | 28 | 0.0022 | 1 | U | K.QYMDRGDLVPDEVTIGIVR.E + Oxidation (M) |
| 577 | 25 - 43 | 731.3668 | 2191.0785 | 2191.0943 | -7.25 | 1 | 22 | 0.0091 | 1 | U | K.QYMDRGDLVPDEVTIGIVR.E + Oxidation (M) |
| 578 | 25 - 43 | 731.3671 | 2191.0795 | 2191.0943 | -6.78 | 1 | 38 | 0.00022 | 1 | U | K.QYMDRGDLVPDEVTIGIVR.E + Oxidation (M) |
| 579 | 25 - 43 | 731.3675 | 2191.0807 | 2191.0943 | -6.23 | 1 | 36 | 0.00035 | 1 | U | K.QYMDRGDLVPDEVTIGIVR.E + Oxidation (M) |
| 580 | 25 - 43 | 731.3698 | 2191.0875 | 2191.0943 | -3.14 | 1 | 16 | 0.033 | 1 | U | K.QYMDRGDLVPDEVTIGIVR.E + Oxidation (M) |
| 287 | 30 - 43 | 741.9060 | 1481.7975 | 1481.8090 | -7.74 | 0 | 29 | 0.0012 | 1 | U | R.GDLVPDEVTIGIVR.E |
| 288 | 30 - 43 | 741.9064 | 1481.7983 | 1481.8090 | -7.20 | 0 | 38 | 0.00017 | 1 | U | R.GDLVPDEVTIGIVR.E |
| 289 | 30 - 43 | 741.9074 | 1481.8003 | 1481.8090 | -5.85 | 0 | 32 | 0.00059 | 1 | U | R.GDLVPDEVTIGIVR.E |
| 290 | 30 - 43 | 741.9077 | 1481.8009 | 1481.8090 | -5.43 | 0 | 48 | 1.7e-005 | 1 | U | R.GDLVPDEVTIGIVR.E |
| 291 | 30 - 43 | 741.9078 | 1481.8010 | 1481.8090 | -5.38 | 0 | 66 | 2.7e-007 | 1 | U | R.GDLVPDEVTIGIVR.E |
| 294 | 30 - 43 | 741.9084 | 1481.8022 | 1481.8090 | -4.57 | 0 | 1 | 0.82 | 1 | U | R.GDLVPDEVTIGIVR.E |
| 295 | 30 - 43 | 741.9084 | 1481.8023 | 1481.8090 | -4.51 | 0 | 42 | 5.6e-005 | 1 | U | R.GDLVPDEVTIGIVR.E |
| 296 | 30 - 43 | 741.9087 | 1481.8028 | 1481.8090 | -4.15 | 0 | 50 | 9.2e-006 | 1 | U | R.GDLVPDEVTIGIVR.E |
| 298 | 30 - 43 | 741.9090 | 1481.8034 | 1481.8090 | -3.76 | 0 | 60 | 9e-007 | 1 | U | R.GDLVPDEVTIGIVR.E |
| 299 | 30 - 43 | 741.9091 | 1481.8036 | 1481.8090 | -3.65 | 0 | 53 | 4.9e-006 | 1 | U | R.GDLVPDEVTIGIVR.E |

| Query | Start – End | Observed | Mr(expt) | Mr(calc) | ppm | M | Score | Expect | Rank | U | Peptide |
|---|---|---|---|---|---|---|---|---|---|---|---|
| 300 | 30 – 43 | 741.9098 | 1481.8050 | 1481.8090 | -2.66 | 0 | 43 | 4.8e-005 | 1 | U | R.GDLVPDEVTIGIVR.E |
| 301 | 30 – 43 | 741.9136 | 1481.8126 | 1481.8090 | 2.42 | 0 | 11 | 0.081 | 1 | U | R.GDLVPDEVTIGIVR.E |
| 388 | 63 – 79 | 894.9566 | 1787.8986 | 1787.9087 | -5.66 | 0 | 52 | 6.6e-006 | 1 | U | R.TVAQAEALETMLADIGR.K |
| 389 | 63 – 79 | 894.9569 | 1787.8992 | 1787.9087 | -5.30 | 0 | 17 | 0.018 | 1 | U | R.TVAQAEALETMLADIGR.K |
| 391 | 63 – 79 | 894.9609 | 1787.9072 | 1787.9087 | -0.86 | 0 | 70 | 1e-007 | 1 | U | R.TVAQAEALETMLADIGR.K |
| 392 | 63 – 79 | 602.2977 | 1803.8714 | 1803.9036 | -17.9 | 0 | 20 | 0.011 | 1 | U | R.TVAQAEALETMLADIGR.K + Oxidation (M) |
| 393 | 63 – 79 | 602.3001 | 1803.8786 | 1803.9036 | -13.9 | 0 | 11 | 0.075 | 1 | U | R.TVAQAEALETMLADIGR.K + Oxidation (M) |
| 394 | 63 – 79 | 602.3015 | 1803.8827 | 1803.9036 | -11.6 | 0 | 39 | 0.00012 | 1 | U | R.TVAQAEALETMLADIGR.K + Oxidation (M) |
| 395 | 63 – 79 | 902.9505 | 1803.8865 | 1803.9036 | -9.51 | 0 | 24 | 0.0042 | 1 | U | R.TVAQAEALETMLADIGR.K + Oxidation (M) |
| 396 | 63 – 79 | 602.3031 | 1803.8875 | 1803.9036 | -8.93 | 0 | 28 | 0.0015 | 1 | U | R.TVAQAEALETMLADIGR.K + Oxidation (M) |
| 397 | 63 – 79 | 902.9518 | 1803.8891 | 1803.9036 | -8.06 | 0 | 59 | 1.2e-006 | 1 | U | R.TVAQAEALETMLADIGR.K + Oxidation (M) |
| 398 | 63 – 79 | 602.3044 | 1803.8914 | 1803.9036 | -6.77 | 0 | 22 | 0.0059 | 1 | U | R.TVAQAEALETMLADIGR.K + Oxidation (M) |
| 399 | 63 – 79 | 902.9536 | 1803.8926 | 1803.9036 | -6.13 | 0 | 23 | 0.0047 | 1 | U | R.TVAQAEALETMLADIGR.K + Oxidation (M) |
| 400 | 63 – 79 | 902.9537 | 1803.8929 | 1803.9036 | -5.95 | 0 | 25 | 0.0031 | 1 | U | R.TVAQAEALETMLADIGR.K + Oxidation (M) |
| 401 | 63 – 79 | 902.9540 | 1803.8934 | 1803.9036 | -5.69 | 0 | 47 | 1.8e-005 | 1 | U | R.TVAQAEALETMLADIGR.K + Oxidation (M) |
| 402 | 63 – 79 | 902.9540 | 1803.8934 | 1803.9036 | -5.69 | 0 | 85 | 3.2e-009 | 1 | U | R.TVAQAEALETMLADIGR.K + Oxidation (M) |
| 403 | 63 – 79 | 902.9545 | 1803.8945 | 1803.9036 | -5.07 | 0 | 7 | 0.22 | 1 | U | R.TVAQAEALETMLADIGR.K + Oxidation (M) |
| 404 | 63 – 79 | 902.9549 | 1803.8952 | 1803.9036 | -4.65 | 0 | 59 | 1.4e-006 | 1 | U | R.TVAQAEALETMLADIGR.K + Oxidation (M) |
| 405 | 63 – 79 | 902.9549 | 1803.8952 | 1803.9036 | -4.65 | 0 | 48 | 1.7e-005 | 1 | U | R.TVAQAEALETMLADIGR.K + Oxidation (M) |
| 406 | 63 – 79 | 902.9559 | 1803.8972 | 1803.9036 | -3.57 | 0 | 60 | 1e-006 | 1 | U | R.TVAQAEALETMLADIGR.K + Oxidation (M) |
| 407 | 63 – 79 | 902.9560 | 1803.8974 | 1803.9036 | -3.46 | 0 | 21 | 0.0077 | 1 | U | R.TVAQAEALETMLADIGR.K + Oxidation (M) |
| 408 | 63 – 79 | 902.9561 | 1803.8976 | 1803.9036 | -3.37 | 0 | 66 | 2.7e-007 | 1 | U | R.TVAQAEALETMLADIGR.K + Oxidation (M) |
| 409 | 63 – 79 | 602.3065 | 1803.8977 | 1803.9036 | -3.31 | 0 | 17 | 0.02 | 1 | U | R.TVAQAEALETMLADIGR.K + Oxidation (M) |
| 410 | 63 – 79 | 902.9579 | 1803.9013 | 1803.9036 | -1.28 | 0 | 60 | 9.5e-007 | 1 | U | R.TVAQAEALETMLADIGR.K + Oxidation (M) |
| 411 | 63 – 79 | 902.9579 | 1803.9013 | 1803.9036 | -1.28 | 0 | 72 | 6.4e-008 | 1 | U | R.TVAQAEALETMLADIGR.K + Oxidation (M) |
| 255 | 80 – 90 | 457.5899 | 1369.7480 | 1369.7718 | -17.3 | 1 | 4 | 0.37 | 1 | U | R.KLDYVIHIDVR.Q |
| 256 | 80 – 90 | 457.5917 | 1369.7531 | 1369.7718 | -13.6 | 1 | 4 | 0.37 | 1 | U | R.KLDYVIHIDVR.Q |
| 257 | 80 – 90 | 457.5919 | 1369.7537 | 1369.7718 | -13.2 | 1 | 3 | 0.46 | 1 | U | R.KLDYVIHIDVR.Q |
| 258 | 80 – 90 | 457.5920 | 1369.7540 | 1369.7718 | -13.0 | 1 | 24 | 0.004 | 1 | U | R.KLDYVIHIDVR.Q |
| 260 | 80 – 90 | 457.5923 | 1369.7550 | 1369.7718 | -12.2 | 1 | 19 | 0.012 | 1 | U | R.KLDYVIHIDVR.Q |
| 261 | 80 – 90 | 457.5928 | 1369.7567 | 1369.7718 | -11.0 | 1 | 20 | 0.01 | 1 | U | R.KLDYVIHIDVR.Q |
| 262 | 80 – 90 | 457.5929 | 1369.7570 | 1369.7718 | -10.8 | 1 | 4 | 0.43 | 1 | U | R.KLDYVIHIDVR.Q |
| 263 | 80 – 90 | 457.5931 | 1369.7576 | 1369.7718 | -10.4 | 1 | 6 | 0.27 | 1 | U | R.KLDYVIHIDVR.Q |
| 264 | 80 – 90 | 457.5936 | 1369.7591 | 1369.7718 | -9.26 | 1 | 18 | 0.015 | 1 | U | R.KLDYVIHIDVR.Q |
| 266 | 80 – 90 | 457.5940 | 1369.7602 | 1369.7718 | -8.47 | 1 | 6 | 0.25 | 1 | U | R.KLDYVIHIDVR.Q |
| 267 | 80 – 90 | 685.8875 | 1369.7604 | 1369.7718 | -8.30 | 1 | 17 | 0.023 | 1 | U | R.KLDYVIHIDVR.Q |
| 268 | 80 – 90 | 685.8876 | 1369.7607 | 1369.7718 | -8.06 | 1 | 23 | 0.0058 | 1 | U | R.KLDYVIHIDVR.Q |
| 269 | 80 – 90 | 457.5942 | 1369.7607 | 1369.7718 | -8.05 | 1 | 12 | 0.073 | 1 | U | R.KLDYVIHIDVR.Q |
| 270 | 80 – 90 | 685.8877 | 1369.7608 | 1369.7718 | -8.03 | 1 | 31 | 0.0009 | 1 | U | R.KLDYVIHIDVR.Q |
| 272 | 80 – 90 | 457.5944 | 1369.7613 | 1369.7718 | -7.68 | 1 | 4 | 0.46 | 1 | U | R.KLDYVIHIDVR.Q |
| 273 | 80 – 90 | 685.8880 | 1369.7614 | 1369.7718 | -7.54 | 1 | 23 | 0.0057 | 1 | U | R.KLDYVIHIDVR.Q |
| 274 | 80 – 90 | 685.8882 | 1369.7618 | 1369.7718 | -7.28 | 1 | 18 | 0.017 | 1 | U | R.KLDYVIHIDVR.Q |
| 276 | 80 – 90 | 457.5949 | 1369.7628 | 1369.7718 | -6.56 | 1 | 3 | 0.48 | 1 | U | R.KLDYVIHIDVR.Q |
| 277 | 80 – 90 | 685.8887 | 1369.7628 | 1369.7718 | -6.55 | 1 | 5 | 0.32 | 1 | U | R.KLDYVIHIDVR.Q |
| 278 | 80 – 90 | 457.5949 | 1369.7630 | 1369.7718 | -6.41 | 1 | 22 | 0.0071 | 1 | U | R.KLDYVIHIDVR.Q |
| 279 | 80 – 90 | 457.5953 | 1369.7640 | 1369.7718 | -5.69 | 1 | 12 | 0.067 | 1 | U | R.KLDYVIHIDVR.Q |
| 280 | 80 – 90 | 685.8893 | 1369.7641 | 1369.7718 | -5.60 | 1 | 24 | 0.0044 | 1 | U | R.KLDYVIHIDVR.Q |
| 281 | 80 – 90 | 685.8895 | 1369.7645 | 1369.7718 | -5.29 | 1 | 33 | 0.00057 | 1 | U | R.KLDYVIHIDVR.Q |
| 282 | 80 – 90 | 685.8896 | 1369.7647 | 1369.7718 | -5.19 | 1 | 41 | 7.5e-005 | 1 | U | R.KLDYVIHIDVR.Q |
| 285 | 80 – 90 | 685.8938 | 1369.7730 | 1369.7718 | 0.90 | 1 | 6 | 0.27 | 1 | U | R.KLDYVIHIDVR.Q |
| 594 | 80 – 97 | 753.4001 | 2257.1785 | 2257.1889 | -4.59 | 2 | 1 | 0.9 | 1 | U | R.KLDYVIHIDVRQDVLMER.L + Oxidation (M) |
| 135 | 91 – 97 | 453.7145 | 905.4144 | 905.4277 | -14.6 | 0 | 19 | 0.013 | 1 | U | R.QDVLMER.L + Oxidation (M) |
| 136 | 91 – 97 | 453.7172 | 905.4198 | 905.4277 | -8.67 | 0 | 7 | 0.21 | 1 | U | R.QDVLMER.L + Oxidation (M) |
| 137 | 91 – 97 | 453.7172 | 905.4199 | 905.4277 | -8.58 | 0 | 3 | 0.5 | 1 | U | R.QDVLMER.L + Oxidation (M) |
| 138 | 91 – 97 | 453.7181 | 905.4216 | 905.4277 | -6.73 | 0 | 15 | 0.034 | 1 | U | R.QDVLMER.L + Oxidation (M) |
| 139 | 91 – 97 | 453.7193 | 905.4240 | 905.4277 | -4.03 | 0 | 11 | 0.074 | 1 | U | R.QDVLMER.L + Oxidation (M) |
| 140 | 91 – 97 | 453.7194 | 905.4242 | 905.4277 | -3.81 | 0 | 12 | 0.064 | 1 | U | R.QDVLMER.L + Oxidation (M) |

| Query | Start - End | Observed | Mr(expt) | Mr(calc) | ppm | M | Score | Expect | Rank | U | Peptide |
|---|---|---|---|---|---|---|---|---|---|---|---|
| 141 | 91 - 97 | 453.7196 | 905.4246 | 905.4277 | -3.39 | 0 | 5 | 0.33 | 1 | U | R.QDVLMER.L + Oxidation (M) |
| 142 | 91 - 97 | 453.7201 | 905.4256 | 905.4277 | -2.22 | 0 | 26 | 0.0025 | 1 | U | R.QDVLMER.L + Oxidation (M) |
| 238 | 135 - 145 | 588.2623 | 1174.5101 | 1174.5214 | -9.63 | 0 | 0 | 0.94 | 1 | U | R.ADDNEATVANR.L |
| 239 | 135 - 145 | 588.2625 | 1174.5104 | 1174.5214 | -9.40 | 0 | 24 | 0.0041 | 1 | U | R.ADDNEATVANR.L |
| 240 | 135 - 145 | 588.2626 | 1174.5107 | 1174.5214 | -9.16 | 0 | 4 | 0.36 | 1 | U | R.ADDNEATVANR.L |
| 241 | 135 - 145 | 588.2643 | 1174.5141 | 1174.5214 | -6.25 | 0 | 11 | 0.086 | 1 | U | R.ADDNEATVANR.L |
| 243 | 135 - 145 | 588.2664 | 1174.5182 | 1174.5214 | -2.75 | 0 | 8 | 0.16 | 1 | U | R.ADDNEATVANR.L |
| 244 | 135 - 145 | 588.2685 | 1174.5224 | 1174.5214 | 0.84 | 0 | 14 | 0.041 | 1 | U | R.ADDNEATVANR.L |
| 439 | 135 - 151 | 635.9627 | 1904.8663 | 1904.8898 | -12.3 | 1 | 26 | 0.0027 | 1 | U | R.ADDNEATVANRLEVNMK.Q + Oxidation (M) |
| 440 | 135 - 151 | 635.9630 | 1904.8673 | 1904.8898 | -11.8 | 1 | 15 | 0.034 | 1 | U | R.ADDNEATVANRLEVNMK.Q + Oxidation (M) |
| 441 | 135 - 151 | 635.9635 | 1904.8686 | 1904.8898 | -11.1 | 1 | 30 | 0.0011 | 1 | U | R.ADDNEATVANRLEVNMK.Q + Oxidation (M) |
| 442 | 135 - 151 | 635.9638 | 1904.8695 | 1904.8898 | -10.7 | 1 | 12 | 0.072 | 1 | U | R.ADDNEATVANRLEVNMK.Q + Oxidation (M) |
| 443 | 135 - 151 | 953.4425 | 1904.8705 | 1904.8898 | -10.1 | 1 | 6 | 0.28 | 1 | U | R.ADDNEATVANRLEVNMK.Q + Oxidation (M) |
| 444 | 135 - 151 | 635.9644 | 1904.8713 | 1904.8898 | -9.69 | 1 | 30 | 0.0011 | 1 | U | R.ADDNEATVANRLEVNMK.Q + Oxidation (M) |
| 445 | 135 - 151 | 635.9651 | 1904.8735 | 1904.8898 | -8.56 | 1 | 13 | 0.058 | 1 | U | R.ADDNEATVANRLEVNMK.Q + Oxidation (M) |
| 446 | 135 - 151 | 635.9653 | 1904.8740 | 1904.8898 | -8.29 | 1 | 31 | 0.0009 | 1 | U | R.ADDNEATVANRLEVNMK.Q + Oxidation (M) |
| 448 | 135 - 151 | 953.4453 | 1904.8761 | 1904.8898 | -7.17 | 1 | 12 | 0.063 | 1 | U | R.ADDNEATVANRLEVNMK.Q + Oxidation (M) |
| 449 | 135 - 151 | 635.9662 | 1904.8768 | 1904.8898 | -6.79 | 1 | 44 | 4.7e-005 | 1 | U | R.ADDNEATVANRLEVNMK.Q + Oxidation (M) |
| 450 | 135 - 151 | 635.9664 | 1904.8773 | 1904.8898 | -6.54 | 1 | 39 | 0.00012 | 1 | U | R.ADDNEATVANRLEVNMK.Q + Oxidation (M) |
| 451 | 135 - 151 | 953.4461 | 1904.8777 | 1904.8898 | -6.31 | 1 | 11 | 0.094 | 1 | U | R.ADDNEATVANRLEVNMK.Q + Oxidation (M) |
| 453 | 135 - 151 | 635.9667 | 1904.8782 | 1904.8898 | -6.08 | 1 | 33 | 0.00052 | 1 | U | R.ADDNEATVANRLEVNMK.Q + Oxidation (M) |
| 454 | 135 - 151 | 953.4464 | 1904.8782 | 1904.8898 | -6.05 | 1 | 16 | 0.028 | 1 | U | R.ADDNEATVANRLEVNMK.Q + Oxidation (M) |
| 455 | 135 - 151 | 635.9672 | 1904.8797 | 1904.8898 | -5.30 | 1 | 19 | 0.016 | 1 | U | R.ADDNEATVANRLEVNMK.Q + Oxidation (M) |
| 338 | 152 - 163 | 514.5866 | 1540.7379 | 1540.7596 | -14.1 | 1 | 0 | 2 | 1 | U | K.QMKPLVDFYEQK.G + Oxidation (M) |
| 341 | 152 - 163 | 514.5881 | 1540.7424 | 1540.7596 | -11.1 | 1 | 6 | 0.46 | 1 | U | K.QMKPLVDFYEQK.G + Oxidation (M) |
| 342 | 152 - 163 | 771.3786 | 1540.7427 | 1540.7596 | -10.9 | 1 | 19 | 0.022 | 1 | U | K.QMKPLVDFYEQK.G + Oxidation (M) |
| 345 | 152 - 163 | 514.5893 | 1540.7460 | 1540.7596 | -8.77 | 1 | 7 | 0.42 | 1 | U | K.QMKPLVDFYEQK.G + Oxidation (M) |
| 347 | 152 - 163 | 771.3812 | 1540.7478 | 1540.7596 | -7.61 | 1 | 14 | 0.08 | 1 | U | K.QMKPLVDFYEQK.G + Oxidation (M) |
| 348 | 152 - 163 | 514.5899 | 1540.7480 | 1540.7596 | -7.52 | 1 | 16 | 0.05 | 1 | U | K.QMKPLVDFYEQK.G + Oxidation (M) |
| 349 | 152 - 163 | 771.3815 | 1540.7485 | 1540.7596 | -7.14 | 1 | 2 | 1.3 | 1 | U | K.QMKPLVDFYEQK.G + Oxidation (M) |
| 350 | 152 - 163 | 771.3827 | 1540.7508 | 1540.7596 | -5.65 | 1 | 10 | 0.22 | 1 | U | K.QMKPLVDFYEQK.G + Oxidation (M) |
| 351 | 152 - 163 | 771.3835 | 1540.7523 | 1540.7596 | -4.68 | 1 | 24 | 0.0068 | 1 | U | K.QMKPLVDFYEQK.G + Oxidation (M) |
| 352 | 152 - 163 | 771.3845 | 1540.7545 | 1540.7596 | -3.26 | 1 | 21 | 0.013 | 1 | U | K.QMKPLVDFYEQK.G + Oxidation (M) |
| 353 | 152 - 163 | 771.3849 | 1540.7552 | 1540.7596 | -2.85 | 1 | 14 | 0.059 | 1 | U | K.QMKPLVDFYEQK.G + Oxidation (M) |
| 354 | 152 - 163 | 771.3901 | 1540.7656 | 1540.7596 | 3.90 | 1 | 5 | 0.47 | 1 | U | K.QMKPLVDFYEQK.G + Oxidation (M) |
| 422 | 168 - 183 | 632.3038 | 1893.8897 | 1893.8891 | 0.32 | 1 | 5 | 0.31 | 1 | U | R.NINGEQDMEKVFADIR.E + Oxidation (M) |
| 423 | 168 - 183 | 632.6251 | 1894.8534 | 1893.8891 | 509 | 1 | 4 | 0.4 | 1 | U | R.NINGEQDMEKVFADIR.E + Oxidation (M) |
| 95 | 184 - 191 | 828.4891 | 827.4818 | 827.4865 | -5.64 | 0 | 6 | 0.27 | 1 | U | R.ELLGGLAR.A |
| 96 | 184 - 191 | 828.4898 | 827.4826 | 827.4865 | -4.75 | 0 | 13 | 0.049 | 1 | U | R.ELLGGLAR.A |
| 97 | 184 - 191 | 828.4956 | 827.4883 | 827.4865 | 2.22 | 0 | 6 | 0.22 | 1 | U | R.ELLGGLAR.A |

Error: try setting browser cache to automatic.

---

```
LOCUS       QNL33370                  37 aa            linear   BCT 04-SEP-2020
DEFINITION  hypothetical protein (plasmid) [Escherichia coli].
ACCESSION   QNL33370
VERSION     QNL33370.1
DBSOURCE    accession MT180430.1
KEYWORDS    .
SOURCE      Escherichia coli
```

```
  ORGANISM  Escherichia coli
            Bacteria; Proteobacteria; Gammaproteobacteria; Enterobacterales;
            Enterobacteriaceae; Escherichia.
REFERENCE   1  (residues 1 to 37)
  AUTHORS   Tarabai,H., Wyrsch,E.R., Bitar,I., Djordjevic,S.P. and Dolejska,M.
  TITLE     Comparative analysis of multidrug resistant Escherichia coli ST216
            isolates from silver gulls in Australia
  JOURNAL   Unpublished
REFERENCE   2  (residues 1 to 37)
  AUTHORS   Tarabai,H., Wyrsch,E.R., Bitar,I., Djordjevic,S.P. and Dolejska,M.
  TITLE     Direct Submission
  JOURNAL   Submitted (09-MAR-2020) Department of Biology and Wildlife
            Diseases, University of Veterinary and Pharmaceutical Sciences
            Brno, Palackeho tr. 1946/1, Brno, Brno 61242, Czech Republic
COMMENT     ##Assembly-Data-START##
            Assembly Method       :: SMRT Link v. 8.0
            Assembly Name         :: pCE1681-A
            Coverage              :: 326X
            Sequencing Technology :: PacBio
            ##Assembly-Data-END##
FEATURES             Location/Qualifiers
     source          1..37
                     /organism="Escherichia coli"
                     /strain="CE1681"
                     /host="Chroicocephalus novaehollandiae"
                     /db_xref="taxon:562"
                     /plasmid="pCE1681-A"
                     /country="Australia"
                     /collection_date="2012"
                     /note="Closed IncHI2-ST3/IncN fusion plasmid;
                     type: ST216"
     Protein         1..37
                     /product="hypothetical protein"
     CDS             1..37
                     /coded_by="complement(MT180430.1:247749..247862)"
                     /note="hypothetical protein"
                     /transl_table=11
                     /db_xref="SEED:fig|6666666.506717.peg.327"
```

### Species 6

#### MATRIX SCIENCE MASCOT Search Results

##### Protein View: QNL33370.1

gsAK-cp45 [Escherichia coli]

| | |
|---|---|
| Database: | NCBIcustTC |
| Score: | 223 |
| Monoisotopic mass ($M_r$): | 26627 |
| Calculated pI: | 6.45 |

Sequence similarity is available as an NCBI BLAST search of QNL33370.1 against nr.

##### Search parameters

| | |
|---|---|
| MS data file: | 09182020_TC-5_001.temp.mgf |
| Enzyme: | Trypsin/P: cuts C-term side of KR. |
| Variable modifications: | Oxidation (M), Propionamide (C), Propionamide (K), Propionamide (N-term) |

##### Protein sequence coverage: 38%

Matched peptides shown in **bold red**.

```
  1 MGFRIYRETL SRFSCAAQLG LQAKQYMDRG DLVPDEVTIG IVRERLSKDD
 51 CQNGFLLDGF PRTVAQAEAL ETMLADIGRK LDYVIHIDVR QDVLMERLTG
101 RRICRNCGAT YHLIFHPPAK PGVCDKCGGE LYQRADDNEA TVANRLEVNM
151 KQMKPLVDFY EQKGYLRNIN GEQDMEKVFA DIRELLGGLA RAAAMNLVLM
201 GLPGAGKGTQ AEKIVAAYGI PHISTGDMFR AAMKEGTPLG
```

Unformatted sequence string: 240 residues (for pasting into other applications).

Sort by ◉ residue number ○ increasing mass ○ decreasing mass
Show ◉ matched peptides only ○ predicted peptides also
Show ◉ uncorrected delta ○ delta corrected for 13C

| Query | Start - End | Observed | Mr(expt) | Mr(calc) | ppm | M | Score | Expect | Rank | U | Peptide |
|---|---|---|---|---|---|---|---|---|---|---|---|
| 409 | 25 - 43 | 731.3625 | 2191.0657 | 2191.0943 | -13.1 | 1 | 4 | 0.46 | 1 | U | K.QYMDRGDLVPDEVTIGIVR.E + Oxidation (M) |
| 410 | 25 - 43 | 731.3656 | 2191.0751 | 2191.0943 | -8.79 | 1 | 16 | 0.035 | 1 | U | K.QYMDRGDLVPDEVTIGIVR.E + Oxidation (M) |
| 412 | 25 - 43 | 731.3659 | 2191.0757 | 2191.0943 | -8.49 | 1 | 6 | 0.3 | 1 | U | K.QYMDRGDLVPDEVTIGIVR.E + Oxidation (M) |
| 413 | 25 - 43 | 731.3676 | 2191.0809 | 2191.0943 | -6.14 | 1 | 12 | 0.084 | 1 | U | K.QYMDRGDLVPDEVTIGIVR.E + Oxidation (M) |
| 414 | 25 - 43 | 731.3685 | 2191.0837 | 2191.0943 | -4.85 | 1 | 19 | 0.018 | 1 | U | K.QYMDRGDLVPDEVTIGIVR.E + Oxidation (M) |
| 415 | 25 - 43 | 731.3685 | 2191.0837 | 2191.0943 | -4.85 | 1 | 2 | 0.85 | 1 | U | K.QYMDRGDLVPDEVTIGIVR.E + Oxidation (M) |
| 242 | 30 - 43 | 741.9053 | 1481.7960 | 1481.8090 | -8.78 | 0 | 18 | 0.018 | 1 | U | R.GDLVPDEVTIGIVR.E |
| 243 | 30 - 43 | 741.9078 | 1481.8011 | 1481.8090 | -5.30 | 0 | 14 | 0.042 | 1 | U | R.GDLVPDEVTIGIVR.E |
| 244 | 30 - 43 | 741.9080 | 1481.8014 | 1481.8090 | -5.11 | 0 | 36 | 0.00026 | 1 | U | R.GDLVPDEVTIGIVR.E |
| 245 | 30 - 43 | 741.9086 | 1481.8026 | 1481.8090 | -4.31 | 0 | 14 | 0.038 | 1 | U | R.GDLVPDEVTIGIVR.E |
| 246 | 30 - 43 | 741.9103 | 1481.8061 | 1481.8090 | -1.92 | 0 | 9 | 0.12 | 1 | U | R.GDLVPDEVTIGIVR.E |
| 239 | 80 - 90 | 457.5929 | 1369.7570 | 1369.7718 | -10.8 | 1 | 16 | 0.024 | 1 | U | R.KLDYVIHIDVR.Q |
| 319 | 135 - 151 | 635.9602 | 1904.8587 | 1904.8898 | -16.3 | 1 | 0 | 0.98 | 1 | U | R.ADDNEATVANRLEVNMK.Q + Oxidation (M) |
| 320 | 135 - 151 | 635.9641 | 1904.8704 | 1904.8898 | -10.1 | 1 | 22 | 0.0062 | 1 | U | R.ADDNEATVANRLEVNMK.Q + Oxidation (M) |
| 321 | 135 - 151 | 635.9641 | 1904.8704 | 1904.8898 | -10.1 | 1 | 33 | 0.0005 | 1 | U | R.ADDNEATVANRLEVNMK.Q + Oxidation (M) |
| 322 | 135 - 151 | 635.9642 | 1904.8706 | 1904.8898 | -10.1 | 1 | 17 | 0.019 | 1 | U | R.ADDNEATVANRLEVNMK.Q + Oxidation (M) |
| 324 | 135 - 151 | 635.9647 | 1904.8723 | 1904.8898 | -9.17 | 1 | 2 | 0.67 | 1 | U | R.ADDNEATVANRLEVNMK.Q + Oxidation (M) |
| 325 | 135 - 151 | 635.9647 | 1904.8723 | 1904.8898 | -9.17 | 1 | 20 | 0.0097 | 1 | U | R.ADDNEATVANRLEVNMK.Q + Oxidation (M) |
| 327 | 135 - 151 | 635.9657 | 1904.8752 | 1904.8898 | -7.63 | 1 | 29 | 0.0012 | 1 | U | R.ADDNEATVANRLEVNMK.Q + Oxidation (M) |
| 328 | 135 - 151 | 635.9657 | 1904.8753 | 1904.8898 | -7.58 | 1 | 27 | 0.0022 | 1 | U | R.ADDNEATVANRLEVNMK.Q + Oxidation (M) |
| 329 | 135 - 151 | 635.9662 | 1904.8768 | 1904.8898 | -6.81 | 1 | 30 | 0.0011 | 1 | U | R.ADDNEATVANRLEVNMK.Q + Oxidation (M) |
| 330 | 135 - 151 | 635.9679 | 1904.8819 | 1904.8898 | -4.13 | 1 | 22 | 0.0074 | 1 | U | R.ADDNEATVANRLEVNMK.Q + Oxidation (M) |
| 331 | 135 - 151 | 635.9679 | 1904.8819 | 1904.8898 | -4.13 | 1 | 13 | 0.055 | 1 | U | R.ADDNEATVANRLEVNMK.Q + Oxidation (M) |
| 259 | 152 - 163 | 514.5874 | 1540.7403 | 1540.7596 | -12.5 | 1 | 1 | 1.5 | 1 | U | K.QMKPLVDFYEQK.G + Oxidation (M) |
| 262 | 192 - 207 | 773.4035 | 1544.7924 | 1544.8055 | -8.43 | 0 | 29 | 0.0021 | 1 | U | R.AAAMNLVLMGLPGAGK.G + 2 Oxidation (M) |
| 263 | 192 - 207 | 773.4035 | 1544.7924 | 1544.8055 | -8.43 | 0 | 24 | 0.006 | 1 | U | R.AAAMNLVLMGLPGAGK.G + 2 Oxidation (M) |
| 264 | 192 - 207 | 773.4040 | 1544.7934 | 1544.8055 | -7.81 | 0 | 19 | 0.021 | 1 | U | R.AAAMNLVLMGLPGAGK.G + 2 Oxidation (M) |

| Query | Start - End | Observed | Mr(expt) | Mr(calc) | ppm | M | Score | Expect | Rank | U | Peptide |
|---|---|---|---|---|---|---|---|---|---|---|---|
| 265 | 192 - 207 | 773.4047 | 1544.7948 | 1544.8055 | -6.90 | 0 | 6 | 0.4 | 1 | U | R.AAAMNLVLMGLPGAGK.G + 2 Oxidation (M) |
| 267 | 192 - 207 | 773.4052 | 1544.7959 | 1544.8055 | -6.19 | 0 | 17 | 0.031 | 1 | U | R.AAAMNLVLMGLPGAGK.G + 2 Oxidation (M) |
| 268 | 192 - 207 | 773.4056 | 1544.7966 | 1544.8055 | -5.72 | 0 | 26 | 0.0037 | 1 | U | R.AAAMNLVLMGLPGAGK.G + 2 Oxidation (M) |
| 269 | 192 - 207 | 773.4060 | 1544.7975 | 1544.8055 | -5.15 | 0 | 3 | 0.81 | 1 | U | R.AAAMNLVLMGLPGAGK.G + 2 Oxidation (M) |
| 270 | 192 - 207 | 773.4068 | 1544.7990 | 1544.8055 | -4.18 | 0 | 6 | 0.4 | 1 | U | R.AAAMNLVLMGLPGAGK.G + 2 Oxidation (M) |
| 271 | 192 - 207 | 773.4070 | 1544.7993 | 1544.8055 | -3.95 | 0 | 6 | 0.41 | 1 | U | R.AAAMNLVLMGLPGAGK.G + 2 Oxidation (M) |
| 272 | 192 - 207 | 773.4075 | 1544.8004 | 1544.8055 | -3.26 | 0 | 19 | 0.019 | 1 | U | R.AAAMNLVLMGLPGAGK.G + 2 Oxidation (M) |
| 273 | 192 - 207 | 773.4078 | 1544.8011 | 1544.8055 | -2.82 | 0 | 23 | 0.0073 | 1 | U | R.AAAMNLVLMGLPGAGK.G + 2 Oxidation (M) |
| 274 | 192 - 207 | 773.4084 | 1544.8023 | 1544.8055 | -2.03 | 0 | 22 | 0.011 | 1 | U | R.AAAMNLVLMGLPGAGK.G + 2 Oxidation (M) |
| 275 | 192 - 207 | 773.4084 | 1544.8023 | 1544.8055 | -2.03 | 0 | 20 | 0.015 | 1 | U | R.AAAMNLVLMGLPGAGK.G + 2 Oxidation (M) |
| 318 | 214 - 230 | 621.9836 | 1862.9289 | 1862.9349 | -3.23 | 0 | 2 | 0.66 | 1 | U | K.IVAAYGIPHISTGDMFR.A + Oxidation (M) |

Error: try setting browser cache to automatic.

---

```
LOCUS       QNL33370                  37 aa            linear   BCT 04-SEP-2020
DEFINITION  hypothetical protein (plasmid) [Escherichia coli].
ACCESSION   QNL33370
VERSION     QNL33370.1
DBSOURCE    accession MT180430.1
KEYWORDS    .
SOURCE      Escherichia coli
  ORGANISM  Escherichia coli
            Bacteria; Proteobacteria; Gammaproteobacteria; Enterobacterales;
            Enterobacteriaceae; Escherichia.
REFERENCE   1  (residues 1 to 37)
  AUTHORS   Tarabai,H., Wyrsch,E.R., Bitar,I., Djordjevic,S.P. and Dolejska,M.
  TITLE     Comparative analysis of multidrug resistant Escherichia coli ST216
            isolates from silver gulls in Australia
  JOURNAL   Unpublished
REFERENCE   2  (residues 1 to 37)
  AUTHORS   Tarabai,H., Wyrsch,E.R., Bitar,I., Djordjevic,S.P. and Dolejska,M.
  TITLE     Direct Submission
  JOURNAL   Submitted (09-MAR-2020) Department of Biology and Wildlife
            Diseases, University of Veterinary and Pharmaceutical Sciences
            Brno, Palackeho tr. 1946/1, Brno, Brno 61242, Czech Republic
COMMENT     ##Assembly-Data-START##
            Assembly Method       :: SMRT Link v. 8.0
            Assembly Name         :: pCE1681-A
            Coverage              :: 326X
            Sequencing Technology :: PacBio
            ##Assembly-Data-END##
FEATURES             Location/Qualifiers
     source          1..37
                     /organism="Escherichia coli"
                     /strain="CE1681"
                     /host="Chroicocephalus novaehollandiae"
                     /db_xref="taxon:562"
                     /plasmid="pCE1681-A"
                     /country="Australia"
                     /collection_date="2012"
                     /note="Closed IncHI2-ST3/IncN fusion plasmid;
                     type: ST216"
     Protein         1..37
                     /product="hypothetical protein"
     CDS             1..37
                     /coded_by="complement(MT180430.1:247749..247862)"
                     /note="hypothetical protein"
                     /transl_table=11
                     /db_xref="SEED:fig|6666666.506717.peg.327"
```

### Species 7

#### MATRIX SCIENCE MASCOT Search Results

##### Protein View: QNL33349.1

bsAK [Escherichia coli]

| | |
|---|---|
| Database: | NCBIcustTC |
| Score: | 2677 |
| Monoisotopic mass ($M_r$): | 24104 |
| Calculated pI: | 4.65 |

Sequencesimilarity is availableas an NCBI BLAST search of QNL33349.1 against nr.

##### Search parameters

| | |
|---|---|
| MS data file: | 09182020_TC-8_001.temp.mgf |
| Enzyme: | Trypsin/P: cuts C-term side of KR. |
| Variable modifications: | Oxidation (M), Propionamide (C), Propionamide (K), Propionamide (N-term), Carbamidomethyl (C) |

##### Protein sequence coverage: 71%

Matched peptides shown in **bold red**.

```
  1 MNLVLMGLPG AGKGTQGERI VEDYGIPHIS TGDMFRAAMK EETPLGLEAK
 51 SYIDKGELVP DEVTIGIVKE RLGKDDCERG FLLDGFPRTV AQAEALEEIL
101 EEYGKPIDYV INIEVDKDVL MERLTGRRIC SVCGTTYHLV FNPPKTPGIC
151 DKDGGELYQR ADDNEETVSK RLEVNMKQTQ PLLDFYSEKG YLANVNGQQD
201 IQDVYADVKD LLGGLKK
```

Unformatted sequence string: 217 residues (for pasting into other applications).

Sort by (●) residue number ( ) increasing mass ( ) decreasing mass
Show (●) matched peptides only ( ) predicted peptides also
Show (●) uncorrected delta ( ) delta corrected for 13C

| Query | Start - End | Observed | Mr(expt) | Mr(calc) | ppm | M | Score | Expect | Rank | U | Peptide |
|---|---|---|---|---|---|---|---|---|---|---|---|
| 525 | 20 - 36 | 650.6460 | 1948.9162 | 1948.9353 | -9.79 | 0 | 3 | 0.81 | 1 | U | R.IVEDYGIPHISTGDMFR.A |
| 526 | 20 - 36 | 650.6461 | 1948.9164 | 1948.9353 | -9.71 | 0 | 19 | 0.024 | 1 | U | R.IVEDYGIPHISTGDMFR.A |
| 527 | 20 - 36 | 650.6464 | 1948.9173 | 1948.9353 | -9.26 | 0 | 4 | 0.7 | 1 | U | R.IVEDYGIPHISTGDMFR.A |
| 528 | 20 - 36 | 650.6464 | 1948.9174 | 1948.9353 | -9.17 | 0 | 39 | 0.00022 | 1 | U | R.IVEDYGIPHISTGDMFR.A |
| 529 | 20 - 36 | 650.6467 | 1948.9183 | 1948.9353 | -8.71 | 0 | 39 | 0.00022 | 1 | U | R.IVEDYGIPHISTGDMFR.A |
| 530 | 20 - 36 | 650.6468 | 1948.9186 | 1948.9353 | -8.55 | 0 | 8 | 0.31 | 1 | U | R.IVEDYGIPHISTGDMFR.A |
| 531 | 20 - 36 | 650.6471 | 1948.9196 | 1948.9353 | -8.06 | 0 | 21 | 0.018 | 1 | U | R.IVEDYGIPHISTGDMFR.A |
| 532 | 20 - 36 | 650.6473 | 1948.9200 | 1948.9353 | -7.88 | 0 | 30 | 0.0019 | 1 | U | R.IVEDYGIPHISTGDMFR.A |
| 533 | 20 - 36 | 650.6474 | 1948.9204 | 1948.9353 | -7.63 | 0 | 53 | 9.4e-006 | 1 | U | R.IVEDYGIPHISTGDMFR.A |
| 534 | 20 - 36 | 650.6475 | 1948.9207 | 1948.9353 | -7.49 | 0 | 22 | 0.012 | 1 | U | R.IVEDYGIPHISTGDMFR.A |
| 535 | 20 - 36 | 650.6478 | 1948.9215 | 1948.9353 | -7.09 | 0 | 22 | 0.013 | 1 | U | R.IVEDYGIPHISTGDMFR.A |
| 536 | 20 - 36 | 975.4686 | 1948.9227 | 1948.9353 | -6.45 | 0 | 13 | 0.1 | 1 | U | R.IVEDYGIPHISTGDMFR.A |
| 537 | 20 - 36 | 650.6482 | 1948.9228 | 1948.9353 | -6.43 | 0 | 28 | 0.0032 | 1 | U | R.IVEDYGIPHISTGDMFR.A |
| 538 | 20 - 36 | 650.6484 | 1948.9233 | 1948.9353 | -6.18 | 0 | 26 | 0.006 | 1 | U | R.IVEDYGIPHISTGDMFR.A |
| 539 | 20 - 36 | 975.4691 | 1948.9236 | 1948.9353 | -5.99 | 0 | 21 | 0.017 | 1 | U | R.IVEDYGIPHISTGDMFR.A |
| 541 | 20 - 36 | 975.4720 | 1948.9294 | 1948.9353 | -3.00 | 0 | 15 | 0.069 | 1 | U | R.IVEDYGIPHISTGDMFR.A |
| 542 | 20 - 36 | 975.4739 | 1948.9333 | 1948.9353 | -1.01 | 0 | 18 | 0.034 | 1 | U | R.IVEDYGIPHISTGDMFR.A |
| 543 | 20 - 36 | 983.4596 | 1964.9047 | 1964.9302 | -13.0 | 0 | 22 | 0.0065 | 1 | U | R.IVEDYGIPHISTGDMFR.A + Oxidation (M) |
| 544 | 20 - 36 | 983.4602 | 1964.9058 | 1964.9302 | -12.4 | 0 | 18 | 0.017 | 1 | U | R.IVEDYGIPHISTGDMFR.A + Oxidation (M) |
| 545 | 20 - 36 | 983.4624 | 1964.9103 | 1964.9302 | -10.1 | 0 | 6 | 0.27 | 1 | U | R.IVEDYGIPHISTGDMFR.A + Oxidation (M) |
| 546 | 20 - 36 | 655.9778 | 1964.9117 | 1964.9302 | -9.43 | 0 | 7 | 0.22 | 1 | U | R.IVEDYGIPHISTGDMFR.A + Oxidation (M) |
| 547 | 20 - 36 | 655.9779 | 1964.9120 | 1964.9302 | -9.26 | 0 | 53 | 5.1e-006 | 1 | U | R.IVEDYGIPHISTGDMFR.A + Oxidation (M) |
| 548 | 20 - 36 | 983.4633 | 1964.9121 | 1964.9302 | -9.23 | 0 | 9 | 0.13 | 1 | U | R.IVEDYGIPHISTGDMFR.A + Oxidation (M) |
| 549 | 20 - 36 | 655.9782 | 1964.9127 | 1964.9302 | -8.94 | 0 | 55 | 2.9e-006 | 1 | U | R.IVEDYGIPHISTGDMFR.A + Oxidation (M) |
| 550 | 20 - 36 | 655.9782 | 1964.9128 | 1964.9302 | -8.85 | 0 | 46 | 2.7e-005 | 1 | U | R.IVEDYGIPHISTGDMFR.A + Oxidation (M) |
| 551 | 20 - 36 | 983.4638 | 1964.9130 | 1964.9302 | -8.78 | 0 | 18 | 0.016 | 1 | U | R.IVEDYGIPHISTGDMFR.A + Oxidation (M) |
| 552 | 20 - 36 | 655.9783 | 1964.9132 | 1964.9302 | -8.65 | 0 | 41 | 8.4e-005 | 1 | U | R.IVEDYGIPHISTGDMFR.A + Oxidation (M) |
| 553 | 20 - 36 | 655.9786 | 1964.9139 | 1964.9302 | -8.33 | 0 | 42 | 6.2e-005 | 1 | U | R.IVEDYGIPHISTGDMFR.A + Oxidation (M) |
| 554 | 20 - 36 | 983.4643 | 1964.9141 | 1964.9302 | -8.18 | 0 | 21 | 0.0086 | 1 | U | R.IVEDYGIPHISTGDMFR.A + Oxidation (M) |
| 555 | 20 - 36 | 655.9787 | 1964.9144 | 1964.9302 | -8.07 | 0 | 14 | 0.038 | 1 | U | R.IVEDYGIPHISTGDMFR.A + Oxidation (M) |

| Query | Start - End | Observed | Mr(expt) | Mr(calc) | ppm | M | Score | Expect | Rank | U | Peptide |
|---|---|---|---|---|---|---|---|---|---|---|---|
| 556 | 20 - 36 | 655.9787 | 1964.9144 | 1964.9302 | -8.04 | 0 | 9 | 0.12 | 1 | U | R.IVEDYGIPHISTGDMFR.A + Oxidation (M) |
| 557 | 20 - 36 | 655.9787 | 1964.9144 | 1964.9302 | -8.04 | 0 | 26 | 0.0026 | 1 | U | R.IVEDYGIPHISTGDMFR.A + Oxidation (M) |
| 558 | 20 - 36 | 655.9789 | 1964.9148 | 1964.9302 | -7.86 | 0 | 67 | 2.1e-007 | 1 | U | R.IVEDYGIPHISTGDMFR.A + Oxidation (M) |
| 559 | 20 - 36 | 655.9790 | 1964.9150 | 1964.9302 | -7.74 | 0 | 10 | 0.11 | 1 | U | R.IVEDYGIPHISTGDMFR.A + Oxidation (M) |
| 560 | 20 - 36 | 655.9790 | 1964.9152 | 1964.9302 | -7.64 | 0 | 46 | 2.6e-005 | 1 | U | R.IVEDYGIPHISTGDMFR.A + Oxidation (M) |
| 561 | 20 - 36 | 655.9792 | 1964.9157 | 1964.9302 | -7.37 | 0 | 22 | 0.0067 | 1 | U | R.IVEDYGIPHISTGDMFR.A + Oxidation (M) |
| 562 | 20 - 36 | 655.9792 | 1964.9157 | 1964.9302 | -7.37 | 0 | 32 | 0.00062 | 1 | U | R.IVEDYGIPHISTGDMFR.A + Oxidation (M) |
| 563 | 20 - 36 | 655.9793 | 1964.9160 | 1964.9302 | -7.22 | 0 | 35 | 0.00029 | 1 | U | R.IVEDYGIPHISTGDMFR.A + Oxidation (M) |
| 564 | 20 - 36 | 655.9793 | 1964.9160 | 1964.9302 | -7.22 | 0 | 40 | 9.2e-005 | 1 | U | R.IVEDYGIPHISTGDMFR.A + Oxidation (M) |
| 565 | 20 - 36 | 655.9794 | 1964.9163 | 1964.9302 | -7.09 | 0 | 47 | 1.9e-005 | 1 | U | R.IVEDYGIPHISTGDMFR.A + Oxidation (M) |
| 566 | 20 - 36 | 983.4654 | 1964.9163 | 1964.9302 | -7.09 | 0 | 26 | 0.0026 | 1 | U | R.IVEDYGIPHISTGDMFR.A + Oxidation (M) |
| 567 | 20 - 36 | 655.9795 | 1964.9166 | 1964.9302 | -6.93 | 0 | 9 | 0.12 | 1 | U | R.IVEDYGIPHISTGDMFR.A + Oxidation (M) |
| 568 | 20 - 36 | 655.9795 | 1964.9167 | 1964.9302 | -6.88 | 0 | 53 | 4.9e-006 | 1 | U | R.IVEDYGIPHISTGDMFR.A + Oxidation (M) |
| 569 | 20 - 36 | 655.9795 | 1964.9167 | 1964.9302 | -6.87 | 0 | 18 | 0.016 | 1 | U | R.IVEDYGIPHISTGDMFR.A + Oxidation (M) |
| 571 | 20 - 36 | 655.9800 | 1964.9180 | 1964.9302 | -6.21 | 0 | 41 | 7.7e-005 | 1 | U | R.IVEDYGIPHISTGDMFR.A + Oxidation (M) |
| 572 | 20 - 36 | 983.4668 | 1964.9190 | 1964.9302 | -5.72 | 0 | 3 | 0.52 | 1 | U | R.IVEDYGIPHISTGDMFR.A + Oxidation (M) |
| 573 | 20 - 36 | 655.9803 | 1964.9191 | 1964.9302 | -5.67 | 0 | 52 | 6.2e-006 | 1 | U | R.IVEDYGIPHISTGDMFR.A + Oxidation (M) |
| 574 | 20 - 36 | 655.9803 | 1964.9192 | 1964.9302 | -5.63 | 0 | 34 | 0.00043 | 1 | U | R.IVEDYGIPHISTGDMFR.A + Oxidation (M) |
| 575 | 20 - 36 | 983.4672 | 1964.9198 | 1964.9302 | -5.29 | 0 | 32 | 0.00059 | 1 | U | R.IVEDYGIPHISTGDMFR.A + Oxidation (M) |
| 576 | 20 - 36 | 983.4677 | 1964.9207 | 1964.9302 | -4.82 | 0 | 5 | 0.31 | 1 | U | R.IVEDYGIPHISTGDMFR.A + Oxidation (M) |
| 577 | 20 - 36 | 983.4679 | 1964.9212 | 1964.9302 | -4.57 | 0 | 5 | 0.33 | 1 | U | R.IVEDYGIPHISTGDMFR.A + Oxidation (M) |
| 578 | 20 - 36 | 655.9812 | 1964.9217 | 1964.9302 | -4.32 | 0 | 17 | 0.02 | 1 | U | R.IVEDYGIPHISTGDMFR.A + Oxidation (M) |
| 580 | 20 - 36 | 983.4689 | 1964.9233 | 1964.9302 | -3.54 | 0 | 20 | 0.0098 | 1 | U | R.IVEDYGIPHISTGDMFR.A + Oxidation (M) |
| 581 | 20 - 36 | 983.4692 | 1964.9238 | 1964.9302 | -3.28 | 0 | 17 | 0.018 | 1 | U | R.IVEDYGIPHISTGDMFR.A + Oxidation (M) |
| 582 | 20 - 36 | 983.4693 | 1964.9241 | 1964.9302 | -3.12 | 0 | 18 | 0.016 | 1 | U | R.IVEDYGIPHISTGDMFR.A + Oxidation (M) |
| 583 | 20 - 36 | 983.4698 | 1964.9251 | 1964.9302 | -2.62 | 0 | 8 | 0.17 | 1 | U | R.IVEDYGIPHISTGDMFR.A + Oxidation (M) |
| 584 | 20 - 36 | 983.4709 | 1964.9272 | 1964.9302 | -1.52 | 0 | 23 | 0.0049 | 1 | U | R.IVEDYGIPHISTGDMFR.A + Oxidation (M) |
| 585 | 20 - 36 | 983.4711 | 1964.9276 | 1964.9302 | -1.31 | 0 | 30 | 0.00095 | 1 | U | R.IVEDYGIPHISTGDMFR.A + Oxidation (M) |
| 586 | 20 - 36 | 983.4728 | 1964.9311 | 1964.9302 | 0.46 | 0 | 19 | 0.014 | 1 | U | R.IVEDYGIPHISTGDMFR.A + Oxidation (M) |
| 399 | 37 - 50 | 744.3852 | 1486.7559 | 1486.7701 | -9.53 | 1 | 0 | 0.94 | 1 | U | R.AAMKEETPLGLEAK.S |
| 400 | 37 - 50 | 744.3861 | 1486.7577 | 1486.7701 | -8.34 | 1 | 4 | 0.4 | 1 | U | R.AAMKEETPLGLEAK.S |
| 401 | 37 - 50 | 744.3865 | 1486.7584 | 1486.7701 | -7.88 | 1 | 21 | 0.0087 | 1 | U | R.AAMKEETPLGLEAK.S |
| 402 | 37 - 50 | 744.3866 | 1486.7585 | 1486.7701 | -7.76 | 1 | 1 | 0.79 | 1 | U | R.AAMKEETPLGLEAK.S |
| 403 | 37 - 50 | 744.3871 | 1486.7596 | 1486.7701 | -7.03 | 1 | 3 | 0.51 | 1 | U | R.AAMKEETPLGLEAK.S |
| 407 | 37 - 50 | 501.9223 | 1502.7452 | 1502.7650 | -13.2 | 1 | 5 | 0.29 | 1 | U | R.AAMKEETPLGLEAK.S + Oxidation (M) |
| 408 | 37 - 50 | 501.9237 | 1502.7493 | 1502.7650 | -10.4 | 1 | 6 | 0.24 | 1 | U | R.AAMKEETPLGLEAK.S + Oxidation (M) |
| 409 | 37 - 50 | 752.3820 | 1502.7495 | 1502.7650 | -10.3 | 1 | 10 | 0.091 | 1 | U | R.AAMKEETPLGLEAK.S + Oxidation (M) |
| 410 | 37 - 50 | 501.9238 | 1502.7495 | 1502.7650 | -10.3 | 1 | 28 | 0.0015 | 1 | U | R.AAMKEETPLGLEAK.S + Oxidation (M) |
| 411 | 37 - 50 | 501.9242 | 1502.7508 | 1502.7650 | -9.43 | 1 | 9 | 0.13 | 1 | U | R.AAMKEETPLGLEAK.S + Oxidation (M) |
| 414 | 37 - 50 | 752.3830 | 1502.7514 | 1502.7650 | -9.07 | 1 | 6 | 0.24 | 1 | U | R.AAMKEETPLGLEAK.S + Oxidation (M) |
| 415 | 37 - 50 | 752.3831 | 1502.7517 | 1502.7650 | -8.87 | 1 | 22 | 0.0065 | 1 | U | R.AAMKEETPLGLEAK.S + Oxidation (M) |
| 416 | 37 - 50 | 501.9248 | 1502.7526 | 1502.7650 | -8.25 | 1 | 16 | 0.024 | 1 | U | R.AAMKEETPLGLEAK.S + Oxidation (M) |
| 417 | 37 - 50 | 501.9253 | 1502.7541 | 1502.7650 | -7.25 | 1 | 5 | 0.34 | 1 | U | R.AAMKEETPLGLEAK.S + Oxidation (M) |
| 418 | 37 - 50 | 752.3844 | 1502.7541 | 1502.7650 | -7.22 | 1 | 24 | 0.0038 | 1 | U | R.AAMKEETPLGLEAK.S + Oxidation (M) |
| 419 | 37 - 50 | 752.3844 | 1502.7542 | 1502.7650 | -7.21 | 1 | 46 | 2.5e-005 | 1 | U | R.AAMKEETPLGLEAK.S + Oxidation (M) |
| 420 | 37 - 50 | 752.3846 | 1502.7546 | 1502.7650 | -6.94 | 1 | 24 | 0.0044 | 1 | U | R.AAMKEETPLGLEAK.S + Oxidation (M) |
| 421 | 37 - 50 | 752.3850 | 1502.7553 | 1502.7650 | -6.42 | 1 | 24 | 0.0041 | 1 | U | R.AAMKEETPLGLEAK.S + Oxidation (M) |
| 422 | 37 - 50 | 752.3856 | 1502.7566 | 1502.7650 | -5.57 | 1 | 5 | 0.31 | 1 | U | R.AAMKEETPLGLEAK.S + Oxidation (M) |
| 423 | 37 - 50 | 752.3858 | 1502.7570 | 1502.7650 | -5.34 | 1 | 1 | 0.78 | 1 | U | R.AAMKEETPLGLEAK.S + Oxidation (M) |
| 424 | 37 - 50 | 752.3858 | 1502.7570 | 1502.7650 | -5.29 | 1 | 16 | 0.024 | 1 | U | R.AAMKEETPLGLEAK.S + Oxidation (M) |
| 425 | 37 - 50 | 752.3858 | 1502.7571 | 1502.7650 | -5.25 | 1 | 25 | 0.0034 | 1 | U | R.AAMKEETPLGLEAK.S + Oxidation (M) |
| 426 | 37 - 50 | 752.3858 | 1502.7571 | 1502.7650 | -5.24 | 1 | 1 | 0.84 | 1 | U | R.AAMKEETPLGLEAK.S + Oxidation (M) |
| 427 | 37 - 50 | 752.3859 | 1502.7573 | 1502.7650 | -5.14 | 1 | 10 | 0.11 | 1 | U | R.AAMKEETPLGLEAK.S + Oxidation (M) |
| 428 | 37 - 50 | 752.3860 | 1502.7574 | 1502.7650 | -5.02 | 1 | 34 | 0.00036 | 1 | U | R.AAMKEETPLGLEAK.S + Oxidation (M) |
| 429 | 37 - 50 | 752.3861 | 1502.7576 | 1502.7650 | -4.90 | 1 | 16 | 0.025 | 1 | U | R.AAMKEETPLGLEAK.S + Oxidation (M) |
| 430 | 37 - 50 | 752.3861 | 1502.7576 | 1502.7650 | -4.90 | 1 | 25 | 0.003 | 1 | U | R.AAMKEETPLGLEAK.S + Oxidation (M) |
| 431 | 37 - 50 | 752.3865 | 1502.7584 | 1502.7650 | -4.41 | 1 | 18 | 0.015 | 1 | U | R.AAMKEETPLGLEAK.S + Oxidation (M) |
| 432 | 37 - 50 | 752.3865 | 1502.7584 | 1502.7650 | -4.39 | 1 | 11 | 0.082 | 1 | U | R.AAMKEETPLGLEAK.S + Oxidation (M) |
| 433 | 37 - 50 | 752.3873 | 1502.7601 | 1502.7650 | -3.25 | 1 | 33 | 0.00053 | 1 | U | R.AAMKEETPLGLEAK.S + Oxidation (M) |
| 434 | 37 - 50 | 752.3877 | 1502.7607 | 1502.7650 | -2.83 | 1 | 27 | 0.002 | 1 | U | R.AAMKEETPLGLEAK.S + Oxidation (M) |
| 435 | 37 - 50 | 752.3879 | 1502.7612 | 1502.7650 | -2.50 | 1 | 1 | 0.8 | 1 | U | R.AAMKEETPLGLEAK.S + Oxidation (M) |
| 436 | 37 - 50 | 752.3892 | 1502.7639 | 1502.7650 | -0.75 | 1 | 2 | 0.62 | 1 | U | R.AAMKEETPLGLEAK.S + Oxidation (M) |
| 612 | 51 - 69 | 692.3722 | 2074.0947 | 2074.1198 | -12.1 | 1 | 7 | 0.39 | 1 | U | K.SYIDKGELVPDEVTIGIVK.E |
| 613 | 51 - 69 | 692.3725 | 2074.0958 | 2074.1198 | -11.6 | 1 | 21 | 0.016 | 1 | U | K.SYIDKGELVPDEVTIGIVK.E |
| 614 | 51 - 69 | 692.3727 | 2074.0963 | 2074.1198 | -11.3 | 1 | 12 | 0.1 | 1 | U | K.SYIDKGELVPDEVTIGIVK.E |
| 615 | 51 - 69 | 1038.0575 | 2074.1004 | 2074.1198 | -9.32 | 1 | 48 | 2.8e-005 | 1 | U | K.SYIDKGELVPDEVTIGIVK.E |
| 616 | 51 - 69 | 692.3742 | 2074.1008 | 2074.1198 | -9.16 | 1 | 38 | 0.00027 | 1 | U | K.SYIDKGELVPDEVTIGIVK.E |
| 617 | 51 - 69 | 692.3744 | 2074.1013 | 2074.1198 | -8.93 | 1 | 49 | 2.1e-005 | 1 | U | K.SYIDKGELVPDEVTIGIVK.E |

| Query | Start – End | Observed | Mr(expt) | Mr(calc) | ppm | M | Score | Expect | Rank | U | Peptide |
|---|---|---|---|---|---|---|---|---|---|---|---|
| 618 | 51 – 69 | 692.3744 | 2074.1014 | 2074.1198 | -8.85 | 1 | 26 | 0.0039 | 1 | U | K.SYIDKGELVPDEVTIGIVK.E |
| 619 | 51 – 69 | 692.3745 | 2074.1016 | 2074.1198 | -8.77 | 1 | 57 | 3.2e-006 | 1 | U | K.SYIDKGELVPDEVTIGIVK.E |
| 620 | 51 – 69 | 692.3750 | 2074.1031 | 2074.1198 | -8.04 | 1 | 14 | 0.062 | 1 | U | K.SYIDKGELVPDEVTIGIVK.E |
| 621 | 51 – 69 | 692.3752 | 2074.1038 | 2074.1198 | -7.69 | 1 | 44 | 6.1e-005 | 1 | U | K.SYIDKGELVPDEVTIGIVK.E |
| 622 | 51 – 69 | 692.3756 | 2074.1051 | 2074.1198 | -7.07 | 1 | 49 | 1.8e-005 | 1 | U | K.SYIDKGELVPDEVTIGIVK.E |
| 623 | 51 – 69 | 692.3757 | 2074.1052 | 2074.1198 | -7.02 | 1 | 49 | 2e-005 | 1 | U | K.SYIDKGELVPDEVTIGIVK.E |
| 624 | 51 – 69 | 692.3758 | 2074.1055 | 2074.1198 | -6.89 | 1 | 48 | 2.2e-005 | 1 | U | K.SYIDKGELVPDEVTIGIVK.E |
| 625 | 51 – 69 | 692.3762 | 2074.1066 | 2074.1198 | -6.34 | 1 | 61 | 1.1e-006 | 1 | U | K.SYIDKGELVPDEVTIGIVK.E |
| 626 | 51 – 69 | 1038.0611 | 2074.1076 | 2074.1198 | -5.85 | 1 | 73 | 8.5e-008 | 1 | U | K.SYIDKGELVPDEVTIGIVK.E |
| 627 | 51 – 69 | 692.3766 | 2074.1080 | 2074.1198 | -5.68 | 1 | 53 | 7.4e-006 | 1 | U | K.SYIDKGELVPDEVTIGIVK.E |
| 628 | 51 – 69 | 1038.0620 | 2074.1094 | 2074.1198 | -4.98 | 1 | 29 | 0.0019 | 1 | U | K.SYIDKGELVPDEVTIGIVK.E |
| 629 | 51 – 69 | 1038.0621 | 2074.1096 | 2074.1198 | -4.89 | 1 | 45 | 4.2e-005 | 1 | U | K.SYIDKGELVPDEVTIGIVK.E |
| 630 | 51 – 69 | 1038.0623 | 2074.1100 | 2074.1198 | -4.69 | 1 | 51 | 1e-005 | 1 | U | K.SYIDKGELVPDEVTIGIVK.E |
| 632 | 51 – 69 | 1038.0631 | 2074.1116 | 2074.1198 | -3.92 | 1 | 43 | 6.2e-005 | 1 | U | K.SYIDKGELVPDEVTIGIVK.E |
| 633 | 51 – 69 | 1038.0652 | 2074.1158 | 2074.1198 | -1.90 | 1 | 48 | 1.6e-005 | 1 | U | K.SYIDKGELVPDEVTIGIVK.E |
| 277 | 80 – 88 | 511.2694 | 1020.5242 | 1020.5393 | -14.8 | 0 | 37 | 0.00018 | 1 | U | R.GFLLDGFPR.T |
| 278 | 80 – 88 | 511.2700 | 1020.5254 | 1020.5393 | -13.6 | 0 | 24 | 0.0037 | 1 | U | R.GFLLDGFPR.T |
| 279 | 80 – 88 | 511.2704 | 1020.5263 | 1020.5393 | -12.7 | 0 | 46 | 2.6e-005 | 1 | U | R.GFLLDGFPR.T |
| 280 | 80 – 88 | 511.2710 | 1020.5274 | 1020.5393 | -11.6 | 0 | 31 | 0.00084 | 1 | U | R.GFLLDGFPR.T |
| 281 | 80 – 88 | 511.2713 | 1020.5281 | 1020.5393 | -11.0 | 0 | 24 | 0.0042 | 1 | U | R.GFLLDGFPR.T |
| 282 | 80 – 88 | 511.2718 | 1020.5290 | 1020.5393 | -10.0 | 0 | 47 | 2.1e-005 | 1 | U | R.GFLLDGFPR.T |
| 283 | 80 – 88 | 511.2720 | 1020.5295 | 1020.5393 | -9.53 | 0 | 52 | 6.2e-006 | 1 | U | R.GFLLDGFPR.T |
| 284 | 80 – 88 | 511.2726 | 1020.5307 | 1020.5393 | -8.42 | 0 | 31 | 0.0008 | 1 | U | R.GFLLDGFPR.T |
| 285 | 80 – 88 | 511.2727 | 1020.5309 | 1020.5393 | -8.22 | 0 | 36 | 0.00022 | 1 | U | R.GFLLDGFPR.T |
| 286 | 80 – 88 | 511.2727 | 1020.5309 | 1020.5393 | -8.18 | 0 | 37 | 0.0002 | 1 | U | R.GFLLDGFPR.T |
| 287 | 80 – 88 | 511.2728 | 1020.5311 | 1020.5393 | -8.00 | 0 | 32 | 0.00063 | 1 | U | R.GFLLDGFPR.T |
| 288 | 80 – 88 | 511.2733 | 1020.5320 | 1020.5393 | -7.14 | 0 | 51 | 7.7e-006 | 1 | U | R.GFLLDGFPR.T |
| 289 | 80 – 88 | 511.2737 | 1020.5329 | 1020.5393 | -6.22 | 0 | 24 | 0.0043 | 1 | U | R.GFLLDGFPR.T |
| 290 | 80 – 88 | 511.2745 | 1020.5345 | 1020.5393 | -4.67 | 0 | 26 | 0.0026 | 1 | U | R.GFLLDGFPR.T |
| 291 | 80 – 88 | 511.2745 | 1020.5345 | 1020.5393 | -4.63 | 0 | 35 | 0.00035 | 1 | U | R.GFLLDGFPR.T |
| 292 | 80 – 88 | 511.2750 | 1020.5354 | 1020.5393 | -3.83 | 0 | 35 | 0.00032 | 1 | U | R.GFLLDGFPR.T |
| 293 | 80 – 88 | 511.2753 | 1020.5361 | 1020.5393 | -3.14 | 0 | 42 | 6.3e-005 | 1 | U | R.GFLLDGFPR.T |
| 294 | 80 – 88 | 511.2756 | 1020.5367 | 1020.5393 | -2.56 | 0 | 39 | 0.00013 | 1 | U | R.GFLLDGFPR.T |
| 295 | 80 – 88 | 511.2761 | 1020.5377 | 1020.5393 | -1.54 | 0 | 35 | 0.0003 | 1 | U | R.GFLLDGFPR.T |
| 951 | 89 – 123 | 1345.6783 | 4034.0131 | 4034.0445 | -7.79 | 2 | 3 | 0.66 | 1 | U | R.TVAQAEALEEILEEYGKPIDYVINIEVDKDVLMER.L |
| 952 | 89 – 123 | 1346.0164 | 4035.0274 | 4034.0445 | 244 | 2 | 12 | 0.1 | 1 | U | R.TVAQAEALEEILEEYGKPIDYVINIEVDKDVLMER.L |
| 954 | 89 – 123 | 1351.0029 | 4049.9869 | 4050.0394 | -13.0 | 2 | 10 | 0.14 | 1 | U | R.TVAQAEALEEILEEYGKPIDYVINIEVDKDVLMER.L + Oxidation (M) |
| 955 | 89 – 123 | 1351.0029 | 4049.9869 | 4050.0394 | -13.0 | 2 | 10 | 0.12 | 1 | U | R.TVAQAEALEEILEEYGKPIDYVINIEVDKDVLMER.L + Oxidation (M) |
| 956 | 89 – 123 | 1351.0047 | 4049.9923 | 4050.0394 | -11.6 | 2 | 13 | 0.065 | 1 | U | R.TVAQAEALEEILEEYGKPIDYVINIEVDKDVLMER.L + Oxidation (M) |
| 957 | 89 – 123 | 1351.0047 | 4049.9923 | 4050.0394 | -11.6 | 2 | 23 | 0.0069 | 1 | U | R.TVAQAEALEEILEEYGKPIDYVINIEVDKDVLMER.L + Oxidation (M) |
| 958 | 89 – 123 | 1351.0070 | 4049.9992 | 4050.0394 | -9.94 | 2 | 12 | 0.078 | 1 | U | R.TVAQAEALEEILEEYGKPIDYVINIEVDKDVLMER.L + Oxidation (M) |
| 959 | 89 – 123 | 1351.0086 | 4050.0040 | 4050.0394 | -8.75 | 2 | 19 | 0.015 | 1 | U | R.TVAQAEALEEILEEYGKPIDYVINIEVDKDVLMER.L + Oxidation (M) |
| 960 | 89 – 123 | 1351.0086 | 4050.0040 | 4050.0394 | -8.75 | 2 | 12 | 0.074 | 1 | U | R.TVAQAEALEEILEEYGKPIDYVINIEVDKDVLMER.L + Oxidation (M) |
| 961 | 89 – 123 | 1351.0091 | 4050.0055 | 4050.0394 | -8.38 | 2 | 3 | 0.61 | 1 | U | R.TVAQAEALEEILEEYGKPIDYVINIEVDKDVLMER.L + Oxidation (M) |
| 962 | 89 – 123 | 1351.0092 | 4050.0058 | 4050.0394 | -8.31 | 2 | 20 | 0.014 | 1 | U | R.TVAQAEALEEILEEYGKPIDYVINIEVDKDVLMER.L + Oxidation (M) |
| 963 | 89 – 123 | 1351.0092 | 4050.0058 | 4050.0394 | -8.31 | 2 | 24 | 0.0048 | 1 | U | R.TVAQAEALEEILEEYGKPIDYVINIEVDKDVLMER.L + Oxidation (M) |
| 964 | 89 – 123 | 1351.0127 | 4050.0163 | 4050.0394 | -5.72 | 2 | 5 | 0.45 | 1 | U | R.TVAQAEALEEILEEYGKPIDYVINIEVDKDVLMER.L + Oxidation (M) |
| 965 | 89 – 123 | 1351.0134 | 4050.0184 | 4050.0394 | -5.20 | 2 | 10 | 0.12 | 1 | U | R.TVAQAEALEEILEEYGKPIDYVINIEVDKDVLMER.L + Oxidation (M) |
| 966 | 89 – 123 | 1351.0134 | 4050.0184 | 4050.0394 | -5.20 | 2 | 25 | 0.0047 | 1 | U | R.TVAQAEALEEILEEYGKPIDYVINIEVDKDVLMER.L + Oxidation (M) |
| 967 | 89 – 123 | 1351.0142 | 4050.0208 | 4050.0394 | -4.61 | 2 | 2 | 0.64 | 1 | U | R.TVAQAEALEEILEEYGKPIDYVINIEVDKDVLMER.L + Oxidation (M) |
| 968 | 89 – 123 | 1351.0249 | 4050.0529 | 4050.0394 | 3.32 | 2 | 11 | 0.078 | 1 | U | R.TVAQAEALEEILEEYGKPIDYVINIEVDKDVLMER.L + Oxidation (M) |
| 970 | 89 – 123 | 1351.3424 | 4051.0054 | 4050.0394 | 239 | 2 | 11 | 0.1 | 1 | U | R.TVAQAEALEEILEEYGKPIDYVINIEVDKDVLMER.L + Oxidation (M) |
| 971 | 89 – 123 | 1351.3489 | 4051.0249 | 4050.0394 | 243 | 2 | 16 | 0.029 | 1 | U | R.TVAQAEALEEILEEYGKPIDYVINIEVDKDVLMER.L + Oxidation (M) |
| 972 | 89 – 123 | 1351.3489 | 4051.0249 | 4050.0394 | 243 | 2 | 26 | 0.0025 | 1 | U | R.TVAQAEALEEILEEYGKPIDYVINIEVDKDVLMER.L + Oxidation (M) |
| 973 | 89 – 123 | 1351.3637 | 4051.0693 | 4050.0394 | 254 | 2 | 16 | 0.023 | 1 | U | R.TVAQAEALEEILEEYGKPIDYVINIEVDKDVLMER.L + Oxidation (M) |
| 974 | 89 – 123 | 1351.3637 | 4051.0693 | 4050.0394 | 254 | 2 | 12 | 0.06 | 1 | U | R.TVAQAEALEEILEEYGKPIDYVINIEVDKDVLMER.L + Oxidation (M) |
| 486 | 146 – 160 | 574.9312 | 1721.7717 | 1721.8043 | -18.9 | 1 | 14 | 0.051 | 1 | U | K.TPGICDKDGGELYQR.A + Propionamide (C) |
| 487 | 146 – 160 | 574.9325 | 1721.7758 | 1721.8043 | -16.6 | 1 | 4 | 0.49 | 1 | U | K.TPGICDKDGGELYQR.A + Propionamide (K) |
| 488 | 146 – 160 | 574.9353 | 1721.7840 | 1721.8043 | -11.8 | 1 | 0 | 1.4 | 1 | U | K.TPGICDKDGGELYQR.A + Propionamide (K) |
| 493 | 146 – 160 | 574.9397 | 1721.7973 | 1721.8043 | -4.05 | 1 | 14 | 0.051 | 1 | U | K.TPGICDKDGGELYQR.A + Propionamide (C) |
| 161 | 171 – 177 | 445.2463 | 888.4781 | 888.4851 | -7.86 | 1 | 17 | 0.02 | 1 | U | K.RLEVNMK.Q |
| 162 | 171 – 177 | 445.2468 | 888.4790 | 888.4851 | -6.83 | 1 | 11 | 0.075 | 1 | U | K.RLEVNMK.Q |
| 163 | 171 – 177 | 445.2469 | 888.4792 | 888.4851 | -6.62 | 1 | 3 | 0.54 | 1 | U | K.RLEVNMK.Q |
| 164 | 171 – 177 | 445.2471 | 888.4797 | 888.4851 | -6.06 | 1 | 22 | 0.0062 | 1 | U | K.RLEVNMK.Q |
| 165 | 171 – 177 | 445.2472 | 888.4798 | 888.4851 | -5.90 | 1 | 2 | 0.57 | 1 | U | K.RLEVNMK.Q |
| 166 | 171 – 177 | 445.2476 | 888.4806 | 888.4851 | -5.07 | 1 | 20 | 0.011 | 1 | U | K.RLEVNMK.Q |
| 175 | 171 – 177 | 453.2411 | 904.4677 | 904.4800 | -13.6 | 1 | 3 | 0.52 | 1 | U | K.RLEVNMK.Q + Oxidation (M) |
| 177 | 171 – 177 | 453.2417 | 904.4688 | 904.4800 | -12.4 | 1 | 12 | 0.056 | 1 | U | K.RLEVNMK.Q + Oxidation (M) |
| 179 | 171 – 177 | 453.2421 | 904.4696 | 904.4800 | -11.5 | 1 | 4 | 0.39 | 1 | U | K.RLEVNMK.Q + Oxidation (M) |

| Query | Start – End | Observed | Mr(expt) | Mr(calc) | ppm | M | Score | Expect | Rank | U | Peptide |
|---|---|---|---|---|---|---|---|---|---|---|---|
| 180 | 171 – 177 | 453.2421 | 904.4696 | 904.4800 | -11.5 | 1 | 21 | 0.0076 | 1 | U | K.RLEVNMK.Q + Oxidation (M) |
| 181 | 171 – 177 | 453.2424 | 904.4703 | 904.4800 | -10.7 | 1 | 14 | 0.036 | 1 | U | K.RLEVNMK.Q + Oxidation (M) |
| 185 | 171 – 177 | 453.2427 | 904.4709 | 904.4800 | -10.1 | 1 | 5 | 0.35 | 1 | U | K.RLEVNMK.Q + Oxidation (M) |
| 186 | 171 – 177 | 453.2428 | 904.4711 | 904.4800 | -9.82 | 1 | 9 | 0.12 | 1 | U | K.RLEVNMK.Q + Oxidation (M) |
| 187 | 171 – 177 | 453.2428 | 904.4711 | 904.4800 | -9.82 | 1 | 26 | 0.0025 | 1 | U | K.RLEVNMK.Q + Oxidation (M) |
| 188 | 171 – 177 | 453.2429 | 904.4712 | 904.4800 | -9.71 | 1 | 11 | 0.08 | 1 | U | K.RLEVNMK.Q + Oxidation (M) |
| 190 | 171 – 177 | 453.2429 | 904.4713 | 904.4800 | -9.62 | 1 | 8 | 0.16 | 1 | U | K.RLEVNMK.Q + Oxidation (M) |
| 191 | 171 – 177 | 453.2433 | 904.4720 | 904.4800 | -8.87 | 1 | 8 | 0.15 | 1 | U | K.RLEVNMK.Q + Oxidation (M) |
| 192 | 171 – 177 | 453.2435 | 904.4724 | 904.4800 | -8.36 | 1 | 7 | 0.19 | 1 | U | K.RLEVNMK.Q + Oxidation (M) |
| 196 | 171 – 177 | 453.2448 | 904.4751 | 904.4800 | -5.42 | 1 | 13 | 0.049 | 1 | U | K.RLEVNMK.Q + Oxidation (M) |
| 198 | 171 – 177 | 453.2451 | 904.4757 | 904.4800 | -4.78 | 1 | 2 | 0.56 | 1 | U | K.RLEVNMK.Q + Oxidation (M) |
| 375 | 178 – 189 | 734.8622 | 1467.7098 | 1467.7245 | -10.1 | 0 | 41 | 7.2e-005 | 1 | U | K.QTQPLLDFYSEK.G |
| 376 | 178 – 189 | 734.8631 | 1467.7117 | 1467.7245 | -8.73 | 0 | 35 | 0.00029 | 1 | U | K.QTQPLLDFYSEK.G |
| 377 | 178 – 189 | 734.8632 | 1467.7118 | 1467.7245 | -8.65 | 0 | 32 | 0.00059 | 1 | U | K.QTQPLLDFYSEK.G |
| 378 | 178 – 189 | 734.8632 | 1467.7119 | 1467.7245 | -8.64 | 0 | 49 | 1.2e-005 | 1 | U | K.QTQPLLDFYSEK.G |
| 379 | 178 – 189 | 734.8632 | 1467.7119 | 1467.7245 | -8.58 | 0 | 26 | 0.0023 | 1 | U | K.QTQPLLDFYSEK.G |
| 380 | 178 – 189 | 734.8636 | 1467.7127 | 1467.7245 | -8.08 | 0 | 35 | 0.00035 | 1 | U | K.QTQPLLDFYSEK.G |
| 381 | 178 – 189 | 734.8638 | 1467.7131 | 1467.7245 | -7.79 | 0 | 28 | 0.0016 | 1 | U | K.QTQPLLDFYSEK.G |
| 383 | 178 – 189 | 734.8640 | 1467.7135 | 1467.7245 | -7.52 | 0 | 33 | 0.00051 | 1 | U | K.QTQPLLDFYSEK.G |
| 384 | 178 – 189 | 734.8651 | 1467.7157 | 1467.7245 | -6.04 | 0 | 11 | 0.088 | 1 | U | K.QTQPLLDFYSEK.G |
| 385 | 178 – 189 | 734.8652 | 1467.7159 | 1467.7245 | -5.87 | 0 | 30 | 0.0009 | 1 | U | K.QTQPLLDFYSEK.G |
| 386 | 178 – 189 | 734.8653 | 1467.7161 | 1467.7245 | -5.78 | 0 | 33 | 0.00045 | 1 | U | K.QTQPLLDFYSEK.G |
| 387 | 178 – 189 | 734.8658 | 1467.7171 | 1467.7245 | -5.07 | 0 | 25 | 0.003 | 1 | U | K.QTQPLLDFYSEK.G |
| 388 | 178 – 189 | 734.8663 | 1467.7180 | 1467.7245 | -4.44 | 0 | 20 | 0.01 | 1 | U | K.QTQPLLDFYSEK.G |
| 389 | 178 – 189 | 734.8668 | 1467.7191 | 1467.7245 | -3.73 | 0 | 30 | 0.00092 | 1 | U | K.QTQPLLDFYSEK.G |
| 390 | 178 – 189 | 734.8676 | 1467.7206 | 1467.7245 | -2.70 | 0 | 34 | 0.00038 | 1 | U | K.QTQPLLDFYSEK.G |
| 391 | 178 – 189 | 734.8682 | 1467.7218 | 1467.7245 | -1.84 | 0 | 26 | 0.0026 | 1 | U | K.QTQPLLDFYSEK.G |
| 392 | 178 – 189 | 734.8683 | 1467.7220 | 1467.7245 | -1.76 | 0 | 20 | 0.0097 | 1 | U | K.QTQPLLDFYSEK.G |
| 393 | 178 – 189 | 734.8683 | 1467.7220 | 1467.7245 | -1.76 | 0 | 20 | 0.01 | 1 | U | K.QTQPLLDFYSEK.G |
| 394 | 178 – 189 | 734.8683 | 1467.7221 | 1467.7245 | -1.68 | 0 | 13 | 0.057 | 1 | U | K.QTQPLLDFYSEK.G |
| 395 | 178 – 189 | 734.8700 | 1467.7254 | 1467.7245 | 0.57 | 0 | 30 | 0.00092 | 1 | U | K.QTQPLLDFYSEK.G |
| 396 | 178 – 189 | 734.8752 | 1467.7359 | 1467.7245 | 7.75 | 0 | 7 | 0.24 | 1 | U | K.QTQPLLDFYSEK.G |
| 671 | 190 – 209 | 737.3549 | 2209.0427 | 2209.0651 | -10.1 | 0 | 1 | 1 | 1 | U | K.GYLANVNGQQDIQDVYADVK.D |
| 672 | 190 – 209 | 1105.5292 | 2209.0438 | 2209.0651 | -9.63 | 0 | 41 | 9.6e-005 | 1 | U | K.GYLANVNGQQDIQDVYADVK.D |
| 673 | 190 – 209 | 737.3560 | 2209.0463 | 2209.0651 | -8.53 | 0 | 6 | 0.37 | 1 | U | K.GYLANVNGQQDIQDVYADVK.D |
| 674 | 190 – 209 | 737.3560 | 2209.0463 | 2209.0651 | -8.53 | 0 | 0 | 1.2 | 1 | U | K.GYLANVNGQQDIQDVYADVK.D |
| 675 | 190 – 209 | 1105.5328 | 2209.0510 | 2209.0651 | -6.37 | 0 | 45 | 4.1e-005 | 1 | U | K.GYLANVNGQQDIQDVYADVK.D |
| 676 | 190 – 209 | 1105.5329 | 2209.0512 | 2209.0651 | -6.28 | 0 | 51 | 1.1e-005 | 1 | U | K.GYLANVNGQQDIQDVYADVK.D |
| 677 | 190 – 209 | 1105.5360 | 2209.0574 | 2209.0651 | -3.48 | 0 | 50 | 1.6e-005 | 1 | U | K.GYLANVNGQQDIQDVYADVK.D |
| 678 | 190 – 209 | 1106.0243 | 2210.0340 | 2209.0651 | 439 | 0 | 51 | 1.6e-005 | 1 | U | K.GYLANVNGQQDIQDVYADVK.D |
| 679 | 190 – 209 | 1106.0243 | 2210.0340 | 2209.0651 | 439 | 0 | 44 | 7.4e-005 | 1 | U | K.GYLANVNGQQDIQDVYADVK.D |
| 680 | 190 – 209 | 1106.0259 | 2210.0372 | 2209.0651 | 440 | 0 | 24 | 0.0075 | 1 | U | K.GYLANVNGQQDIQDVYADVK.D |
| 681 | 190 – 209 | 1106.0259 | 2210.0372 | 2209.0651 | 440 | 0 | 5 | 0.66 | 1 | U | K.GYLANVNGQQDIQDVYADVK.D |
| 683 | 190 – 209 | 1106.0267 | 2210.0388 | 2209.0651 | 441 | 0 | 5 | 0.54 | 1 | U | K.GYLANVNGQQDIQDVYADVK.D |
| 684 | 190 – 209 | 1106.0276 | 2210.0406 | 2209.0651 | 442 | 0 | 14 | 0.076 | 1 | U | K.GYLANVNGQQDIQDVYADVK.D |
| 685 | 190 – 209 | 1106.0278 | 2210.0410 | 2209.0651 | 442 | 0 | 20 | 0.017 | 1 | U | K.GYLANVNGQQDIQDVYADVK.D |
| 686 | 190 – 209 | 1106.0281 | 2210.0416 | 2209.0651 | 442 | 0 | 39 | 0.00021 | 1 | U | K.GYLANVNGQQDIQDVYADVK.D |
| 687 | 190 – 209 | 1106.0305 | 2210.0464 | 2209.0651 | 444 | 0 | 28 | 0.0027 | 1 | U | K.GYLANVNGQQDIQDVYADVK.D |
| 795 | 190 – 216 | 969.4919 | 2905.4538 | 2905.4821 | -9.76 | 1 | 1 | 1.2 | 1 | U | K.GYLANVNGQQDIQDVYADVKDLLGGLK.K |
| 799 | 190 – 216 | 969.4948 | 2905.4626 | 2905.4821 | -6.72 | 1 | 1 | 1.4 | 1 | U | K.GYLANVNGQQDIQDVYADVKDLLGGLK.K |
| 800 | 190 – 216 | 969.4991 | 2905.4754 | 2905.4821 | -2.32 | 1 | 4 | 0.69 | 1 | U | K.GYLANVNGQQDIQDVYADVKDLLGGLK.K |
| 803 | 190 – 216 | 969.5026 | 2905.4861 | 2905.4821 | 1.35 | 1 | 10 | 0.12 | 1 | U | K.GYLANVNGQQDIQDVYADVKDLLGGLK.K |
| 807 | 190 – 216 | 969.8206 | 2906.4399 | 2905.4821 | 330 | 1 | 0 | 1.9 | 1 | U | K.GYLANVNGQQDIQDVYADVKDLLGGLK.K |
| 830 | 190 – 217 | 1012.1872 | 3033.5398 | 3033.5771 | -12.3 | 2 | 11 | 0.15 | 1 | U | K.GYLANVNGQQDIQDVYADVKDLLGGLKK.- |
| 832 | 190 – 217 | 1012.1899 | 3033.5479 | 3033.5771 | -9.64 | 2 | 2 | 0.99 | 1 | U | K.GYLANVNGQQDIQDVYADVKDLLGGLKK.- |
| 835 | 190 – 217 | 1012.1954 | 3033.5644 | 3033.5771 | -4.20 | 2 | 3 | 0.67 | 1 | U | K.GYLANVNGQQDIQDVYADVKDLLGGLKK.- |
| 836 | 190 – 217 | 1012.1955 | 3033.5647 | 3033.5771 | -4.10 | 2 | 4 | 0.59 | 1 | U | K.GYLANVNGQQDIQDVYADVKDLLGGLKK.- |
| 838 | 190 – 217 | 1012.1966 | 3033.5680 | 3033.5771 | -3.01 | 2 | 1 | 1.1 | 1 | U | K.GYLANVNGQQDIQDVYADVKDLLGGLKK.- |
| 839 | 190 – 217 | 1012.1976 | 3033.5710 | 3033.5771 | -2.02 | 2 | 21 | 0.0098 | 1 | U | K.GYLANVNGQQDIQDVYADVKDLLGGLKK.- |
| 120 | 210 – 217 | 422.2625 | 842.5105 | 842.5225 | -14.2 | 1 | 18 | 0.017 | 1 | U | K.DLLGGLKK.- |
| 121 | 210 – 217 | 422.2625 | 842.5105 | 842.5225 | -14.2 | 1 | 8 | 0.16 | 1 | U | K.DLLGGLKK.- |
| 122 | 210 – 217 | 422.2638 | 842.5130 | 842.5225 | -11.4 | 1 | 10 | 0.096 | 1 | U | K.DLLGGLKK.- |
| 123 | 210 – 217 | 422.2646 | 842.5146 | 842.5225 | -9.41 | 1 | 9 | 0.14 | 1 | U | K.DLLGGLKK.- |
| 125 | 210 – 217 | 422.2653 | 842.5160 | 842.5225 | -7.75 | 1 | 12 | 0.07 | 1 | U | K.DLLGGLKK.- |
| 126 | 210 – 217 | 422.2656 | 842.5165 | 842.5225 | -7.11 | 1 | 3 | 0.48 | 1 | U | K.DLLGGLKK.- |
| 127 | 210 – 217 | 422.2658 | 842.5170 | 842.5225 | -6.54 | 1 | 2 | 0.7 | 1 | U | K.DLLGGLKK.- |
| 128 | 210 – 217 | 422.2659 | 842.5172 | 842.5225 | -6.35 | 1 | 3 | 0.45 | 1 | U | K.DLLGGLKK.- |
| 129 | 210 – 217 | 422.2659 | 842.5172 | 842.5225 | -6.35 | 1 | 10 | 0.1 | 1 | U | K.DLLGGLKK.- |
| 130 | 210 – 217 | 422.2660 | 842.5174 | 842.5225 | -6.09 | 1 | 15 | 0.035 | 1 | U | K.DLLGGLKK.- |

| Query | Start - End | Observed | Mr(expt) | Mr(calc) | ppm | M | Score | Expect | Rank | U | Peptide |
|---|---|---|---|---|---|---|---|---|---|---|---|
| 131 | 210 - 217 | 422.2660 | 842.5174 | 842.5225 | -6.09 | 1 | 12 | 0.064 | 1 | U | K.DLLGGLKK.- |
| 132 | 210 - 217 | 422.2662 | 842.5179 | 842.5225 | -5.52 | 1 | 10 | 0.1 | 1 | U | K.DLLGGLKK.- |
| 133 | 210 - 217 | 422.2662 | 842.5179 | 842.5225 | -5.52 | 1 | 13 | 0.048 | 1 | U | K.DLLGGLKK.- |
| 134 | 210 - 217 | 843.5259 | 842.5187 | 842.5225 | -4.61 | 1 | 8 | 0.17 | 1 | U | K.DLLGGLKK.- |
| 136 | 210 - 217 | 843.5277 | 842.5204 | 842.5225 | -2.53 | 1 | 5 | 0.32 | 1 | U | K.DLLGGLKK.- |
| 137 | 210 - 217 | 843.5296 | 842.5223 | 842.5225 | -0.25 | 1 | 6 | 0.26 | 1 | U | K.DLLGGLKK.- |
| 139 | 210 - 217 | 843.5369 | 842.5297 | 842.5225 | 8.46 | 1 | 2 | 0.67 | 1 | U | K.DLLGGLKK.- |

Error: try setting browser cache to automatic.

---

```
LOCUS       QNL33349                  47 aa            linear   BCT 04-SEP-2020
DEFINITION  hypothetical protein (plasmid) [Escherichia coli].
ACCESSION   QNL33349
VERSION     QNL33349.1
DBSOURCE    accession MT180430.1
KEYWORDS    .
SOURCE      Escherichia coli
  ORGANISM  Escherichia coli
            Bacteria; Proteobacteria; Gammaproteobacteria; Enterobacterales;
            Enterobacteriaceae; Escherichia.
REFERENCE   1  (residues 1 to 47)
  AUTHORS   Tarabai,H., Wyrsch,E.R., Bitar,I., Djordjevic,S.P. and Dolejska,M.
  TITLE     Comparative analysis of multidrug resistant Escherichia coli ST216
            isolates from silver gulls in Australia
  JOURNAL   Unpublished
REFERENCE   2  (residues 1 to 47)
  AUTHORS   Tarabai,H., Wyrsch,E.R., Bitar,I., Djordjevic,S.P. and Dolejska,M.
  TITLE     Direct Submission
  JOURNAL   Submitted (09-MAR-2020) Department of Biology and Wildlife
            Diseases, University of Veterinary and Pharmaceutical Sciences
            Brno, Palackeho tr. 1946/1, Brno, Brno 61242, Czech Republic
COMMENT     ##Assembly-Data-START##
            Assembly Method       :: SMRT Link v. 8.0
            Assembly Name         :: pCE1681-A
            Coverage              :: 326X
            Sequencing Technology :: PacBio
            ##Assembly-Data-END##
FEATURES             Location/Qualifiers
     source          1..47
                     /organism="Escherichia coli"
                     /strain="CE1681"
                     /host="Chroicocephalus novaehollandiae"
                     /db_xref="taxon:562"
                     /plasmid="pCE1681-A"
                     /country="Australia"
                     /collection_date="2012"
                     /note="Closed IncHI2-ST3/IncN fusion plasmid;
                     type: ST216"
     Protein         1..47
                     /product="hypothetical protein"
     CDS             1..47
                     /coded_by="MT180430.1:235999..236142"
                     /note="hypothetical protein"
                     /transl_table=11
                     /db_xref="SEED:fig|6666666.506717.peg.306"
```

### Species 8

#### MASCOT Search Results

##### Protein View: QNL33349.1

bsAK [Escherichia coli]

| | |
|---|---|
| Database: | NCBIcustTC |
| Score: | 150 |
| Monoisotopic mass ($M_r$): | 24104 |
| Calculated pI: | 4.65 |

Sequencesimilarity is availableas an NCBI BLAST search of QNL33349.1 against nr.

##### Search parameters

| | |
|---|---|
| MS data file: | 09182020_TC-9_001.temp.mgf |
| Enzyme: | Trypsin/P: cuts C-term side of KR. |
| Variable modifications: | Oxidation (M), Propionamide (C), Propionamide (K), Propionamide (N-term) |

##### Protein sequence coverage: 35%

Matched peptides shown in **bold red**.

```
  1 MNLVLMGLPG AGKGTQGERI VEDYGIPHIS TGDMFRAAMK EETPLGLEAK
 51 SYIDKGELVP DEVTIGIVKE RLGKDDCERG FLLDGFPRTV AQAEALEEIL
101 EEYGKPIDYV INIEVDKDVL MERLTGRRIC SVCGTTYHLV FNPPKTPGIC
151 DKDGGELYQR ADDNEETVSK RLEVNMKQTQ PLLDFYSEKG YLANVNGQQD
201 IQDVYADVKD LLGGLKK
```

Unformatted sequencestring: 217 residues (for pasting into other applications).

Sort by (•) residue number ( ) increasing mass ( ) decreasing mass
Show (•) matched peptides only ( ) predicted peptides also
Show (•) uncorrected delta ( ) delta corrected for 13C

| Query | Start – End | Observed | Mr(expt) | Mr(calc) | ppm | M | Score | Expect | Rank | U | Peptide |
|---|---|---|---|---|---|---|---|---|---|---|---|
| 173 | 20 – 36 | 655.9745 | 1964.9016 | 1964.9302 | -14.6 | 0 | 8 | 0.17 | 1 | U | R.IVEDYGIPHISTGDMFR.A + Oxidation (M) |
| 174 | 20 – 36 | 655.9753 | 1964.9041 | 1964.9302 | -13.3 | 0 | 30 | 0.0011 | 1 | U | R.IVEDYGIPHISTGDMFR.A + Oxidation (M) |
| 175 | 20 – 36 | 655.9760 | 1964.9063 | 1964.9302 | -12.2 | 0 | 15 | 0.033 | 1 | U | R.IVEDYGIPHISTGDMFR.A + Oxidation (M) |
| 176 | 20 – 36 | 655.9762 | 1964.9068 | 1964.9302 | -11.9 | 0 | 20 | 0.01 | 1 | U | R.IVEDYGIPHISTGDMFR.A + Oxidation (M) |
| 177 | 20 – 36 | 655.9764 | 1964.9074 | 1964.9302 | -11.6 | 0 | 6 | 0.23 | 1 | U | R.IVEDYGIPHISTGDMFR.A + Oxidation (M) |
| 178 | 20 – 36 | 655.9830 | 1964.9271 | 1964.9302 | -1.61 | 0 | 17 | 0.021 | 1 | U | R.IVEDYGIPHISTGDMFR.A + Oxidation (M) |
| 189 | 51 – 69 | 692.3730 | 2074.0971 | 2074.1198 | -10.9 | 1 | 20 | 0.015 | 1 | U | K.SYIDKGELVPDEVTIGIVK.E |
| 191 | 51 – 69 | 692.3736 | 2074.0989 | 2074.1198 | -10.1 | 1 | 24 | 0.0059 | 1 | U | K.SYIDKGELVPDEVTIGIVK.E |
| 193 | 51 – 69 | 692.3743 | 2074.1011 | 2074.1198 | -8.99 | 1 | 10 | 0.16 | 1 | U | K.SYIDKGELVPDEVTIGIVK.E |
| 195 | 51 – 69 | 692.3746 | 2074.1021 | 2074.1198 | -8.51 | 1 | 18 | 0.026 | 1 | U | K.SYIDKGELVPDEVTIGIVK.E |
| 196 | 51 – 69 | 692.3751 | 2074.1035 | 2074.1198 | -7.83 | 1 | 14 | 0.059 | 1 | U | K.SYIDKGELVPDEVTIGIVK.E |
| 197 | 51 – 69 | 692.3752 | 2074.1038 | 2074.1198 | -7.69 | 1 | 11 | 0.14 | 1 | U | K.SYIDKGELVPDEVTIGIVK.E |
| 198 | 51 – 69 | 1038.0604 | 2074.1062 | 2074.1198 | -6.52 | 1 | 17 | 0.031 | 1 | U | K.SYIDKGELVPDEVTIGIVK.E |
| 84 | 80 – 88 | 511.2732 | 1020.5318 | 1020.5393 | -7.36 | 0 | 5 | 0.34 | 1 | U | R.GFLLDGFPR.T |
| 85 | 80 – 88 | 511.2744 | 1020.5343 | 1020.5393 | -4.91 | 0 | 23 | 0.0051 | 1 | U | R.GFLLDGFPR.T |
| 151 | 178 – 189 | 734.8595 | 1467.7044 | 1467.7245 | -13.7 | 0 | 23 | 0.0052 | 1 | U | K.QTQPLLDFYSEK.G |
| 152 | 178 – 189 | 734.8614 | 1467.7083 | 1467.7245 | -11.1 | 0 | 24 | 0.0042 | 1 | U | K.QTQPLLDFYSEK.G |
| 153 | 178 – 189 | 734.8636 | 1467.7126 | 1467.7245 | -8.14 | 0 | 2 | 0.58 | 1 | U | K.QTQPLLDFYSEK.G |
| 154 | 178 – 189 | 734.8639 | 1467.7133 | 1467.7245 | -7.66 | 0 | 38 | 0.00016 | 1 | U | K.QTQPLLDFYSEK.G |
| 155 | 178 – 189 | 734.8648 | 1467.7150 | 1467.7245 | -6.53 | 0 | 5 | 0.31 | 1 | U | K.QTQPLLDFYSEK.G |
| 156 | 178 – 189 | 734.8652 | 1467.7158 | 1467.7245 | -5.95 | 0 | 19 | 0.013 | 1 | U | K.QTQPLLDFYSEK.G |
| 157 | 178 – 189 | 734.8668 | 1467.7190 | 1467.7245 | -3.80 | 0 | 21 | 0.0085 | 1 | U | K.QTQPLLDFYSEK.G |
| 158 | 178 – 189 | 734.8682 | 1467.7219 | 1467.7245 | -1.81 | 0 | 28 | 0.0015 | 1 | U | K.QTQPLLDFYSEK.G |
| 235 | 190 – 209 | 1105.5278 | 2209.0410 | 2209.0651 | -10.9 | 0 | 3 | 0.51 | 1 | U | K.GYLANVNGQQDIQDVYADVK.D |
| 238 | 190 – 209 | 1105.5307 | 2209.0468 | 2209.0651 | -8.27 | 0 | 8 | 0.18 | 1 | U | K.GYLANVNGQQDIQDVYADVK.D |
| 239 | 190 – 209 | 1105.5345 | 2209.0544 | 2209.0651 | -4.83 | 0 | 5 | 0.32 | 1 | U | K.GYLANVNGQQDIQDVYADVK.D |
| 241 | 190 – 209 | 1106.0259 | 2210.0372 | 2209.0651 | 440 | 0 | 7 | 0.24 | 1 | U | K.GYLANVNGQQDIQDVYADVK.D |

| Query | Start - End | Observed | Mr(expt) | Mr(calc) | ppm | M | Score | Expect | Rank | U | Peptide |
|---|---|---|---|---|---|---|---|---|---|---|---|
| 242 | 190 - 209 | 1106.0343 | 2210.0540 | 2209.0651 | 448 | 0 | 2 | 0.79 | 1 | U | K.GYLANVNGQQDIQDVYADVK.D |

Error: try setting browser cache to automatic.

```
LOCUS       QNL33349                  47 aa            linear   BCT 04-SEP-2020
DEFINITION  hypothetical protein (plasmid) [Escherichia coli].
ACCESSION   QNL33349
VERSION     QNL33349.1
DBSOURCE    accession MT180430.1
KEYWORDS    .
SOURCE      Escherichia coli
  ORGANISM  Escherichia coli
            Bacteria; Proteobacteria; Gammaproteobacteria; Enterobacterales;
            Enterobacteriaceae; Escherichia.
REFERENCE   1  (residues 1 to 47)
  AUTHORS   Tarabai,H., Wyrsch,E.R., Bitar,I., Djordjevic,S.P. and Dolejska,M.
  TITLE     Comparative analysis of multidrug resistant Escherichia coli ST216
            isolates from silver gulls in Australia
  JOURNAL   Unpublished
REFERENCE   2  (residues 1 to 47)
  AUTHORS   Tarabai,H., Wyrsch,E.R., Bitar,I., Djordjevic,S.P. and Dolejska,M.
  TITLE     Direct Submission
  JOURNAL   Submitted (09-MAR-2020) Department of Biology and Wildlife
            Diseases, University of Veterinary and Pharmaceutical Sciences
            Brno, Palackeho tr. 1946/1, Brno, Brno 61242, Czech Republic
COMMENT     ##Assembly-Data-START##
            Assembly Method       :: SMRT Link v. 8.0
            Assembly Name         :: pCE1681-A
            Coverage              :: 326X
            Sequencing Technology :: PacBio
            ##Assembly-Data-END##
FEATURES             Location/Qualifiers
     source          1..47
                     /organism="Escherichia coli"
                     /strain="CE1681"
                     /host="Chroicocephalus novaehollandiae"
                     /db_xref="taxon:562"
                     /plasmid="pCE1681-A"
                     /country="Australia"
                     /collection_date="2012"
                     /note="Closed IncHI2-ST3/IncN fusion plasmid;
                     type: ST216"
     Protein         1..47
                     /product="hypothetical protein"
     CDS             1..47
                     /coded_by="MT180430.1:235999..236142"
                     /note="hypothetical protein"
                     /transl_table=11
                     /db_xref="SEED:fig|6666666.506717.peg.306"
```

### Species 9

#### MATRIX SCIENCE MASCOT Search Results

##### Protein View: QNL33346.1

bsAK-cp45 [Escherichia coli]

| | |
|---|---|
| Database: | NCBIcustTC |
| Score: | 2975 |
| Monoisotopic mass ($M_r$): | 26604 |
| Calculated pI: | 4.78 |

Sequence similarity is available as an NCBI BLAST search of QNL33346.1 against nr.

##### Search parameters

| | |
|---|---|
| MS data file: | 09182020_TC-11_001.temp.mgf |
| Enzyme: | Trypsin/P: cuts C-term side of KR. |
| Variable modifications: | Oxidation (M), Propionamide (C), Propionamide (K), Propionamide (N-term) |

##### Protein sequence coverage: 75%

Matched peptides shown in **bold red**.

```
  1 MGFRIYRETL SRFSCAAQLG LEAKSYIDKG ELVPDEVTIG IVKERLGKDD
 51 CERGFLLDGF PRTVAQAEAL EEILEEYGKP IDYVINIEVD KDVLMERLTG
101 RRICSVCGTT YHLVFNPPKT PGICDKDGGE LYQRADDNEE TVSKRLEVNM
151 KQTQPLLDFY SEKGYLANVN GQQDIQDVYA DVKDLLGGLK KAAAMNLVLM
201 GLPGAGKGTQ GERIVEDYGI PHISTGDMFR AAMKEETPLG
```

Unformatted sequence string: 240 residues (for pasting into other applications).

Sort by ⦿ residue number ○ increasing mass ○ decreasing mass
Show ⦿ matched peptides only ○ predicted peptides also
Show ⦿ uncorrected delta ○ delta corrected for 13C

| Query | Start – End | Observed | Mr(expt) | Mr(calc) | ppm | M | Score | Expect | Rank | U | Peptide |
|---|---|---|---|---|---|---|---|---|---|---|---|
| 267 | 13 – 24 | 619.3089 | 1236.6032 | 1236.6172 | -11.4 | 0 | 3 | 0.53 | 1 | U | R.FSCAAQLGLEAK.S |
| 268 | 13 – 24 | 619.3124 | 1236.6103 | 1236.6172 | -5.63 | 0 | 0 | 0.9 | 1 | U | R.FSCAAQLGLEAK.S |
| 269 | 13 – 24 | 619.3157 | 1236.6169 | 1236.6172 | -0.30 | 0 | 4 | 0.47 | 1 | U | R.FSCAAQLGLEAK.S |
| 271 | 13 – 24 | 654.8296 | 1307.6446 | 1307.6543 | -7.43 | 0 | 1 | 0.98 | 1 | U | R.FSCAAQLGLEAK.S + Propionamide (C) |
| 272 | 13 – 24 | 654.8301 | 1307.6455 | 1307.6543 | -6.73 | 0 | 3 | 0.7 | 1 | U | R.FSCAAQLGLEAK.S + Propionamide (C) |
| 273 | 13 – 24 | 654.8306 | 1307.6467 | 1307.6543 | -5.82 | 0 | 21 | 0.01 | 1 | U | R.FSCAAQLGLEAK.S + Propionamide (C) |
| 274 | 13 – 24 | 654.8316 | 1307.6487 | 1307.6543 | -4.32 | 0 | 16 | 0.035 | 1 | U | R.FSCAAQLGLEAK.S + Propionamide (C) |
| 275 | 13 – 24 | 654.8340 | 1307.6535 | 1307.6543 | -0.67 | 0 | 16 | 0.039 | 1 | U | R.FSCAAQLGLEAK.S + Propionamide (C) |
| 477 | 25 – 43 | 692.3715 | 2074.0927 | 2074.1198 | -13.0 | 1 | 49 | 2.1e-005 | 1 | U | K.SYIDKGELVPDEVTIGIVK.E |
| 478 | 25 – 43 | 692.3734 | 2074.0984 | 2074.1198 | -10.3 | 1 | 39 | 0.0002 | 1 | U | K.SYIDKGELVPDEVTIGIVK.E |
| 479 | 25 – 43 | 692.3735 | 2074.0988 | 2074.1198 | -10.1 | 1 | 37 | 0.00028 | 1 | U | K.SYIDKGELVPDEVTIGIVK.E |
| 480 | 25 – 43 | 692.3740 | 2074.1001 | 2074.1198 | -9.47 | 1 | 20 | 0.015 | 1 | U | K.SYIDKGELVPDEVTIGIVK.E |
| 481 | 25 – 43 | 1038.0575 | 2074.1004 | 2074.1198 | -9.32 | 1 | 39 | 0.00018 | 1 | U | K.SYIDKGELVPDEVTIGIVK.E |
| 482 | 25 – 43 | 692.3745 | 2074.1016 | 2074.1198 | -8.77 | 1 | 5 | 0.44 | 1 | U | K.SYIDKGELVPDEVTIGIVK.E |
| 483 | 25 – 43 | 692.3746 | 2074.1019 | 2074.1198 | -8.62 | 1 | 20 | 0.015 | 1 | U | K.SYIDKGELVPDEVTIGIVK.E |
| 484 | 25 – 43 | 1038.0586 | 2074.1026 | 2074.1198 | -8.26 | 1 | 26 | 0.0041 | 1 | U | K.SYIDKGELVPDEVTIGIVK.E |
| 485 | 25 – 43 | 1038.0596 | 2074.1046 | 2074.1198 | -7.30 | 1 | 25 | 0.0042 | 1 | U | K.SYIDKGELVPDEVTIGIVK.E |
| 486 | 25 – 43 | 692.3755 | 2074.1047 | 2074.1198 | -7.27 | 1 | 64 | 5.8e-007 | 1 | U | K.SYIDKGELVPDEVTIGIVK.E |
| 487 | 25 – 43 | 692.3757 | 2074.1053 | 2074.1198 | -6.97 | 1 | 27 | 0.0028 | 1 | U | K.SYIDKGELVPDEVTIGIVK.E |
| 488 | 25 – 43 | 692.3758 | 2074.1054 | 2074.1198 | -6.92 | 1 | 63 | 7.1e-007 | 1 | U | K.SYIDKGELVPDEVTIGIVK.E |
| 489 | 25 – 43 | 692.3758 | 2074.1055 | 2074.1198 | -6.91 | 1 | 42 | 9.5e-005 | 1 | U | K.SYIDKGELVPDEVTIGIVK.E |
| 490 | 25 – 43 | 1038.0605 | 2074.1064 | 2074.1198 | -6.43 | 1 | 46 | 4e-005 | 1 | U | K.SYIDKGELVPDEVTIGIVK.E |
| 491 | 25 – 43 | 1038.0616 | 2074.1086 | 2074.1198 | -5.37 | 1 | 30 | 0.0012 | 1 | U | K.SYIDKGELVPDEVTIGIVK.E |
| 492 | 25 – 43 | 1038.0623 | 2074.1100 | 2074.1198 | -4.69 | 1 | 51 | 1e-005 | 1 | U | K.SYIDKGELVPDEVTIGIVK.E |
| 493 | 25 – 43 | 1038.0623 | 2074.1100 | 2074.1198 | -4.69 | 1 | 45 | 3.8e-005 | 1 | U | K.SYIDKGELVPDEVTIGIVK.E |
| 494 | 25 – 43 | 692.3776 | 2074.1109 | 2074.1198 | -4.26 | 1 | 50 | 1.1e-005 | 1 | U | K.SYIDKGELVPDEVTIGIVK.E |
| 495 | 25 – 43 | 692.3784 | 2074.1134 | 2074.1198 | -3.08 | 1 | 61 | 8.6e-007 | 1 | U | K.SYIDKGELVPDEVTIGIVK.E |
| 496 | 25 – 43 | 692.3785 | 2074.1138 | 2074.1198 | -2.89 | 1 | 46 | 2.9e-005 | 1 | U | K.SYIDKGELVPDEVTIGIVK.E |
| 497 | 25 – 43 | 1038.0643 | 2074.1140 | 2074.1198 | -2.76 | 1 | 14 | 0.046 | 1 | U | K.SYIDKGELVPDEVTIGIVK.E |
| 498 | 25 – 43 | 1038.0648 | 2074.1150 | 2074.1198 | -2.28 | 1 | 0 | 0.95 | 1 | U | K.SYIDKGELVPDEVTIGIVK.E |

| Query | Start – End | Observed | Mr(expt) | Mr(calc) | ppm | M | Score | Expect | Rank | U | Peptide |
|---|---|---|---|---|---|---|---|---|---|---|---|
| 499 | 25 – 43 | 1038.0651 | 2074.1156 | 2074.1198 | -1.99 | 1 | 57 | 1.8e-006 | 1 | U | K.SYIDKGELVPDEVTIGIVK.E |
| 205 | 54 – 62 | 511.2700 | 1020.5255 | 1020.5393 | -13.5 | 0 | 31 | 0.00085 | 1 | U | R.GFLLDGFPR.T |
| 206 | 54 – 62 | 511.2702 | 1020.5259 | 1020.5393 | -13.1 | 0 | 52 | 6.3e-006 | 1 | U | R.GFLLDGFPR.T |
| 207 | 54 – 62 | 511.2706 | 1020.5266 | 1020.5393 | -12.4 | 0 | 22 | 0.0068 | 1 | U | R.GFLLDGFPR.T |
| 208 | 54 – 62 | 511.2713 | 1020.5279 | 1020.5393 | -11.1 | 0 | 24 | 0.0041 | 1 | U | R.GFLLDGFPR.T |
| 209 | 54 – 62 | 511.2716 | 1020.5286 | 1020.5393 | -10.4 | 0 | 23 | 0.0054 | 1 | U | R.GFLLDGFPR.T |
| 210 | 54 – 62 | 511.2718 | 1020.5290 | 1020.5393 | -10.1 | 0 | 37 | 0.00022 | 1 | U | R.GFLLDGFPR.T |
| 211 | 54 – 62 | 511.2727 | 1020.5308 | 1020.5393 | -8.34 | 0 | 52 | 5.7e-006 | 1 | U | R.GFLLDGFPR.T |
| 212 | 54 – 62 | 511.2730 | 1020.5315 | 1020.5393 | -7.61 | 0 | 36 | 0.00028 | 1 | U | R.GFLLDGFPR.T |
| 213 | 54 – 62 | 511.2730 | 1020.5315 | 1020.5393 | -7.59 | 0 | 26 | 0.0025 | 1 | U | R.GFLLDGFPR.T |
| 214 | 54 – 62 | 511.2731 | 1020.5316 | 1020.5393 | -7.51 | 0 | 53 | 4.9e-006 | 1 | U | R.GFLLDGFPR.T |
| 215 | 54 – 62 | 511.2731 | 1020.5317 | 1020.5393 | -7.38 | 0 | 52 | 6.4e-006 | 1 | U | R.GFLLDGFPR.T |
| 216 | 54 – 62 | 511.2732 | 1020.5318 | 1020.5393 | -7.32 | 0 | 26 | 0.0024 | 1 | U | R.GFLLDGFPR.T |
| 217 | 54 – 62 | 511.2734 | 1020.5323 | 1020.5393 | -6.81 | 0 | 33 | 0.00048 | 1 | U | R.GFLLDGFPR.T |
| 218 | 54 – 62 | 511.2735 | 1020.5324 | 1020.5393 | -6.73 | 0 | 24 | 0.0043 | 1 | U | R.GFLLDGFPR.T |
| 219 | 54 – 62 | 511.2740 | 1020.5334 | 1020.5393 | -5.73 | 0 | 33 | 0.00046 | 1 | U | R.GFLLDGFPR.T |
| 220 | 54 – 62 | 511.2740 | 1020.5335 | 1020.5393 | -5.65 | 0 | 32 | 0.00063 | 1 | U | R.GFLLDGFPR.T |
| 743 | 63 – 97 | 1351.0077 | 4050.0013 | 4050.0394 | -9.42 | 2 | 5 | 0.33 | 1 | U | R.TVAQAEALEEILEEYGKPIDYVINIEVDKDVLMER.L + Oxidation (M) |
| 745 | 63 – 97 | 1351.0120 | 4050.0142 | 4050.0394 | -6.24 | 2 | 6 | 0.28 | 1 | U | R.TVAQAEALEEILEEYGKPIDYVINIEVDKDVLMER.L + Oxidation (M) |
| 746 | 63 – 97 | 1351.0127 | 4050.0163 | 4050.0394 | -5.72 | 2 | 7 | 0.22 | 1 | U | R.TVAQAEALEEILEEYGKPIDYVINIEVDKDVLMER.L + Oxidation (M) |
| 747 | 63 – 97 | 1351.0138 | 4050.0196 | 4050.0394 | -4.90 | 2 | 3 | 0.46 | 1 | U | R.TVAQAEALEEILEEYGKPIDYVINIEVDKDVLMER.L + Oxidation (M) |
| 748 | 63 – 97 | 1351.0150 | 4050.0232 | 4050.0394 | -4.01 | 2 | 5 | 0.35 | 1 | U | R.TVAQAEALEEILEEYGKPIDYVINIEVDKDVLMER.L + Oxidation (M) |
| 749 | 63 – 97 | 1351.0196 | 4050.0370 | 4050.0394 | -0.61 | 2 | 2 | 0.62 | 1 | U | R.TVAQAEALEEILEEYGKPIDYVINIEVDKDVLMER.L + Oxidation (M) |
| 750 | 63 – 97 | 1351.0196 | 4050.0370 | 4050.0394 | -0.61 | 2 | 21 | 0.0081 | 1 | U | R.TVAQAEALEEILEEYGKPIDYVINIEVDKDVLMER.L + Oxidation (M) |
| 751 | 63 – 97 | 1351.0203 | 4050.0391 | 4050.0394 | -0.088 | 2 | 2 | 0.65 | 1 | U | R.TVAQAEALEEILEEYGKPIDYVINIEVDKDVLMER.L + Oxidation (M) |
| 752 | 63 – 97 | 1351.0203 | 4050.0391 | 4050.0394 | -0.088 | 2 | 15 | 0.034 | 1 | U | R.TVAQAEALEEILEEYGKPIDYVINIEVDKDVLMER.L + Oxidation (M) |
| 754 | 63 – 97 | 1351.0255 | 4050.0547 | 4050.0394 | 3.76 | 2 | 13 | 0.054 | 1 | U | R.TVAQAEALEEILEEYGKPIDYVINIEVDKDVLMER.L + Oxidation (M) |
| 755 | 63 – 97 | 1351.0255 | 4050.0547 | 4050.0394 | 3.76 | 2 | 9 | 0.13 | 1 | U | R.TVAQAEALEEILEEYGKPIDYVINIEVDKDVLMER.L + Oxidation (M) |
| 756 | 63 – 97 | 1351.0305 | 4050.0697 | 4050.0394 | 7.47 | 2 | 2 | 0.6 | 1 | U | R.TVAQAEALEEILEEYGKPIDYVINIEVDKDVLMER.L + Oxidation (M) |
| 758 | 63 – 97 | 1351.3418 | 4051.0036 | 4050.0394 | 238 | 2 | 2 | 0.67 | 1 | U | R.TVAQAEALEEILEEYGKPIDYVINIEVDKDVLMER.L + Oxidation (M) |
| 759 | 63 – 97 | 1351.3674 | 4051.0804 | 4050.0394 | 257 | 2 | 4 | 0.43 | 1 | U | R.TVAQAEALEEILEEYGKPIDYVINIEVDKDVLMER.L + Oxidation (M) |
| 381 | 120 – 134 | 574.9338 | 1721.7795 | 1721.8043 | -14.4 | 1 | 18 | 0.014 | 1 | U | K.TPGICDKDGGELYQR.A + Propionamide (K) |
| 384 | 120 – 134 | 574.9365 | 1721.7876 | 1721.8043 | -9.66 | 1 | 2 | 0.65 | 1 | U | K.TPGICDKDGGELYQR.A + Propionamide (C) |
| 385 | 120 – 134 | 574.9376 | 1721.7910 | 1721.8043 | -7.71 | 1 | 10 | 0.12 | 1 | U | K.TPGICDKDGGELYQR.A + Propionamide (C) |
| 386 | 120 – 134 | 574.9378 | 1721.7914 | 1721.8043 | -7.47 | 1 | 20 | 0.01 | 1 | U | K.TPGICDKDGGELYQR.A + Propionamide (K) |
| 387 | 120 – 134 | 574.9378 | 1721.7914 | 1721.8043 | -7.47 | 1 | 2 | 0.69 | 1 | U | K.TPGICDKDGGELYQR.A + Propionamide (K) |
| 121 | 145 – 151 | 445.2455 | 888.4765 | 888.4851 | -9.68 | 1 | 32 | 0.00066 | 1 | U | K.RLEVNMK.Q |
| 122 | 145 – 151 | 445.2475 | 888.4805 | 888.4851 | -5.18 | 1 | 13 | 0.046 | 1 | U | K.RLEVNMK.Q |
| 131 | 145 – 151 | 453.2393 | 904.4641 | 904.4800 | -17.6 | 1 | 18 | 0.015 | 1 | U | K.RLEVNMK.Q + Oxidation (M) |
| 133 | 145 – 151 | 453.2416 | 904.4687 | 904.4800 | -12.5 | 1 | 9 | 0.14 | 1 | U | K.RLEVNMK.Q + Oxidation (M) |
| 134 | 145 – 151 | 453.2419 | 904.4692 | 904.4800 | -11.9 | 1 | 30 | 0.00094 | 1 | U | K.RLEVNMK.Q + Oxidation (M) |
| 135 | 145 – 151 | 453.2422 | 904.4698 | 904.4800 | -11.2 | 1 | 24 | 0.0036 | 1 | U | K.RLEVNMK.Q + Oxidation (M) |
| 136 | 145 – 151 | 453.2423 | 904.4701 | 904.4800 | -11.0 | 1 | 14 | 0.043 | 1 | U | K.RLEVNMK.Q + Oxidation (M) |
| 138 | 145 – 151 | 453.2426 | 904.4706 | 904.4800 | -10.4 | 1 | 11 | 0.089 | 1 | U | K.RLEVNMK.Q + Oxidation (M) |
| 139 | 145 – 151 | 453.2428 | 904.4709 | 904.4800 | -10.0 | 1 | 12 | 0.058 | 1 | U | K.RLEVNMK.Q + Oxidation (M) |
| 140 | 145 – 151 | 453.2428 | 904.4711 | 904.4800 | -9.84 | 1 | 24 | 0.004 | 1 | U | K.RLEVNMK.Q + Oxidation (M) |
| 141 | 145 – 151 | 453.2435 | 904.4723 | 904.4800 | -8.47 | 1 | 11 | 0.072 | 1 | U | K.RLEVNMK.Q + Oxidation (M) |
| 142 | 145 – 151 | 453.2436 | 904.4726 | 904.4800 | -8.14 | 1 | 15 | 0.034 | 1 | U | K.RLEVNMK.Q + Oxidation (M) |
| 143 | 145 – 151 | 453.2436 | 904.4726 | 904.4800 | -8.14 | 1 | 29 | 0.0014 | 1 | U | K.RLEVNMK.Q + Oxidation (M) |
| 144 | 145 – 151 | 453.2436 | 904.4727 | 904.4800 | -8.12 | 1 | 3 | 0.52 | 1 | U | K.RLEVNMK.Q + Oxidation (M) |
| 145 | 145 – 151 | 453.2436 | 904.4727 | 904.4800 | -8.12 | 1 | 1 | 0.87 | 1 | U | K.RLEVNMK.Q + Oxidation (M) |
| 147 | 145 – 151 | 453.2439 | 904.4733 | 904.4800 | -7.37 | 1 | 36 | 0.00026 | 1 | U | K.RLEVNMK.Q + Oxidation (M) |
| 148 | 145 – 151 | 453.2439 | 904.4733 | 904.4800 | -7.37 | 1 | 18 | 0.017 | 1 | U | K.RLEVNMK.Q + Oxidation (M) |
| 149 | 145 – 151 | 453.2440 | 904.4735 | 904.4800 | -7.23 | 1 | 31 | 0.00086 | 1 | U | K.RLEVNMK.Q + Oxidation (M) |
| 150 | 145 – 151 | 453.2444 | 904.4742 | 904.4800 | -6.42 | 1 | 4 | 0.38 | 1 | U | K.RLEVNMK.Q + Oxidation (M) |
| 151 | 145 – 151 | 453.2444 | 904.4742 | 904.4800 | -6.37 | 1 | 10 | 0.091 | 1 | U | K.RLEVNMK.Q + Oxidation (M) |
| 152 | 145 – 151 | 453.2449 | 904.4753 | 904.4800 | -5.22 | 1 | 14 | 0.044 | 1 | U | K.RLEVNMK.Q + Oxidation (M) |
| 154 | 145 – 151 | 453.2459 | 904.4773 | 904.4800 | -2.97 | 1 | 10 | 0.11 | 1 | U | K.RLEVNMK.Q + Oxidation (M) |
| 155 | 145 – 151 | 453.2463 | 904.4781 | 904.4800 | -2.08 | 1 | 23 | 0.0047 | 1 | U | K.RLEVNMK.Q + Oxidation (M) |
| 289 | 152 – 163 | 734.8625 | 1467.7104 | 1467.7245 | -9.63 | 0 | 19 | 0.014 | 1 | U | K.QTQPLLDFYSEK.G |
| 290 | 152 – 163 | 734.8627 | 1467.7108 | 1467.7245 | -9.33 | 0 | 50 | 1e-005 | 1 | U | K.QTQPLLDFYSEK.G |
| 291 | 152 – 163 | 734.8635 | 1467.7124 | 1467.7245 | -8.24 | 0 | 46 | 2.7e-005 | 1 | U | K.QTQPLLDFYSEK.G |
| 292 | 152 – 163 | 734.8639 | 1467.7132 | 1467.7245 | -7.71 | 0 | 17 | 0.022 | 1 | U | K.QTQPLLDFYSEK.G |
| 293 | 152 – 163 | 734.8642 | 1467.7138 | 1467.7245 | -7.33 | 0 | 25 | 0.0035 | 1 | U | K.QTQPLLDFYSEK.G |
| 294 | 152 – 163 | 734.8643 | 1467.7141 | 1467.7245 | -7.14 | 0 | 22 | 0.006 | 1 | U | K.QTQPLLDFYSEK.G |
| 295 | 152 – 163 | 734.8643 | 1467.7141 | 1467.7245 | -7.10 | 0 | 25 | 0.0032 | 1 | U | K.QTQPLLDFYSEK.G |
| 296 | 152 – 163 | 734.8646 | 1467.7146 | 1467.7245 | -6.76 | 0 | 3 | 0.52 | 1 | U | K.QTQPLLDFYSEK.G |
| 297 | 152 – 163 | 734.8651 | 1467.7155 | 1467.7245 | -6.13 | 0 | 31 | 0.00081 | 1 | U | K.QTQPLLDFYSEK.G |
| 298 | 152 – 163 | 734.8653 | 1467.7161 | 1467.7245 | -5.74 | 0 | 40 | 9.4e-005 | 1 | U | K.QTQPLLDFYSEK.G |

| Query | Start – End | Observed | Mr(expt) | Mr(calc) | ppm | M | Score | Expect | Rank | U | Peptide |
|---|---|---|---|---|---|---|---|---|---|---|---|
| 299 | 152 – 163 | 734.8655 | 1467.7164 | 1467.7245 | -5.56 | 0 | 24 | 0.0043 | 1 | U | K.QTQPLLDFYSEK.G |
| 300 | 152 – 163 | 734.8655 | 1467.7164 | 1467.7245 | -5.56 | 0 | 32 | 0.00059 | 1 | U | K.QTQPLLDFYSEK.G |
| 301 | 152 – 163 | 734.8655 | 1467.7164 | 1467.7245 | -5.52 | 0 | 43 | 5.4e-005 | 1 | U | K.QTQPLLDFYSEK.G |
| 302 | 152 – 163 | 734.8659 | 1467.7172 | 1467.7245 | -5.00 | 0 | 22 | 0.0064 | 1 | U | K.QTQPLLDFYSEK.G |
| 303 | 152 – 163 | 734.8662 | 1467.7178 | 1467.7245 | -4.61 | 0 | 24 | 0.0037 | 1 | U | K.QTQPLLDFYSEK.G |
| 304 | 152 – 163 | 734.8663 | 1467.7180 | 1467.7245 | -4.46 | 0 | 26 | 0.0024 | 1 | U | K.QTQPLLDFYSEK.G |
| 305 | 152 – 163 | 734.8664 | 1467.7181 | 1467.7245 | -4.36 | 0 | 31 | 0.00073 | 1 | U | K.QTQPLLDFYSEK.G |
| 306 | 152 – 163 | 734.8676 | 1467.7207 | 1467.7245 | -2.60 | 0 | 46 | 2.4e-005 | 1 | U | K.QTQPLLDFYSEK.G |
| 307 | 152 – 163 | 734.8680 | 1467.7214 | 1467.7245 | -2.15 | 0 | 34 | 0.00038 | 1 | U | K.QTQPLLDFYSEK.G |
| 308 | 152 – 163 | 734.8740 | 1467.7335 | 1467.7245 | 6.12 | 0 | 5 | 0.35 | 1 | U | K.QTQPLLDFYSEK.G |
| 737 | 152 – 183 | 1220.5916 | 3658.7530 | 3658.7791 | -7.15 | 1 | 0 | 1.3 | 1 | U | K.QTQPLLDFYSEKGYLANVNGQQDIQDVYADVK.D |
| 738 | 152 – 183 | 1220.5926 | 3658.7560 | 3658.7791 | -6.33 | 1 | 1 | 1.1 | 1 | U | K.QTQPLLDFYSEKGYLANVNGQQDIQDVYADVK.D |
| 740 | 152 – 183 | 1220.9242 | 3659.7508 | 3658.7791 | 266 | 1 | 2 | 0.76 | 1 | U | K.QTQPLLDFYSEKGYLANVNGQQDIQDVYADVK.D |
| 510 | 164 – 183 | 737.3506 | 2209.0298 | 2209.0651 | -16.0 | 0 | 4 | 0.36 | 1 | U | K.GYLANVNGQQDIQDVYADVK.D |
| 511 | 164 – 183 | 1105.5291 | 2209.0436 | 2209.0651 | -9.72 | 0 | 35 | 0.00034 | 1 | U | K.GYLANVNGQQDIQDVYADVK.D |
| 512 | 164 – 183 | 1105.5291 | 2209.0436 | 2209.0651 | -9.72 | 0 | 34 | 0.00041 | 1 | U | K.GYLANVNGQQDIQDVYADVK.D |
| 513 | 164 – 183 | 1105.5324 | 2209.0502 | 2209.0651 | -6.73 | 0 | 38 | 0.00017 | 1 | U | K.GYLANVNGQQDIQDVYADVK.D |
| 514 | 164 – 183 | 1105.5359 | 2209.0572 | 2209.0651 | -3.57 | 0 | 40 | 0.00012 | 1 | U | K.GYLANVNGQQDIQDVYADVK.D |
| 517 | 164 – 183 | 1106.0253 | 2210.0360 | 2209.0651 | 440 | 0 | 33 | 0.00054 | 1 | U | K.GYLANVNGQQDIQDVYADVK.D |
| 519 | 164 – 183 | 1106.0270 | 2210.0394 | 2209.0651 | 441 | 0 | 29 | 0.0015 | 1 | U | K.GYLANVNGQQDIQDVYADVK.D |
| 520 | 164 – 183 | 1106.0272 | 2210.0398 | 2209.0651 | 441 | 0 | 15 | 0.034 | 1 | U | K.GYLANVNGQQDIQDVYADVK.D |
| 521 | 164 – 183 | 1106.0278 | 2210.0410 | 2209.0651 | 442 | 0 | 11 | 0.1 | 1 | U | K.GYLANVNGQQDIQDVYADVK.D |
| 522 | 164 – 183 | 1106.0282 | 2210.0418 | 2209.0651 | 442 | 0 | 54 | 4.8e-006 | 1 | U | K.GYLANVNGQQDIQDVYADVK.D |
| 523 | 164 – 183 | 1106.0294 | 2210.0442 | 2209.0651 | 443 | 0 | 44 | 5.3e-005 | 1 | U | K.GYLANVNGQQDIQDVYADVK.D |
| 524 | 164 – 183 | 1106.0298 | 2210.0450 | 2209.0651 | 444 | 0 | 36 | 0.00028 | 1 | U | K.GYLANVNGQQDIQDVYADVK.D |
| 525 | 164 – 183 | 1106.0298 | 2210.0450 | 2209.0651 | 444 | 0 | 37 | 0.00023 | 1 | U | K.GYLANVNGQQDIQDVYADVK.D |
| 526 | 164 – 183 | 1106.0300 | 2210.0454 | 2209.0651 | 444 | 0 | 8 | 0.19 | 1 | U | K.GYLANVNGQQDIQDVYADVK.D |
| 527 | 164 – 183 | 1106.0300 | 2210.0454 | 2209.0651 | 444 | 0 | 39 | 0.00014 | 1 | U | K.GYLANVNGQQDIQDVYADVK.D |
| 528 | 164 – 183 | 1106.0328 | 2210.0510 | 2209.0651 | 446 | 0 | 14 | 0.044 | 1 | U | K.GYLANVNGQQDIQDVYADVK.D |
| 602 | 164 – 190 | 969.4942 | 2905.4607 | 2905.4821 | -7.39 | 1 | 3 | 0.62 | 1 | U | K.GYLANVNGQQDIQDVYADVKDLLGGLK.K |
| 607 | 164 – 190 | 969.4968 | 2905.4686 | 2905.4821 | -4.65 | 1 | 1 | 0.99 | 1 | U | K.GYLANVNGQQDIQDVYADVKDLLGGLK.K |
| 614 | 164 – 190 | 969.8246 | 2906.4519 | 2905.4821 | 334 | 1 | 3 | 0.53 | 1 | U | K.GYLANVNGQQDIQDVYADVKDLLGGLK.K |
| 615 | 164 – 190 | 969.8258 | 2906.4555 | 2905.4821 | 335 | 1 | 5 | 0.28 | 1 | U | K.GYLANVNGQQDIQDVYADVKDLLGGLK.K |
| 635 | 164 – 191 | 1012.1881 | 3033.5425 | 3033.5771 | -11.4 | 2 | 2 | 0.82 | 1 | U | K.GYLANVNGQQDIQDVYADVKDLLGGLKK.A |
| 636 | 164 – 191 | 1012.1903 | 3033.5491 | 3033.5771 | -9.24 | 2 | 1 | 1 | 1 | U | K.GYLANVNGQQDIQDVYADVKDLLGGLKK.A |
| 637 | 164 – 191 | 1012.1904 | 3033.5494 | 3033.5771 | -9.14 | 2 | 1 | 1.1 | 1 | U | K.GYLANVNGQQDIQDVYADVKDLLGGLKK.A |
| 642 | 164 – 191 | 1012.1937 | 3033.5593 | 3033.5771 | -5.88 | 2 | 12 | 0.088 | 1 | U | K.GYLANVNGQQDIQDVYADVKDLLGGLKK.A |
| 27 | 184 – 190 | 715.4305 | 714.4233 | 714.4276 | -6.06 | 0 | 7 | 0.21 | 1 | U | K.DLLGGLK.K |
| 106 | 184 – 191 | 422.2635 | 842.5124 | 842.5225 | -12.0 | 1 | 8 | 0.15 | 1 | U | K.DLLGGLKK.A |
| 107 | 184 – 191 | 843.5214 | 842.5141 | 842.5225 | -9.99 | 1 | 2 | 0.59 | 1 | U | K.DLLGGLKK.A |
| 108 | 184 – 191 | 422.2646 | 842.5147 | 842.5225 | -9.27 | 1 | 1 | 0.77 | 1 | U | K.DLLGGLKK.A |
| 109 | 184 – 191 | 422.2651 | 842.5157 | 842.5225 | -8.15 | 1 | 8 | 0.14 | 1 | U | K.DLLGGLKK.A |
| 110 | 184 – 191 | 422.2655 | 842.5164 | 842.5225 | -7.23 | 1 | 8 | 0.18 | 1 | U | K.DLLGGLKK.A |
| 111 | 184 – 191 | 843.5251 | 842.5178 | 842.5225 | -5.59 | 1 | 7 | 0.22 | 1 | U | K.DLLGGLKK.A |
| 316 | 192 – 207 | 757.4083 | 1512.8021 | 1512.8156 | -8.94 | 0 | 37 | 0.00021 | 1 | U | K.AAAMNLVLMGLPGAGK.G |
| 317 | 192 – 207 | 757.4099 | 1512.8053 | 1512.8156 | -6.84 | 0 | 44 | 4e-005 | 1 | U | K.AAAMNLVLMGLPGAGK.G |
| 318 | 192 – 207 | 757.4124 | 1512.8102 | 1512.8156 | -3.59 | 0 | 15 | 0.03 | 1 | U | K.AAAMNLVLMGLPGAGK.G |
| 319 | 192 – 207 | 757.4126 | 1512.8106 | 1512.8156 | -3.31 | 0 | 34 | 0.00042 | 1 | U | K.AAAMNLVLMGLPGAGK.G |
| 320 | 192 – 207 | 757.4127 | 1512.8108 | 1512.8156 | -3.18 | 0 | 15 | 0.032 | 1 | U | K.AAAMNLVLMGLPGAGK.G |
| 321 | 192 – 207 | 757.4136 | 1512.8126 | 1512.8156 | -2.01 | 0 | 23 | 0.0052 | 1 | U | K.AAAMNLVLMGLPGAGK.G |
| 322 | 192 – 207 | 757.4146 | 1512.8146 | 1512.8156 | -0.66 | 0 | 22 | 0.0068 | 1 | U | K.AAAMNLVLMGLPGAGK.G |
| 323 | 192 – 207 | 757.4162 | 1512.8179 | 1512.8156 | 1.49 | 0 | 23 | 0.0054 | 1 | U | K.AAAMNLVLMGLPGAGK.G |
| 328 | 192 – 207 | 765.4040 | 1528.7934 | 1528.8105 | -11.2 | 0 | 32 | 0.00057 | 1 | U | K.AAAMNLVLMGLPGAGK.G + Oxidation (M) |
| 329 | 192 – 207 | 765.4049 | 1528.7952 | 1528.8105 | -10.0 | 0 | 23 | 0.0053 | 1 | U | K.AAAMNLVLMGLPGAGK.G + Oxidation (M) |
| 330 | 192 – 207 | 765.4067 | 1528.7989 | 1528.8105 | -7.60 | 0 | 16 | 0.027 | 1 | U | K.AAAMNLVLMGLPGAGK.G + Oxidation (M) |
| 331 | 192 – 207 | 765.4071 | 1528.7996 | 1528.8105 | -7.13 | 0 | 5 | 0.33 | 1 | U | K.AAAMNLVLMGLPGAGK.G + Oxidation (M) |
| 332 | 192 – 207 | 765.4085 | 1528.8025 | 1528.8105 | -5.27 | 0 | 24 | 0.0037 | 1 | U | K.AAAMNLVLMGLPGAGK.G + Oxidation (M) |
| 333 | 192 – 207 | 765.4088 | 1528.8030 | 1528.8105 | -4.90 | 0 | 6 | 0.25 | 1 | U | K.AAAMNLVLMGLPGAGK.G + Oxidation (M) |
| 334 | 192 – 207 | 765.4090 | 1528.8034 | 1528.8105 | -4.69 | 0 | 22 | 0.0068 | 1 | U | K.AAAMNLVLMGLPGAGK.G + Oxidation (M) |
| 335 | 192 – 207 | 765.4094 | 1528.8043 | 1528.8105 | -4.06 | 0 | 13 | 0.048 | 1 | U | K.AAAMNLVLMGLPGAGK.G + Oxidation (M) |
| 336 | 192 – 207 | 765.4096 | 1528.8046 | 1528.8105 | -3.86 | 0 | 25 | 0.0035 | 1 | U | K.AAAMNLVLMGLPGAGK.G + Oxidation (M) |
| 337 | 192 – 207 | 765.4114 | 1528.8082 | 1528.8105 | -1.55 | 0 | 14 | 0.043 | 1 | U | K.AAAMNLVLMGLPGAGK.G + Oxidation (M) |
| 338 | 192 – 207 | 765.4119 | 1528.8093 | 1528.8105 | -0.83 | 0 | 37 | 0.0002 | 1 | U | K.AAAMNLVLMGLPGAGK.G + Oxidation (M) |
| 339 | 192 – 207 | 765.4141 | 1528.8136 | 1528.8105 | 2.00 | 0 | 38 | 0.00018 | 1 | U | K.AAAMNLVLMGLPGAGK.G + Oxidation (M) |
| 340 | 192 – 207 | 773.4030 | 1544.7915 | 1544.8055 | -9.05 | 0 | 22 | 0.0093 | 1 | U | K.AAAMNLVLMGLPGAGK.G + 2 Oxidation (M) |
| 341 | 192 – 207 | 773.4031 | 1544.7916 | 1544.8055 | -8.97 | 0 | 11 | 0.12 | 1 | U | K.AAAMNLVLMGLPGAGK.G + 2 Oxidation (M) |
| 342 | 192 – 207 | 773.4040 | 1544.7934 | 1544.8055 | -7.78 | 0 | 40 | 0.00014 | 1 | U | K.AAAMNLVLMGLPGAGK.G + 2 Oxidation (M) |
| 343 | 192 – 207 | 773.4053 | 1544.7960 | 1544.8055 | -6.14 | 0 | 23 | 0.0078 | 1 | U | K.AAAMNLVLMGLPGAGK.G + 2 Oxidation (M) |
| 344 | 192 – 207 | 773.4054 | 1544.7961 | 1544.8055 | -6.02 | 0 | 38 | 0.00023 | 1 | U | K.AAAMNLVLMGLPGAGK.G + 2 Oxidation (M) |

| Query | Start – End | Observed | Mr(expt) | Mr(calc) | ppm | M | Score | Expect | Rank | U | Peptide |
|---|---|---|---|---|---|---|---|---|---|---|---|
| 345 | 192 - 207 | 773.4054 | 1544.7963 | 1544.8055 | -5.94 | 0 | 37 | 0.00033 | 1 | U | K.AAAMNLVLMGLPGAGK.G + 2 Oxidation (M) |
| 346 | 192 - 207 | 773.4055 | 1544.7964 | 1544.8055 | -5.87 | 0 | 45 | 5.1e-005 | 1 | U | K.AAAMNLVLMGLPGAGK.G + 2 Oxidation (M) |
| 347 | 192 - 207 | 773.4055 | 1544.7964 | 1544.8055 | -5.87 | 0 | 36 | 0.00039 | 1 | U | K.AAAMNLVLMGLPGAGK.G + 2 Oxidation (M) |
| 348 | 192 - 207 | 773.4058 | 1544.7969 | 1544.8055 | -5.50 | 0 | 27 | 0.003 | 1 | U | K.AAAMNLVLMGLPGAGK.G + 2 Oxidation (M) |
| 349 | 192 - 207 | 773.4059 | 1544.7973 | 1544.8055 | -5.27 | 0 | 32 | 0.00099 | 1 | U | K.AAAMNLVLMGLPGAGK.G + 2 Oxidation (M) |
| 350 | 192 - 207 | 773.4065 | 1544.7984 | 1544.8055 | -4.56 | 0 | 43 | 8.2e-005 | 1 | U | K.AAAMNLVLMGLPGAGK.G + 2 Oxidation (M) |
| 351 | 192 - 207 | 773.4065 | 1544.7984 | 1544.8055 | -4.56 | 0 | 37 | 0.00031 | 1 | U | K.AAAMNLVLMGLPGAGK.G + 2 Oxidation (M) |
| 352 | 192 - 207 | 773.4067 | 1544.7989 | 1544.8055 | -4.26 | 0 | 35 | 0.00052 | 1 | U | K.AAAMNLVLMGLPGAGK.G + 2 Oxidation (M) |
| 353 | 192 - 207 | 773.4077 | 1544.8009 | 1544.8055 | -2.95 | 0 | 38 | 0.00024 | 1 | U | K.AAAMNLVLMGLPGAGK.G + 2 Oxidation (M) |
| 354 | 192 - 207 | 773.4077 | 1544.8009 | 1544.8055 | -2.95 | 0 | 43 | 7.9e-005 | 1 | U | K.AAAMNLVLMGLPGAGK.G + 2 Oxidation (M) |
| 355 | 192 - 207 | 773.4078 | 1544.8011 | 1544.8055 | -2.84 | 0 | 39 | 0.00019 | 1 | U | K.AAAMNLVLMGLPGAGK.G + 2 Oxidation (M) |
| 356 | 192 - 207 | 773.4078 | 1544.8011 | 1544.8055 | -2.84 | 0 | 46 | 4e-005 | 1 | U | K.AAAMNLVLMGLPGAGK.G + 2 Oxidation (M) |
| 357 | 192 - 207 | 773.4080 | 1544.8015 | 1544.8055 | -2.56 | 0 | 17 | 0.027 | 1 | U | K.AAAMNLVLMGLPGAGK.G + 2 Oxidation (M) |
| 412 | 214 - 230 | 650.6468 | 1948.9185 | 1948.9353 | -8.60 | 0 | 19 | 0.02 | 1 | U | R.IVEDYGIPHISTGDMFR.A |
| 413 | 214 - 230 | 650.6468 | 1948.9187 | 1948.9353 | -8.51 | 0 | 11 | 0.11 | 1 | U | R.IVEDYGIPHISTGDMFR.A |
| 414 | 214 - 230 | 650.6470 | 1948.9191 | 1948.9353 | -8.29 | 0 | 30 | 0.0017 | 1 | U | R.IVEDYGIPHISTGDMFR.A |
| 415 | 214 - 230 | 650.6475 | 1948.9205 | 1948.9353 | -7.58 | 0 | 25 | 0.0057 | 1 | U | R.IVEDYGIPHISTGDMFR.A |
| 416 | 214 - 230 | 650.6477 | 1948.9214 | 1948.9353 | -7.15 | 0 | 26 | 0.0048 | 1 | U | R.IVEDYGIPHISTGDMFR.A |
| 417 | 214 - 230 | 650.6482 | 1948.9228 | 1948.9353 | -6.41 | 0 | 4 | 0.69 | 1 | U | R.IVEDYGIPHISTGDMFR.A |
| 418 | 214 - 230 | 975.4689 | 1948.9233 | 1948.9353 | -6.16 | 0 | 8 | 0.29 | 1 | U | R.IVEDYGIPHISTGDMFR.A |
| 419 | 214 - 230 | 650.6484 | 1948.9233 | 1948.9353 | -6.14 | 0 | 44 | 7.5e-005 | 1 | U | R.IVEDYGIPHISTGDMFR.A |
| 420 | 214 - 230 | 975.4690 | 1948.9234 | 1948.9353 | -6.10 | 0 | 27 | 0.0034 | 1 | U | R.IVEDYGIPHISTGDMFR.A |
| 421 | 214 - 230 | 650.6488 | 1948.9245 | 1948.9353 | -5.55 | 0 | 21 | 0.013 | 1 | U | R.IVEDYGIPHISTGDMFR.A |
| 422 | 214 - 230 | 650.6489 | 1948.9250 | 1948.9353 | -5.29 | 0 | 27 | 0.0038 | 1 | U | R.IVEDYGIPHISTGDMFR.A |
| 423 | 214 - 230 | 650.6493 | 1948.9260 | 1948.9353 | -4.80 | 0 | 15 | 0.06 | 1 | U | R.IVEDYGIPHISTGDMFR.A |
| 424 | 214 - 230 | 650.6496 | 1948.9269 | 1948.9353 | -4.29 | 0 | 26 | 0.0045 | 1 | U | R.IVEDYGIPHISTGDMFR.A |
| 425 | 214 - 230 | 975.4717 | 1948.9288 | 1948.9353 | -3.32 | 0 | 7 | 0.35 | 1 | U | R.IVEDYGIPHISTGDMFR.A |
| 426 | 214 - 230 | 650.6505 | 1948.9297 | 1948.9353 | -2.87 | 0 | 19 | 0.022 | 1 | U | R.IVEDYGIPHISTGDMFR.A |
| 427 | 214 - 230 | 975.4724 | 1948.9302 | 1948.9353 | -2.63 | 0 | 30 | 0.0018 | 1 | U | R.IVEDYGIPHISTGDMFR.A |
| 428 | 214 - 230 | 975.4727 | 1948.9308 | 1948.9353 | -2.32 | 0 | 16 | 0.04 | 1 | U | R.IVEDYGIPHISTGDMFR.A |
| 429 | 214 - 230 | 983.4606 | 1964.9065 | 1964.9302 | -12.0 | 0 | 5 | 0.31 | 1 | U | R.IVEDYGIPHISTGDMFR.A + Oxidation (M) |
| 430 | 214 - 230 | 655.9765 | 1964.9078 | 1964.9302 | -11.4 | 0 | 22 | 0.0061 | 1 | U | R.IVEDYGIPHISTGDMFR.A + Oxidation (M) |
| 431 | 214 - 230 | 983.4612 | 1964.9079 | 1964.9302 | -11.4 | 0 | 18 | 0.017 | 1 | U | R.IVEDYGIPHISTGDMFR.A + Oxidation (M) |
| 432 | 214 - 230 | 655.9766 | 1964.9079 | 1964.9302 | -11.4 | 0 | 26 | 0.0024 | 1 | U | R.IVEDYGIPHISTGDMFR.A + Oxidation (M) |
| 433 | 214 - 230 | 655.9767 | 1964.9083 | 1964.9302 | -11.2 | 0 | 40 | 9.7e-005 | 1 | U | R.IVEDYGIPHISTGDMFR.A + Oxidation (M) |
| 434 | 214 - 230 | 655.9768 | 1964.9085 | 1964.9302 | -11.1 | 0 | 32 | 0.00067 | 1 | U | R.IVEDYGIPHISTGDMFR.A + Oxidation (M) |
| 435 | 214 - 230 | 983.4619 | 1964.9092 | 1964.9302 | -10.7 | 0 | 21 | 0.0072 | 1 | U | R.IVEDYGIPHISTGDMFR.A + Oxidation (M) |
| 436 | 214 - 230 | 983.4629 | 1964.9112 | 1964.9302 | -9.69 | 0 | 21 | 0.0089 | 1 | U | R.IVEDYGIPHISTGDMFR.A + Oxidation (M) |
| 437 | 214 - 230 | 983.4631 | 1964.9117 | 1964.9302 | -9.44 | 0 | 27 | 0.0021 | 1 | U | R.IVEDYGIPHISTGDMFR.A + Oxidation (M) |
| 438 | 214 - 230 | 655.9779 | 1964.9118 | 1964.9302 | -9.35 | 0 | 55 | 3.3e-006 | 1 | U | R.IVEDYGIPHISTGDMFR.A + Oxidation (M) |
| 439 | 214 - 230 | 655.9781 | 1964.9123 | 1964.9302 | -9.11 | 0 | 35 | 0.00032 | 1 | U | R.IVEDYGIPHISTGDMFR.A + Oxidation (M) |
| 440 | 214 - 230 | 655.9781 | 1964.9126 | 1964.9302 | -8.99 | 0 | 50 | 1e-005 | 1 | U | R.IVEDYGIPHISTGDMFR.A + Oxidation (M) |
| 441 | 214 - 230 | 655.9782 | 1964.9127 | 1964.9302 | -8.94 | 0 | 40 | 0.00011 | 1 | U | R.IVEDYGIPHISTGDMFR.A + Oxidation (M) |
| 442 | 214 - 230 | 655.9784 | 1964.9134 | 1964.9302 | -8.54 | 0 | 25 | 0.0033 | 1 | U | R.IVEDYGIPHISTGDMFR.A + Oxidation (M) |
| 443 | 214 - 230 | 655.9785 | 1964.9136 | 1964.9302 | -8.47 | 0 | 40 | 0.00011 | 1 | U | R.IVEDYGIPHISTGDMFR.A + Oxidation (M) |
| 444 | 214 - 230 | 983.4641 | 1964.9137 | 1964.9302 | -8.41 | 0 | 21 | 0.0085 | 1 | U | R.IVEDYGIPHISTGDMFR.A + Oxidation (M) |
| 445 | 214 - 230 | 655.9786 | 1964.9139 | 1964.9302 | -8.30 | 0 | 3 | 0.45 | 1 | U | R.IVEDYGIPHISTGDMFR.A + Oxidation (M) |
| 446 | 214 - 230 | 983.4644 | 1964.9143 | 1964.9302 | -8.11 | 0 | 24 | 0.0039 | 1 | U | R.IVEDYGIPHISTGDMFR.A + Oxidation (M) |
| 447 | 214 - 230 | 655.9788 | 1964.9145 | 1964.9302 | -8.03 | 0 | 13 | 0.048 | 1 | U | R.IVEDYGIPHISTGDMFR.A + Oxidation (M) |
| 448 | 214 - 230 | 655.9788 | 1964.9145 | 1964.9302 | -8.00 | 0 | 63 | 5e-007 | 1 | U | R.IVEDYGIPHISTGDMFR.A + Oxidation (M) |
| 449 | 214 - 230 | 655.9789 | 1964.9148 | 1964.9302 | -7.86 | 0 | 54 | 3.9e-006 | 1 | U | R.IVEDYGIPHISTGDMFR.A + Oxidation (M) |
| 450 | 214 - 230 | 655.9793 | 1964.9162 | 1964.9302 | -7.16 | 0 | 46 | 2.4e-005 | 1 | U | R.IVEDYGIPHISTGDMFR.A + Oxidation (M) |
| 451 | 214 - 230 | 655.9795 | 1964.9168 | 1964.9302 | -6.85 | 0 | 63 | 5e-007 | 1 | U | R.IVEDYGIPHISTGDMFR.A + Oxidation (M) |
| 452 | 214 - 230 | 655.9797 | 1964.9174 | 1964.9302 | -6.53 | 0 | 35 | 0.00032 | 1 | U | R.IVEDYGIPHISTGDMFR.A + Oxidation (M) |
| 453 | 214 - 230 | 655.9802 | 1964.9188 | 1964.9302 | -5.81 | 0 | 6 | 0.23 | 1 | U | R.IVEDYGIPHISTGDMFR.A + Oxidation (M) |
| 454 | 214 - 230 | 983.4678 | 1964.9210 | 1964.9302 | -4.70 | 0 | 5 | 0.3 | 1 | U | R.IVEDYGIPHISTGDMFR.A + Oxidation (M) |
| 455 | 214 - 230 | 655.9811 | 1964.9216 | 1964.9302 | -4.41 | 0 | 45 | 3.1e-005 | 1 | U | R.IVEDYGIPHISTGDMFR.A + Oxidation (M) |
| 456 | 214 - 230 | 983.4689 | 1964.9233 | 1964.9302 | -3.52 | 0 | 20 | 0.011 | 1 | U | R.IVEDYGIPHISTGDMFR.A + Oxidation (M) |
| 457 | 214 - 230 | 983.4690 | 1964.9234 | 1964.9302 | -3.45 | 0 | 14 | 0.037 | 1 | U | R.IVEDYGIPHISTGDMFR.A + Oxidation (M) |
| 458 | 214 - 230 | 655.9819 | 1964.9240 | 1964.9302 | -3.16 | 0 | 37 | 0.00018 | 1 | U | R.IVEDYGIPHISTGDMFR.A + Oxidation (M) |
| 459 | 214 - 230 | 983.4696 | 1964.9247 | 1964.9302 | -2.81 | 0 | 15 | 0.031 | 1 | U | R.IVEDYGIPHISTGDMFR.A + Oxidation (M) |
| 460 | 214 - 230 | 983.4702 | 1964.9258 | 1964.9302 | -2.26 | 0 | 17 | 0.022 | 1 | U | R.IVEDYGIPHISTGDMFR.A + Oxidation (M) |
| 461 | 214 - 230 | 983.4724 | 1964.9303 | 1964.9302 | 0.044 | 0 | 19 | 0.012 | 1 | U | R.IVEDYGIPHISTGDMFR.A + Oxidation (M) |
| 221 | 231 - 240 | 1062.4964 | 1061.4891 | 1061.5063 | -16.2 | 1 | 3 | 0.46 | 1 | U | R.AAMKEETPLG.- + Oxidation (M) |
| 222 | 231 - 240 | 531.7534 | 1061.4922 | 1061.5063 | -13.2 | 1 | 1 | 0.86 | 1 | U | R.AAMKEETPLG.- + Oxidation (M) |
| 225 | 231 - 240 | 531.7554 | 1061.4963 | 1061.5063 | -9.40 | 1 | 20 | 0.01 | 1 | U | R.AAMKEETPLG.- + Oxidation (M) |
| 226 | 231 - 240 | 531.7557 | 1061.4968 | 1061.5063 | -8.95 | 1 | 9 | 0.14 | 1 | U | R.AAMKEETPLG.- + Oxidation (M) |
| 227 | 231 - 240 | 531.7557 | 1061.4969 | 1061.5063 | -8.84 | 1 | 4 | 0.37 | 1 | U | R.AAMKEETPLG.- + Oxidation (M) |

| Query | Start - End | Observed | Mr(expt) | Mr(calc) | ppm | M | Score | Expect | Rank | U | Peptide |
|---|---|---|---|---|---|---|---|---|---|---|---|
| 229 | 231 - 240 | 531.7561 | 1061.4976 | 1061.5063 | -8.16 | 1 | 2 | 0.59 | 1 | U | R.AAMKEETPLG.- + Oxidation (M) |
| 231 | 231 - 240 | 531.7570 | 1061.4995 | 1061.5063 | -6.41 | 1 | 7 | 0.21 | 1 | U | R.AAMKEETPLG.- + Oxidation (M) |
| 233 | 231 - 240 | 531.7580 | 1061.5015 | 1061.5063 | -4.48 | 1 | 8 | 0.17 | 1 | U | R.AAMKEETPLG.- + Oxidation (M) |
| 236 | 231 - 240 | 1062.5250 | 1061.5177 | 1061.5063 | 10.8 | 1 | 4 | 0.42 | 1 | U | R.AAMKEETPLG.- + Oxidation (M) |

Error: try setting browser cache to automatic.

---

```
LOCUS       QNL33346                  48 aa            linear   BCT 04-SEP-2020
DEFINITION  hypothetical protein (plasmid) [Escherichia coli].
ACCESSION   QNL33346
VERSION     QNL33346.1
DBSOURCE    accession MT180430.1
KEYWORDS    .
SOURCE      Escherichia coli
  ORGANISM  Escherichia coli
            Bacteria; Proteobacteria; Gammaproteobacteria; Enterobacterales;
            Enterobacteriaceae; Escherichia.
REFERENCE   1  (residues 1 to 48)
  AUTHORS   Tarabai,H., Wyrsch,E.R., Bitar,I., Djordjevic,S.P. and Dolejska,M.
  TITLE     Comparative analysis of multidrug resistant Escherichia coli ST216
            isolates from silver gulls in Australia
  JOURNAL   Unpublished
REFERENCE   2  (residues 1 to 48)
  AUTHORS   Tarabai,H., Wyrsch,E.R., Bitar,I., Djordjevic,S.P. and Dolejska,M.
  TITLE     Direct Submission
  JOURNAL   Submitted (09-MAR-2020) Department of Biology and Wildlife
            Diseases, University of Veterinary and Pharmaceutical Sciences
            Brno, Palackeho tr. 1946/1, Brno, Brno 61242, Czech Republic
COMMENT     ##Assembly-Data-START##
            Assembly Method       :: SMRT Link v. 8.0
            Assembly Name         :: pCE1681-A
            Coverage              :: 326X
            Sequencing Technology :: PacBio
            ##Assembly-Data-END##
FEATURES             Location/Qualifiers
     source          1..48
                     /organism="Escherichia coli"
                     /strain="CE1681"
                     /host="Chroicocephalus novaehollandiae"
                     /db_xref="taxon:562"
                     /plasmid="pCE1681-A"
                     /country="Australia"
                     /collection_date="2012"
                     /note="Closed IncHI2-ST3/IncN fusion plasmid;
                     type: ST216"
     Protein         1..48
                     /product="hypothetical protein"
     CDS             1..48
                     /coded_by="MT180430.1:234558..234704"
                     /note="hypothetical protein"
                     /transl_table=11
                     /db_xref="SEED:fig|6666666.506717.peg.303"
```

### Species 10

#### MATRIX SCIENCE MASCOT Search Results

##### Protein View: QNL33346.1

bsAK-cp45 [Escherichia coli]

| | |
|---|---|
| Database: | NCBIcustTC |
| Score: | 439 |
| Monoisotopic mass ($M_r$): | 26604 |
| Calculated pI: | 4.78 |

Sequence similarity is available as an NCBI BLAST search of QNL33346.1 against nr.

##### Search parameters

| | |
|---|---|
| MS data file: | 09182020_TC-12_001.temp.mgf |
| Enzyme: | Trypsin/P: cuts C-term side of KR. |
| Variable modifications: | Oxidation (M), Propionamide (C), Propionamide (K), Propionamide (N-term) |

##### Protein sequence coverage: 38%

Matched peptides shown in **bold red**.

```
  1 MGFRIYRETL SRFSCAAQLG LEAKSYIDKG ELVPDEVTIG IVKERLGKDD
 51 CERGFLLDGF PRTVAQAEAL EEILEEYGKP IDYVINIEVD KDVLMERLTG
101 RRICSVCGTT YHLVFNPPKT PGICDKDGGE LYQRADDNEE TVSKRLEVNM
151 KQTQPLLDFY SEKGYLANVN GQQDIQDVYA DVKDLLGGLK KAAAMNLVLM
201 GLPGAGKGTQ GERIVEDYGI PHISTGDMFR AAMKEETPLG
```

Unformatted sequence string: 240 residues (for pasting into other applications).

Sort by (•) residue number ( ) increasing mass ( ) decreasing mass
Show (•) matched peptides only ( ) predicted peptides also
Show (•) uncorrected delta ( ) delta corrected for 13C

| Query | Start - End | Observed | Mr(expt) | Mr(calc) | ppm | M | Score | Expect | Rank | U | Peptide |
|---|---|---|---|---|---|---|---|---|---|---|---|
| 146 | 25 - 43 | 1038.0534 | 2074.0922 | 2074.1198 | -13.3 | 1 | 24 | 0.0071 | 1 | U | K.SYIDKGELVPDEVTIGIVK.E |
| 148 | 25 - 43 | 692.3724 | 2074.0953 | 2074.1198 | -11.8 | 1 | 16 | 0.038 | 1 | U | K.SYIDKGELVPDEVTIGIVK.E |
| 149 | 25 - 43 | 1038.0554 | 2074.0962 | 2074.1198 | -11.3 | 1 | 16 | 0.039 | 1 | U | K.SYIDKGELVPDEVTIGIVK.E |
| 150 | 25 - 43 | 692.3735 | 2074.0985 | 2074.1198 | -10.2 | 1 | 4 | 0.56 | 1 | U | K.SYIDKGELVPDEVTIGIVK.E |
| 151 | 25 - 43 | 692.3740 | 2074.1003 | 2074.1198 | -9.41 | 1 | 27 | 0.0032 | 1 | U | K.SYIDKGELVPDEVTIGIVK.E |
| 152 | 25 - 43 | 692.3741 | 2074.1004 | 2074.1198 | -9.32 | 1 | 9 | 0.22 | 1 | U | K.SYIDKGELVPDEVTIGIVK.E |
| 153 | 25 - 43 | 692.3741 | 2074.1005 | 2074.1198 | -9.30 | 1 | 24 | 0.0066 | 1 | U | K.SYIDKGELVPDEVTIGIVK.E |
| 154 | 25 - 43 | 692.3741 | 2074.1006 | 2074.1198 | -9.24 | 1 | 14 | 0.066 | 1 | U | K.SYIDKGELVPDEVTIGIVK.E |
| 155 | 25 - 43 | 692.3745 | 2074.1018 | 2074.1198 | -8.69 | 1 | 10 | 0.15 | 1 | U | K.SYIDKGELVPDEVTIGIVK.E |
| 156 | 25 - 43 | 692.3747 | 2074.1024 | 2074.1198 | -8.37 | 1 | 18 | 0.027 | 1 | U | K.SYIDKGELVPDEVTIGIVK.E |
| 157 | 25 - 43 | 692.3748 | 2074.1027 | 2074.1198 | -8.24 | 1 | 34 | 0.00055 | 1 | U | K.SYIDKGELVPDEVTIGIVK.E |
| 159 | 25 - 43 | 1038.0599 | 2074.1052 | 2074.1198 | -7.01 | 1 | 21 | 0.012 | 1 | U | K.SYIDKGELVPDEVTIGIVK.E |
| 160 | 25 - 43 | 1038.0635 | 2074.1124 | 2074.1198 | -3.54 | 1 | 9 | 0.13 | 1 | U | K.SYIDKGELVPDEVTIGIVK.E |
| 161 | 25 - 43 | 692.3791 | 2074.1154 | 2074.1198 | -2.14 | 1 | 10 | 0.11 | 1 | U | K.SYIDKGELVPDEVTIGIVK.E |
| 162 | 25 - 43 | 1038.0653 | 2074.1160 | 2074.1198 | -1.80 | 1 | 22 | 0.0058 | 1 | U | K.SYIDKGELVPDEVTIGIVK.E |
| 163 | 25 - 43 | 1038.0696 | 2074.1246 | 2074.1198 | 2.35 | 1 | 10 | 0.097 | 1 | U | K.SYIDKGELVPDEVTIGIVK.E |
| 65 | 54 - 62 | 511.2691 | 1020.5236 | 1020.5393 | -15.3 | 0 | 24 | 0.0041 | 1 | U | R.GFLLDGFPR.T |
| 66 | 54 - 62 | 511.2721 | 1020.5297 | 1020.5393 | -9.40 | 0 | 23 | 0.0045 | 1 | U | R.GFLLDGFPR.T |
| 67 | 54 - 62 | 511.2746 | 1020.5346 | 1020.5393 | -4.61 | 0 | 23 | 0.005 | 1 | U | R.GFLLDGFPR.T |
| 87 | 152 - 163 | 734.8622 | 1467.7099 | 1467.7245 | -9.96 | 0 | 26 | 0.0023 | 1 | U | K.QTQPLLDFYSEK.G |
| 88 | 152 - 163 | 734.8630 | 1467.7114 | 1467.7245 | -8.94 | 0 | 11 | 0.084 | 1 | U | K.QTQPLLDFYSEK.G |
| 89 | 152 - 163 | 734.8633 | 1467.7121 | 1467.7245 | -8.49 | 0 | 19 | 0.013 | 1 | U | K.QTQPLLDFYSEK.G |
| 90 | 152 - 163 | 734.8650 | 1467.7154 | 1467.7245 | -6.20 | 0 | 36 | 0.00028 | 1 | U | K.QTQPLLDFYSEK.G |
| 91 | 152 - 163 | 734.8660 | 1467.7175 | 1467.7245 | -4.82 | 0 | 15 | 0.033 | 1 | U | K.QTQPLLDFYSEK.G |
| 92 | 152 - 163 | 734.8663 | 1467.7181 | 1467.7245 | -4.39 | 0 | 20 | 0.009 | 1 | U | K.QTQPLLDFYSEK.G |
| 93 | 152 - 163 | 734.8666 | 1467.7187 | 1467.7245 | -4.01 | 0 | 24 | 0.0036 | 1 | U | K.QTQPLLDFYSEK.G |
| 94 | 152 - 163 | 734.8671 | 1467.7196 | 1467.7245 | -3.38 | 0 | 4 | 0.44 | 1 | U | K.QTQPLLDFYSEK.G |

| Query | Start – End | Observed | Mr(expt) | Mr(calc) | ppm | M | Score | Expect | Rank | U | Peptide |
|---|---|---|---|---|---|---|---|---|---|---|---|
| 95 | 152 – 163 | 734.8677 | 1467.7209 | 1467.7245 | -2.49 | 0 | 37 | 0.00019 | 1 | U | K.QTQPLLDFYSEK.G |
| 168 | 164 – 183 | 1105.5263 | 2209.0380 | 2209.0651 | -12.3 | 0 | 3 | 0.49 | 1 | U | K.GYLANVNGQQDIQDVYADVK.D |
| 170 | 164 – 183 | 1105.5266 | 2209.0386 | 2209.0651 | -12.0 | 0 | 1 | 0.85 | 1 | U | K.GYLANVNGQQDIQDVYADVK.D |
| 171 | 164 – 183 | 1105.5267 | 2209.0388 | 2209.0651 | -11.9 | 0 | 4 | 0.42 | 1 | U | K.GYLANVNGQQDIQDVYADVK.D |
| 172 | 164 – 183 | 1105.5283 | 2209.0420 | 2209.0651 | -10.4 | 0 | 13 | 0.048 | 1 | U | K.GYLANVNGQQDIQDVYADVK.D |
| 173 | 164 – 183 | 1105.5286 | 2209.0426 | 2209.0651 | -10.2 | 0 | 17 | 0.018 | 1 | U | K.GYLANVNGQQDIQDVYADVK.D |
| 174 | 164 – 183 | 1105.5322 | 2209.0498 | 2209.0651 | -6.92 | 0 | 3 | 0.49 | 1 | U | K.GYLANVNGQQDIQDVYADVK.D |
| 175 | 164 – 183 | 1105.5358 | 2209.0570 | 2209.0651 | -3.66 | 0 | 6 | 0.35 | 1 | U | K.GYLANVNGQQDIQDVYADVK.D |
| 177 | 164 – 183 | 1106.0272 | 2210.0398 | 2209.0651 | 441 | 0 | 9 | 0.17 | 1 | U | K.GYLANVNGQQDIQDVYADVK.D |
| 96 | 192 – 207 | 773.4008 | 1544.7870 | 1544.8055 | -11.9 | 0 | 5 | 0.41 | 1 | U | K.AAAMNLVLMGLPGAGK.G + 2 Oxidation (M) |
| 97 | 192 – 207 | 773.4041 | 1544.7936 | 1544.8055 | -7.64 | 0 | 18 | 0.023 | 1 | U | K.AAAMNLVLMGLPGAGK.G + 2 Oxidation (M) |
| 98 | 192 – 207 | 773.4041 | 1544.7936 | 1544.8055 | -7.64 | 0 | 26 | 0.0038 | 1 | U | K.AAAMNLVLMGLPGAGK.G + 2 Oxidation (M) |
| 99 | 192 – 207 | 773.4045 | 1544.7944 | 1544.8055 | -7.17 | 0 | 6 | 0.4 | 1 | U | K.AAAMNLVLMGLPGAGK.G + 2 Oxidation (M) |
| 100 | 192 – 207 | 773.4045 | 1544.7944 | 1544.8055 | -7.17 | 0 | 16 | 0.036 | 1 | U | K.AAAMNLVLMGLPGAGK.G + 2 Oxidation (M) |
| 101 | 192 – 207 | 773.4048 | 1544.7951 | 1544.8055 | -6.72 | 0 | 23 | 0.0072 | 1 | U | K.AAAMNLVLMGLPGAGK.G + 2 Oxidation (M) |
| 102 | 192 – 207 | 773.4051 | 1544.7956 | 1544.8055 | -6.38 | 0 | 5 | 0.54 | 1 | U | K.AAAMNLVLMGLPGAGK.G + 2 Oxidation (M) |
| 103 | 192 – 207 | 773.4052 | 1544.7957 | 1544.8055 | -6.28 | 0 | 15 | 0.051 | 1 | U | K.AAAMNLVLMGLPGAGK.G + 2 Oxidation (M) |
| 104 | 192 – 207 | 773.4060 | 1544.7974 | 1544.8055 | -5.21 | 0 | 10 | 0.16 | 1 | U | K.AAAMNLVLMGLPGAGK.G + 2 Oxidation (M) |
| 105 | 192 – 207 | 773.4061 | 1544.7977 | 1544.8055 | -5.04 | 0 | 33 | 0.00073 | 1 | U | K.AAAMNLVLMGLPGAGK.G + 2 Oxidation (M) |
| 106 | 192 – 207 | 773.4061 | 1544.7977 | 1544.8055 | -5.04 | 0 | 32 | 0.00096 | 1 | U | K.AAAMNLVLMGLPGAGK.G + 2 Oxidation (M) |
| 107 | 192 – 207 | 773.4064 | 1544.7982 | 1544.8055 | -4.69 | 0 | 15 | 0.049 | 1 | U | K.AAAMNLVLMGLPGAGK.G + 2 Oxidation (M) |
| 108 | 192 – 207 | 773.4066 | 1544.7985 | 1544.8055 | -4.47 | 0 | 7 | 0.33 | 1 | U | K.AAAMNLVLMGLPGAGK.G + 2 Oxidation (M) |
| 109 | 192 – 207 | 773.4073 | 1544.8000 | 1544.8055 | -3.52 | 0 | 5 | 0.51 | 1 | U | K.AAAMNLVLMGLPGAGK.G + 2 Oxidation (M) |
| 110 | 192 – 207 | 773.4125 | 1544.8105 | 1544.8055 | 3.27 | 0 | 5 | 0.65 | 1 | U | K.AAAMNLVLMGLPGAGK.G + 2 Oxidation (M) |
| 118 | 214 – 230 | 983.4564 | 1964.8982 | 1964.9302 | -16.3 | 0 | 7 | 0.21 | 1 | U | R.IVEDYGIPHISTGDMFR.A + Oxidation (M) |
| 119 | 214 – 230 | 655.9751 | 1964.9033 | 1964.9302 | -13.7 | 0 | 15 | 0.033 | 1 | U | R.IVEDYGIPHISTGDMFR.A + Oxidation (M) |
| 120 | 214 – 230 | 655.9762 | 1964.9068 | 1964.9302 | -11.9 | 0 | 16 | 0.028 | 1 | U | R.IVEDYGIPHISTGDMFR.A + Oxidation (M) |
| 121 | 214 – 230 | 655.9765 | 1964.9076 | 1964.9302 | -11.5 | 0 | 22 | 0.0067 | 1 | U | R.IVEDYGIPHISTGDMFR.A + Oxidation (M) |
| 122 | 214 – 230 | 983.4615 | 1964.9084 | 1964.9302 | -11.1 | 0 | 26 | 0.0026 | 1 | U | R.IVEDYGIPHISTGDMFR.A + Oxidation (M) |
| 123 | 214 – 230 | 655.9771 | 1964.9094 | 1964.9302 | -10.6 | 0 | 29 | 0.0013 | 1 | U | R.IVEDYGIPHISTGDMFR.A + Oxidation (M) |
| 124 | 214 – 230 | 655.9781 | 1964.9123 | 1964.9302 | -9.11 | 0 | 22 | 0.0069 | 1 | U | R.IVEDYGIPHISTGDMFR.A + Oxidation (M) |
| 125 | 214 – 230 | 655.9782 | 1964.9129 | 1964.9302 | -8.82 | 0 | 34 | 0.0004 | 1 | U | R.IVEDYGIPHISTGDMFR.A + Oxidation (M) |
| 126 | 214 – 230 | 655.9784 | 1964.9135 | 1964.9302 | -8.53 | 0 | 14 | 0.043 | 1 | U | R.IVEDYGIPHISTGDMFR.A + Oxidation (M) |
| 127 | 214 – 230 | 655.9786 | 1964.9139 | 1964.9302 | -8.33 | 0 | 25 | 0.003 | 1 | U | R.IVEDYGIPHISTGDMFR.A + Oxidation (M) |
| 128 | 214 – 230 | 655.9786 | 1964.9139 | 1964.9302 | -8.30 | 0 | 33 | 0.00052 | 1 | U | R.IVEDYGIPHISTGDMFR.A + Oxidation (M) |
| 129 | 214 – 230 | 983.4644 | 1964.9142 | 1964.9302 | -8.14 | 0 | 5 | 0.31 | 1 | U | R.IVEDYGIPHISTGDMFR.A + Oxidation (M) |
| 130 | 214 – 230 | 655.9792 | 1964.9157 | 1964.9302 | -7.37 | 0 | 49 | 1.4e-005 | 1 | U | R.IVEDYGIPHISTGDMFR.A + Oxidation (M) |
| 131 | 214 – 230 | 655.9792 | 1964.9158 | 1964.9302 | -7.35 | 0 | 22 | 0.007 | 1 | U | R.IVEDYGIPHISTGDMFR.A + Oxidation (M) |
| 132 | 214 – 230 | 655.9798 | 1964.9175 | 1964.9302 | -6.48 | 0 | 7 | 0.2 | 1 | U | R.IVEDYGIPHISTGDMFR.A + Oxidation (M) |
| 133 | 214 – 230 | 983.4665 | 1964.9185 | 1964.9302 | -5.95 | 0 | 17 | 0.022 | 1 | U | R.IVEDYGIPHISTGDMFR.A + Oxidation (M) |
| 134 | 214 – 230 | 983.4674 | 1964.9203 | 1964.9302 | -5.03 | 0 | 13 | 0.052 | 1 | U | R.IVEDYGIPHISTGDMFR.A + Oxidation (M) |
| 135 | 214 – 230 | 983.4677 | 1964.9209 | 1964.9302 | -4.75 | 0 | 3 | 0.47 | 1 | U | R.IVEDYGIPHISTGDMFR.A + Oxidation (M) |
| 136 | 214 – 230 | 655.9820 | 1964.9241 | 1964.9302 | -3.12 | 0 | 16 | 0.026 | 1 | U | R.IVEDYGIPHISTGDMFR.A + Oxidation (M) |
| 137 | 214 – 230 | 983.4701 | 1964.9257 | 1964.9302 | -2.32 | 0 | 19 | 0.012 | 1 | U | R.IVEDYGIPHISTGDMFR.A + Oxidation (M) |

Error: try setting browser cache to automatic.

---

```
LOCUS       QNL33346                  48 aa            linear   BCT 04-SEP-2020
DEFINITION  hypothetical protein (plasmid) [Escherichia coli].
ACCESSION   QNL33346
VERSION     QNL33346.1
DBSOURCE    accession MT180430.1
KEYWORDS    .
SOURCE      Escherichia coli
  ORGANISM  Escherichia coli
            Bacteria; Proteobacteria; Gammaproteobacteria; Enterobacterales;
            Enterobacteriaceae; Escherichia.
REFERENCE   1  (residues 1 to 48)
```

```
  AUTHORS   Tarabai,H., Wyrsch,E.R., Bitar,I., Djordjevic,S.P. and Dolejska,M.
  TITLE     Comparative analysis of multidrug resistant Escherichia coli ST216
            isolates from silver gulls in Australia
  JOURNAL   Unpublished
REFERENCE   2  (residues 1 to 48)
  AUTHORS   Tarabai,H., Wyrsch,E.R., Bitar,I., Djordjevic,S.P. and Dolejska,M.
  TITLE     Direct Submission
  JOURNAL   Submitted (09-MAR-2020) Department of Biology and Wildlife
            Diseases, University of Veterinary and Pharmaceutical Sciences
            Brno, Palackeho tr. 1946/1, Brno, Brno 61242, Czech Republic
COMMENT     ##Assembly-Data-START##
            Assembly Method       :: SMRT Link v. 8.0
            Assembly Name         :: pCE1681-A
            Coverage              :: 326X
            Sequencing Technology :: PacBio
            ##Assembly-Data-END##
FEATURES             Location/Qualifiers
     source          1..48
                     /organism="Escherichia coli"
                     /strain="CE1681"
                     /host="Chroicocephalus novaehollandiae"
                     /db_xref="taxon:562"
                     /plasmid="pCE1681-A"
                     /country="Australia"
                     /collection_date="2012"
                     /note="Closed IncHI2-ST3/IncN fusion plasmid;
                     type: ST216"
     Protein         1..48
                     /product="hypothetical protein"
     CDS             1..48
                     /coded_by="MT180430.1:234558..234704"
                     /note="hypothetical protein"
                     /transl_table=11
                     /db_xref="SEED:fig|6666666.506717.peg.303"
```

### Species 11

#### MASCOT Search Results

##### Protein View: QNL33346.1

bsAK-cp45 [Escherichia coli]

| | |
|---|---|
| Database: | NCBIcustTC |
| Score: | 1245 |
| Monoisotopic mass ($M_r$): | 26604 |
| Calculated pI: | 4.78 |

Sequencesimilarity is availableas an NCBI BLAST search of QNL33346.1 against nr.

##### Search parameters

| | |
|---|---|
| MS data file: | 09182020_TC-14_001.temp.mgf |
| Enzyme: | Trypsin/P: cuts C-term side of KR. |
| Variable modifications: | Oxidation (M), Propionamide (C), Propionamide (K), Propionamide (N-term) |

##### Protein sequence coverage: 65%

Matched peptides shown in **bold red**.

```
  1 MGFRIYRETL SRFSCAAQLG LEAKSYIDKG ELVPDEVTIG IVKERLGKDD
 51 CERGFLLDGF PRTVAQAEAL EEILEEYGKP IDYVINIEVD KDVLMERLTG
101 RRICSVCGTT YHLVFNPPKT PGICDKDGGE LYQRADDNEE TVSKRLEVNM
151 KQTQPLLDFY SEKGYLANVN GQQDIQDVYA DVKDLLGGLK KAAAMNLVLM
201 GLPGAGKGTQ GERIVEDYGI PHISTGDMFR AAMKEETPLG
```

Unformatted sequencestring: 240 residues (for pasting into other applications).

Sort by ◉ residue number ○ increasing mass ○ decreasing mass
Show ◉ matched peptides only ○ predicted peptides also
Show ◉ uncorrected delta ○ delta corrected for 13C

| Query | Start - End | Observed | Mr(expt) | Mr(calc) | ppm | M | Score | Expect | Rank | U | Peptide |
|---|---|---|---|---|---|---|---|---|---|---|---|
| 137 | 13 - 24 | 619.3106 | 1236.6065 | 1236.6172 | -8.64 | 0 | 13 | 0.049 | 1 | U | R.FSCAAQLGLEAK.S |
| 256 | 25 - 43 | 692.3716 | 2074.0931 | 2074.1198 | -12.9 | 1 | 28 | 0.003 | 1 | U | K.SYIDKGELVPDEVTIGIVK.E |
| 257 | 25 - 43 | 1038.0555 | 2074.0964 | 2074.1198 | -11.2 | 1 | 49 | 1.9e-005 | 1 | U | K.SYIDKGELVPDEVTIGIVK.E |
| 258 | 25 - 43 | 692.3730 | 2074.0971 | 2074.1198 | -10.9 | 1 | 16 | 0.042 | 1 | U | K.SYIDKGELVPDEVTIGIVK.E |
| 259 | 25 - 43 | 692.3738 | 2074.0995 | 2074.1198 | -9.80 | 1 | 40 | 0.00016 | 1 | U | K.SYIDKGELVPDEVTIGIVK.E |
| 260 | 25 - 43 | 692.3738 | 2074.0997 | 2074.1198 | -9.67 | 1 | 49 | 1.8e-005 | 1 | U | K.SYIDKGELVPDEVTIGIVK.E |
| 261 | 25 - 43 | 692.3744 | 2074.1014 | 2074.1198 | -8.86 | 1 | 37 | 0.00032 | 1 | U | K.SYIDKGELVPDEVTIGIVK.E |
| 262 | 25 - 43 | 692.3750 | 2074.1031 | 2074.1198 | -8.05 | 1 | 20 | 0.014 | 1 | U | K.SYIDKGELVPDEVTIGIVK.E |
| 263 | 25 - 43 | 692.3750 | 2074.1031 | 2074.1198 | -8.02 | 1 | 64 | 6.2e-007 | 1 | U | K.SYIDKGELVPDEVTIGIVK.E |
| 264 | 25 - 43 | 692.3750 | 2074.1033 | 2074.1198 | -7.95 | 1 | 42 | 0.00011 | 1 | U | K.SYIDKGELVPDEVTIGIVK.E |
| 266 | 25 - 43 | 692.3753 | 2074.1040 | 2074.1198 | -7.60 | 1 | 18 | 0.027 | 1 | U | K.SYIDKGELVPDEVTIGIVK.E |
| 267 | 25 - 43 | 1038.0606 | 2074.1066 | 2074.1198 | -6.33 | 1 | 15 | 0.046 | 1 | U | K.SYIDKGELVPDEVTIGIVK.E |
| 268 | 25 - 43 | 1038.0609 | 2074.1072 | 2074.1198 | -6.04 | 1 | 27 | 0.0026 | 1 | U | K.SYIDKGELVPDEVTIGIVK.E |
| 269 | 25 - 43 | 1038.0621 | 2074.1096 | 2074.1198 | -4.89 | 1 | 28 | 0.0019 | 1 | U | K.SYIDKGELVPDEVTIGIVK.E |
| 270 | 25 - 43 | 692.3772 | 2074.1097 | 2074.1198 | -4.84 | 1 | 19 | 0.018 | 1 | U | K.SYIDKGELVPDEVTIGIVK.E |
| 271 | 25 - 43 | 1038.0623 | 2074.1100 | 2074.1198 | -4.69 | 1 | 11 | 0.11 | 1 | U | K.SYIDKGELVPDEVTIGIVK.E |
| 272 | 25 - 43 | 1038.0635 | 2074.1124 | 2074.1198 | -3.54 | 1 | 25 | 0.0034 | 1 | U | K.SYIDKGELVPDEVTIGIVK.E |
| 273 | 25 - 43 | 1038.0673 | 2074.1200 | 2074.1198 | 0.13 | 1 | 19 | 0.014 | 1 | U | K.SYIDKGELVPDEVTIGIVK.E |
| 274 | 25 - 43 | 1038.0729 | 2074.1312 | 2074.1198 | 5.53 | 1 | 15 | 0.031 | 1 | U | K.SYIDKGELVPDEVTIGIVK.E |
| 97 | 54 - 62 | 511.2698 | 1020.5251 | 1020.5393 | -13.9 | 0 | 29 | 0.0012 | 1 | U | R.GFLLDGFPR.T |
| 98 | 54 - 62 | 511.2704 | 1020.5262 | 1020.5393 | -12.8 | 0 | 24 | 0.0045 | 1 | U | R.GFLLDGFPR.T |
| 99 | 54 - 62 | 511.2713 | 1020.5281 | 1020.5393 | -11.0 | 0 | 33 | 0.00054 | 1 | U | R.GFLLDGFPR.T |
| 100 | 54 - 62 | 511.2716 | 1020.5286 | 1020.5393 | -10.5 | 0 | 23 | 0.0047 | 1 | U | R.GFLLDGFPR.T |
| 101 | 54 - 62 | 511.2716 | 1020.5287 | 1020.5393 | -10.4 | 0 | 31 | 0.00085 | 1 | U | R.GFLLDGFPR.T |
| 102 | 54 - 62 | 511.2723 | 1020.5300 | 1020.5393 | -9.10 | 0 | 24 | 0.0042 | 1 | U | R.GFLLDGFPR.T |
| 103 | 54 - 62 | 511.2725 | 1020.5305 | 1020.5393 | -8.57 | 0 | 21 | 0.0078 | 1 | U | R.GFLLDGFPR.T |
| 104 | 54 - 62 | 511.2735 | 1020.5325 | 1020.5393 | -6.65 | 0 | 20 | 0.009 | 1 | U | R.GFLLDGFPR.T |
| 105 | 54 - 62 | 511.2737 | 1020.5328 | 1020.5393 | -6.36 | 0 | 30 | 0.00095 | 1 | U | R.GFLLDGFPR.T |
| 106 | 54 - 62 | 511.2747 | 1020.5349 | 1020.5393 | -4.24 | 0 | 26 | 0.0028 | 1 | U | R.GFLLDGFPR.T |
| 107 | 54 - 62 | 511.2748 | 1020.5351 | 1020.5393 | -4.10 | 0 | 34 | 0.00041 | 1 | U | R.GFLLDGFPR.T |

| Query | Start - End | Observed | Mr(expt) | Mr(calc) | ppm | M | Score | Expect | Rank | U | Peptide |
|---|---|---|---|---|---|---|---|---|---|---|---|
| 321 | 63 - 97 | 1351.0334 | 4050.0784 | 4050.0394 | 9.62 | 2 | 5 | 0.34 | 1 | U | R.TVAQAEALEEILEEYGKPIDYVINIEVDKDVLMER.L + Oxidation (M) |
| 322 | 63 - 97 | 1351.3408 | 4051.0006 | 4050.0394 | 237 | 2 | 9 | 0.14 | 1 | U | R.TVAQAEALEEILEEYGKPIDYVINIEVDKDVLMER.L + Oxidation (M) |
| 56 | 145 - 151 | 453.2406 | 904.4666 | 904.4800 | -14.8 | 1 | 5 | 0.3 | 1 | U | K.RLEVNMK.Q + Oxidation (M) |
| 59 | 145 - 151 | 453.2427 | 904.4708 | 904.4800 | -10.2 | 1 | 13 | 0.048 | 1 | U | K.RLEVNMK.Q + Oxidation (M) |
| 148 | 152 - 163 | 734.8615 | 1467.7085 | 1467.7245 | -10.9 | 0 | 19 | 0.014 | 1 | U | K.QTQPLLDFYSEK.G |
| 149 | 152 - 163 | 734.8640 | 1467.7133 | 1467.7245 | -7.63 | 0 | 17 | 0.018 | 1 | U | K.QTQPLLDFYSEK.G |
| 150 | 152 - 163 | 734.8644 | 1467.7143 | 1467.7245 | -6.96 | 0 | 34 | 0.00044 | 1 | U | K.QTQPLLDFYSEK.G |
| 151 | 152 - 163 | 734.8650 | 1467.7154 | 1467.7245 | -6.21 | 0 | 14 | 0.036 | 1 | U | K.QTQPLLDFYSEK.G |
| 153 | 152 - 163 | 734.8653 | 1467.7161 | 1467.7245 | -5.76 | 0 | 34 | 0.0004 | 1 | U | K.QTQPLLDFYSEK.G |
| 154 | 152 - 163 | 734.8653 | 1467.7161 | 1467.7245 | -5.75 | 0 | 45 | 3.2e-005 | 1 | U | K.QTQPLLDFYSEK.G |
| 155 | 152 - 163 | 734.8654 | 1467.7162 | 1467.7245 | -5.70 | 0 | 34 | 0.00036 | 1 | U | K.QTQPLLDFYSEK.G |
| 156 | 152 - 163 | 734.8655 | 1467.7164 | 1467.7245 | -5.53 | 0 | 19 | 0.013 | 1 | U | K.QTQPLLDFYSEK.G |
| 157 | 152 - 163 | 734.8658 | 1467.7171 | 1467.7245 | -5.10 | 0 | 14 | 0.04 | 1 | U | K.QTQPLLDFYSEK.G |
| 158 | 152 - 163 | 734.8660 | 1467.7175 | 1467.7245 | -4.78 | 0 | 13 | 0.053 | 1 | U | K.QTQPLLDFYSEK.G |
| 159 | 152 - 163 | 734.8664 | 1467.7182 | 1467.7245 | -4.29 | 0 | 38 | 0.00015 | 1 | U | K.QTQPLLDFYSEK.G |
| 160 | 152 - 163 | 734.8669 | 1467.7193 | 1467.7245 | -3.56 | 0 | 24 | 0.0035 | 1 | U | K.QTQPLLDFYSEK.G |
| 161 | 152 - 163 | 734.8672 | 1467.7198 | 1467.7245 | -3.22 | 0 | 21 | 0.0089 | 1 | U | K.QTQPLLDFYSEK.G |
| 162 | 152 - 163 | 734.8680 | 1467.7214 | 1467.7245 | -2.11 | 0 | 28 | 0.0017 | 1 | U | K.QTQPLLDFYSEK.G |
| 163 | 152 - 163 | 734.8695 | 1467.7245 | 1467.7245 | -0.055 | 0 | 5 | 0.32 | 1 | U | K.QTQPLLDFYSEK.G |
| 278 | 164 - 183 | 1105.5280 | 2209.0414 | 2209.0651 | -10.7 | 0 | 48 | 1.8e-005 | 1 | U | K.GYLANVNGQQDIQDVYADVK.D |
| 279 | 164 - 183 | 1105.5302 | 2209.0458 | 2209.0651 | -8.73 | 0 | 6 | 0.26 | 1 | U | K.GYLANVNGQQDIQDVYADVK.D |
| 280 | 164 - 183 | 1105.5310 | 2209.0474 | 2209.0651 | -8.00 | 0 | 17 | 0.022 | 1 | U | K.GYLANVNGQQDIQDVYADVK.D |
| 281 | 164 - 183 | 1105.5316 | 2209.0486 | 2209.0651 | -7.46 | 0 | 8 | 0.16 | 1 | U | K.GYLANVNGQQDIQDVYADVK.D |
| 282 | 164 - 183 | 737.3584 | 2209.0535 | 2209.0651 | -5.27 | 0 | 2 | 0.71 | 1 | U | K.GYLANVNGQQDIQDVYADVK.D |
| 284 | 164 - 183 | 737.3640 | 2209.0702 | 2209.0651 | 2.28 | 0 | 3 | 0.84 | 1 | U | K.GYLANVNGQQDIQDVYADVK.D |
| 285 | 164 - 183 | 1105.5436 | 2209.0726 | 2209.0651 | 3.41 | 0 | 5 | 0.48 | 1 | U | K.GYLANVNGQQDIQDVYADVK.D |
| 286 | 164 - 183 | 1106.0260 | 2210.0374 | 2209.0651 | 440 | 0 | 2 | 0.79 | 1 | U | K.GYLANVNGQQDIQDVYADVK.D |
| 287 | 164 - 183 | 1106.0268 | 2210.0390 | 2209.0651 | 441 | 0 | 37 | 0.00021 | 1 | U | K.GYLANVNGQQDIQDVYADVK.D |
| 289 | 164 - 183 | 1106.0290 | 2210.0434 | 2209.0651 | 443 | 0 | 15 | 0.038 | 1 | U | K.GYLANVNGQQDIQDVYADVK.D |
| 290 | 164 - 183 | 1106.0301 | 2210.0456 | 2209.0651 | 444 | 0 | 9 | 0.14 | 1 | U | K.GYLANVNGQQDIQDVYADVK.D |
| 291 | 164 - 183 | 1106.0314 | 2210.0482 | 2209.0651 | 445 | 0 | 14 | 0.047 | 1 | U | K.GYLANVNGQQDIQDVYADVK.D |
| 292 | 164 - 183 | 1106.0322 | 2210.0498 | 2209.0651 | 446 | 0 | 5 | 0.4 | 1 | U | K.GYLANVNGQQDIQDVYADVK.D |
| 293 | 164 - 183 | 1106.0333 | 2210.0520 | 2209.0651 | 447 | 0 | 16 | 0.028 | 1 | U | K.GYLANVNGQQDIQDVYADVK.D |
| 169 | 192 - 207 | 765.4041 | 1528.7937 | 1528.8105 | -11.0 | 0 | 5 | 0.29 | 1 | U | K.AAAMNLVLMGLPGAGK.G + Oxidation (M) |
| 170 | 192 - 207 | 765.4042 | 1528.7939 | 1528.8105 | -10.9 | 0 | 17 | 0.021 | 1 | U | K.AAAMNLVLMGLPGAGK.G + Oxidation (M) |
| 171 | 192 - 207 | 765.4049 | 1528.7952 | 1528.8105 | -10.0 | 0 | 14 | 0.043 | 1 | U | K.AAAMNLVLMGLPGAGK.G + Oxidation (M) |
| 172 | 192 - 207 | 765.4062 | 1528.7978 | 1528.8105 | -8.31 | 0 | 2 | 0.67 | 1 | U | K.AAAMNLVLMGLPGAGK.G + Oxidation (M) |
| 174 | 192 - 207 | 773.4028 | 1544.7910 | 1544.8055 | -9.35 | 0 | 1 | 1.3 | 1 | U | K.AAAMNLVLMGLPGAGK.G + 2 Oxidation (M) |
| 175 | 192 - 207 | 773.4028 | 1544.7910 | 1544.8055 | -9.32 | 0 | 4 | 0.59 | 1 | U | K.AAAMNLVLMGLPGAGK.G + 2 Oxidation (M) |
| 176 | 192 - 207 | 773.4029 | 1544.7912 | 1544.8055 | -9.19 | 0 | 21 | 0.011 | 1 | U | K.AAAMNLVLMGLPGAGK.G + 2 Oxidation (M) |
| 177 | 192 - 207 | 773.4029 | 1544.7912 | 1544.8055 | -9.19 | 0 | 13 | 0.07 | 1 | U | K.AAAMNLVLMGLPGAGK.G + 2 Oxidation (M) |
| 178 | 192 - 207 | 773.4034 | 1544.7922 | 1544.8055 | -8.56 | 0 | 21 | 0.012 | 1 | U | K.AAAMNLVLMGLPGAGK.G + 2 Oxidation (M) |
| 179 | 192 - 207 | 773.4034 | 1544.7922 | 1544.8055 | -8.56 | 0 | 29 | 0.002 | 1 | U | K.AAAMNLVLMGLPGAGK.G + 2 Oxidation (M) |
| 180 | 192 - 207 | 773.4038 | 1544.7930 | 1544.8055 | -8.05 | 0 | 40 | 0.00015 | 1 | U | K.AAAMNLVLMGLPGAGK.G + 2 Oxidation (M) |
| 181 | 192 - 207 | 773.4045 | 1544.7945 | 1544.8055 | -7.11 | 0 | 43 | 8e-005 | 1 | U | K.AAAMNLVLMGLPGAGK.G + 2 Oxidation (M) |
| 182 | 192 - 207 | 773.4045 | 1544.7945 | 1544.8055 | -7.11 | 0 | 32 | 0.00088 | 1 | U | K.AAAMNLVLMGLPGAGK.G + 2 Oxidation (M) |
| 183 | 192 - 207 | 773.4048 | 1544.7951 | 1544.8055 | -6.72 | 0 | 47 | 3.3e-005 | 1 | U | K.AAAMNLVLMGLPGAGK.G + 2 Oxidation (M) |
| 184 | 192 - 207 | 773.4048 | 1544.7951 | 1544.8055 | -6.72 | 0 | 36 | 0.0004 | 1 | U | K.AAAMNLVLMGLPGAGK.G + 2 Oxidation (M) |
| 185 | 192 - 207 | 773.4050 | 1544.7954 | 1544.8055 | -6.49 | 0 | 19 | 0.019 | 1 | U | K.AAAMNLVLMGLPGAGK.G + 2 Oxidation (M) |
| 186 | 192 - 207 | 773.4051 | 1544.7956 | 1544.8055 | -6.35 | 0 | 21 | 0.012 | 1 | U | K.AAAMNLVLMGLPGAGK.G + 2 Oxidation (M) |
| 187 | 192 - 207 | 773.4056 | 1544.7967 | 1544.8055 | -5.67 | 0 | 33 | 0.00077 | 1 | U | K.AAAMNLVLMGLPGAGK.G + 2 Oxidation (M) |
| 188 | 192 - 207 | 773.4057 | 1544.7968 | 1544.8055 | -5.61 | 0 | 36 | 0.00042 | 1 | U | K.AAAMNLVLMGLPGAGK.G + 2 Oxidation (M) |
| 189 | 192 - 207 | 773.4060 | 1544.7974 | 1544.8055 | -5.22 | 0 | 27 | 0.0028 | 1 | U | K.AAAMNLVLMGLPGAGK.G + 2 Oxidation (M) |
| 190 | 192 - 207 | 773.4072 | 1544.7999 | 1544.8055 | -3.61 | 0 | 5 | 0.54 | 1 | U | K.AAAMNLVLMGLPGAGK.G + 2 Oxidation (M) |
| 191 | 192 - 207 | 773.4079 | 1544.8012 | 1544.8055 | -2.75 | 0 | 13 | 0.08 | 1 | U | K.AAAMNLVLMGLPGAGK.G + 2 Oxidation (M) |
| 212 | 214 - 230 | 650.6440 | 1948.9102 | 1948.9353 | -12.9 | 0 | 13 | 0.073 | 1 | U | R.IVEDYGIPHISTGDMFR.A |
| 213 | 214 - 230 | 650.6447 | 1948.9122 | 1948.9353 | -11.9 | 0 | 3 | 0.8 | 1 | U | R.IVEDYGIPHISTGDMFR.A |
| 214 | 214 - 230 | 650.6459 | 1948.9159 | 1948.9353 | -9.95 | 0 | 1 | 1.1 | 1 | U | R.IVEDYGIPHISTGDMFR.A |
| 215 | 214 - 230 | 650.6464 | 1948.9174 | 1948.9353 | -9.20 | 0 | 25 | 0.0048 | 1 | U | R.IVEDYGIPHISTGDMFR.A |
| 216 | 214 - 230 | 650.6468 | 1948.9187 | 1948.9353 | -8.52 | 0 | 13 | 0.078 | 1 | U | R.IVEDYGIPHISTGDMFR.A |
| 217 | 214 - 230 | 650.6469 | 1948.9188 | 1948.9353 | -8.46 | 0 | 6 | 0.4 | 1 | U | R.IVEDYGIPHISTGDMFR.A |
| 218 | 214 - 230 | 650.6470 | 1948.9191 | 1948.9353 | -8.32 | 0 | 13 | 0.077 | 1 | U | R.IVEDYGIPHISTGDMFR.A |
| 219 | 214 - 230 | 650.6473 | 1948.9201 | 1948.9353 | -7.80 | 0 | 3 | 0.87 | 1 | U | R.IVEDYGIPHISTGDMFR.A |
| 220 | 214 - 230 | 650.6477 | 1948.9213 | 1948.9353 | -7.17 | 0 | 24 | 0.0075 | 1 | U | R.IVEDYGIPHISTGDMFR.A |
| 221 | 214 - 230 | 975.4735 | 1948.9324 | 1948.9353 | -1.50 | 0 | 5 | 0.56 | 1 | U | R.IVEDYGIPHISTGDMFR.A |
| 222 | 214 - 230 | 975.4828 | 1948.9511 | 1948.9353 | 8.09 | 0 | 18 | 0.039 | 1 | U | R.IVEDYGIPHISTGDMFR.A |
| 223 | 214 - 230 | 655.9761 | 1964.9066 | 1964.9302 | -12.0 | 0 | 64 | 3.8e-007 | 1 | U | R.IVEDYGIPHISTGDMFR.A + Oxidation (M) |
| 224 | 214 - 230 | 655.9762 | 1964.9067 | 1964.9302 | -12.0 | 0 | 4 | 0.4 | 1 | U | R.IVEDYGIPHISTGDMFR.A + Oxidation (M) |
| 225 | 214 - 230 | 655.9768 | 1964.9087 | 1964.9302 | -11.0 | 0 | 11 | 0.076 | 1 | U | R.IVEDYGIPHISTGDMFR.A + Oxidation (M) |

| Query | Start – End | Observed | Mr(expt) | Mr(calc) | ppm | M | Score | Expect | Rank | U | Peptide |
|---|---|---|---|---|---|---|---|---|---|---|---|
| 226 | 214 – 230 | 983.4631 | 1964.9117 | 1964.9302 | -9.43 | 0 | 11 | 0.087 | 1 | U | R.IVEDYGIPHISTGDMFR.A + Oxidation (M) |
| 227 | 214 – 230 | 655.9778 | 1964.9117 | 1964.9302 | -9.42 | 0 | 5 | 0.34 | 1 | U | R.IVEDYGIPHISTGDMFR.A + Oxidation (M) |
| 228 | 214 – 230 | 655.9779 | 1964.9120 | 1964.9302 | -9.26 | 0 | 38 | 0.00015 | 1 | U | R.IVEDYGIPHISTGDMFR.A + Oxidation (M) |
| 229 | 214 – 230 | 655.9780 | 1964.9122 | 1964.9302 | -9.17 | 0 | 31 | 0.00079 | 1 | U | R.IVEDYGIPHISTGDMFR.A + Oxidation (M) |
| 230 | 214 – 230 | 655.9780 | 1964.9123 | 1964.9302 | -9.14 | 0 | 36 | 0.00023 | 1 | U | R.IVEDYGIPHISTGDMFR.A + Oxidation (M) |
| 231 | 214 – 230 | 655.9781 | 1964.9125 | 1964.9302 | -9.00 | 0 | 42 | 6e-005 | 1 | U | R.IVEDYGIPHISTGDMFR.A + Oxidation (M) |
| 232 | 214 – 230 | 983.4639 | 1964.9132 | 1964.9302 | -8.64 | 0 | 7 | 0.2 | 1 | U | R.IVEDYGIPHISTGDMFR.A + Oxidation (M) |
| 233 | 214 – 230 | 655.9787 | 1964.9143 | 1964.9302 | -8.10 | 0 | 50 | 1e-005 | 1 | U | R.IVEDYGIPHISTGDMFR.A + Oxidation (M) |
| 234 | 214 – 230 | 655.9788 | 1964.9145 | 1964.9302 | -8.00 | 0 | 23 | 0.0048 | 1 | U | R.IVEDYGIPHISTGDMFR.A + Oxidation (M) |
| 235 | 214 – 230 | 655.9789 | 1964.9148 | 1964.9302 | -7.86 | 0 | 46 | 2.2e-005 | 1 | U | R.IVEDYGIPHISTGDMFR.A + Oxidation (M) |
| 236 | 214 – 230 | 655.9789 | 1964.9149 | 1964.9302 | -7.78 | 0 | 35 | 0.00029 | 1 | U | R.IVEDYGIPHISTGDMFR.A + Oxidation (M) |
| 237 | 214 – 230 | 655.9791 | 1964.9154 | 1964.9302 | -7.57 | 0 | 29 | 0.0012 | 1 | U | R.IVEDYGIPHISTGDMFR.A + Oxidation (M) |
| 239 | 214 – 230 | 655.9793 | 1964.9161 | 1964.9302 | -7.20 | 0 | 23 | 0.0052 | 1 | U | R.IVEDYGIPHISTGDMFR.A + Oxidation (M) |
| 240 | 214 – 230 | 655.9800 | 1964.9183 | 1964.9302 | -6.09 | 0 | 6 | 0.27 | 1 | U | R.IVEDYGIPHISTGDMFR.A + Oxidation (M) |
| 241 | 214 – 230 | 983.4666 | 1964.9187 | 1964.9302 | -5.88 | 0 | 20 | 0.0091 | 1 | U | R.IVEDYGIPHISTGDMFR.A + Oxidation (M) |
| 242 | 214 – 230 | 983.4668 | 1964.9191 | 1964.9302 | -5.65 | 0 | 6 | 0.27 | 1 | U | R.IVEDYGIPHISTGDMFR.A + Oxidation (M) |
| 243 | 214 – 230 | 983.4670 | 1964.9195 | 1964.9302 | -5.44 | 0 | 20 | 0.011 | 1 | U | R.IVEDYGIPHISTGDMFR.A + Oxidation (M) |
| 244 | 214 – 230 | 655.9806 | 1964.9201 | 1964.9302 | -5.16 | 0 | 23 | 0.005 | 1 | U | R.IVEDYGIPHISTGDMFR.A + Oxidation (M) |
| 245 | 214 – 230 | 983.4677 | 1964.9209 | 1964.9302 | -4.73 | 0 | 26 | 0.0023 | 1 | U | R.IVEDYGIPHISTGDMFR.A + Oxidation (M) |
| 246 | 214 – 230 | 655.9810 | 1964.9211 | 1964.9302 | -4.67 | 0 | 4 | 0.4 | 1 | U | R.IVEDYGIPHISTGDMFR.A + Oxidation (M) |
| 247 | 214 – 230 | 983.4679 | 1964.9212 | 1964.9302 | -4.60 | 0 | 6 | 0.26 | 1 | U | R.IVEDYGIPHISTGDMFR.A + Oxidation (M) |
| 249 | 214 – 230 | 983.4719 | 1964.9292 | 1964.9302 | -0.51 | 0 | 6 | 0.22 | 1 | U | R.IVEDYGIPHISTGDMFR.A + Oxidation (M) |
| 250 | 214 – 230 | 983.4723 | 1964.9301 | 1964.9302 | -0.068 | 0 | 9 | 0.11 | 1 | U | R.IVEDYGIPHISTGDMFR.A + Oxidation (M) |
| 109 | 231 – 240 | 531.7518 | 1061.4891 | 1061.5063 | -16.2 | 1 | 1 | 0.89 | 1 | U | R.AAMKEETPLG.- + Oxidation (M) |
| 114 | 231 – 240 | 531.7568 | 1061.4991 | 1061.5063 | -6.74 | 1 | 1 | 0.84 | 1 | U | R.AAMKEETPLG.- + Oxidation (M) |

Error: try setting browser cache to automatic.

---

```
LOCUS       QNL33346                  48 aa            linear   BCT 04-SEP-2020
DEFINITION  hypothetical protein (plasmid) [Escherichia coli].
ACCESSION   QNL33346
VERSION     QNL33346.1
DBSOURCE    accession MT180430.1
KEYWORDS    .
SOURCE      Escherichia coli
  ORGANISM  Escherichia coli
            Bacteria; Proteobacteria; Gammaproteobacteria; Enterobacterales;
            Enterobacteriaceae; Escherichia.
REFERENCE   1  (residues 1 to 48)
  AUTHORS   Tarabai,H., Wyrsch,E.R., Bitar,I., Djordjevic,S.P. and Dolejska,M.
  TITLE     Comparative analysis of multidrug resistant Escherichia coli ST216
            isolates from silver gulls in Australia
  JOURNAL   Unpublished
REFERENCE   2  (residues 1 to 48)
  AUTHORS   Tarabai,H., Wyrsch,E.R., Bitar,I., Djordjevic,S.P. and Dolejska,M.
  TITLE     Direct Submission
  JOURNAL   Submitted (09-MAR-2020) Department of Biology and Wildlife
            Diseases, University of Veterinary and Pharmaceutical Sciences
            Brno, Palackeho tr. 1946/1, Brno, Brno 61242, Czech Republic
COMMENT     ##Assembly-Data-START##
            Assembly Method       :: SMRT Link v. 8.0
            Assembly Name         :: pCE1681-A
            Coverage              :: 326X
            Sequencing Technology :: PacBio
            ##Assembly-Data-END##
FEATURES             Location/Qualifiers
     source          1..48
                     /organism="Escherichia coli"
                     /strain="CE1681"
                     /host="Chroicocephalus novaehollandiae"
                     /db_xref="taxon:562"
                     /plasmid="pCE1681-A"
                     /country="Australia"
                     /collection_date="2012"
                     /note="Closed IncHI2-ST3/IncN fusion plasmid;
```

```
                type: ST216"
Protein         1..48
                /product="hypothetical protein"
CDS             1..48
                /coded_by="MT180430.1:234558..234704"
                /note="hypothetical protein"
                /transl_table=11
                /db_xref="SEED:fig|6666666.506717.peg.303"
```

### Species 12

#### MATRIX SCIENCE MASCOT Search Results

##### Protein View: QNL33346.1

bsAK-cp45 [Escherichia coli]

| | |
|---|---|
| Database: | NCBIcustTC |
| Score: | 1331 |
| Monoisotopic mass ($M_r$): | 26604 |
| Calculated pI: | 4.78 |

Sequence similarity is available as an NCBI BLAST search of QNL33346.1 against nr.

##### Search parameters

| | |
|---|---|
| MS data file: | 09182020_TC-13_001.temp.mgf |
| Enzyme: | Trypsin/P: cuts C-term side of KR. |
| Variable modifications: | Oxidation (M), Propionamide (C), Propionamide (K), Propionamide (N-term) |

##### Protein sequence coverage: 59%

Matched peptides shown in **bold red**.

```
  1 MGFRIYRETL SRFSCAAQLG LEAKSYIDKG ELVPDEVTIG IVKERLGKDD
 51 CERGFLLDGF PRTVAQAEAL EEILEEYGKP IDYVINIEVD KDVLMERLTG
101 RRICSVCGTT YHLVFNPPKT PGICDKDGGE LYQRADDNEE TVSKRLEVNM
151 KQTQPLLDFY SEKGYLANVN GQQDIQDVYA DVKDLLGGLK KAAAMNLVLM
201 GLPGAGKGTQ GERIVEDYGI PHISTGDMFR AAMKEETPLG
```

Unformatted sequence string: 240 residues (for pasting into other applications).

Sort by ◉ residue number ○ increasing mass ○ decreasing mass
Show ◉ matched peptides only ○ predicted peptides also
Show ◉ uncorrected delta ○ delta corrected for 13C

| Query | Start – End | Observed | Mr(expt) | Mr(calc) | ppm | M | Score | Expect | Rank | U | Peptide |
|---|---|---|---|---|---|---|---|---|---|---|---|
| 326 | 25 – 43 | 692.3719 | 2074.0940 | 2074.1198 | -12.4 | 1 | 22 | 0.011 | 1 | U | K.SYIDKGELVPDEVTIGIVK.E |
| 327 | 25 – 43 | 692.3722 | 2074.0948 | 2074.1198 | -12.1 | 1 | 29 | 0.0021 | 1 | U | K.SYIDKGELVPDEVTIGIVK.E |
| 328 | 25 – 43 | 1038.0550 | 2074.0954 | 2074.1198 | -11.7 | 1 | 29 | 0.0021 | 1 | U | K.SYIDKGELVPDEVTIGIVK.E |
| 329 | 25 – 43 | 692.3735 | 2074.0987 | 2074.1198 | -10.2 | 1 | 19 | 0.019 | 1 | U | K.SYIDKGELVPDEVTIGIVK.E |
| 330 | 25 – 43 | 692.3737 | 2074.0993 | 2074.1198 | -9.87 | 1 | 16 | 0.038 | 1 | U | K.SYIDKGELVPDEVTIGIVK.E |
| 331 | 25 – 43 | 1038.0586 | 2074.1026 | 2074.1198 | -8.26 | 1 | 13 | 0.071 | 1 | U | K.SYIDKGELVPDEVTIGIVK.E |
| 332 | 25 – 43 | 1038.0594 | 2074.1042 | 2074.1198 | -7.49 | 1 | 7 | 0.33 | 1 | U | K.SYIDKGELVPDEVTIGIVK.E |
| 333 | 25 – 43 | 692.3754 | 2074.1043 | 2074.1198 | -7.47 | 1 | 24 | 0.0068 | 1 | U | K.SYIDKGELVPDEVTIGIVK.E |
| 334 | 25 – 43 | 692.3755 | 2074.1047 | 2074.1198 | -7.28 | 1 | 32 | 0.00096 | 1 | U | K.SYIDKGELVPDEVTIGIVK.E |
| 335 | 25 – 43 | 692.3755 | 2074.1048 | 2074.1198 | -7.24 | 1 | 39 | 0.00017 | 1 | U | K.SYIDKGELVPDEVTIGIVK.E |
| 336 | 25 – 43 | 1038.0615 | 2074.1084 | 2074.1198 | -5.46 | 1 | 33 | 0.00073 | 1 | U | K.SYIDKGELVPDEVTIGIVK.E |
| 337 | 25 – 43 | 1038.0619 | 2074.1092 | 2074.1198 | -5.08 | 1 | 3 | 0.63 | 1 | U | K.SYIDKGELVPDEVTIGIVK.E |
| 339 | 25 – 43 | 692.3782 | 2074.1126 | 2074.1198 | -3.45 | 1 | 22 | 0.0081 | 1 | U | K.SYIDKGELVPDEVTIGIVK.E |
| 340 | 25 – 43 | 692.3782 | 2074.1129 | 2074.1198 | -3.32 | 1 | 20 | 0.011 | 1 | U | K.SYIDKGELVPDEVTIGIVK.E |
| 341 | 25 – 43 | 1038.0666 | 2074.1186 | 2074.1198 | -0.55 | 1 | 33 | 0.0005 | 1 | U | K.SYIDKGELVPDEVTIGIVK.E |
| 203 | 30 – 43 | 734.9086 | 1467.8027 | 1467.8185 | -10.7 | 0 | 6 | 0.26 | 1 | U | K.GELVPDEVTIGIVK.E |
| 136 | 54 – 62 | 511.2695 | 1020.5245 | 1020.5393 | -14.5 | 0 | 5 | 0.3 | 1 | U | R.GFLLDGFPR.T |
| 137 | 54 – 62 | 511.2701 | 1020.5256 | 1020.5393 | -13.4 | 0 | 35 | 0.00029 | 1 | U | R.GFLLDGFPR.T |
| 138 | 54 – 62 | 511.2706 | 1020.5266 | 1020.5393 | -12.4 | 0 | 26 | 0.0025 | 1 | U | R.GFLLDGFPR.T |
| 139 | 54 – 62 | 511.2706 | 1020.5267 | 1020.5393 | -12.4 | 0 | 19 | 0.014 | 1 | U | R.GFLLDGFPR.T |
| 140 | 54 – 62 | 511.2717 | 1020.5287 | 1020.5393 | -10.3 | 0 | 28 | 0.0014 | 1 | U | R.GFLLDGFPR.T |
| 141 | 54 – 62 | 511.2738 | 1020.5330 | 1020.5393 | -6.14 | 0 | 24 | 0.0043 | 1 | U | R.GFLLDGFPR.T |
| 142 | 54 – 62 | 511.2738 | 1020.5331 | 1020.5393 | -6.00 | 0 | 35 | 0.00031 | 1 | U | R.GFLLDGFPR.T |
| 143 | 54 – 62 | 511.2740 | 1020.5334 | 1020.5393 | -5.77 | 0 | 40 | 0.00011 | 1 | U | R.GFLLDGFPR.T |
| 144 | 54 – 62 | 511.2742 | 1020.5339 | 1020.5393 | -5.28 | 0 | 23 | 0.0052 | 1 | U | R.GFLLDGFPR.T |
| 145 | 54 – 62 | 511.2753 | 1020.5361 | 1020.5393 | -3.10 | 0 | 34 | 0.00038 | 1 | U | R.GFLLDGFPR.T |
| 146 | 54 – 62 | 511.2766 | 1020.5387 | 1020.5393 | -0.56 | 0 | 35 | 0.00031 | 1 | U | R.GFLLDGFPR.T |
| 434 | 63 – 97 | 1351.0043 | 4049.9911 | 4050.0394 | -11.9 | 2 | 3 | 0.54 | 1 | U | R.TVAQAEALEEILEEYGKPIDYVINIEVDKDVLMER.L + Oxidation (M) |
| 435 | 63 – 97 | 1351.0043 | 4049.9911 | 4050.0394 | -11.9 | 2 | 0 | 1.1 | 1 | U | R.TVAQAEALEEILEEYGKPIDYVINIEVDKDVLMER.L + Oxidation (M) |
| 437 | 63 – 97 | 1351.0175 | 4050.0307 | 4050.0394 | -2.16 | 2 | 1 | 0.78 | 1 | U | R.TVAQAEALEEILEEYGKPIDYVINIEVDKDVLMER.L + Oxidation (M) |

| Query | Start – End | Observed | Mr(expt) | Mr(calc) | ppm | M | Score | Expect | Rank | U | Peptide |
|---|---|---|---|---|---|---|---|---|---|---|---|
| 438 | 63 – 97 | 1351.3465 | 4051.0177 | 4050.0394 | 242 | 2 | 8 | 0.17 | 1 | U | R.TVAQAEALEEILEEYGKPIDYVINIEVDKDVLMER.L + Oxidation (M) |
| 440 | 63 – 97 | 1351.3502 | 4051.0288 | 4050.0394 | 244 | 2 | 4 | 0.43 | 1 | U | R.TVAQAEALEEILEEYGKPIDYVINIEVDKDVLMER.L + Oxidation (M) |
| 441 | 63 – 97 | 1351.3544 | 4051.0414 | 4050.0394 | 247 | 2 | 6 | 0.25 | 1 | U | R.TVAQAEALEEILEEYGKPIDYVINIEVDKDVLMER.L + Oxidation (M) |
| 76 | 145 – 151 | 453.2413 | 904.4681 | 904.4800 | -13.1 | 1 | 15 | 0.034 | 1 | U | K.RLEVNMK.Q + Oxidation (M) |
| 77 | 145 – 151 | 453.2416 | 904.4686 | 904.4800 | -12.6 | 1 | 22 | 0.0057 | 1 | U | K.RLEVNMK.Q + Oxidation (M) |
| 79 | 145 – 151 | 453.2449 | 904.4752 | 904.4800 | -5.31 | 1 | 5 | 0.34 | 1 | U | K.RLEVNMK.Q + Oxidation (M) |
| 80 | 145 – 151 | 453.2457 | 904.4767 | 904.4800 | -3.61 | 1 | 4 | 0.4 | 1 | U | K.RLEVNMK.Q + Oxidation (M) |
| 189 | 152 – 163 | 734.8626 | 1467.7107 | 1467.7245 | -9.42 | 0 | 15 | 0.035 | 1 | U | K.QTQPLLDFYSEK.G |
| 190 | 152 – 163 | 734.8629 | 1467.7113 | 1467.7245 | -9.03 | 0 | 15 | 0.034 | 1 | U | K.QTQPLLDFYSEK.G |
| 191 | 152 – 163 | 734.8631 | 1467.7116 | 1467.7245 | -8.79 | 0 | 20 | 0.0091 | 1 | U | K.QTQPLLDFYSEK.G |
| 192 | 152 – 163 | 734.8639 | 1467.7133 | 1467.7245 | -7.66 | 0 | 19 | 0.014 | 1 | U | K.QTQPLLDFYSEK.G |
| 193 | 152 – 163 | 734.8641 | 1467.7136 | 1467.7245 | -7.45 | 0 | 20 | 0.011 | 1 | U | K.QTQPLLDFYSEK.G |
| 194 | 152 – 163 | 734.8641 | 1467.7136 | 1467.7245 | -7.43 | 0 | 20 | 0.0095 | 1 | U | K.QTQPLLDFYSEK.G |
| 195 | 152 – 163 | 734.8644 | 1467.7142 | 1467.7245 | -7.07 | 0 | 50 | 1e-005 | 1 | U | K.QTQPLLDFYSEK.G |
| 196 | 152 – 163 | 734.8654 | 1467.7162 | 1467.7245 | -5.71 | 0 | 23 | 0.0045 | 1 | U | K.QTQPLLDFYSEK.G |
| 197 | 152 – 163 | 734.8654 | 1467.7163 | 1467.7245 | -5.61 | 0 | 38 | 0.00016 | 1 | U | K.QTQPLLDFYSEK.G |
| 198 | 152 – 163 | 734.8663 | 1467.7180 | 1467.7245 | -4.46 | 0 | 28 | 0.0016 | 1 | U | K.QTQPLLDFYSEK.G |
| 199 | 152 – 163 | 734.8663 | 1467.7181 | 1467.7245 | -4.39 | 0 | 17 | 0.021 | 1 | U | K.QTQPLLDFYSEK.G |
| 200 | 152 – 163 | 734.8665 | 1467.7185 | 1467.7245 | -4.10 | 0 | 34 | 0.00038 | 1 | U | K.QTQPLLDFYSEK.G |
| 201 | 152 – 163 | 734.8676 | 1467.7206 | 1467.7245 | -2.66 | 0 | 24 | 0.0035 | 1 | U | K.QTQPLLDFYSEK.G |
| 350 | 164 – 183 | 1105.5262 | 2209.0378 | 2209.0651 | -12.3 | 0 | 29 | 0.0011 | 1 | U | K.GYLANVNGQQDIQDVYADVK.D |
| 351 | 164 – 183 | 1105.5320 | 2209.0494 | 2209.0651 | -7.10 | 0 | 54 | 4e-006 | 1 | U | K.GYLANVNGQQDIQDVYADVK.D |
| 352 | 164 – 183 | 1105.5325 | 2209.0504 | 2209.0651 | -6.64 | 0 | 50 | 1e-005 | 1 | U | K.GYLANVNGQQDIQDVYADVK.D |
| 353 | 164 – 183 | 1105.5326 | 2209.0506 | 2209.0651 | -6.55 | 0 | 12 | 0.061 | 1 | U | K.GYLANVNGQQDIQDVYADVK.D |
| 355 | 164 – 183 | 1106.0197 | 2210.0248 | 2209.0651 | 434 | 0 | 12 | 0.061 | 1 | U | K.GYLANVNGQQDIQDVYADVK.D |
| 356 | 164 – 183 | 1106.0227 | 2210.0308 | 2209.0651 | 437 | 0 | 3 | 0.67 | 1 | U | K.GYLANVNGQQDIQDVYADVK.D |
| 357 | 164 – 183 | 1106.0239 | 2210.0332 | 2209.0651 | 438 | 0 | 9 | 0.16 | 1 | U | K.GYLANVNGQQDIQDVYADVK.D |
| 359 | 164 – 183 | 1106.0257 | 2210.0368 | 2209.0651 | 440 | 0 | 3 | 0.58 | 1 | U | K.GYLANVNGQQDIQDVYADVK.D |
| 360 | 164 – 183 | 1106.0265 | 2210.0384 | 2209.0651 | 441 | 0 | 17 | 0.026 | 1 | U | K.GYLANVNGQQDIQDVYADVK.D |
| 361 | 164 – 183 | 1106.0265 | 2210.0384 | 2209.0651 | 441 | 0 | 13 | 0.055 | 1 | U | K.GYLANVNGQQDIQDVYADVK.D |
| 365 | 164 – 183 | 1106.0269 | 2210.0392 | 2209.0651 | 441 | 0 | 13 | 0.056 | 1 | U | K.GYLANVNGQQDIQDVYADVK.D |
| 367 | 164 – 183 | 1106.0290 | 2210.0434 | 2209.0651 | 443 | 0 | 3 | 0.64 | 1 | U | K.GYLANVNGQQDIQDVYADVK.D |
| 368 | 164 – 183 | 1106.0319 | 2210.0492 | 2209.0651 | 445 | 0 | 16 | 0.028 | 1 | U | K.GYLANVNGQQDIQDVYADVK.D |
| 393 | 164 – 190 | 969.4937 | 2905.4592 | 2905.4821 | -7.90 | 1 | 7 | 0.24 | 1 | U | K.GYLANVNGQQDIQDVYADVKDLLGGLK.K |
| 61 | 184 – 191 | 422.2643 | 842.5141 | 842.5225 | -10.1 | 1 | 7 | 0.22 | 1 | U | K.DLLGGLKK.A |
| 62 | 184 – 191 | 422.2645 | 842.5143 | 842.5225 | -9.72 | 1 | 16 | 0.025 | 1 | U | K.DLLGGLKK.A |
| 63 | 184 – 191 | 422.2645 | 842.5144 | 842.5225 | -9.67 | 1 | 9 | 0.11 | 1 | U | K.DLLGGLKK.A |
| 65 | 184 – 191 | 422.2698 | 842.5251 | 842.5225 | 3.08 | 1 | 10 | 0.096 | 1 | U | K.DLLGGLKK.A |
| 214 | 192 – 207 | 765.4016 | 1528.7886 | 1528.8105 | -14.3 | 0 | 18 | 0.016 | 1 | U | K.AAAMNLVLMGLPGAGK.G + Oxidation (M) |
| 215 | 192 – 207 | 765.4029 | 1528.7912 | 1528.8105 | -12.6 | 0 | 6 | 0.23 | 1 | U | K.AAAMNLVLMGLPGAGK.G + Oxidation (M) |
| 216 | 192 – 207 | 765.4044 | 1528.7942 | 1528.8105 | -10.7 | 0 | 26 | 0.0024 | 1 | U | K.AAAMNLVLMGLPGAGK.G + Oxidation (M) |
| 217 | 192 – 207 | 765.4049 | 1528.7952 | 1528.8105 | -10.0 | 0 | 1 | 0.88 | 1 | U | K.AAAMNLVLMGLPGAGK.G + Oxidation (M) |
| 218 | 192 – 207 | 765.4065 | 1528.7984 | 1528.8105 | -7.90 | 0 | 12 | 0.057 | 1 | U | K.AAAMNLVLMGLPGAGK.G + Oxidation (M) |
| 219 | 192 – 207 | 765.4066 | 1528.7987 | 1528.8105 | -7.72 | 0 | 7 | 0.2 | 1 | U | K.AAAMNLVLMGLPGAGK.G + Oxidation (M) |
| 220 | 192 – 207 | 765.4095 | 1528.8045 | 1528.8105 | -3.97 | 0 | 4 | 0.4 | 1 | U | K.AAAMNLVLMGLPGAGK.G + Oxidation (M) |
| 222 | 192 – 207 | 765.4127 | 1528.8108 | 1528.8105 | 0.17 | 0 | 6 | 0.26 | 1 | U | K.AAAMNLVLMGLPGAGK.G + Oxidation (M) |
| 223 | 192 – 207 | 765.4135 | 1528.8124 | 1528.8105 | 1.20 | 0 | 15 | 0.031 | 1 | U | K.AAAMNLVLMGLPGAGK.G + Oxidation (M) |
| 224 | 192 – 207 | 765.4164 | 1528.8182 | 1528.8105 | 5.05 | 0 | 12 | 0.06 | 1 | U | K.AAAMNLVLMGLPGAGK.G + Oxidation (M) |
| 225 | 192 – 207 | 773.4040 | 1544.7934 | 1544.8055 | -7.77 | 0 | 20 | 0.017 | 1 | U | K.AAAMNLVLMGLPGAGK.G + 2 Oxidation (M) |
| 226 | 192 – 207 | 773.4049 | 1544.7953 | 1544.8055 | -6.54 | 0 | 29 | 0.0021 | 1 | U | K.AAAMNLVLMGLPGAGK.G + 2 Oxidation (M) |
| 227 | 192 – 207 | 773.4049 | 1544.7953 | 1544.8055 | -6.54 | 0 | 35 | 0.00054 | 1 | U | K.AAAMNLVLMGLPGAGK.G + 2 Oxidation (M) |
| 228 | 192 – 207 | 773.4053 | 1544.7960 | 1544.8055 | -6.14 | 0 | 12 | 0.11 | 1 | U | K.AAAMNLVLMGLPGAGK.G + 2 Oxidation (M) |
| 229 | 192 – 207 | 773.4054 | 1544.7961 | 1544.8055 | -6.02 | 0 | 36 | 0.00037 | 1 | U | K.AAAMNLVLMGLPGAGK.G + 2 Oxidation (M) |
| 230 | 192 – 207 | 773.4055 | 1544.7965 | 1544.8055 | -5.80 | 0 | 47 | 2.9e-005 | 1 | U | K.AAAMNLVLMGLPGAGK.G + 2 Oxidation (M) |
| 231 | 192 – 207 | 773.4055 | 1544.7965 | 1544.8055 | -5.80 | 0 | 37 | 0.00034 | 1 | U | K.AAAMNLVLMGLPGAGK.G + 2 Oxidation (M) |
| 232 | 192 – 207 | 773.4055 | 1544.7965 | 1544.8055 | -5.80 | 0 | 33 | 0.00074 | 1 | U | K.AAAMNLVLMGLPGAGK.G + 2 Oxidation (M) |
| 233 | 192 – 207 | 773.4056 | 1544.7967 | 1544.8055 | -5.67 | 0 | 36 | 0.00035 | 1 | U | K.AAAMNLVLMGLPGAGK.G + 2 Oxidation (M) |
| 234 | 192 – 207 | 773.4056 | 1544.7967 | 1544.8055 | -5.67 | 0 | 41 | 0.00014 | 1 | U | K.AAAMNLVLMGLPGAGK.G + 2 Oxidation (M) |
| 235 | 192 – 207 | 773.4058 | 1544.7970 | 1544.8055 | -5.46 | 0 | 12 | 0.11 | 1 | U | K.AAAMNLVLMGLPGAGK.G + 2 Oxidation (M) |
| 236 | 192 – 207 | 773.4062 | 1544.7978 | 1544.8055 | -4.97 | 0 | 36 | 0.00035 | 1 | U | K.AAAMNLVLMGLPGAGK.G + 2 Oxidation (M) |
| 237 | 192 – 207 | 773.4062 | 1544.7978 | 1544.8055 | -4.97 | 0 | 42 | 0.0001 | 1 | U | K.AAAMNLVLMGLPGAGK.G + 2 Oxidation (M) |
| 238 | 192 – 207 | 773.4062 | 1544.7978 | 1544.8055 | -4.95 | 0 | 40 | 0.00014 | 1 | U | K.AAAMNLVLMGLPGAGK.G + 2 Oxidation (M) |
| 239 | 192 – 207 | 773.4062 | 1544.7978 | 1544.8055 | -4.95 | 0 | 54 | 6.4e-006 | 1 | U | K.AAAMNLVLMGLPGAGK.G + 2 Oxidation (M) |
| 240 | 192 – 207 | 773.4063 | 1544.7981 | 1544.8055 | -4.74 | 0 | 27 | 0.003 | 1 | U | K.AAAMNLVLMGLPGAGK.G + 2 Oxidation (M) |
| 241 | 192 – 207 | 773.4066 | 1544.7986 | 1544.8055 | -4.40 | 0 | 14 | 0.066 | 1 | U | K.AAAMNLVLMGLPGAGK.G + 2 Oxidation (M) |
| 242 | 192 – 207 | 773.4068 | 1544.7990 | 1544.8055 | -4.17 | 0 | 17 | 0.034 | 1 | U | K.AAAMNLVLMGLPGAGK.G + 2 Oxidation (M) |
| 243 | 192 – 207 | 773.4071 | 1544.7996 | 1544.8055 | -3.76 | 0 | 47 | 2.9e-005 | 1 | U | K.AAAMNLVLMGLPGAGK.G + 2 Oxidation (M) |
| 244 | 192 – 207 | 773.4071 | 1544.7996 | 1544.8055 | -3.76 | 0 | 36 | 0.0004 | 1 | U | K.AAAMNLVLMGLPGAGK.G + 2 Oxidation (M) |
| 285 | 214 – 230 | 650.6461 | 1948.9164 | 1948.9353 | -9.68 | 0 | 8 | 0.22 | 1 | U | R.IVEDYGIPHISTGDMFR.A |

| Query | Start – End | Observed | Mr(expt) | Mr(calc) | ppm | M | Score | Expect | Rank | U | Peptide |
|---|---|---|---|---|---|---|---|---|---|---|---|
| 286 | 214 – 230 | 650.6480 | 1948.9220 | 1948.9353 | -6.81 | 0 | 4 | 0.74 | 1 | U | R.IVEDYGIPHISTGDMFR.A |
| 287 | 214 – 230 | 650.6492 | 1948.9258 | 1948.9353 | -4.87 | 0 | 12 | 0.1 | 1 | U | R.IVEDYGIPHISTGDMFR.A |
| 288 | 214 – 230 | 650.9795 | 1949.9166 | 1948.9353 | 503 | 0 | 15 | 0.048 | 1 | U | R.IVEDYGIPHISTGDMFR.A |
| 289 | 214 – 230 | 655.9729 | 1964.8969 | 1964.9302 | -17.0 | 0 | 6 | 0.26 | 1 | U | R.IVEDYGIPHISTGDMFR.A + Oxidation (M) |
| 290 | 214 – 230 | 655.9730 | 1964.8971 | 1964.9302 | -16.9 | 0 | 15 | 0.034 | 1 | U | R.IVEDYGIPHISTGDMFR.A + Oxidation (M) |
| 291 | 214 – 230 | 983.4586 | 1964.9027 | 1964.9302 | -14.0 | 0 | 5 | 0.33 | 1 | U | R.IVEDYGIPHISTGDMFR.A + Oxidation (M) |
| 292 | 214 – 230 | 655.9764 | 1964.9074 | 1964.9302 | -11.6 | 0 | 61 | 8.4e-007 | 1 | U | R.IVEDYGIPHISTGDMFR.A + Oxidation (M) |
| 293 | 214 – 230 | 655.9776 | 1964.9109 | 1964.9302 | -9.84 | 0 | 21 | 0.0087 | 1 | U | R.IVEDYGIPHISTGDMFR.A + Oxidation (M) |
| 294 | 214 – 230 | 655.9777 | 1964.9112 | 1964.9302 | -9.66 | 0 | 25 | 0.0029 | 1 | U | R.IVEDYGIPHISTGDMFR.A + Oxidation (M) |
| 296 | 214 – 230 | 655.9782 | 1964.9127 | 1964.9302 | -8.91 | 0 | 46 | 2.7e-005 | 1 | U | R.IVEDYGIPHISTGDMFR.A + Oxidation (M) |
| 297 | 214 – 230 | 655.9782 | 1964.9128 | 1964.9302 | -8.85 | 0 | 50 | 1e-005 | 1 | U | R.IVEDYGIPHISTGDMFR.A + Oxidation (M) |
| 298 | 214 – 230 | 655.9783 | 1964.9132 | 1964.9302 | -8.68 | 0 | 40 | 9.1e-005 | 1 | U | R.IVEDYGIPHISTGDMFR.A + Oxidation (M) |
| 299 | 214 – 230 | 655.9787 | 1964.9142 | 1964.9302 | -8.15 | 0 | 23 | 0.0045 | 1 | U | R.IVEDYGIPHISTGDMFR.A + Oxidation (M) |
| 300 | 214 – 230 | 655.9787 | 1964.9144 | 1964.9302 | -8.06 | 0 | 55 | 3e-006 | 1 | U | R.IVEDYGIPHISTGDMFR.A + Oxidation (M) |
| 301 | 214 – 230 | 983.4646 | 1964.9146 | 1964.9302 | -7.93 | 0 | 6 | 0.27 | 1 | U | R.IVEDYGIPHISTGDMFR.A + Oxidation (M) |
| 302 | 214 – 230 | 983.4647 | 1964.9147 | 1964.9302 | -7.88 | 0 | 22 | 0.0068 | 1 | U | R.IVEDYGIPHISTGDMFR.A + Oxidation (M) |
| 303 | 214 – 230 | 655.9791 | 1964.9156 | 1964.9302 | -7.46 | 0 | 27 | 0.0022 | 1 | U | R.IVEDYGIPHISTGDMFR.A + Oxidation (M) |
| 304 | 214 – 230 | 983.4651 | 1964.9157 | 1964.9302 | -7.40 | 0 | 13 | 0.045 | 1 | U | R.IVEDYGIPHISTGDMFR.A + Oxidation (M) |
| 305 | 214 – 230 | 655.9792 | 1964.9159 | 1964.9302 | -7.29 | 0 | 21 | 0.0088 | 1 | U | R.IVEDYGIPHISTGDMFR.A + Oxidation (M) |
| 306 | 214 – 230 | 655.9794 | 1964.9163 | 1964.9302 | -7.09 | 0 | 54 | 3.8e-006 | 1 | U | R.IVEDYGIPHISTGDMFR.A + Oxidation (M) |
| 307 | 214 – 230 | 655.9796 | 1964.9171 | 1964.9302 | -6.70 | 0 | 31 | 0.00075 | 1 | U | R.IVEDYGIPHISTGDMFR.A + Oxidation (M) |
| 308 | 214 – 230 | 983.4659 | 1964.9172 | 1964.9302 | -6.60 | 0 | 17 | 0.019 | 1 | U | R.IVEDYGIPHISTGDMFR.A + Oxidation (M) |
| 309 | 214 – 230 | 983.4663 | 1964.9180 | 1964.9302 | -6.23 | 0 | 19 | 0.012 | 1 | U | R.IVEDYGIPHISTGDMFR.A + Oxidation (M) |
| 310 | 214 – 230 | 655.9801 | 1964.9183 | 1964.9302 | -6.06 | 0 | 11 | 0.072 | 1 | U | R.IVEDYGIPHISTGDMFR.A + Oxidation (M) |
| 311 | 214 – 230 | 655.9803 | 1964.9192 | 1964.9302 | -5.61 | 0 | 28 | 0.0017 | 1 | U | R.IVEDYGIPHISTGDMFR.A + Oxidation (M) |
| 312 | 214 – 230 | 655.9807 | 1964.9203 | 1964.9302 | -5.05 | 0 | 4 | 0.36 | 1 | U | R.IVEDYGIPHISTGDMFR.A + Oxidation (M) |
| 313 | 214 – 230 | 983.4687 | 1964.9228 | 1964.9302 | -3.78 | 0 | 0 | 0.95 | 1 | U | R.IVEDYGIPHISTGDMFR.A + Oxidation (M) |
| 314 | 214 – 230 | 655.9817 | 1964.9234 | 1964.9302 | -3.49 | 0 | 9 | 0.12 | 1 | U | R.IVEDYGIPHISTGDMFR.A + Oxidation (M) |
| 315 | 214 – 230 | 983.4742 | 1964.9338 | 1964.9302 | 1.83 | 0 | 4 | 0.36 | 1 | U | R.IVEDYGIPHISTGDMFR.A + Oxidation (M) |
| 316 | 214 – 230 | 983.4745 | 1964.9344 | 1964.9302 | 2.15 | 0 | 7 | 0.21 | 1 | U | R.IVEDYGIPHISTGDMFR.A + Oxidation (M) |
| 317 | 214 – 230 | 983.4776 | 1964.9407 | 1964.9302 | 5.34 | 0 | 13 | 0.06 | 1 | U | R.IVEDYGIPHISTGDMFR.A + Oxidation (M) |

Error: try setting browser cache to automatic.

---

```
LOCUS       QNL33346                  48 aa            linear   BCT 04-SEP-2020
DEFINITION  hypothetical protein (plasmid) [Escherichia coli].
ACCESSION   QNL33346
VERSION     QNL33346.1
DBSOURCE    accession MT180430.1
KEYWORDS    .
SOURCE      Escherichia coli
  ORGANISM  Escherichia coli
            Bacteria; Proteobacteria; Gammaproteobacteria; Enterobacterales;
            Enterobacteriaceae; Escherichia.
REFERENCE   1  (residues 1 to 48)
  AUTHORS   Tarabai,H., Wyrsch,E.R., Bitar,I., Djordjevic,S.P. and Dolejska,M.
  TITLE     Comparative analysis of multidrug resistant Escherichia coli ST216
            isolates from silver gulls in Australia
  JOURNAL   Unpublished
REFERENCE   2  (residues 1 to 48)
  AUTHORS   Tarabai,H., Wyrsch,E.R., Bitar,I., Djordjevic,S.P. and Dolejska,M.
  TITLE     Direct Submission
  JOURNAL   Submitted (09-MAR-2020) Department of Biology and Wildlife
            Diseases, University of Veterinary and Pharmaceutical Sciences
            Brno, Palackeho tr. 1946/1, Brno, Brno 61242, Czech Republic
COMMENT     ##Assembly-Data-START##
            Assembly Method       :: SMRT Link v. 8.0
            Assembly Name         :: pCE1681-A
            Coverage              :: 326X
            Sequencing Technology :: PacBio
            ##Assembly-Data-END##
FEATURES             Location/Qualifiers
     source          1..48
                     /organism="Escherichia coli"
```

```
                     /strain="CE1681"
                     /host="Chroicocephalus novaehollandiae"
                     /db_xref="taxon:562"
                     /plasmid="pCE1681-A"
                     /country="Australia"
                     /collection_date="2012"
                     /note="Closed IncHI2-ST3/IncN fusion plasmid;
                     type: ST216"
     Protein         1..48
                     /product="hypothetical protein"
     CDS             1..48
                     /coded_by="MT180430.1:234558..234704"
                     /note="hypothetical protein"
                     /transl_table=11
                     /db_xref="SEED:fig|6666666.506717.peg.303"
```
